# *PTPN11* Mutation Clonal Hierarchy in Acute Myeloid Leukemia

**DOI:** 10.1101/2024.09.18.612239

**Authors:** Sydney Fobare, Chia Sharpe, Kate Quinn, Kinsey Bryant, Linde A. Miles, Robert L. Bowman, Carolyn Cheney, Casie Furby, Marissa Long, Kaytlynn Fyock, Ben Wronowski, James R. Lerma, Allison Mullaney, Krzysztof Mrózek, Deedra Nicolet, Tom Sesterhenn, Megan E. Johnstone, Shesh N. Rai, Chandrashekhar Pasare, Nives Zimmermann, Andrew J. Carroll, Richard M. Stone, Eunice S. Wang, Jonathan E. Kolitz, Bayard L. Powell, John P. Perentesis, Ann-Kathrin Eisfeld, Erin Hertlein, John C. Byrd

**Affiliations:** Medical Scientist Training Program, The Ohio State University, Columbus, OH; Department of Internal Medicine, University of Cincinnati, Cincinnati, OH; Division of Hematology, Department of Internal Medicine, The Ohio State University Comprehensive Cancer Center, Columbus, OH; Division of Experimental Hematology & Cancer Biology, Cincinnati Children’s Hospital, Cincinnati, OH; Department of Cancer Biology, University of Pennsylvania, Philadelphia, PA; Clara D. Bloomfield Center for Leukemia Outcomes Research, Columbus, The Ohio State University Comprehensive Cancer Center, Columbus, OH; Alliance Statistics and Data Management Center, The Ohio State University Comprehensive Cancer Center, Columbus, OH; Department of Genetics, University of Alabama at Birmingham, Birmingham, AL; Dana-Farber/Partners CancerCare, Boston, MA; Roswell Park Comprehensive Cancer Center, Buffalo, NY; Monter Cancer Center, Zucker School of Medicine at Hofstra/Northwell, Lake Success, NY; Wake Forest Baptist Comprehensive Cancer Center, Winston-Salem, NC; Cancer and Blood Diseases Institute, Cincinnati Children’s Hospital, Cincinnati, OH; Environmental & Public Health Sciences, University of Cincinnati, Cincinnati, OH; Division of Immunobiology, Cincinnati Children’s Hospital Medical Center, Cincinnati, OH; Pathology & Laboratory Medicine, University of Cincinnati, Cincinnati, OH

**Author notes:** Correspondence: John C. Byrd, MD, Chair, Department of Internal Medicine, University of Cincinnati College of Medicine, 231 Albert Sabin Way, ML 0551, Room 6065, Cincinnati OH 45267-0551; Erin Hertlein PhD, Assistant Professor, University of Cincinnati College of Medicine, 231 Albert Sabin Way, ML 0551, Room 7009, Cincinnati OH 45267-0551. Equal contribution as first author (Sydney Fobare, Chia Sharpe). Equal contribution as senior author (Erin Hertlein, John C. Byrd). Presented in part in abstract form at the 64^th^ annual meeting of the American Society of Hematology, 10-13 December 2022.

**Keywords:** Acute myeloid leukemia, *PTPN11* mutations, *NPM1* mutations, clonal evolution, immunophenotyping, single-cell DNA sequencing, plasmacytoid dendritic cell

## Abstract

Mutations in protein tyrosine phosphatase non-receptor type 11 (*PTPN11*) have been considered late acquired mutations in acute myeloid leukemia (AML) development. To interrogate the ontogeny of *PTPN11* mutations, we utilized single-cell DNA sequencing and identified that *PTPN11* mutations can occur as initiating events in some AML patients when accompanied by strong oncogenic drivers, commonly *NPM1* mutations. The co-driver role of *PTPN11* mutations was confirmed in a novel murine model that exhibits an AML phenotype with early expansion of a diverse set of variably differentiated myeloid cells that engrafted into immunodeficient and immunocompetent mice. This immune diversity was reconstituted from early precursor cells when engrafted into immunodeficient mice. Moreover, immune diversity was also observed in the blast component of patient samples with *NPM1* and *PTPN11* mutations, providing novel antigen targets for immune based approaches in this subset of AML that is resistant to multiple targeted therapies.

## Introduction

Acute myeloid leukemia (AML) is the most common acute leukemia in adults and for most patients it has extremely poor outcomes. In younger adult patients, leukemogenesis most commonly involves early acquisition of a distinct balanced translocation or mutations with strong oncogenic potential, followed by acquisition of additional late genetic mutations in signaling genes (e.g., *KIT*, *FLT3*, *NRAS*, *KRAS*, *NF1*, *CBL*, and *PTPN11*)^1^. In contrast, in most older adults, AML is characterized by the early acquisition of a variety of clonal hematopoiesis (CH) mutations (e.g., *DNMT3A, TET2, ASXL1*) that are not by themselves transforming but rather confer a stem cell/progenitor growth advantage, promote inflammation, and increase risk of hematologic malignancy and multiple other complications^2^. CH mutations are then followed by acquisition of additional signaling mutations or cytogenetic abnormalities that lead to the spectrum of neoplastic development: cytopenias without dysplasia, myelodysplastic syndrome (MDS), and/or ultimately AML^3^. Characterization of the impact of gene mutations and cytogenetic abnormalities on clinical outcome in AML has been mostly derived from treatment trials with intensive chemotherapy, and as a result a distinct group of driver and late mutations are well characterized as predictors of poor outcome in the 2022 European LeukemiaNet (ELN) classification of AML^4^.

The development of now approved targeted agents toward FLT3 (midostaurin, gilteritinib, quizartinib), IDH1 (ivosidenib, olutasidenib), IDH2 (enasidenib), and BCL2 (venetoclax) have transformed the treatment landscape in AML. Within these new targeted therapeutic approaches, many established mutations associated with poor prognosis, such as *TP53* mutations, have continued to predict poor outcomes. However, additional mutations such as MAPK pathway activating mutations (e.g., *NRAS*, *KRAS*, *CBL, NF1,* and *PTPN11*) have emerged as specifically impacting the outcomes of patients receiving targeted therapies^5–8^. Indeed, patients with *PTPN11*- mutated AML had the lowest response among patients carrying other genomic abnormalities who received IDH2 inhibitors^9^ and venetoclax/azacitidine^10,11^.

The *PTPN11* gene encodes Src homology region 2 (SH2)-containing protein tyrosine phosphatase 2 (SHP2). SHP2 contains two SH2 domains and a tyrosine phosphatase domain. SHP2 is capable of self-regulation as the N-terminal SH2 domain can bind to the active phosphatase domain to prevent dephosphorylation of SHP2 targets^12^. Pathogenic *PTPN11* mutations tend to occur in regions involved in its self-regulation, and thus, these gain-of-function mutations result in activation of numerous pathways including RAS/ERK1/2, FLT3, JAK/STAT, PI3K/AKT, and NF-kB, mediates immune evasion via PD-1, and overexpression of antiapoptotic proteins such as MCL-1^13–15^. In humans, gain of function *PTPN11* mutations can occur as germline mutations in Noonan syndrome and are associated with a higher frequency of juvenile myelomonocytic leukemia (JMML) suggesting this mutation can contribute to the initiation of myeloid malignancies^16,17^.

These findings have prompted several groups, including our own, to examine the prognostic impact of *PTPN11* mutations in AML. These analyses suggest that *PTPN11* mutations co-occur most commonly with *NPM1* mutations or broadly with chromosome translocations and typically confer poor outcome^18–20^. In the absence of an *NPM1* mutation, *PTPN11* mutations are often associated with cytogenetic abnormalities associated with dysregulation of *EVI1.* However, there are some inconsistencies between studies suggesting the pathogenic contribution of *PTPN11* mutations may depend on its context amongst other driver mutations, such as *NPM1*^18^. Notably, in some patients, *PTPN11* mutations occur at a high variant allele frequency (VAF) suggesting they could be initiating or co-initiating events in leukemogenesis. Previous reports have suggested that dominant *PTPN11* mutations are associated with better outcomes than subclonal mutations in patients treated with intensive chemotherapy^19^. Either an early driver or an acquired subclonal mutation, *PTPN11* mutations have serious implication for predicting therapeutic response and emergence of therapeutic resistance.

To study the role of *PTPN11* gain of function mutations, Xu et al. developed the *Ptpn11^E76K^* mouse that mimics the most common hotspot mutation in both Noonan syndrome and myeloid malignancies^21^. They found that constitutive expression of *Ptpn11^E76K^* is embryonically lethal in mice, but conditional, heterozygous expression of the *Ptpn11^E76K^* mutation under the control of the Mx-Cre, promotes the development of multi-lineage leukemias. Interestingly, the *Ptpn11*^E76K^ mice had decreased Lineage-Sca-1^+^c-Kit^+^ (LSK) cells in the bone marrow with evidence of accelerated activation and differentiation of progenitor cells, including quiescent long-term hematopoietic stem cells (HSCs). *Ptpn11^E76K^* HSCs also have greatly decreased repopulating capabilities in competitive repopulation assays.

Given the lack of knowledge related to clonal evolution of *PTPN11* mutations and the absence of a consistent *Ptpn11* mutant AML mouse model, we initiated both human and murine studies to address these questions. Herein, we demonstrate *PTPN11* can be an early acquired mutation, commonly co-occurring with a strong oncogene (*NPM1* mutation) to promote AML development. The *NPM1* and *PTPN11* mutated subtype of AML has a diverse immune profile in patients that was reproduced in our *Npm1^cA^/Ptpn11^E76K^* animal model. This model provides a novel system to understand the role of early *PTPN11* mutations as it relates to clonal architecture and influence on AML immunophenotype.

## Methods

### Patient samples

Single-cell DNA sequencing (scDNA-seq) was performed on samples from 16 newly diagnosed patients *with de novo* AML with *PTPN11* mutations. Five of these patients had *PTPN11* mutation in the absence of *NPM1* mutation with four patients having a *PTPN11* mutation in the dominant clone (VAF>0.3). Nine patients had *PTPN11* as a dominant mutation (VAF >0.3) and two had *PTPN11* as a subclonal mutation (VAF <0.3) together with *NPM1*. All patients provided written informed consent to participate in the following protocols for the collection of pretreatment peripheral blood and bone marrow samples: CALGB 8461 (cytogenetic studies), CALGB 9665 (leukemia tissue bank), and CALGB 20202 (molecular studies).

### Sequencing Reagents

TotalSeq-D Heme Oncology Panel of oligo-conjugated reagents were purchased from Biolegend (clone information can be found in Supplemental Methods). Primers for the Myeloid Clonal Evolution Panel were purchased from MissionBio.^22^

### Single-cell DNA and protein library preparation and sequencing

Cryopreserved samples were thawed and counted using the Cellaca PLX Image Cytometry System (Nexcelom). A Dead Cell Removal Kit (Miltenyi Biotec) was then used to collect live cells. Cells were recounted and resuspended in Cell Staining Buffer (MissionBio). The remaining staining, cell encapsulation, barcoding, target PCR amplification, PCR cleanup, and quantification steps were performed according to MissionBio’s Tapestri Single-Cell DNA + Protein Sequencing Protocol. Libraries were quantified using a Tapestation (Agilent) prior to pooling. Four pooled samples were run on one S1 lane on an Illumina NovaSeq at the Cincinnati Children’s Hospital Medical Center DNA Sequencing and Genotyping Core.

### Data processing

FASTQ files were analyzed through the Tapestri Pipeline, and the subsequent output was analyzed in R as previously described.^22^ Briefly, reads were aligned to the hg19 genome build^23,24^ and to cell barcodes. Genotypes were called using the GATKv3.7.^25–27^ Any intronic changes were removed from the analysis and only non-synonymous variants were included. The aligned reads of all called variants were visually inspected in the Integrative Genomics Viewer (Broad Institute). All variants with VAFs <0.01 and genotyped in <50% of cells were excluded from the analysis. Clones were manually examined to exclude those that have high allele dropout rates, low genotype quality, or low read depth, and exhibit loss of heterozygosity or improper amplification of mutant alleles. Further filtering and mutational order were determined by current biological knowledge of clonal evolution. For the protein analysis, normalization was performed using DSB normalization.^28^ Dimension reduction, differential expression markers, and data visualization was performed using Seurat in R as previously described.^22,29^ To generate unsupervised heatmaps of protein expression in R, the ComplexHeatmap package from Bioconductor was used.^30,31^

### Experimental animals

*Ptpn11*^E76K^ and *Npm1*^cA^ mice were generous gifts from Dr. Cheng-Kui Qu^21^ at Emory University and Dr. George Vassiliou^32^ from Wellcome Sanger Institute, respectively. B6.Cg-Tg(Mx1-cre)1Cgn/J mice (referred to as Mx-Cre mice; strain# 003556)^33^ and B6.SJL- *Ptprc^a^ Pepc^b^*/BoyJ (referred to as PepBoy mice; strain# 002014) were purchased from The Jackson Laboratory. NOD-Prkdc^em26Cd52^Il2rg^em26Cd22^/NjuCrl (referred to as NCG mice; strain# 572) were purchased from Charles River Laboratories. All animal experiments were conducted after approval by the University of Cincinnati (UC) and The Ohio State University (OSU) Institutional Animal Care and Use Committees (IACUC). To induce expression of the transgenes, mice were injected five times (once every other day) with 20µg/g of polyinosinic-polycytidylic acid [poly(I:C), Sigma Aldrich] via intraperitoneal injection. Mice were bled biweekly to monitor disease development via complete blood counts (CBCs) obtained on the Element HT5 (Heska) and signs of early removal criteria (ERC). ERC was defined as 20% weight loss, partial or full hindlimb paralysis, dehydration, anorexia, anemia, hunched posture, inactivity, lethargy, difficulty breathing, or rough hair coat.

### Histopathology

Samples were preserved in 10% neutral buffered formalin and paraffin embedded. They were then sectioned at 3µm onto glass slides and stained with hematoxylin and eosin. A veterinary pathology resident performed the subsequent tissue analysis with faculty veterinary supervision.

### Murine cell isolation

When a mouse met ERC, spleen, femurs, and tibias were collected in phosphate buffered saline (PBS) and immediately processed. Spleens were dissociated in the gentleMACS Dissociator (Miltenyi Biotec) and bone marrow cells were isolated via centrifugation. Both splenocytes and bone marrow cells underwent red blood cell (RBC) lysis with 1x RBC Lysis buffer (Thermo Fisher Scientific). Samples were then sterilely cryopreserved.

### Peripheral blood, splenocytes, and bone marrow flow cytometry

For the peripheral blood flow cytometry, 20 μL of blood was blocked with TrueStain FcX (anti-mouse CD16/32) (BioLegend) and monocyte block for 5 minutes prior to staining with the 20-antibody cocktail (Supplemental Methods) in Brilliant Stain Buffer Plus (BD Biosciences) and FACS buffer (PBS, 2% FBS, and 0.1% sodium azide). After 30 minutes of incubation, samples were lysed twice with 1x RBC Lysis Buffer (ThermoFisher Scientific). Cells were then fixed in paraformaldehyde for 20 minutes prior to resuspension in FACS buffer. 50,000 events per sample were collected on the Cytek Aurora (Cytek Biosciences).

For the splenocyte and bone marrow immunophenotyping flow cytometry, cryopreserved splenocytes and bone marrow cells were thawed and 1×10^6^ or 10×10^6^ cells, respectively, were obtained for staining. Cells were first stained with Live/Dead Fixable Blue Dead Cell Stain Kit (ThermoFisher Scientific) for 30 minutes. Cells were then washed and blocked with TrueStain FcX (anti-mouse CD16/32) (BioLegend) and monocyte block for 5 minutes. Antibody master mix for the splenocyte or bone marrow panel (Supplemental Methods) with Brilliant Stain Buffer Plus (BD Biosciences) in FACS buffer was then added and cells were incubated for 30 minutes. Cells were then washed 3 times prior to paraformaldehyde fixation for 20 minutes. Cells were then washed and stored at 4°C overnight. Flow cytometry data was acquired the next day on the Cytek Aurora (Cytek Biosciences) with 100,000 events for splenocytes and 1,000,000 events for the bone marrow cells were collected per sample.

### *In vivo* transplantation

One million bulk splenocytes from *Ptpn11*^E76K^/*Npm1*^cA^ mice spleens were injected into NCG or PepBoy mice via tail vein injection. For the LSK engraftment, the EasySep Mouse Hematopoietic Progenitor Cell Isolation Kist (StemCell) was used to enrich for HSCs from the splenocytes. Enriched HSCs were then sorted on the BigFoot Spectral Cell Sorter (Thermo Fisher), and LSKs were collected for engraftment (gating strategy, Supplemental Fig 3). 10,000-50,000 LSKs were engrafted per NCG mouse via tail vein injection. Disease development was monitored through weekly bleeds where we obtained complete blood counts and performed flow cytometry for CD45.2 (clone 104) and CD45.1 (clone A20). At ERC, splenocytes and bone marrow cells were cryopreserved as previously described.

### Statistical Analysis

To analyze differences in overall survival time we fit a Cox proportional-hazards model for Group and use contrast to compare each pair of groups. Due to all subjects died, we also fit ANOVA model for the logarithm of the overall survival time about Group and use contrast to compare each pair of groups. ANOVA for log(Weight) or log(Count) using contrast for each pair of groups was used to determine differences in spleen weight, white blood cells (WBCs), RBCs, and platelets (PLTs) at ERC. ANOVA for log(Percent) was used to compare the frequency of various cell populations from the broad splenocyte and bone marrow immunophenotyping panels. Two-sided t-tests and the Shapiro-Wilk method was used for normality. For multiple-comparisons, the Benjamin and Hochberg method was used to adjust p-values. Full statistical analysis can be found in supplemental data. For Tapestri analysis, differences in expression levels of cell surface markers were calculated based on the Wilcoxon rank sum test using Seurat in R. A *P*-value <0.05 was used to determine significance.

## Results

### PTPN11-mutant AML demonstrates multiple ancestor origins

Previous work from our group suggested that *PTPN11* mutations occur at a wide range of VAFs up to 0.54, suggesting that *PTPN11* mutations may occur early in leukemogenesis^18^. To further elucidate the ontogeny of dominant *PTPN11* mutations, we assembled a cohort of patients enriched with those that had a VAF >0.3 (3 patients had a VAF <0.3) and mutations that fell within the *PTPN11* amplicons generated by the Tapestri scDNA-seq Myeloid Clonal Evolution Published Panel (Mission Bio). Five patients with confirmed *PTPN11* mutations without a co-occurring *NPM1* mutation by bulk sequencing and 11 with concurrent *PTPN11* and *NPM1* mutations were analyzed. The demographics and mutation profile identified by bulk sequencing of the included patients are summarized in Table 1. All 5 patients without *NPM1* mutations had a strong driver translocation or inversion involving *EVI1* together with monosomy 7 (cases AML-1 to AML-4) or t(6;11)(q27;q23) (case AML-5). However, chromosomal alterations are not captured by this scDNA-seq panel, so in these samples a single *PTPN11* mutant clone was identified (representative case AML-3) (Fig. 1A; Extended Data Fig 1A).

**Figure 1.**
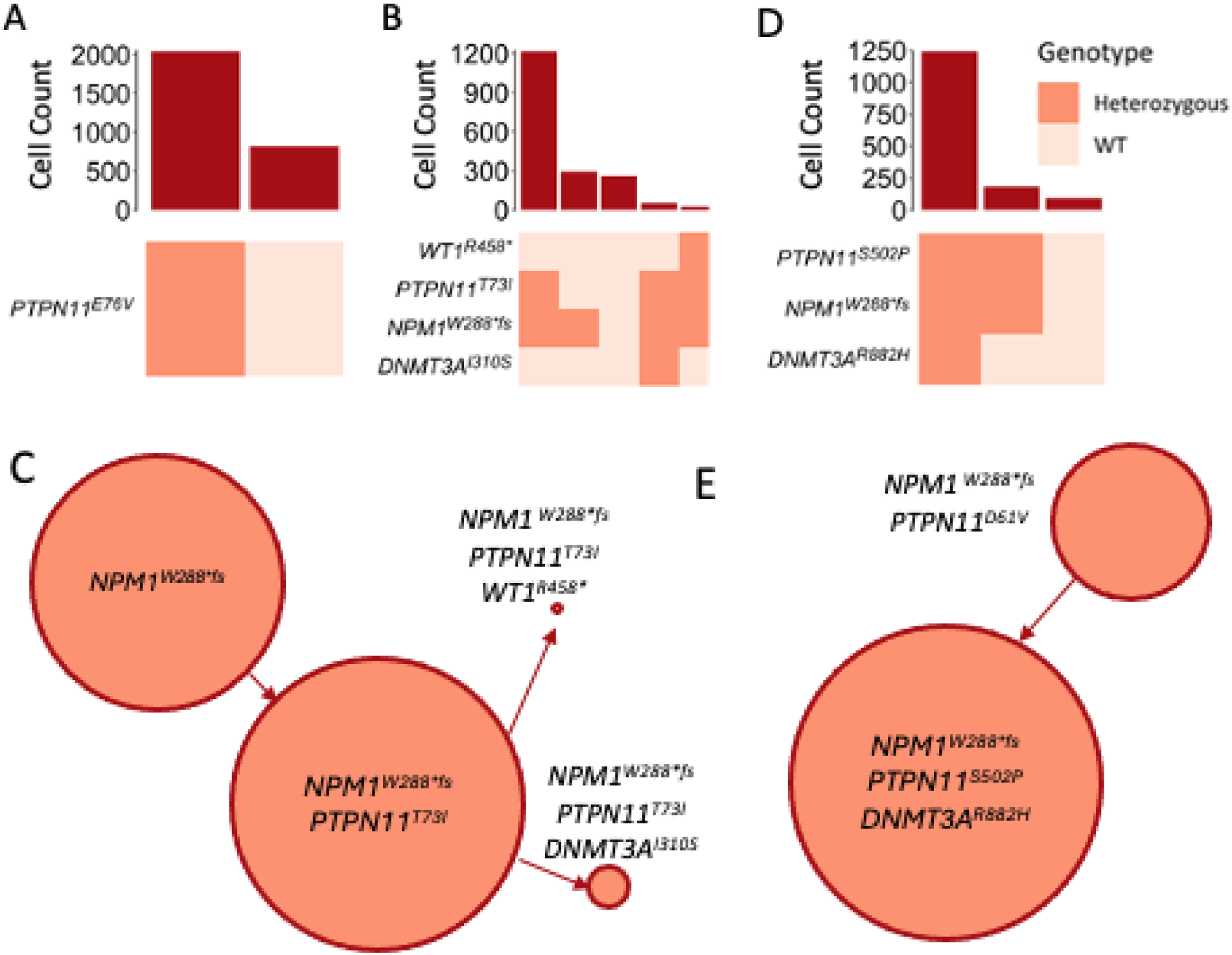
Clonal architecture of PTPN11 mutant AML patient samples. Clonograph demonstrating frequency of co-occurring mutations determined using scDNA-seq representative samples: **A.** *PTPN11* mutation without *NPM1* mutation (AML-3). **B.** Subclonal *PTPN11* mutation (AML-16). **C.** Inferred clonal relationship for **B. D.** Early *PTPN11* mutation (AML-9). **E.** Inferred clonal relationships for **D.**

**Table 1.**
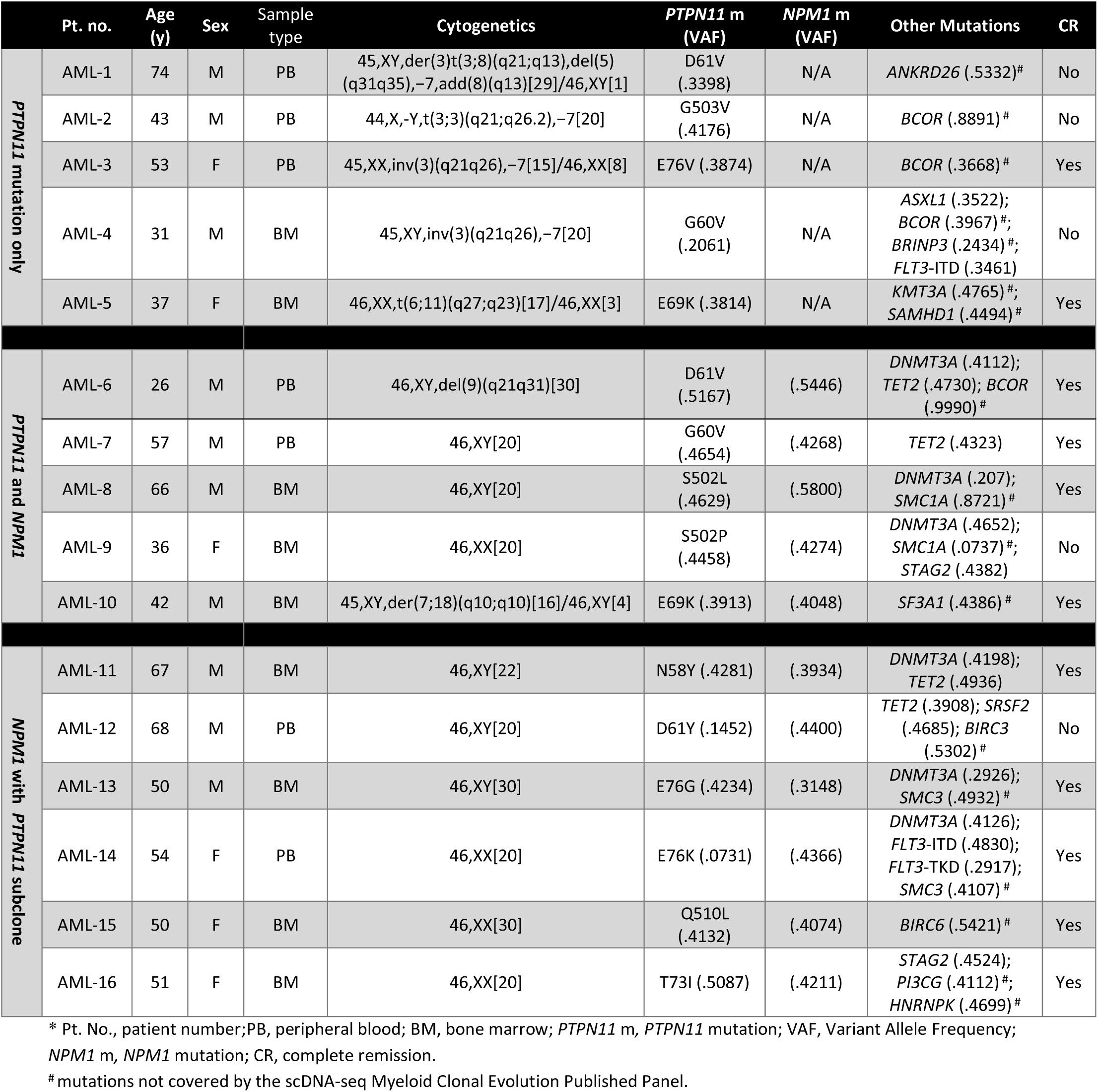
Patients Undergoing Single-Cell Sequencing.

Among the 11 patients with both *PTPN11* and *NPM1* mutations, two distinct patterns emerged. Six patient samples showed evidence of an *NPM1* mutant clone that lacked a *PTPN11* mutation (representative case AML-16; Fig 1B; Extended Data Fig 1B), suggesting that the *NPM1* mutation preceded the *PTPN11* mutation. However, the double mutant cells generated the dominant clone (Fig 1C). The clonal architecture in these samples aligns with the predominant understanding of *PTPN11* mutations as a late acquired signaling mutation akin to *FLT3* or *RAS* family mutations and is supported previous reports of scDNA-seq performed on patients with AML^22,34^. However, in the other 5 patients, we observed *PTPN11* mutations in the earliest detectable clone (representative case AML-9) (Fig 1D; Extended Data Fig 1C). In 4 of these 5 patients, no single *PTPN11* mutant clone was identified, which precludes the ability to confirm that *PTPN11* was an initiating mutation. Nevertheless, this data suggests the early emergence of a *PTPN11* mutation in the clonal evolution of these patients (Fig 1E). Indeed, in 1 patient, we identified 3 mutant clones: *PTPN11* single mutant, *PTPN11* and *DNMT3A* double mutant clone, and a *PTPN11*, *DNMT3A* and *NPM1* triple mutant clone. This suggests that in some patients *PTPN11* mutations may be an early event in AML leukemogenesis and can occur prior to the acquisition of other mutations (AML-8; Extended Data Fig 1C). Collectively, this data supports the conclusion that while *PTPN11* mutations are frequently a late acquired mutation, in contrast to other *RAS* family mutations, they can serve as a driver or co-driver mutation in AML when coupled with mutations or chromosomal abnormalities that are strongly associated with myeloid oncogenesis.

### Npm1^cA^/Ptpn11^E76K^ mice consistently develop acute myeloid leukemia

Our AML patient data suggests that *PTPN11* mutations can constitute a driving transformative event. We therefore aimed to develop a *PTPN11* mutant AML mouse model that would allow further investigation of the role of *PTPN11* mutations in leukemogenesis and pathogenesis. Additionally, such a mouse model provides an important tool for studying response to therapies targeting this mutation, an area of clinically unmet need for AML. We crossed *Ptpn11*^E76K^ ^21^, *Npm1*^cA32^ and Mx-Cre^33^ strains, and all alleles were maintained heterozygous in experimental animals. Mice expressing both mutant alleles (*Npm1*^cA^/*Ptpn11*^E76K^) exhibited a fully penetrant lethal disease phenotype with a median overall survival of 13.4 weeks post poly(I:C) induction, significantly faster than either *Ptpn11*^E76K^/Mx-Cre (*Ptpn11*^E76K^; 39.2 weeks, (FDR)*P*=0.0001) or *Npm1*^cA^/Mx-Cre (*Npm1*^cA^; 35 weeks, (FDR)*P*=0.0001) mice (Fig 2A).

At time of sacrifice, *Npm1*^cA^/*Ptpn11*^E76K^ mice had significant splenomegaly compared to age matched Mx-Cre ((FDR)*P* ≤0.0001) and single gene control mice (Fig 2B; *Npm1*^cA^ (FDR)*P* ≤0.0001; *Ptpn11*^E76K^ (FDR)*P* ≤0.0001). *Npm1*^cA^/*Ptpn11*^E76K^ mice also exhibited significant leukocytosis, anemia, and thrombocytopenia compared with age-matched single gene controls (Fig 2C). Notably, *Npm1*^cA^ mice also showed decreased red cell and platelet counts, consistent with previous reports.^32^ As previously published, the *Ptpn11*^E76K^ mouse can develop multi-lineage leukemias; therefore we wanted to confirm that the addition of an *Npm1*^cA^ mutation would result in a consistent myeloid phenotype. We analyzed blood smears and performed flow cytometric analysis showing that *Npm1*^cA^/*Ptpn11*^E76K^ mice have a clear expansion of CD11b^+^/c-Kit^+^ cells in the blood as early as 4 weeks post-induction which progressively expands throughout disease development (Fig 2D, Extended Data Fig 2A-B). Finally, histopathological analysis of *Npm1*^cA^/*Ptpn11*^E76K^ mice (n=5) with lethal disease showed an infiltration of immature myeloid cells in multiple organs suggesting extramedullary hematopoiesis, another common phenotype associated with AML (Fig 2E; Extended Data Fig 2C).

**Figure 2.**
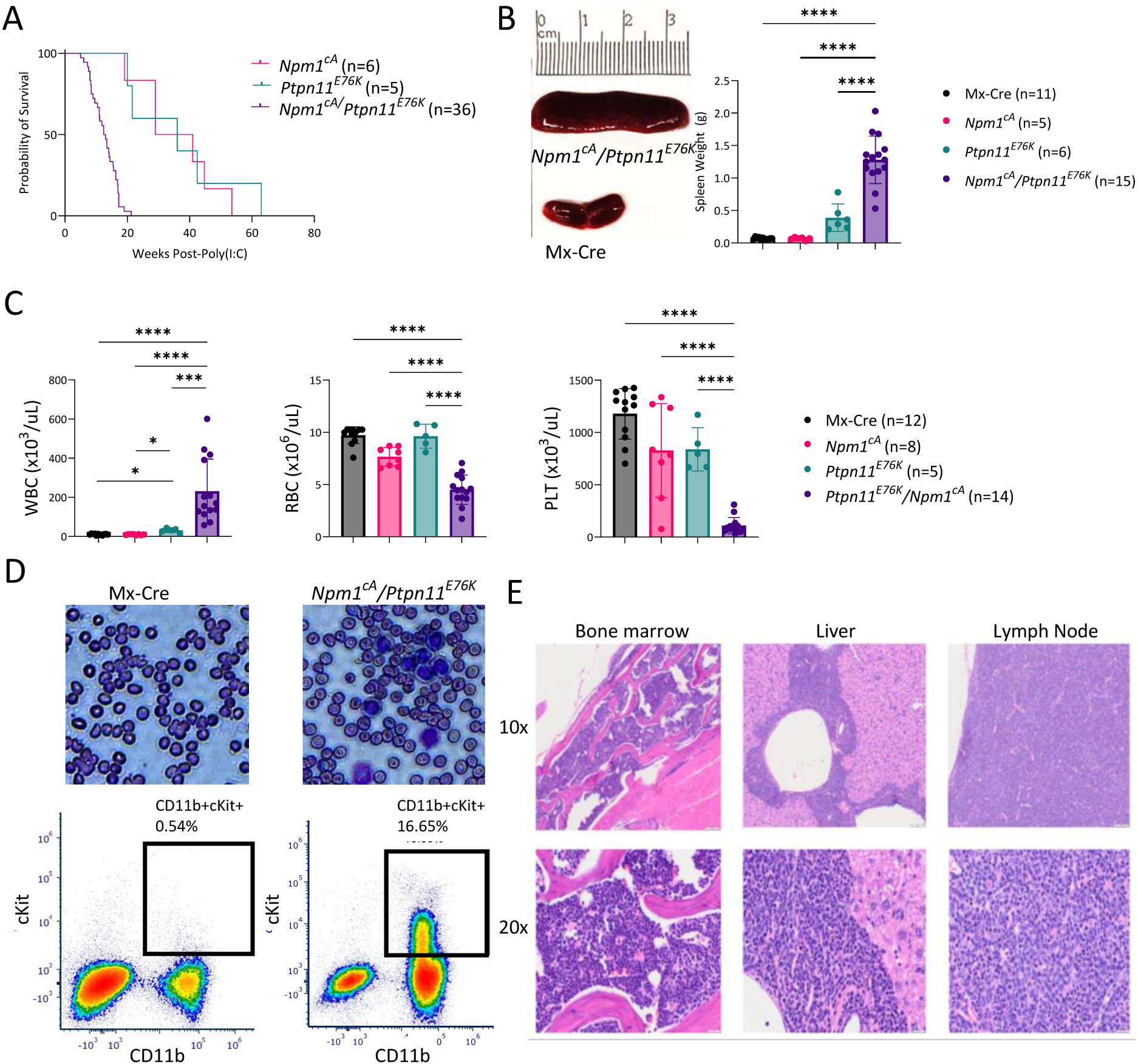
Characterization of a novel Npm1^cA^/Ptpn11^E76K^ mouse model. **A.** Overall survival of *Npm1*^cA^, *Ptpn11*^E76K^, and *Npm1*^cA^/*Ptpn11*^E76K^ mice. **B.** Representative picture of spleen collected from a *Npm1*^cA^/*Ptpn11*^E76K^ mouse at survival endpoint compared to age matched Mx-Cre mouse spleen. Summary of spleen weights in grams of *Npm1*^cA^/*Ptpn11*^E76K^ mice at time of death compared to aged-matched controls. **C.** White blood cell (WBC), red blood cell (RBC), and platelet (PLT) counts of *Npm1*^cA^/*Ptpn11*^E76K^ mice at time of death compared to aged-matched controls. **D.** Representative peripheral blood smears and flow cytometry of plot (gated of CD45^+^ cells) of Mx-Cre and *Ptpn11*^E76K^/*Npm1*^cA^ mice 4 weeks post poly(I:C) induction. **E.** Representative histopathology images of the bone marrow, liver, and lymph node of an *Npm1*^cA^/*Ptpn11*^E76K^ mouse with overt leukemia (n=6). *(FDR)*P*≤0.05, **(FDR)*P*≤0.01, ***(FDR)*P*≤0.001, ****(FDR)*P*≤0.0001.

### Broad immunophenotyping identifies diverse myeloid populations in the Npm1^cA^/Ptpn11^E76K^ mice

While we noted a significant increase in CD11b^+^c-Kit^+^ cells in the peripheral blood that indicated the expansion of an immature myeloid cells consistent with myeloid leukemia, we wanted to comprehensively characterize the *Npm1*^cA^/*Ptpn11*^E76K^ murine disease immunophenotype. To this end, we developed a 36-specificity spectral flow cytometry immunophenotyping panel to characterize both the lymphoid and myeloid populations in the spleen (gating strategy: Supplemental Fig 1A). Compared with age-matched single gene controls, the *Npm1*^cA^/*Ptpn11*^E76K^ mice exhibited an expansion of multiple distinct myeloid populations, including neutrophils (CD11b^+^Ly6G^+^) and monocytic cells (CD11b^+^Ly6C^+^) (Fig 3A, Extended Data Fig 3A). As *PTPN11*-mutated AML is associated with a monocytic phenotype,^20^ it was unsurprising to find that the *Npm1*^cA^/*Ptpn11*^E76K^ mice have an increase in the percentage of monocytes compared with control mice. However, in addition to the expected expansion of the CX3CR1^-^c-Kit^+^ population of ‘immature’ monocytic cells, we also observed a significant expansion of the CX3CR1^+^c-Kit^-^ population of ‘mature’ monocytes (Fig 3B). This suggests that while *Npm1* mutations have been associated with differentiation block, in the context of a co-occurring *Ptpn11* mutation, there is an expansion of variably differentiated myeloid cells including those immunophenotypically consistent with mature monocytes.

**Figure 3.**
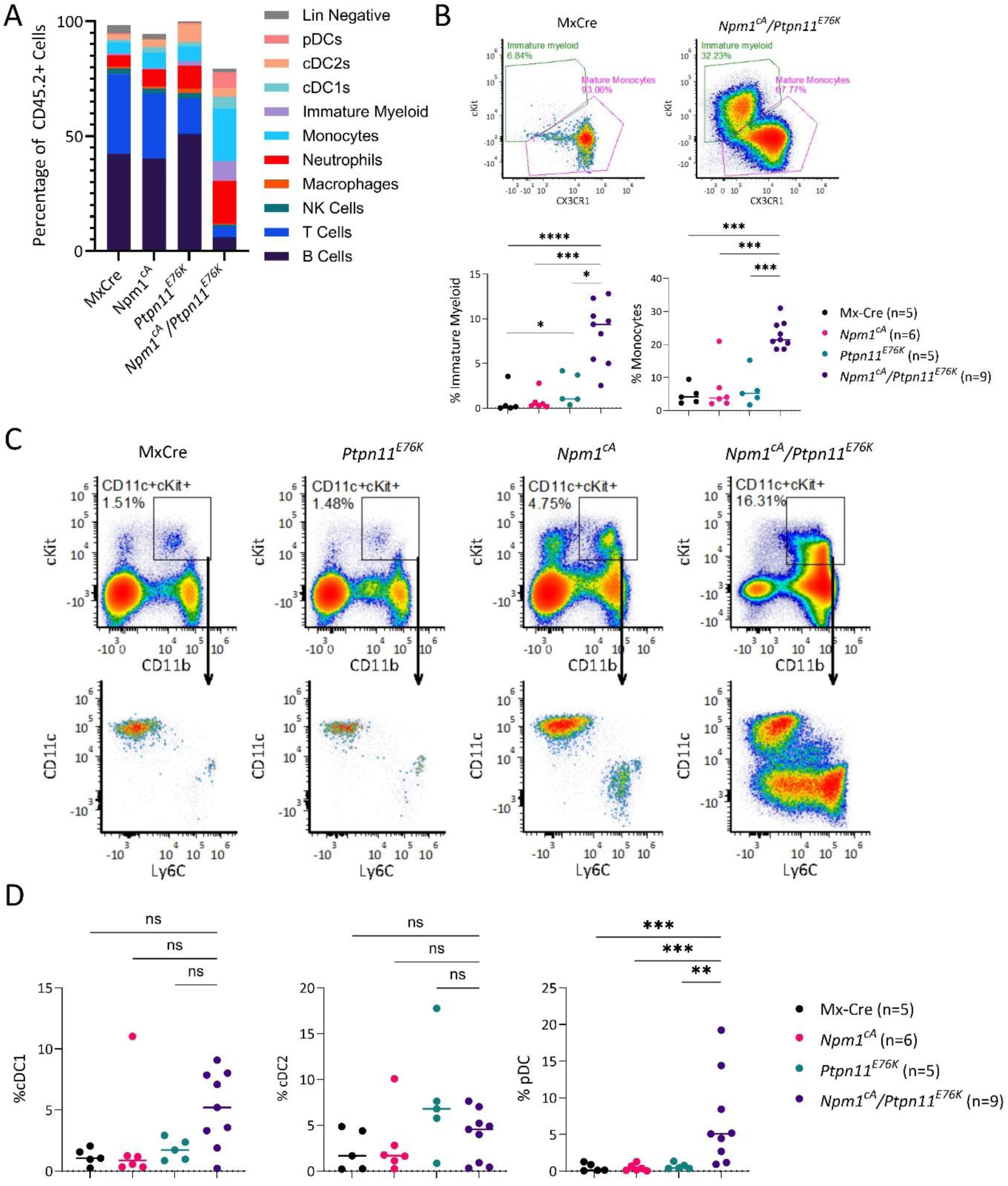
Immunophenotyping of Npm1^cA^/Ptpn11^E76K^ splenocytes. Flow cytometry analysis of splenocytes from Mx-Cre (n=5), *Npm1*^cA^ (n=5), *Ptpn11*^E76K^ (n=5), and *Npm1*^cA^/*Ptpn11*^E76K^ (n=9) mice. Percentages were calculated as fraction of total CD45.2^+^ cells and all control mice were aged-matched. **A.** Percentage of immune subsets in the spleen. **B.** Representative flow plots CX3CR1 and c-Kit expression on monocytes in a representative *Npm1*^cA^/*Ptpn11*^E76K^ mouse. Percentage immature and mature myeloid cells. **C.** Representative flow plots showing proportion of CD11b^+^c-Kit^+^ cells, and monocytic (Ly6C^+^), and dendritic (CD11c^+^) subsets therein. **D.** Percentages of cDC1, cDC2 and pDC populations. *(FDR)*P*≤0.05, **(FDR)*P*≤0.01, ***(FDR)*P*≤0.001, ****(FDR)*P*≤0.0001. Abbreviations: Lineage Negative (Lin Negative), Plasmacytoid Dendritic Cells (pDCs), Conventional Dendritic Cells 2 (cDC2s), Conventional Dendritic Cells 1 (cDC1s), and Natural Killer Cells (NK Cells).

Expansion of c-Kit^+^ or CD11b^+^c-Kit^+^ cells has been used to define a leukemia blast-like phenotype in other AML mouse models^35–38^. To our surprise, the c-Kit^+^ ‘immature’ monocytes composed less than half of the total CD11b^+^c-Kit^+^ cells present in the spleen, with the remaining cells being CD11c^+^ dendritic cells (Fig 3C). While conventional dendritic cells (cDCs) normally express c-Kit^39^, there was no significant expansion of either cDC1 (CD11c^+^CD8a^+^CD24^+^) or cDC2 (CD11c^+^CD11b^+^CD172a^+^) cells in the *Npm1*^cA^/*Ptpn11*^E76K^ mice (Fig 3D). However, there was a significant expansion of plasmacytoid dendritic cells (pDCs) (CD11c^+^B220^+^) in the *Npm1*^cA^/*Ptpn11*^E76K^ mice compared with controls (Fig 3D). The expansion of pDCs was observed in the peripheral blood concurrent with the expansion of other CD11b+cKit+ cells (Extended Data Fig 3B). In the *Npm1*^cA^/*Ptpn11*^E76K^ mice, a subpopulation of pDC phenotype cells also expressed CD11b and c-Kit, which isn’t typical for a normal pDC population. We therefore validated that the CD11c^+^B220^+^ cells also expressed additional markers associated with mature pDCs. Within this CD11c^+^B220^+^ population, there was a subpopulation that expressed both mature pDCs markers Siglec-H and CD317 (Extended Data Fig 3C). Notably, while *Npm1*^cA^ and *Ptpn11*^E76K^ single mutant mice showed an expansion of myeloid populations at survival endpoint, there was no significant expansion of pDCs suggesting this population may be specifically associated with the *Npm1*^cA^/ *Ptpn11*^E76K^ double mutant AML model (Extended Data Fig 3D).

As AML is primarily a disease of the bone marrow, we also developed a 30-specificity bone marrow immunophenotyping panel that both allowed for the characterization of the mature, stem, and progenitor populations (gating strategy: Supplemental Fig 2). Like the spleen, within the lineage positive cells there was an expansion of both immature and mature monocytes as well as a trend toward an expansion of pDCs in the bone marrow compared to age matched controls. However, notable was a significant reduction in the frequency of neutrophils in *Npm1*^cA^/*Ptpn11*^E76K^ mice compared to both Mx-Cre and *Ptpn11*^E76K^ mice (Fig 4A; Extended Data Fig 4A).

**Figure 4.**
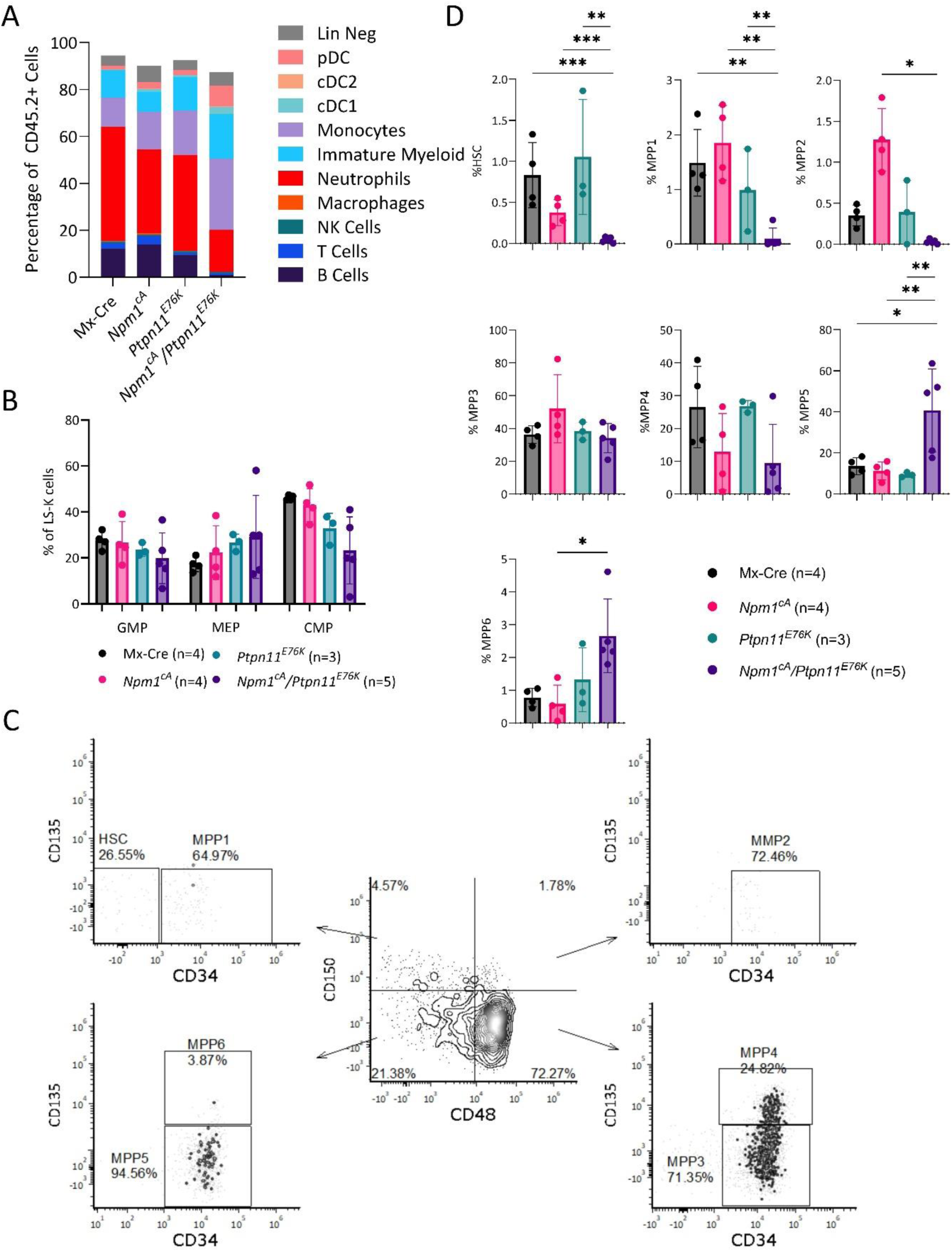
Immunophenotyping of Npm1^cA^/Ptpn11^E76K^ bone marrow. **A.** Percentage of immune subset in the bone marrow of Mx-Cre (n=4), *Npm1*^cA^ (n=4), *Ptpn11*^E76K^ (n=3), and *Npm1*^cA^/*Ptpn11*^E76K^ (n=5) mice. Percentages were calculated based on CD45.2 positivity and all control mice were aged-matched. **B.** Percentage of GMP, MEP and CMP calculated as a percentage of total LSK cells. **C.** Representation of HSC and MPP gating strategy. **D.** HSC and MPPs as a percentage of LSK cells \**P*≤0.05, **(FDR)*P*≤0.01, ***(FDR)*P*≤0.001. Abbreviations: Lineage Negative (Lin Negative), Plasmacytoid Dendritic Cells (pDCs), Conventional Dendritic Cells 2 (cDC2s), Conventional Dendritic Cells 1 (cDC1s), Natural Killer Cells (NK Cells), Lineage^-^Sca1^-^c-Kit^+^ (LS^-^K), Granulocyte-Monocyte Progenitor (GMP), Megakaryocyte-Erythrocyte Progenitor (MEP), Common Myeloid Progenitor (CMP), Hematopoietic Stem Cells (HSCs), and Multipotent Progenitors 1-6 (MPP1-6).

We further characterized the hematopoietic stem and progenitor cells believed to contain leukemia initiating cells (LICs; gating strategy: Supplemental Fig 2B). First, we profiled the progenitor populations within the Lineage^-^Sca1^-^c-Kit^+^ (LS^-^K) population. We defined granulocyte-macrophage progenitors (GMP), megakaryocytic-erythroid progenitors (MEP) and common myeloid progenitor (CMP) cells using CD34 and FcγRII/III (CD16/CD32) expression^40^. *Npm1*^cA^/*Ptpn11*^E76K^ mice had a trend to decreased frequency of GMP and CMP cells compared to Mx-Cre or *Npm1*^cA^ mice (Fig 4B). This contrasts with the *Npm1^cA/+^*/*Nras^G12D/+^* and *Npm1^cA/+^*/*Flt3^ITD/+^* models, in which there was substantial increase in GMPs ^41^.

To further elucidate the stem and multipotent progenitor populations within *Npm1*^cA^/*Ptpn11*^E76K^ AML, we performed comprehensive profiling of the LSK population in the bone marrow (gating strategy: Supplemental Fig 2B). Previous reports of the *Ptpn11*^E76K^ mouse suggested that there was a decrease in the proportion of LSK cells in the bone marrow^21^. In this study, we did not see any decrease in the proportion of LSKs in the bone marrow of *Ptpn11*^E76K^ or *Npm1*^cA^/*Ptpn11*^E76K^ mice, however the low frequency of these cells and the relatively small number of mice analyzed may obscured differences between genotypes (Extended Data Fig 4B). Within the LSK population there was a significant skew away from long-term hematopoietic stem cells (LT-HSCs) (CD48^-^CD150^+^CD34^-^), MPP1 (CD48^-^CD150^+^CD34^+^), MPP2 (CD48^+^CD150^+^CD34^+^) cells that are primarily responsible for megakaryocyte and erythroid differentiation. Conversely, there was an expansion of MPP5 and MPP6 cells, which together are often identified within the short-term hematopoietic stem cell (ST-HSC) population (CD48^-^ CD150^-^) but are differentiated by CD135 expression (Fig 4C-D). Unlike MPP3 and MPP4 cells, which have been shown to have a myeloid and lymphoid bias respectively, it has been suggested that MPP5 cells can differentiate into MPP2-4 and are responsible for emergency myelopoiesis and stable engraftment of lymphocytes, whereas MPP6 (CD135+) display long-term multilineage contribution to hematopoiesis similar to HSCs^42^. The implications of these changes in the MPPs have yet to be fully elucidated in the *Npm1*^cA^/*Ptpn11*^E76K^ mouse. However, they contrast with what has been observed in *Npm1^cA^/Dnmt3a^R878H^*, which is a decrease in the ST-HSC population as well as increased MPP3.^43^ Furthermore, it is notable that we do not see any expansion of the MPP3/4 (CD48^+^CD150^−^) cells that have previously been reported to be the cells responsible for the propagation of the leukemia in secondary recipients in the *Tet2*^−/−^*/Flt3*^ITD^ AML model.^38^

### Npm1^cA^/Ptpn11^E76K^ splenocytes can recapitulate a leukemic phenotype in both immunodeficient and immunocompetent recipients

Thus far, we have demonstrated that co-induction of *Npm1*^cA^ and *Ptpn11*^E76K^ in mice validates our human data and that these two mutations are sufficient to develop a 100% penetrant AML including expansion of immature myeloid populations resulting in bone marrow failure. However, a hallmark of a true AML model requires that it can be adoptively transferred into secondary recipients with preservation of its immunophenotypic features. We therefore wanted to determine if AML cells from the *Npm1*^cA^/*Ptpn11*^E76K^ mice formed a lethal leukemia in a secondary recipient.

As most murine models of AML use fully immunodeficient or heavily preconditioned recipients, we first engrafted immunocompromised Prkdc^em26Cd52^Il2rg^em26Cd22^/NjuCrl (NCG) mice with bulk splenocytes collected from *Npm1*^cA^/*Ptpn11*^E76K^ mice at survival endpoint. We found that these splenocytes were able to recapitulate a leukemic phenotype with an average overall survival of 8.2 weeks post-engraftment (individual donors ranged from 7.5 to 17 weeks, n=8, Fig 5A). While all individual AML donors formed a lethal leukemia in some recipients, 2 of the 8 donors tested (S0283 and S0500) had slightly more inconsistent engraftment (based on the presence of donor tumor cell burden in the peripheral blood, Fig 5B), resulting in failed engraftment in 2 of 5 and 1 of 4 recipients, respectively. However, given that interactions of tumor cells with the immune microenvironment is such an important aspect of tumorigenesis, we decided to investigate if an intact recipient immune system altered the engraftment of the *Npm1*^cA^/*Ptpn11*^E76K^ tumor cells. PepBoy (B6.SJL-*Ptprc^a^ Pepc^b^*/BoyJ) recipients were similarly engrafted with bulk splenocytes from *Npm1*^cA^/*Ptpn11*^E76K^ donors (n=7). For recipient mice that developed disease, the median survival of PepBoy recipients was similar to the NCG recipients (10.4 weeks; individual donors ranging from 9.7 to 17.5 weeks, Fig. 5C). PepBoy recipient mice were consistently repopulated with donor tumor cells (CD45.2+ cells, Fig 5D), and displayed signs and symptoms consistent with the development of leukemia such as splenomegaly, leukocytosis, anemia, and thrombocytopenia (Fig 5E-F). The frequency of engraftment failure was significantly higher in PepBoy recipients with only 16 of the total 36 recipients meeting the survival endpoint (Fig 5C). Mice showing no evidence of engraftment were monitored for up to 20 weeks from engraftment (Fig 5C-D). These findings suggest potential involvement by the immune system in contributing to the failed engraftment in these mice, and we subsequently interrogated changes in the host and donor immune phenotype in this model.

**Figure 5.**
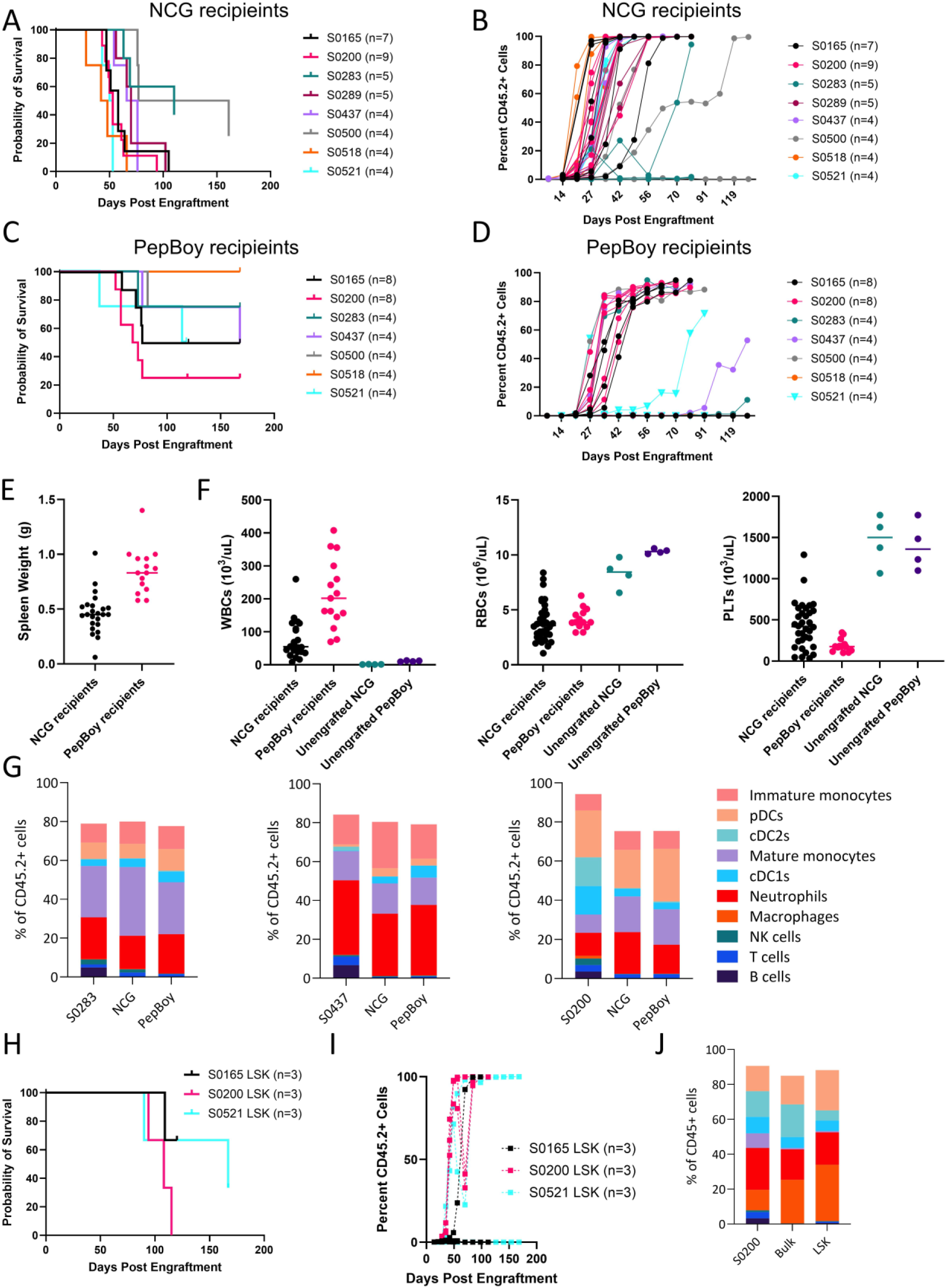
Engraftment of Npm1^cA^/Ptpn11^E76K^ splenocytes into both immunodeficient and immunocompetent mice. Overall survival and percent engraftment over time in NCG (**A-B**) and PepBoy (**C-D**) recipients engrafted with *Npm1*^cA^/*Ptpn11*^E76K^ splenocytes in three independent experiments. **E.** Spleen weights and **F.** White blood cells (WBCs), red blood cells (RBCs) and platelets (PLTs) of NCG and PepBoy recipients at survival endpoint. **G.** Summary of spleen immune subsets as a proportion of total CD45.2^+^ cells in primary AML donors compared to NCG and PepBoy recipients meeting the survival endpoint. **H.** Overall survival of NCG recipients engrafted LSK cells from 3 different donors. **I.** Frequency of donor derived CD45.2+ cells over time in the peripheral blood. **J.** Immune cell subsets in spleen from one donor (S0200) and recipients of either bulk splenocytes or LSK cells at endpoint. The legend for Figure 5J is the same as Figure 5G. Abbreviations: Plasmacytoid Dendritic Cells (pDCs), Conventional Dendritic Cells 2 (cDC2s), Conventional Dendritic Cells 1 (cDC1s), and Natural Killer Cells (NK Cells).

To confirm that the leukemic cells developing in both the immunodeficient and immunocompetent recipients phenotypically matched the donor cells, we performed the immunophenotyping panel on splenocytes from mice with overt leukemia as previously described. When gating on engrafted CD45.2 positive cells in both the NGC and PepBoy recipients, we found the overall diversity of cell subsets shows a similar profile to that of the donor, which suggests that *Npm1*^cA^/*Ptpn11*^E76K^ splenocytes can reproduce a phenotypically comparable AML (Fig 5G). However, given the inconsistent engraftment into PepBoy recipients, we wanted to determine if the acquisition of additional mutations, either in the primary donor or during the engraftment, was responsible for the ability of the *Npm1*^cA^/*Ptpn11*^E76K^ splenocytes to recapitulate a lethal AML phenotype. DNA from splenocytes was sent for targeted DNA sequencing using the Memorial Sloan Kettering Mouse Impact panel^43^. This analysis revealed that 3 of the 6 donors carried a *Flt3* mutation (p.D842G) that has previously been shown to be present in *Npm1*^cA^ AML mouse model^41^. While this mutation was present in both donors that readily form lethal leukemias in PepBoy recipients (S0165 and S0200), it was also present in S0518 that was incapable of engrafting PepBoy recipients. Interestingly, in recipients of S0165 and S0200, the *Flt3* mutation increased from 0.08 and 0.02 in the donors to between 0.38 and 0.49 in the recipients, respectively. Additionally, 4 of the 6 donors showed independent *Nf1* truncating mutations, however this mutation was similarly not associated with improved engraftment but rather was completely lost in the recipient leukemia cells derived from 2 of the 3 successful donors (Supplemental Data Table 1). Therefore, while these mutations may play a role in leukemogenesis, we posit no relationship between additional acquired mutations and engraftment success or disease lethality. Of note, some recipients did acquire additional mutations not identified in the donors, including *Kit*, *Kmt2d*, *Ros1*, and *Vegfa* mutations. However, acquisition of additional mutations was not required for successful engraftment and while each of these mutations have been identified as playing a role in AML pathogenesis, we have insufficient numbers here to understand if the acquisition of any of these mutations accelerated disease development.

### Npm1^cA^/Ptpn11^E76K^ LSKs can recapitulate the immune subset diversity in NCG recipients

With Mx-Cre driven expression, all hematopoietic lineage cells in the *Npm1^cA^/Ptpn11^E76K^* mice express the driver mutations. However, not all leukemia populations are expected to have the same capacity to engraft into secondary recipients. Given the diversity and variable states of differentiation of both the leukemia cells present in the *Npm1*^cA^/*Ptpn11*^E76K^ mice, we wanted to determine if the same profile of mature phenotype cells would develop in mice engrafted with LSK cells. Therefore, LSKs were sorted from the spleens of individual donors (n=3) and injected via the tail vein into NCG recipients (n=3; Sorting Strategy: Supplemental Fig 3). Unexpectedly, the LSK cells engrafted more slowly and had delayed development of lethal disease compared with the bulk splenocytes. Furthermore, there was considerable variability in the engraftment of the LSK cells into NCG recipients with donors producing a lethal leukemia in 1 of 3, 2 of 3, and 3 of 3 recipients (Fig 5H-I). We again performed immune profiling of the splenocytes from recipient mice that developed a lethal leukemia, and the immune subset diversity in LSK engrafted mice was comparable to bulk engrafted recipients (Fig 5J). This suggests that the diverse mature myeloid populations in the *Npm1*^cA^/*Ptpn11*^E76K^ AML are derived from a stem or progenitor population within the LSK cells. However, the increased latency and failure to engraft into NCG recipients suggest that more mature subsets contribute to the efficient propagation of a lethal leukemia in immunodeficient recipients.

### Immunophenotypic characterization of human PTPN11 mutated AML confirms surface expression diversity and identifies recurrent pDC population

Given the immunophenotypic diversity observed within the AML population in the *Npm1*^cA^/*Ptpn11*^E76K^ mouse model and the ability of LSKs to recapitulate the full spectrum of mature myeloid populations in engrafted mice, we sought to understand if this reflected the immunophenotype of human AML patients with *PTPN11* mutations. Concurrent with the scDNA-seq (Fig 1), we performed simultaneous single cell immunophenotyping using a 42-specificity surface antigen antibody panel in primary AML samples. Distinct immunophenotypic clusters were identified using shared nearest neighbor clustering. Heatmaps were generated to visualize the co-expression of markers to allow for cell type identification (Fig 6A, E; Extended Data Fig 5). The clusters identified as T cells, NK cells, and B cells based on CD3, CD4, CD8, CD56, and CD19 expression primarily consisted of wild type (WT) cells. Conversely, cells containing the AML mutations showed substantial diversity including clusters that aligned with monocytic (CD11b, CD14, CD163, CD64, HLA-DR; pink) and pDC-like (CD11c, HLADR, CD123, CD303/CD304; red) in addition to a cluster expressing markers associated with a classic AML blast phenotype (CD34, CD117, CD123, CD33; navy; Fig 6A).

**Figure 6.**
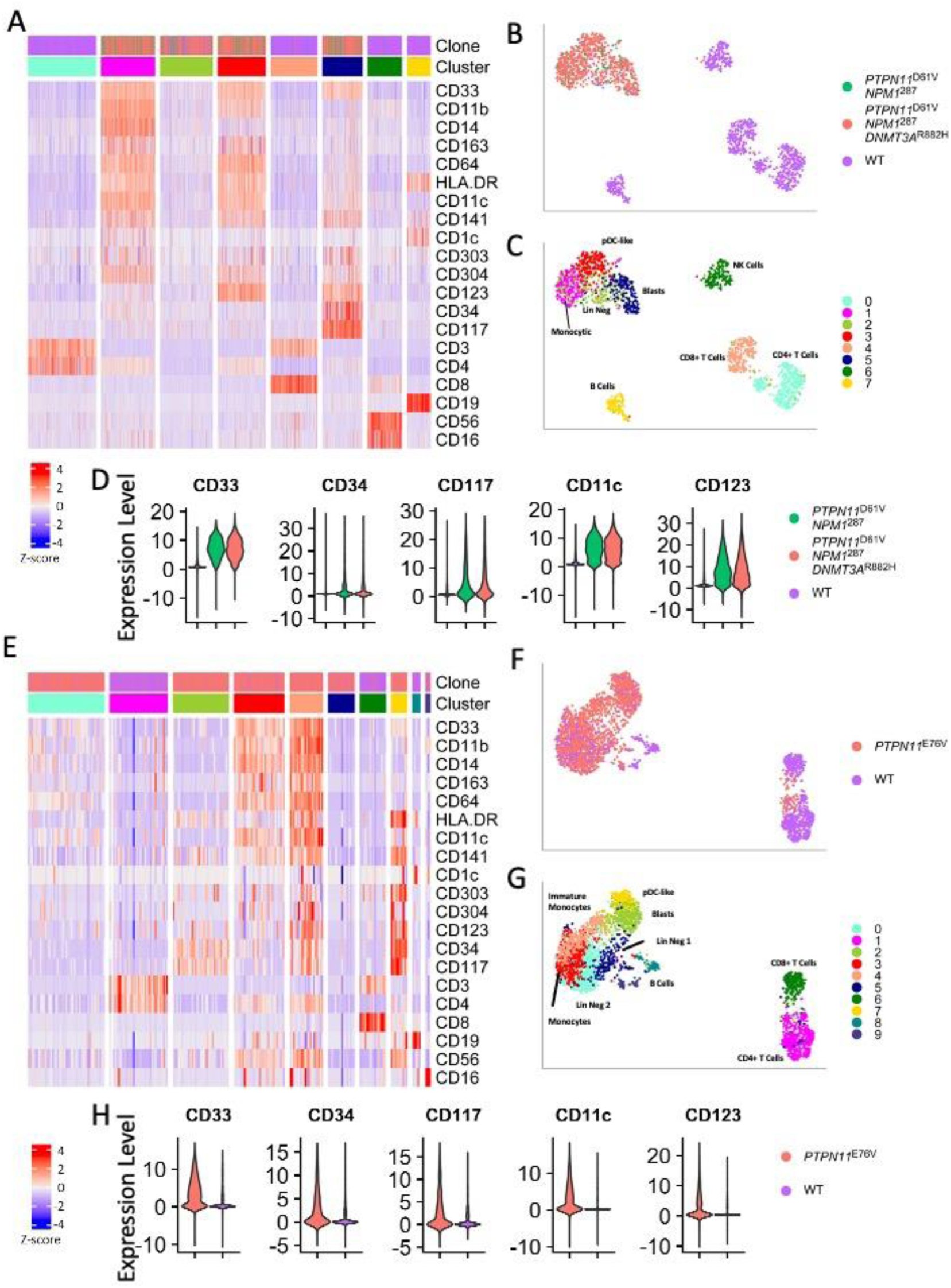
Immunophenotypic diversity of representative early PTPN11 mutant AML. **A-D.** Immunophenotypic clusters were identified using shared nearest neighbor clustering on a representative early *PTPN11* mutant sample (AML-6): **A.** Heatmap of select lineage defining markers. **B.** UMAP overlaid with WT, *NPM1*^287^/*PTPN11*^D61V^ and *NPM1*^287^/*PTPN11*^D61V^/*DNMT3A*^R882H^ clone identity. **C.** UMAP overlaid with cluster identity, cell types were assigned manually based on marker co-expression. **D.** Violin plots of CD33, CD34, CD117, CD11c, and CD123 on AML clones compared to WT. **E-F.** The same analysis was performed on a representative *PTPN11*-mutated sample without *NPM1* mutation (AML-3): **E.** Heatmap of select lineage defining markers. **F.** UMAP overlaid with WT, and *PTPN11*^E76V^ clone identity. **G.** UMAP overlaid with cluster identity, cell types were assigned manually based on marker co-expression. **H.** Violin plots of CD33, CD34, CD117, CD11c, and CD123 on *PTPN11*^mut^ clone compared to WT. Abbreviations: Lineage Negative (Lin Negative), Plasmacytoid Dendritic Cells-Like (pDCs-like), and Natural Killer Cells (NK Cells)

We next generated a UMAP based on the surface marker expression. As expected, the AML clones clustered separately from the WT cells within a patient sample (Fig 6B; Extended data Fig 5). However, we found no distinction between the immunophenotype of subclones (Fig 6C-D). Indeed, in patients with subclonal *PTPN11* mutations, we found no distinction between *NPM1* mutant subclones with or without *PTPN11* suggesting that these mutations do not drive a specific cell type identity (Extended Fig 5). We next compared the expression of AML blast cell markers including CD33, CD34, and CD117 as well as pDC associated markers CD11c and CD123 and saw substantial expression of both blast and pDC markers in the AML populations compared to the WT cells (Fig 6D; Supplemental Data Table 2). We next sought to interrogate if this diverse phenotype was also associated with early concurrent *PTPN11* and *NPM1* mutations. We observed a similar diversity of immunophenotype in a *PTPN11*-mutated patient that lacked an *NPM1* mutation (Fig 6E). Again, in this sample, we identified different populations of WT and leukemia cells including a cluster with expression of markers associated with immature cells (CD34, CD117; light green) as well as pDC-like (CD141, HLADR, CD303; yellow, Fig 6F-H). Our data suggests that AML patients with *PTPN11* mutations have a diverse AML immunophenotype in which AML clones regardless of when the *PTPN11* mutation occurs in AML development.

## Discussion

Herein, using single-cell DNA sequencing of AML patient samples, we present data that for the first time demonstrates that *PTPN11* can act as an early driver mutation in AML when accompanied by a strong oncogene such as *NPM1* mutation or rearrangement of *EVI1* or *KMT2A*. Given the co-association of *PTPN11* and *NPM1* mutations early in leukemogenesis, we then developed a novel transgenic mouse model concurrently expressing the *Ptpn11*^E76K^ and *Npm1*^cA^ alleles to further investigate the disease course of *PTPN11* mutant AML. Mice comprising both *Ptpn11*^E76K^ and *Npm1*^cA^ mutations rapidly developed a fully penetrant lethal myeloid neoplasm that recapitulates many of the features of *PTPN11*-mutated AML in humans. Primary transgenic *Npm1^cA^/Ptpn11^E76K^* mice present with an expansion of both early c-Kit positive precursors and a broad expansion of variably differentiated myeloid cells including cells bearing an immunophenotype of plasmacytoid dendritic cells. Finally, we assessed if the immunophenotypic diversity observed in the *Npm1^cA^/Ptpn11^E76K^* mouse model was also present in the human AML patients with *PTPN11* mutations. Indeed, patients with *PTPN11* mutations showed considerable expression of markers associated with mature myeloid cells. In addition to clusters expressing markers of immature myeloid cells (blasts), we identified clusters of cells with the AML mutations that aligned with both mature monocytic cells and plasmacytoid dendritic cells.

Prior work from our group showing a high variant allele frequency of *PTPN11* mutations in some AML patients suggested that *PTPN11* may be an early co-event in leukemogenesis. The findings of our scDNA-seq analysis corroborate this, confirming that *PTPN11* can be both a late acquired sub-clonal mutation or an early acquired driving mutation together with another strong oncogene (*EVI1, KMT2A,* or *NPM1* mutation). However, two limitations of the scDNA-seq technology utilized limited our ability to fully understand the clonal evolution in these patients. First, the inability to integrate information about chromosomal rearrangements at a single-cell level. This was particularly pertinent in the *PTPN11* mutant patients without a concurrent *NPM1* mutation, which are frequently associated with chromosome rearrangements but had few co-occurring mutations based on the bulk sequencing analysis. In these patients, we could not infer the order of acquisition of the *PTPN11* mutation relative to the chromosome rearrangement. Secondly, due to the limited coverage of the scDNA-seq panel multiple high VAF mutations identified in the bulk sequencing were not captured in the single-cell data. Therefore, while we can infer the order of acquisition for the captured mutations, this information is incomplete. Conversely, the scDNA-seq reliably captured some low frequency mutations not identified in the bulk sequencing, including *NRAS, KRAS*, and *WT1* mutations.

Despite the identification of *PTPN11* mutations being associated with resistance to multiple targeted therapies in AML, there has not been an AML murine model created that can be utilized to study the mechanism of resistance to diverse targeted therapies and to develop new therapeutic agents for this population.^5–8,44,45^ We therefore characterized the *Npm1*^cA^*/Ptpn11*^E76K^ mouse for future use in this context. This model consistently develops an AML phenotype as evidenced by accumulation of myeloid cells in both primary and secondary lymphoid organs, and progressive anemia, thrombocytopenia, and leukocytosis at time of death. The *Npm1*^cA^/*Ptpn11*^E76K^ model has a median survival of approximately 4 months from induction which is shorter than the *Npm1*^cA^*/Dnmt3A*^R878H^ model^43^, longer than the *Npm1*^cA^/*Flt3*^ITD^ model, and comparable to the *Npm1*^cA^/*Nras*^G12D^ model^41^. In addition to similar disease kinetics, the *Npm1*^cA^/*Ptpn11*^E76K^ and *Npm1*^cA^/*Nras*^G12D^ also share a propensity for an AML with maturation phenotype. Comprehensive immune profiling of both splenocytes and bone marrow cells showed considerable diversity in the immunophenotype of *Npm1*^cA^*/Ptpn11*^E76K^ mice. *PTPN11* mutations have been previously associated with a monocytic phenotype^46^, and we indeed identified a significant expansion of a population of immature monocytic cells. However, we also identified expansion of cells with a mature phenotype including a significant expansion of pDCs. In our mouse model, we found that the pDCs expand in the peripheral blood at the same rate as the leukemic blasts and were also identified in the spleen and bone marrow of *Npm1*^cA^/*Ptpn11*^E76K^ mice with overt leukemia. Finally, we found that the mature myeloid populations, including the pDCs, were capable of engraftment in both immunodeficient and fully immunocompetent secondary recipients, and importantly could be derived from the LSK stem/progenitor population.

Following identification of the diversity of the immunophenotype in our mouse model, we specifically looked for these same populations in our patients’ samples and were able to identify similar populations of mature and immature monocytes. One major difference is that we were not able to identify of a population of neutrophils, which is likely related to a difference in cell processing as the human blood and bone marrow samples were mononuclear cells isolated by density gradient centrifugation whereas the mouse cells were not. Finally, we looked for the presence of pDCs in the human samples. In addition to being frequently over expressed in human AML samples, CD123 is expressed on immature myeloid cells as well as pDCs^47,48^. There are only a few studies describing pDCs in AML, and in general have found that the pDCs, defined as cells co-expressing CD123 and HLA-DR, have an otherwise diverse immunophenotype and are commonly associated with mutations associated with poor outcome, particularly *RUNX1*^49–51^. The broad immunophenotyping panel ran in these experiments allowed us to determine if the CD123^+^ cells were also expressing additional pDC markers (HLA-DR and CD303/CD304) or progenitor cell markers (CD117 and CD34). We found that while we could identify CD123 expressing clusters that were consistent with progenitor-like AML cells, we could also identify clusters within the leukemic clone that also expressed the markers of pDCs. Taken together this data indicates that our murine model of *PTPN11* and *NPM1* mutated AML can reliably phenocopy human AML with a pDC differentiation.

Notably, engraftment of LSKs into NSG mice without conditioning recapitulated the broad array of mature myeloid cells including the pDC population corroborating previous reports that the pDCs in AML are derived from the leukemia stem or initiating cells^52^. Interestingly, engraftment of LSK cells was more inconsistent than unselected splenocytes from the same donors suggesting contribution of the more mature myeloid cell populations in efficient pathogenesis of *Ptpn11*^E76K^/*Npm1*^cA^ AML. Further work is needed to delineate how the mature immune populations that comprise the AML contribute to the engraftment of LSK cells. Further work to understand how pDCs contribute to disease development can be pursued using this mouse model. In chronic myelomonocytic leukemia (CMML) patients with *RAS* pathway mutations, pDCs have recently been identified and shown to correlate with an increase in regulatory T cells and acute leukemia transformation^53^. Additional work is needed to understand if and how the pDCs contribute to AML development and potentially therapeutic resistance. Because the *Npm1*^cA^/*Ptpn11*^E76K^ mouse model is phenotypically comparable to what is seen in human disease, this model will be useful for future work understanding the mechanisms of resistance in *PTPN11-* mutated patients and for developing therapies that will be effective in this patient population including testing the therapeutic benefit of targeting the pDC population which may have implication for CD123 targeting therapies.

In conclusion, we have demonstrated with both human and murine studies that *PTPN11* can have a divergent clonal hierarchy but can be a co-initiating factor with other strong oncogenes. Hence, *PTPN11* mutations are both early and late mediators of myeloid leukemia oncogenesis. We have generated a model of *Npm1* and *Ptpn11* mutated AML that can serve to study both the role of pDC expansion on disease progression as well as response to new therapeutics for this subset of AML which is historically very difficult to treat.

## Supporting information

Supplemental Data Statistics

## Acknowledgements

We would like to acknowledge Christopher Manring, Jean Truxall and Shelley Orwick at The Ohio State Comprehensive Cancer Center (OSU CCC). We would also like to acknowledge the OSU CCC Leukemia Tissue Bank Shared Resource (P30CA016058), NIH F30 (SF, F30CA265281), R35 (JCB, R35CA198183) and UG1 (JCB, CA233338-02). SF was supported by a Pelotonia graduate fellowship. All flow cytometry-assisted sorting was performed with the help of and using equipment maintained by the Research Flow Cytometry Facility in the Division of Rheumatology, and the sequencing was performed by the Genomic Sequencing Facility at Cincinnati Children’s Hospital Medical Center. We would also like to thank Troy Robinson and the Levine lab at Memorial Sloan Kettering for his assistance in preforming targeted sequencing of mouse samples.

