## Supplemental Methods and Figures for "*PTPN11* Mutation Clonal Hierarchy in Acute Myeloid Leukemia"

**TotalSeq Heme Oncology Cocktail Clones**

CD64 (clone 10.1), CD34 (clone 581), CD90 (clone 5e10), CD117 (clone 104D2), CD304 (clone 12C2), CD303 (clone 201A), CD45 (clone 2D1), CD16 (clone 3G8), CD56 (clone 5.1H11), CD123 (clone 6H6), CD49d (clone 9F10), FcɛR1ɑ (clone AER-37), CD62P (clone AK4), CD25 (clone BC96), CD30 (clone BY88), CD7 (clone CD7-6B7), CD71 (clone CY1G4). CD62L (clone DREG-56), CD69 (clone FN50), CD163 (clone GHI/61), CD83 (clone HB15e), CD45RA (clone HI100), CD10 (clone HI10a), CD19 (clone HIB19), CD38 (clone HIT2), CD11b (clone ICRF44), CD44 (clone IM7), CD1c (clone L161), HLA-DR (clone L243), CD14 (clone M5E2), CD141 (clone M80), CD138 (clone MI15), CD33 (clone P67.6), CD4 (clone RPA-T4), CD22 (clone S-HCL-1), CD11c (clone S-HCL-3), CD8 (clone SK1), CD2 (clone TS1/8), CD45RO (clone UCHL1), CD3 (clone UCHT1), CD5 (clone UCHT2), CD13 (clone WM15).

**Immunophenotyping Antibodies and Cell Surface Marker Profiles**

The following antibodies, obtained from BD Biosciences, Biolegend, and Thermo Fisher Scientific: CD11b (clone M1/70), LIVE/DEAD Fixable Blue, CD45.2 (clone 104), CD23 (clone B3B4), CD43 (clone S7), CD24 (clone 30-F1), I-A/I-E (clone 2G9, referred to as MHC II), CD27 (clone LG.3A10), Siglec-H (clone 440c), CD93 (clone AA4.1), Ter119 (clone TER-119), Ly6A/E (clone D7 or E13-161.7, referred to as Sca-1), CD45.1 (clone A20), Ly6C (clone HK1.4), NK1.1 (clone PK136), CX3CR1 (clone SA011F11), CD44 (clone IM7), CD8a (clone 5H10-1 or 53-6.7), IgM (R6-60.2), CD138 (clone 281-2), CD4 (clone GK1.5), CD11c (clone N418), Ly6G (clone 1A8), CD21/35 (clone 4E3), CD117 (clone ACK2, referred to as c-Kit), F4-80 (clone QA17A29), CD172a (clone P84), CD19 (clone 6D5), CD49b (clone DX5), CD206 (clone C068C2), CD317 (clone 927), CD3 (clone 17A2), IgD (clone 11-26c.2a), CD86 (clone GL1), CD5 (clone 53-7.3), B220 (RA3-6B2), CD45 (clone I3/2.3), CD48 (clone HM48-1), CD34 (clone SA376A4), CD150 (clone TC15-12F12.2), FcɛR1 (clone MAR-1), CD24 (clone M1/69), CD127 (clone SB/199), CD135 (clone A2F10), and CD16/32 (clone 2.4G2).

Cell surface marker profiles were defined as: neutrophils (CD45.2^+^/CD3^-^/CD19^-^/NK1.1^­-^/Ly6G^+^), macrophages (CD45.2^+^/CD3^-^/CD19^-^/NK1.1^­-^/Ly6G^-^/CD11c^-^/F4-80^+^), monocytes (CD45.2^+^/CD3^-^/CD19^-^/NK1.1^­-^/Ly6G^-^/CD11c^-^/CD11b^+^/F4/80^low^/CX3CR1^+^), cDC1s (CD45.2^+^/CD3^-^/CD19^-^/NK1.1^­-^/Ly6G^-^/CD11c^+^/B220^-^/MHC II^+^/CD172a^+/-^/CD11b^+^), cDC2s (CD45.2^+^/CD3^-^/CD19^-^/NK1.1^­-^/Ly6G^-^/CD11c^+^/B220^-^/MHC II^+^/CD172a^-^/CD11b^-^/CD24^+^/CD8a^+/-^), pDCs (CD45.2^+^/CD3^-^/CD19^-^/NK1.1^­-^/Ly6G^-^/CD11c^+^/MHC II^+^/B220^+^), MPP5 (CD45^+^/Ter119^-^/CD3^-^/B220^-^/Ly6C^-^/CD11b^-^/F4-80^-^/Ly6G^-^/cKit^+^/Sca1^+^/CD135^-^/CD34^+^/CD150^-^/CD48^-^), and MPP6 (CD45^+^/Ter119^-^/CD3^-^/B220^-^/Ly6C^-^/CD11b^-^/F4-80^-^/Ly6G^-^/cKit^+^/Sca1^+^/CD135^-^/CD34^-^/CD150^-^/CD48^-^).

### Supplemental Data

A

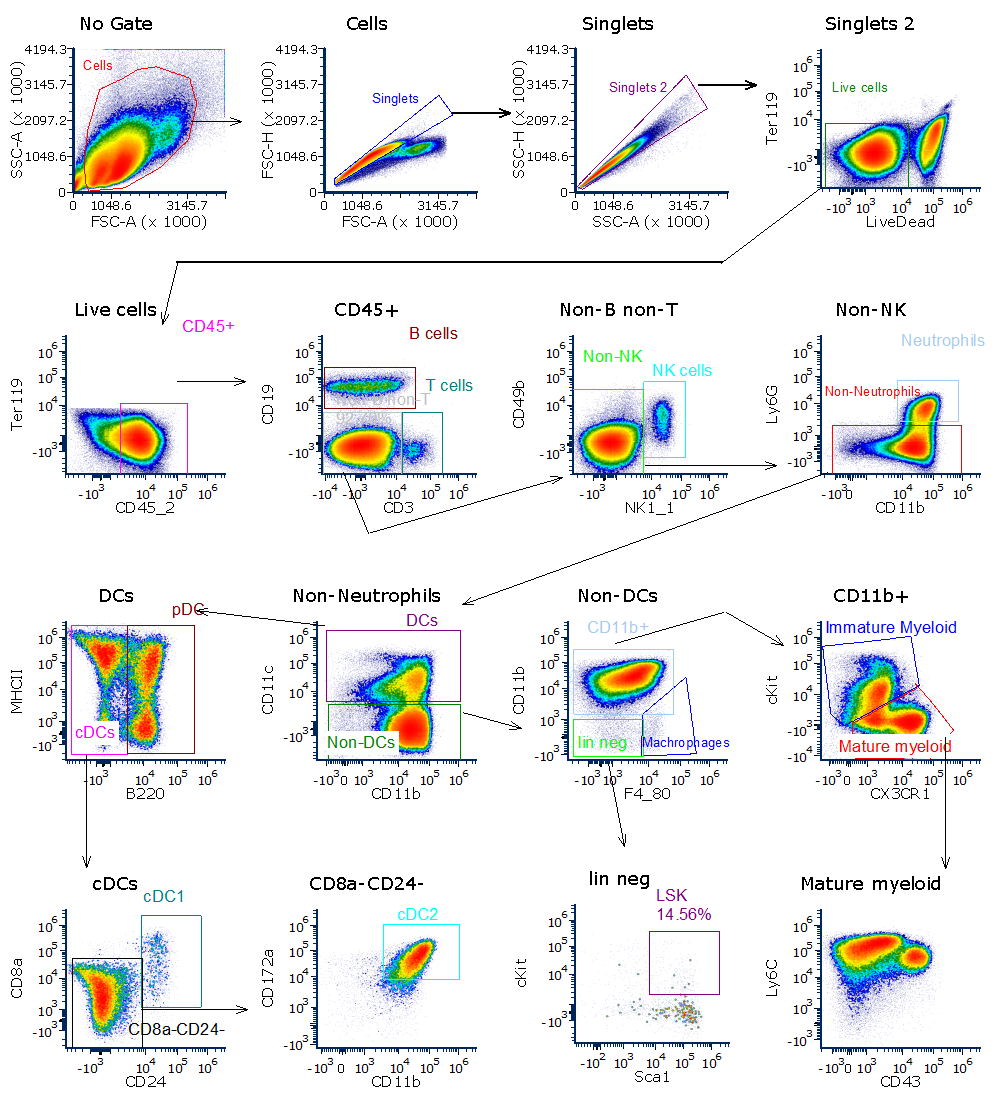

Supplemental Figure 1. Gating strategy for spleen immunophenotyping panel for leukocyte populations and LSK cells.

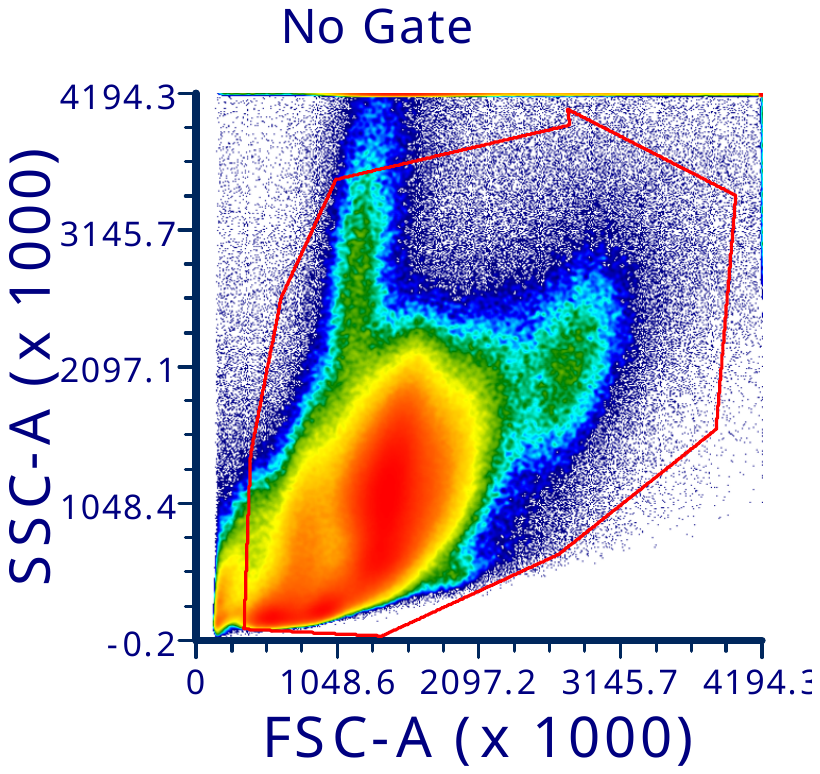

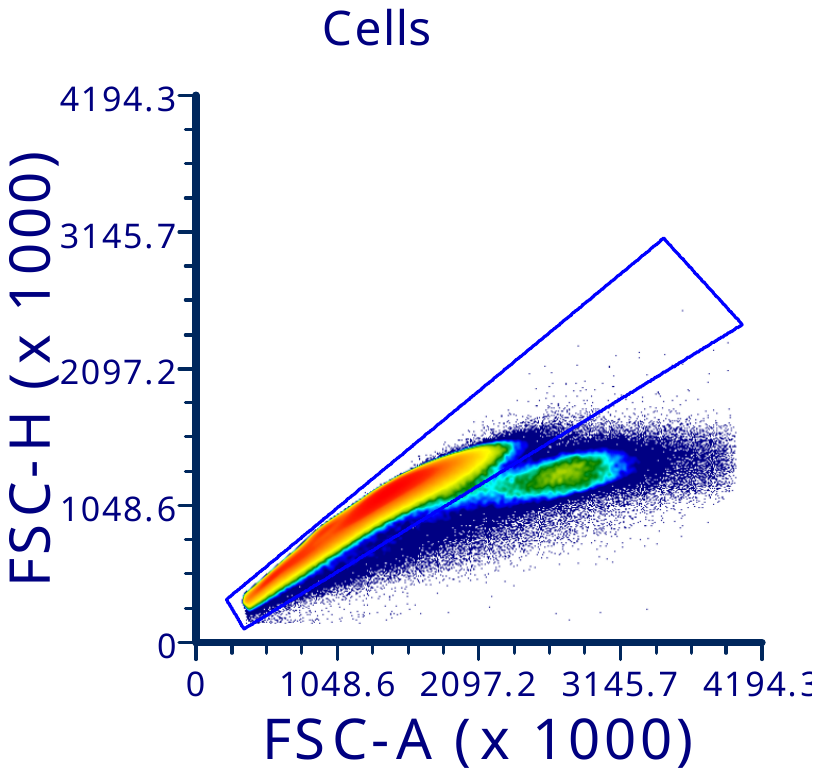

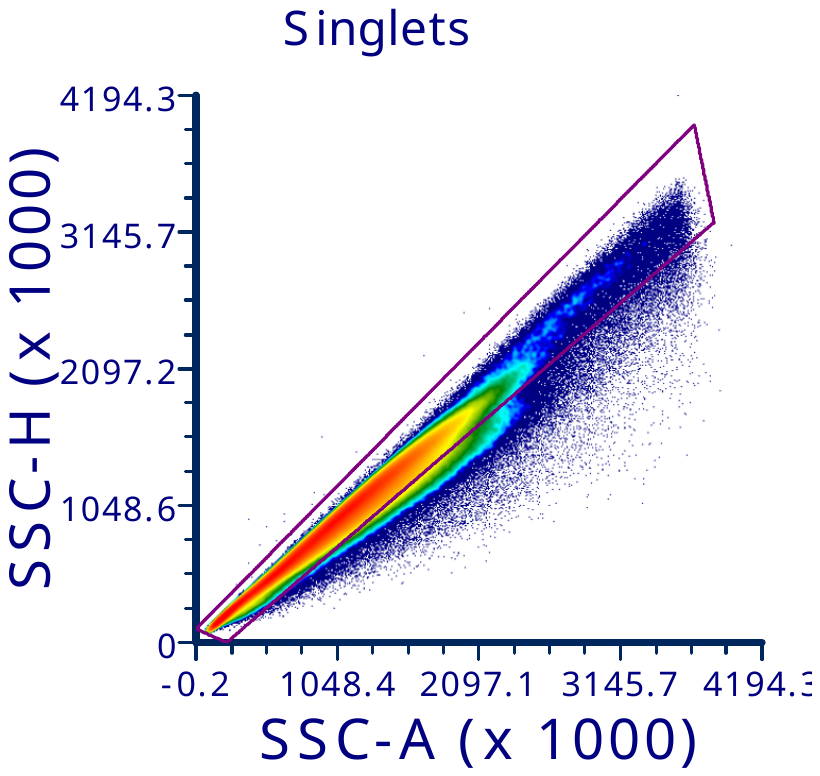

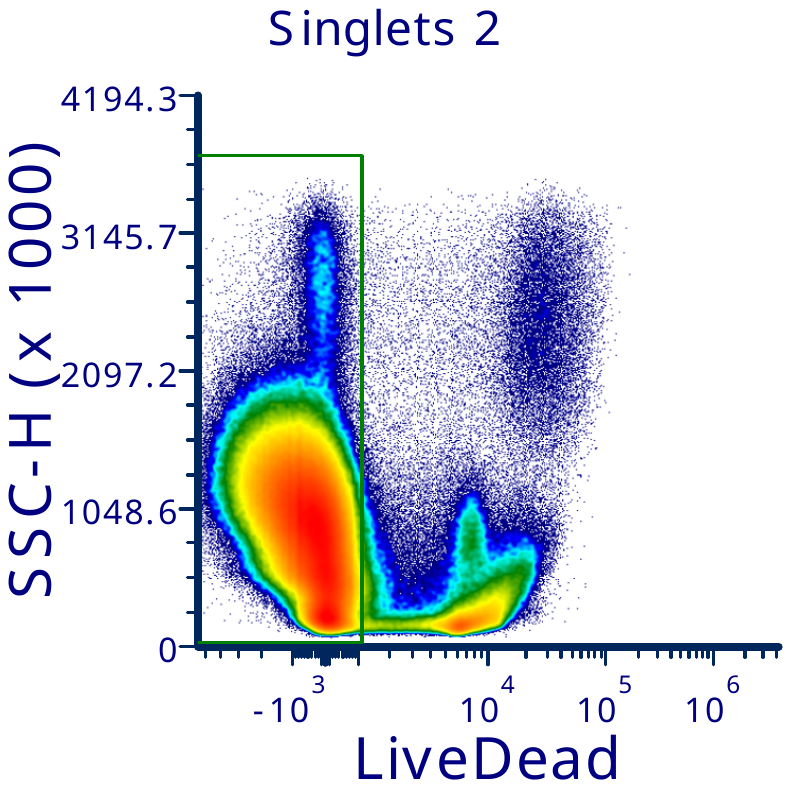

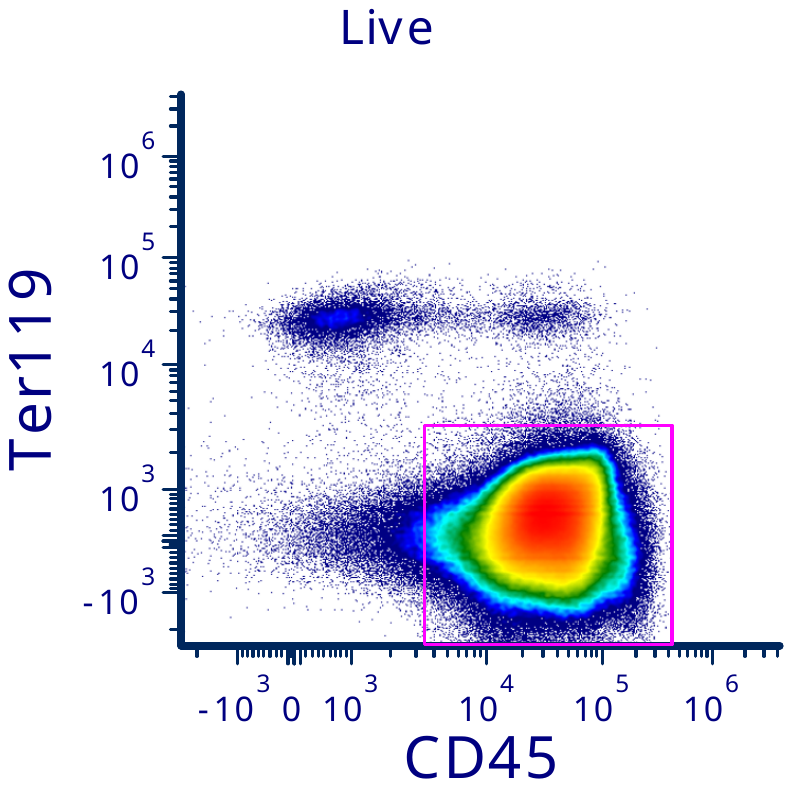

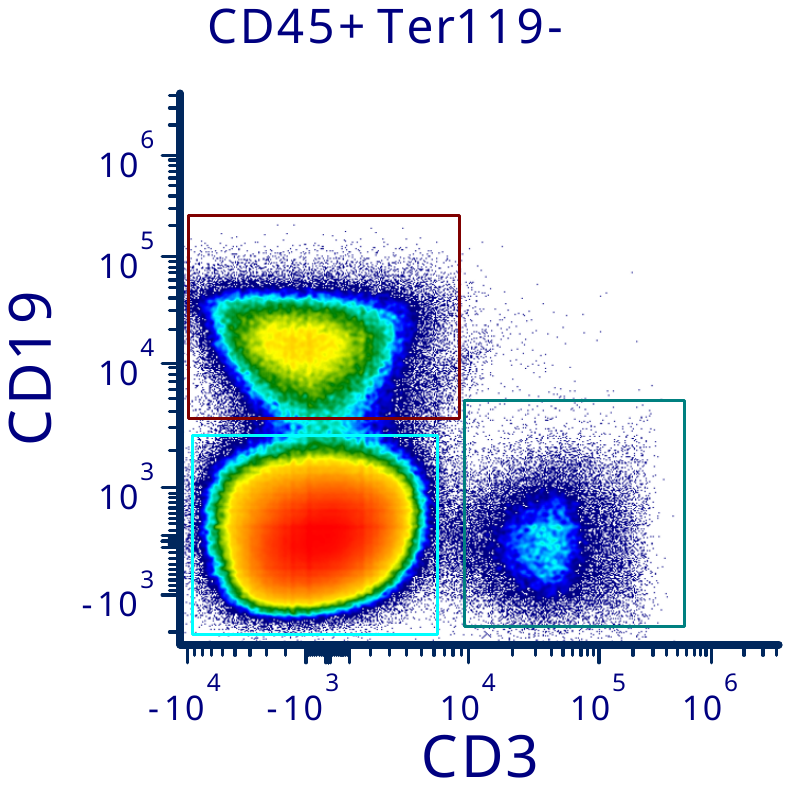

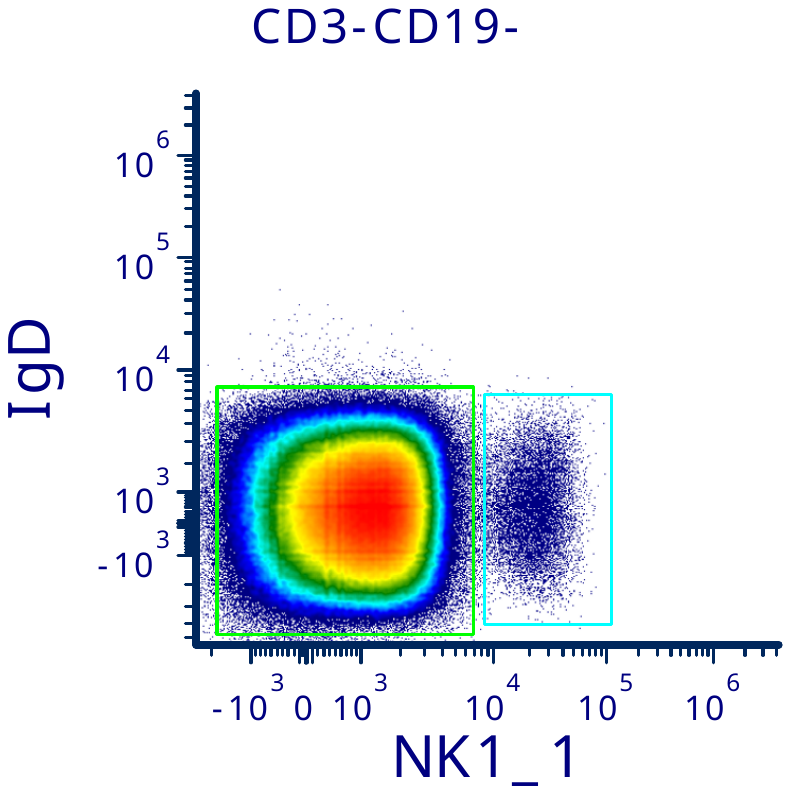

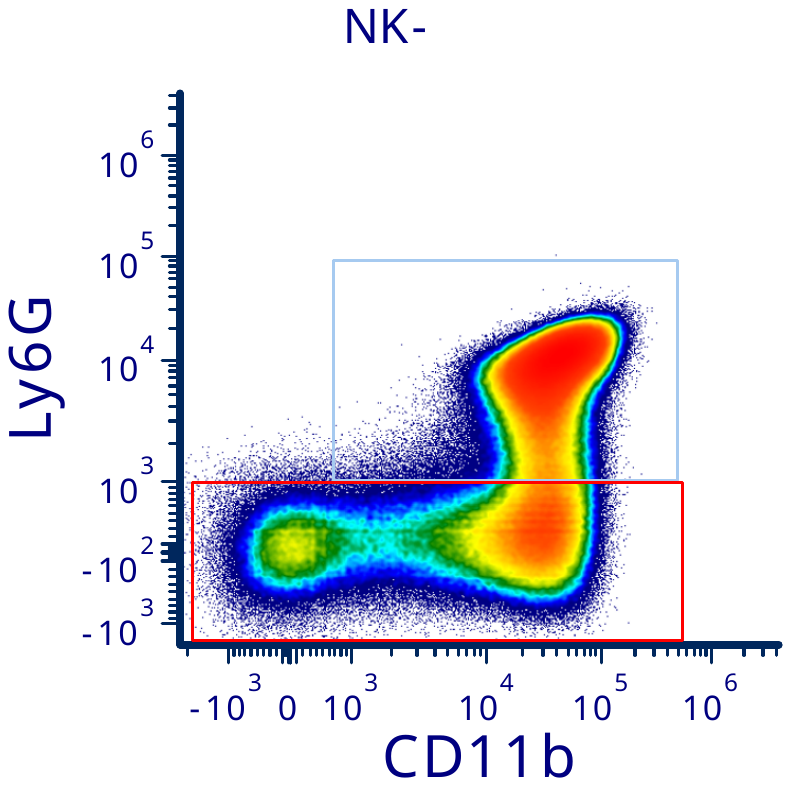

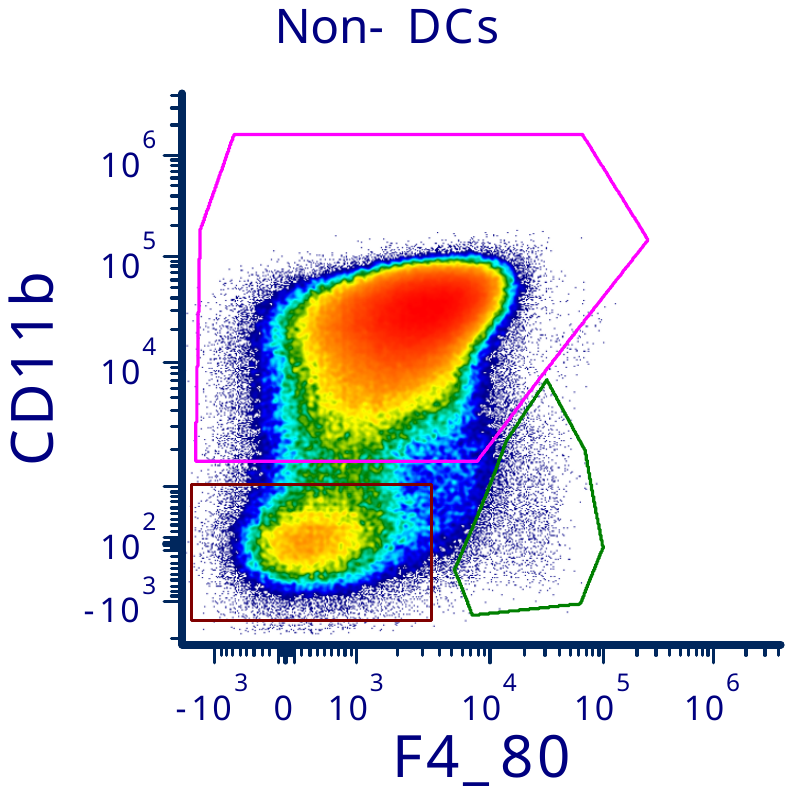

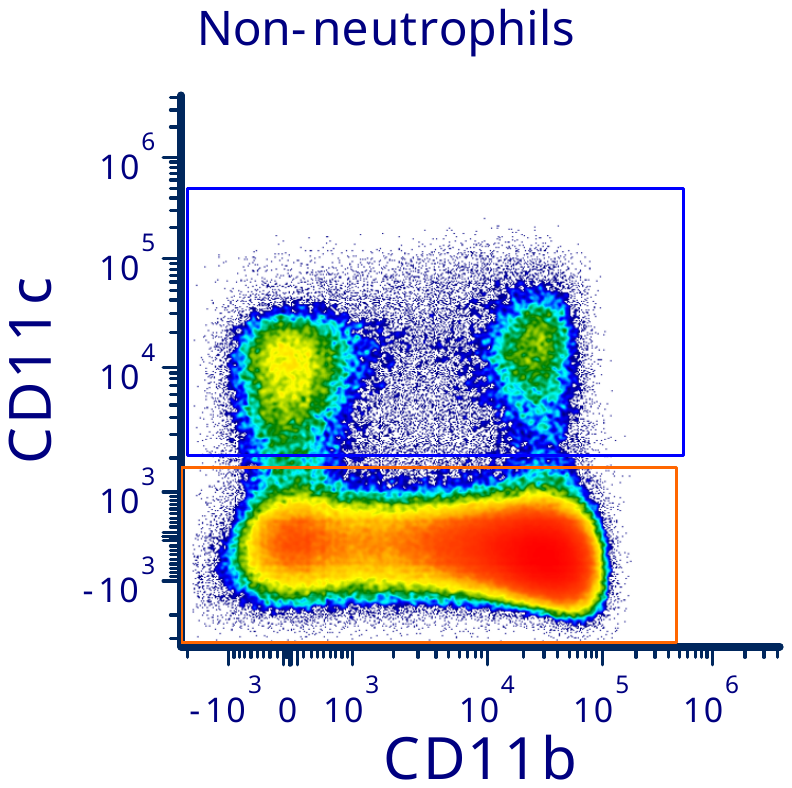

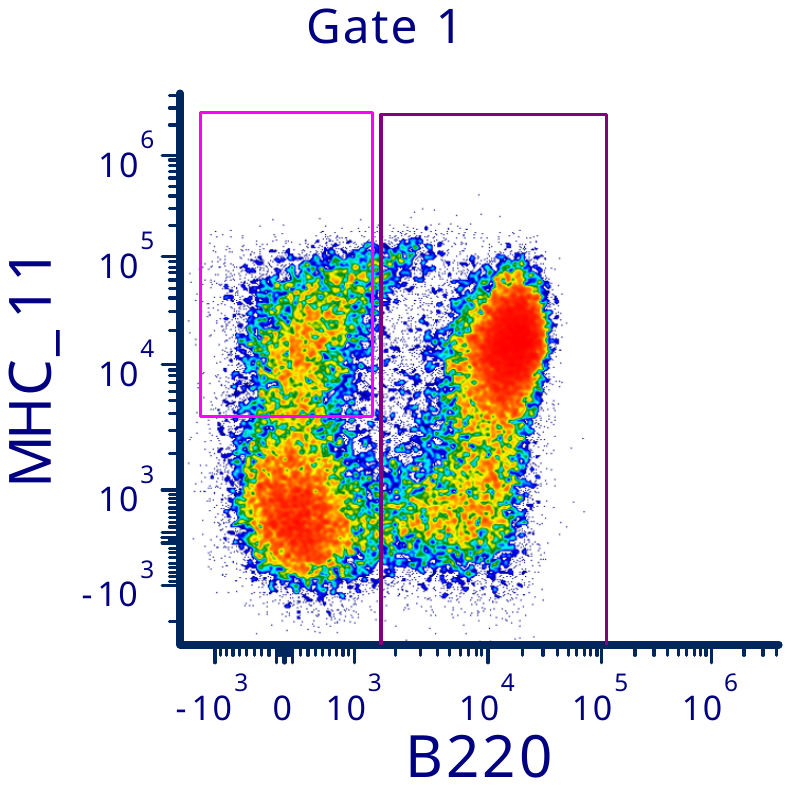

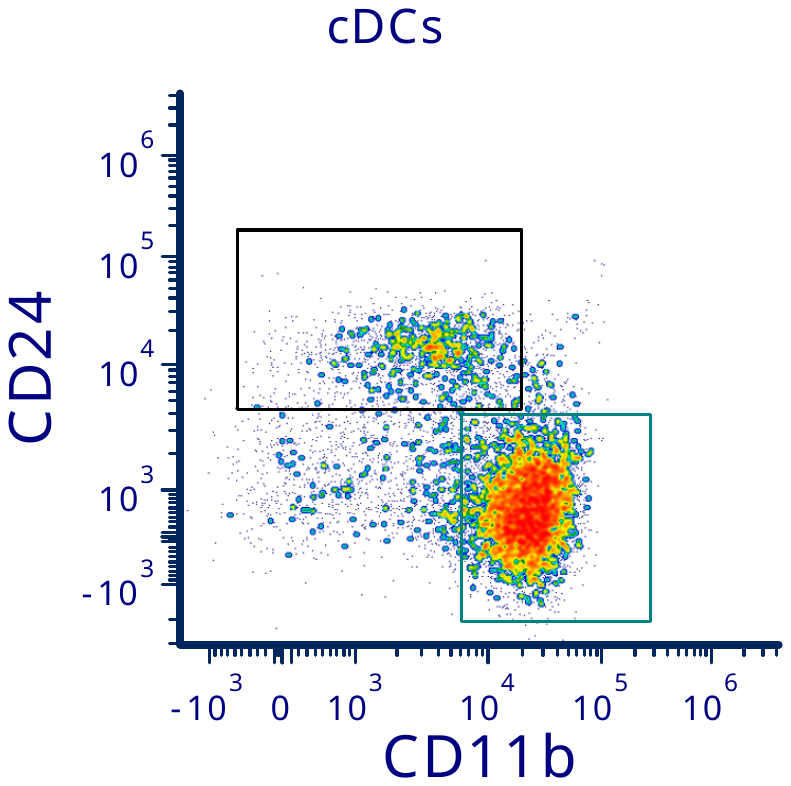

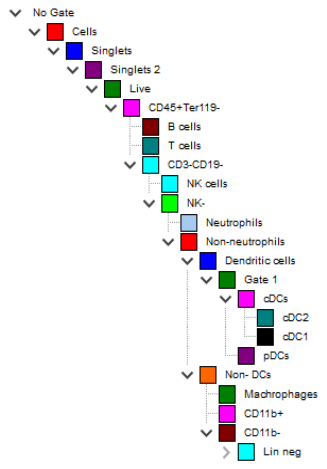

A

**Supplemental Figure 2**. Gating strategy for bone marrow immunophenotyping panel for **A.** mature white blood cells and **B.** subtypes of lineage negative cells.

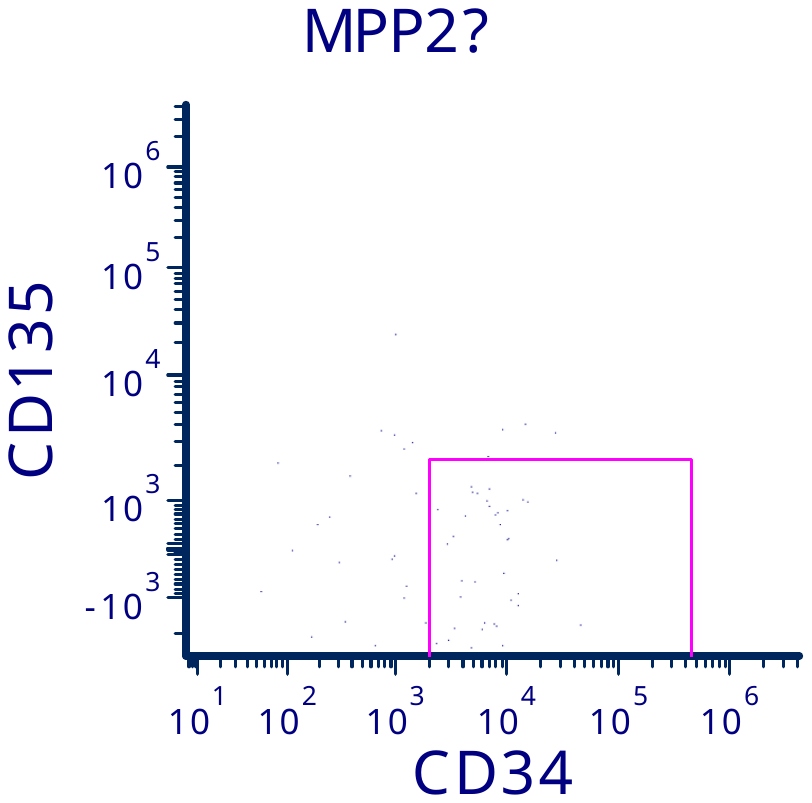

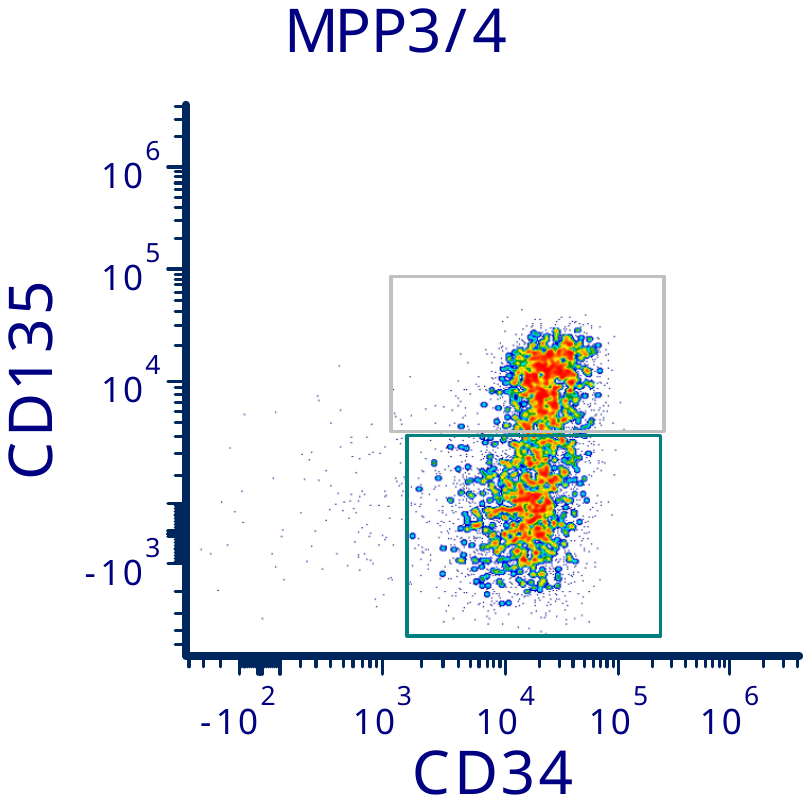

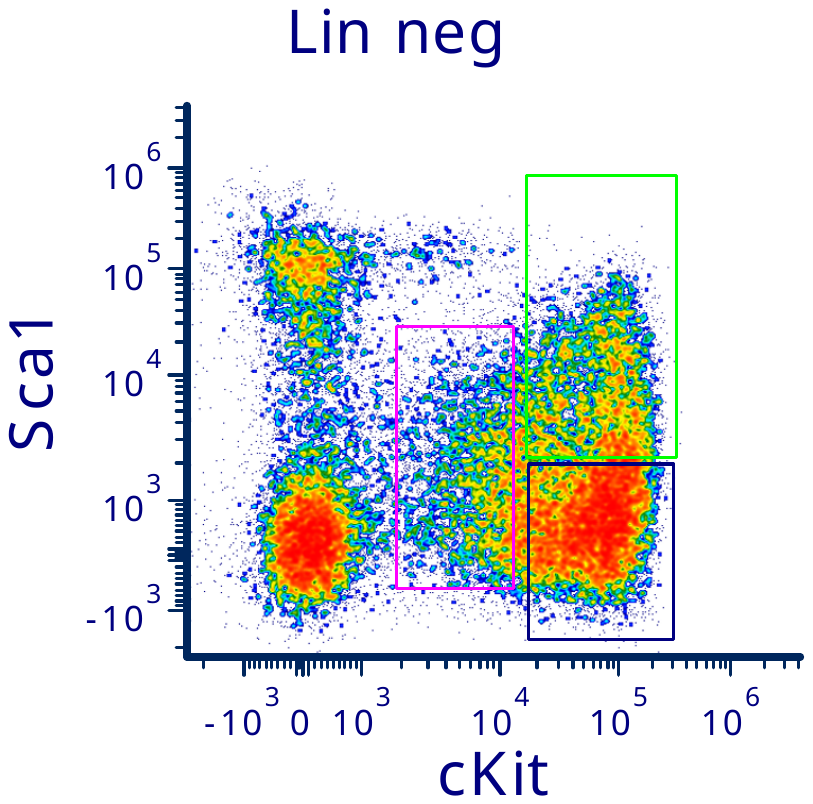

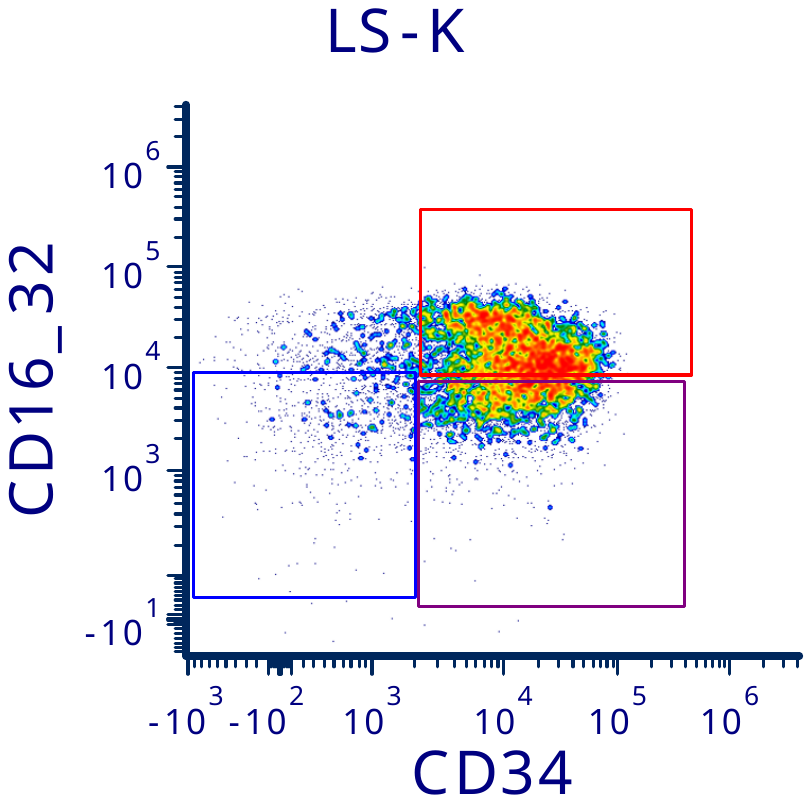

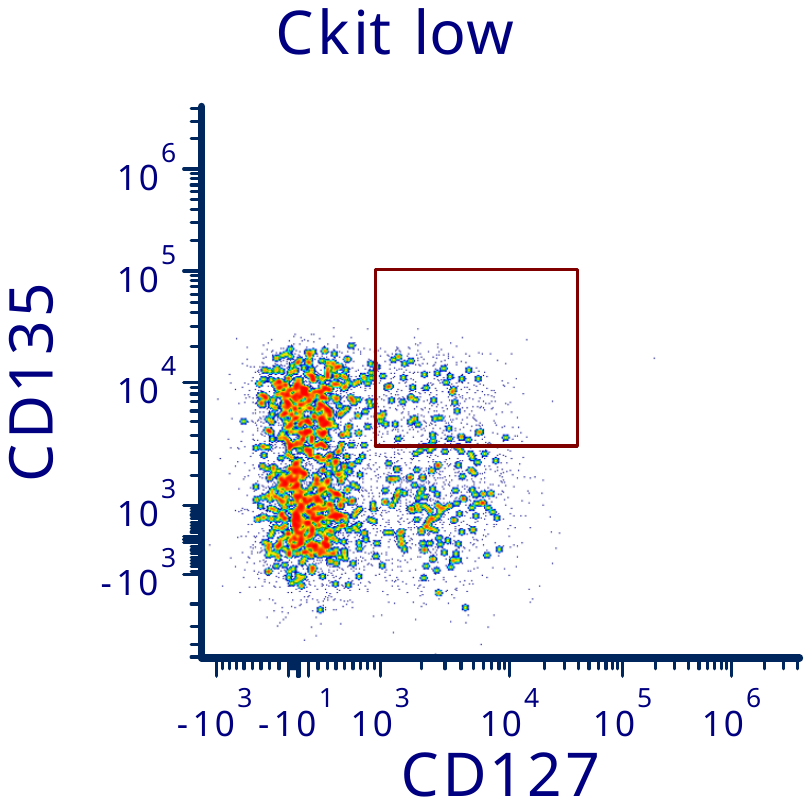

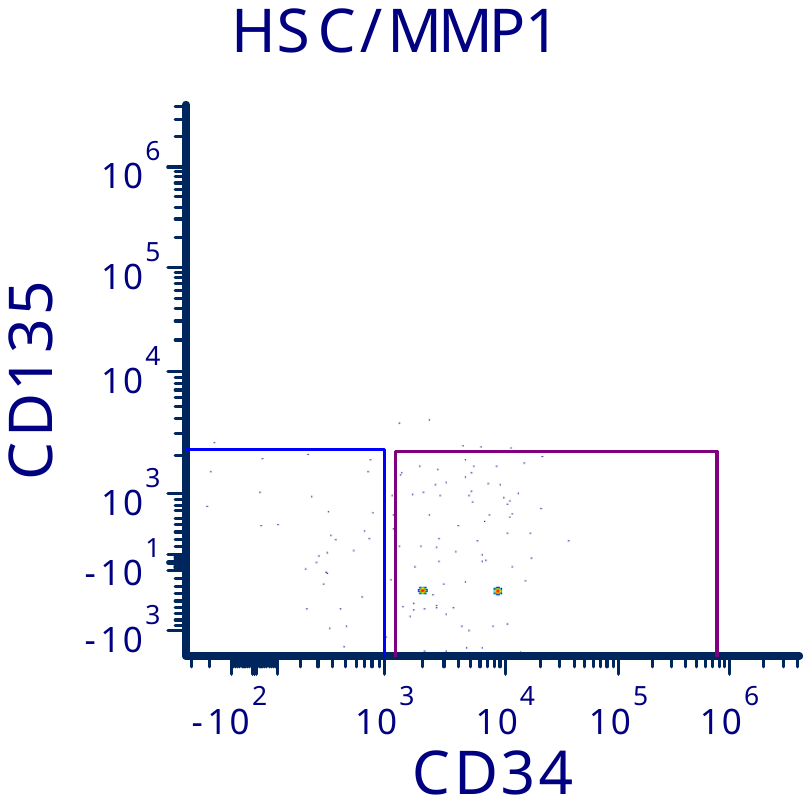

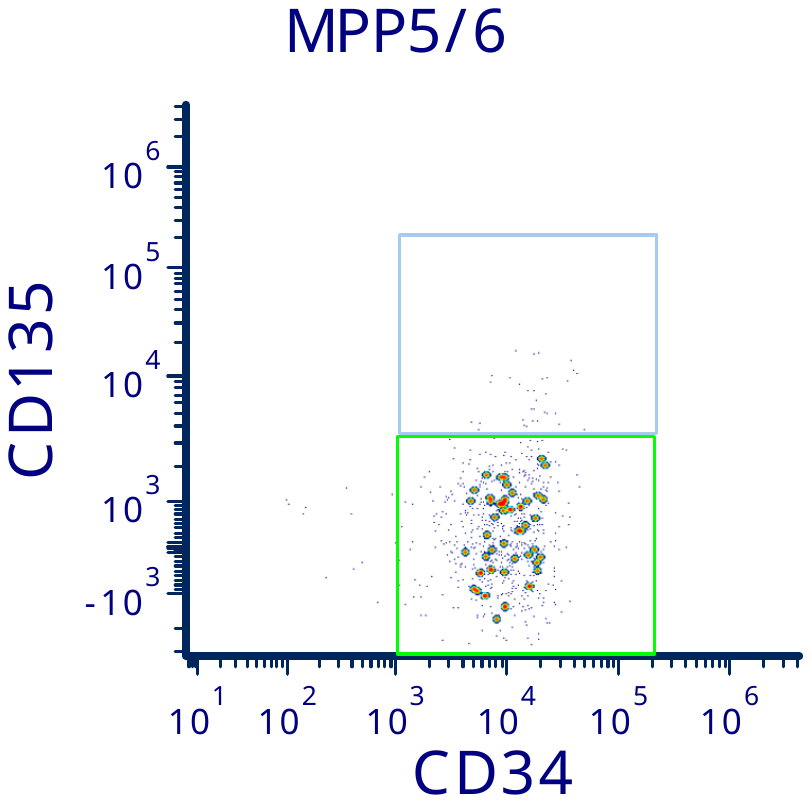

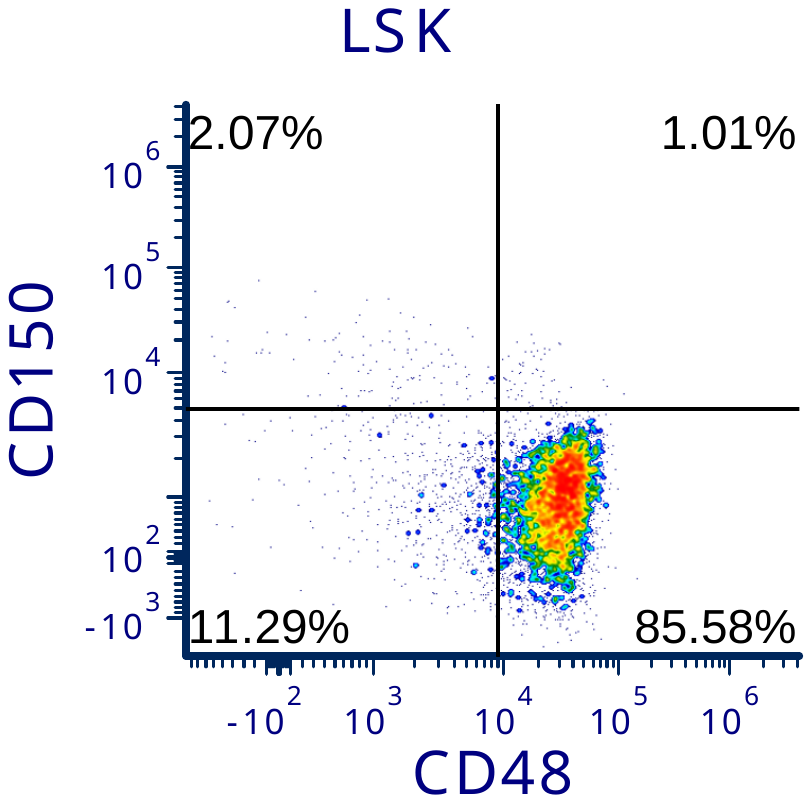

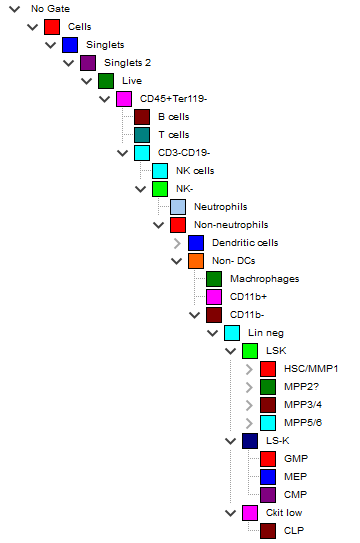

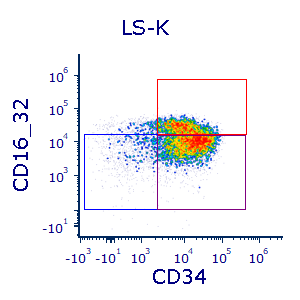

B

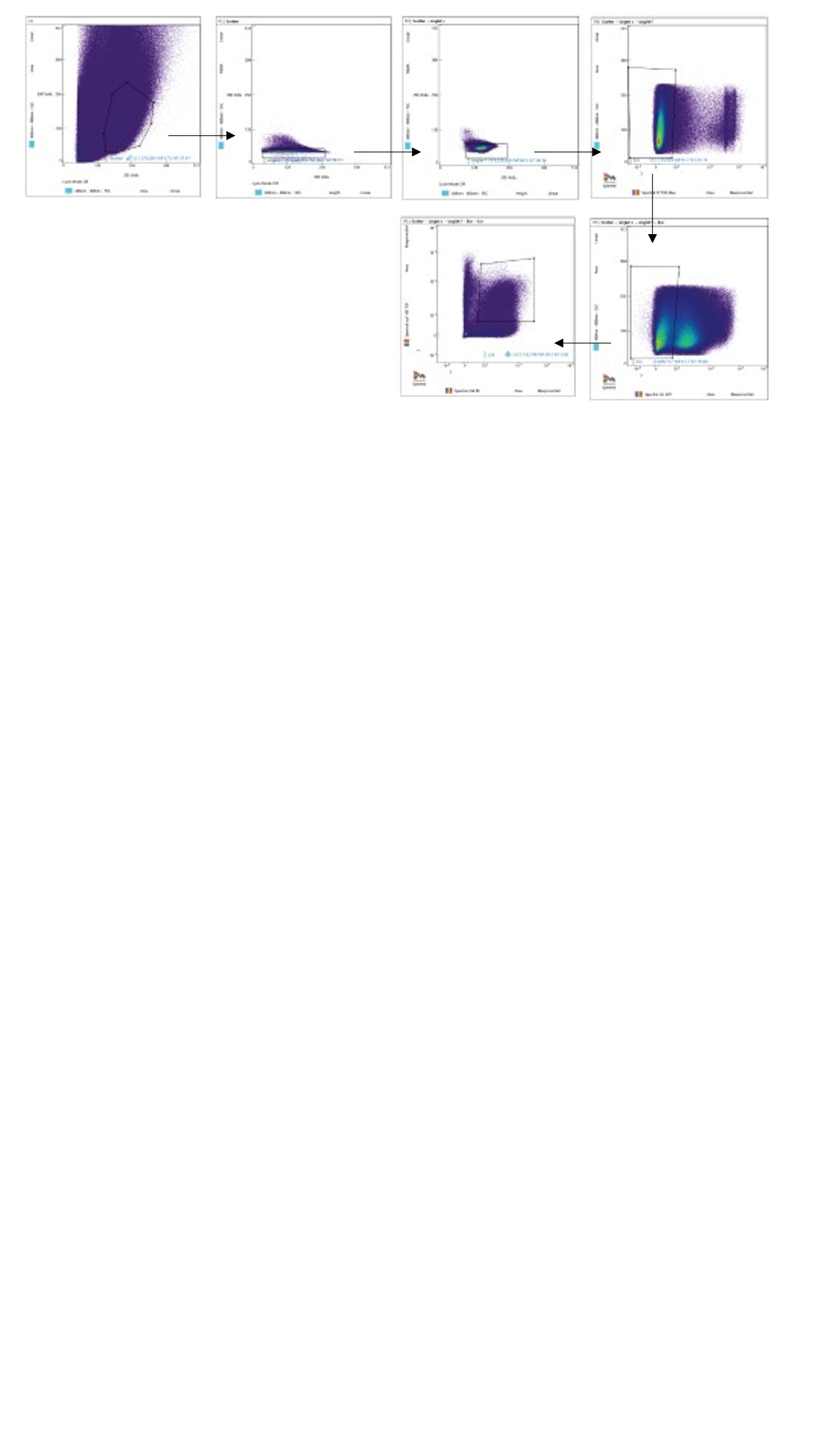

Supplemental Figure 3. Sorting strategy for LSK engraftment experiments.
