## Supplemental Data Table 1 for "*PTPN11* Mutation Clonal Hierarchy in Acute Myeloid Leukemia"

| Hugo_Symbol | HGNC.symbol | Chromosome | Start_Po | End_Pos | Variant_Classification | Variar | Re | Tu | Tu |
|---|---|---|---|---|---|---|---|---|---|
| **Ankrd11** | ANKRD11 | chr8 | 1E+08 | 1E+08 | Missense_Mutation | SNP | C | C | G |
| **Cbl** | CBL | chr9 | 4E+07 | 4E+07 | Missense_Mutation | SNP | A | A | T |
| **Flt3** | FLT3 | chr5 | 1E+08 | 1E+08 | Missense_Mutation | SNP | T | T | C |
| **Npm1** | NPM1 | chr11 | 3E+07 | 3E+07 | Frame_Shift_Ins | INS | - | - | CA |
| **Ptpn11** | PTPN11 | chr5 | 1E+08 | 1E+08 | Missense_Mutation | SNP | C | C | T |
| **Samhd1** | SAMHD1 | chr2 | 2E+08 | 2E+08 | Silent | SNP | G | G | T |
| **Flt3** | FLT3 | chr5 | 1E+08 | 1E+08 | Missense_Mutation | SNP | T | T | C |
| **Nf1** | NF1 | chr11 | 8E+07 | 8E+07 | Missense_Mutation | SNP | C | C | A |
| **Nf1** | NF1 | chr11 | 8E+07 | 8E+07 | Nonsense_Mutation | SNP | C | C | T |
| **Npm1** | NPM1 | chr11 | 3E+07 | 3E+07 | Frame_Shift_Ins | INS | - | - | CA |
| **Ptpn11** | PTPN11 | chr5 | 1E+08 | 1E+08 | Missense_Mutation | SNP | C | C | T |
| **Nf1** | NF1 | chr11 | 8E+07 | 8E+07 | Frame_Shift_Del | DEL | CC | CC | - |
| **Npm1** | NPM1 | chr11 | 3E+07 | 3E+07 | Frame_Shift_Ins | INS | - | - | CA |
| **Ptpn11** | PTPN11 | chr5 | 1E+08 | 1E+08 | Missense_Mutation | SNP | C | C | T |
| **Serpinb3b** | SERPINB3 | chr1 | 1E+08 | 1E+08 | Missense_Mutation | SNP | A | A | G |
| **Npm1** | NPM1 | chr11 | 3E+07 | 3E+07 | Frame_Shift_Ins | INS | - | - | CA |
| **Ptpn11** | PTPN11 | chr5 | 1E+08 | 1E+08 | Missense_Mutation | SNP | C | C | T |
| **Flt3** | FLT3 | chr5 | 1E+08 | 1E+08 | Missense_Mutation | SNP | T | T | C |
| **Nf1** | NF1 | chr11 | 8E+07 | 8E+07 | Nonsense_Mutation | SNP | C | C | T |
| **Npm1** | NPM1 | chr11 | 3E+07 | 3E+07 | Frame_Shift_Ins | INS | - | - | CA |
| **Ptpn11** | PTPN11 | chr5 | 1E+08 | 1E+08 | Missense_Mutation | SNP | C | C | T |
| **Nf1** | NF1 | chr11 | 8E+07 | 8E+07 | Nonsense_Mutation | SNP | G | G | T |
| **Npm1** | NPM1 | chr11 | 3E+07 | 3E+07 | Frame_Shift_Ins | INS | - | - | CA |
| **Ptpn11** | PTPN11 | chr5 | 1E+08 | 1E+08 | Missense_Mutation | SNP | C | C | T |
| **Flt3** | FLT3 | chr5 | 1E+08 | 1E+08 | Missense_Mutation | SNP | T | T | C |
| **Npm1** | NPM1 | chr11 | 3E+07 | 3E+07 | Frame_Shift_Ins | INS | - | - | CA |
| **Ptpn11** | PTPN11 | chr5 | 1E+08 | 1E+08 | Missense_Mutation | SNP | C | C | T |
| **Npm1** | NPM1 | chr11 | 3E+07 | 3E+07 | Frame_Shift_Ins | INS | - | - | CA |
| **Ptpn11** | PTPN11 | chr5 | 1E+08 | 1E+08 | Missense_Mutation | SNP | C | C | T |
| **Vegfa** | VEGFA | chr17 | 5E+07 | 5E+07 | Missense_Mutation | SNP | C | C | T |
| **Erg** | ERG | chr16 | 1E+08 | 1E+08 | Silent | SNP | C | C | A |
| **Flt3** | FLT3 | chr5 | 1E+08 | 1E+08 | Missense_Mutation | SNP | T | T | C |
| **Npm1** | NPM1 | chr11 | 3E+07 | 3E+07 | Frame_Shift_Ins | INS | - | - | CA |
| **Ptpn11** | PTPN11 | chr5 | 1E+08 | 1E+08 | Missense_Mutation | SNP | C | C | T |
| **Ankrd11** | ANKRD11 | chr8 | 1E+08 | 1E+08 | Missense_Mutation | SNP | C | C | G |
| **Flt3** | FLT3 | chr5 | 1E+08 | 1E+08 | Missense_Mutation | SNP | T | T | C |
| **Npm1** | NPM1 | chr11 | 3E+07 | 3E+07 | Frame_Shift_Ins | INS | - | - | CA |
| **Ptpn11** | PTPN11 | chr5 | 1E+08 | 1E+08 | Missense_Mutation | SNP | C | C | T |
| **Rnf43** | RNF43 | chr11 | 9E+07 | 9E+07 | Silent | SNP | A | A | C |
| **Ros1** | ROS1 | chr10 | 5E+07 | 5E+07 | Missense_Mutation | SNP | C | C | A |
| **Samhd1** | SAMHD1 | chr2 | 2E+08 | 2E+08 | Silent | SNP | G | G | T |
| **Flt3** | FLT3 | chr5 | 1E+08 | 1E+08 | Missense_Mutation | SNP | T | T | C |
| **Npm1** | NPM1 | chr11 | 3E+07 | 3E+07 | Frame_Shift_Ins | INS | - | - | CA |
| **Ptpn11** | PTPN11 | chr5 | 1E+08 | 1E+08 | Missense_Mutation | SNP | C | C | T |
| **Npm1** | NPM1 | chr11 | 3E+07 | 3E+07 | Frame_Shift_Ins | INS | - | - | CA |
| **Ptpn11** | PTPN11 | chr5 | 1E+08 | 1E+08 | Missense_Mutation | SNP | C | C | T |
| **Kit** | KIT | chr5 | 8E+07 | 8E+07 | Missense_Mutation | SNP | G | G | T |
| **Lrp1b** | LRP1B | chr2 | 4E+07 | 4E+07 | Silent | SNP | C | C | T |
| **Npm1** | NPM1 | chr11 | 3E+07 | 3E+07 | Frame_Shift_Ins | INS | - | - | CA |

| | | | | | | | | | |
|---|---|---|---|---|---|---|---|---|---|
| **Ptpn11** | PTPN11 | chr5 | 1E+08 | 1E+08 | Missense_Mutation | SNP | C | C | T |
| **Serpinb3b** | SERPINB3 | chr1 | 1E+08 | 1E+08 | Missense_Mutation | SNP | A | A | G |
| **Ankrd11** | ANKRD11 | chr8 | 1E+08 | 1E+08 | Missense_Mutation | SNP | C | C | G |
| **Flt3** | FLT3 | chr5 | 1E+08 | 1E+08 | Missense_Mutation | SNP | T | T | C |
| **Npm1** | NPM1 | chr11 | 3E+07 | 3E+07 | Frame_Shift_Ins | INS | - | - | C/ |
| **Ptpn11** | PTPN11 | chr5 | 1E+08 | 1E+08 | Missense_Mutation | SNP | C | C | T |
| **Samhd1** | SAMHD1 | chr2 | 2E+08 | 2E+08 | Silent | SNP | G | G | T |
| **Ankrd11** | ANKRD11 | chr8 | 1E+08 | 1E+08 | Missense_Mutation | SNP | C | C | G |
| **Flt3** | FLT3 | chr5 | 1E+08 | 1E+08 | Missense_Mutation | SNP | T | T | C |
| **Kmt2d** | KMT2D | chr15 | 1E+08 | 1E+08 | In_Frame_Del | DEL | T( | T( | - |
| **Npm1** | NPM1 | chr11 | 3E+07 | 3E+07 | Frame_Shift_Ins | INS | - | - | C/ |
| **Ptpn11** | PTPN11 | chr5 | 1E+08 | 1E+08 | Missense_Mutation | SNP | C | C | T |
| **Samhd1** | SAMHD1 | chr2 | 2E+08 | 2E+08 | Silent | SNP | G | G | T |
| **Zmym3** | ZMYM3 | chrX | 1E+08 | 1E+08 | Silent | SNP | T | T | C |
| **Kmt2d** | KMT2D | chr15 | 1E+08 | 1E+08 | In_Frame_Del | DEL | T( | T( | - |
| **Nf1** | NF1 | chr11 | 8E+07 | 8E+07 | Nonsense_Mutation | SNP | G | G | T |
| **Npm1** | NPM1 | chr11 | 3E+07 | 3E+07 | Frame_Shift_Ins | INS | - | - | C/ |
| **Ptpn11** | PTPN11 | chr5 | 1E+08 | 1E+08 | Missense_Mutation | SNP | C | C | T |

| dbSNP_F | Tumor_Sample_(do | Matched_Nor | Matc | Ma | t_dept | t_ref_c | t_alt_c | n_dept | n_ref_ | n_alt_c |
|---|---|---|---|---|---|---|---|---|---|---|
| novel | **s_S0165** | s_Mouse-Pool | C | C | 303 | 147 | 156 | 136 | 136 | 0 |
| novel | **s_S0165** | s_Mouse-Pool | A | A | 410 | 399 | 11 | 135 | 135 | 0 |
| novel | **s_S0165** | s_Mouse-Pool | T | T | 397 | 366 | 30 | 145 | 145 | 0 |
| novel | **s_S0165** | s_Mouse-Pool | - | - | 621 | 365 | 256 | 358 | 358 | 0 |
| novel | **s_S0165** | s_Mouse-Pool | C | C | 537 | 268 | 269 | 224 | 224 | 0 |
| novel | **s_S0165** | s_Mouse-Pool | G | G | 584 | 421 | 162 | 218 | 218 | 0 |
| novel | **s_S0200** | s_Mouse-Pool | T | T | 391 | 383 | 8 | 145 | 145 | 0 |
| novel | **s_S0200** | s_Mouse-Pool | C | C | 479 | 325 | 154 | 225 | 225 | 0 |
| novel | **s_S0200** | s_Mouse-Pool | C | C | 482 | 313 | 169 | 213 | 213 | 0 |
| novel | **s_S0200** | s_Mouse-Pool | - | - | 686 | 411 | 275 | 358 | 358 | 0 |
| novel | **s_S0200** | s_Mouse-Pool | C | C | 589 | 309 | 280 | 224 | 224 | 0 |
| novel | **s_S0283** | s_Mouse-Pool | CGCT | CG | 918 | 837 | 81 | 266 | 266 | 0 |
| novel | **s_S0283** | s_Mouse-Pool | - | - | 830 | 503 | 327 | 358 | 358 | 0 |
| novel | **s_S0283** | s_Mouse-Pool | C | C | 617 | 323 | 294 | 224 | 224 | 0 |
| novel | **s_S0283** | s_Mouse-Pool | A | A | 952 | 484 | 467 | 305 | 305 | 0 |
| novel | **s_S0437** | s_Mouse-Pool | - | - | 813 | 510 | 303 | 358 | 358 | 0 |
| novel | **s_S0437** | s_Mouse-Pool | C | C | 736 | 362 | 374 | 224 | 224 | 0 |
| novel | **s_S0518** | s_Mouse-Pool | T | T | 367 | 310 | 57 | 145 | 145 | 0 |
| novel | **s_S0518** | s_Mouse-Pool | C | C | 522 | 498 | 23 | 189 | 189 | 0 |
| novel | **s_S0518** | s_Mouse-Pool | - | - | 714 | 419 | 295 | 358 | 358 | 0 |
| novel | **s_S0518** | s_Mouse-Pool | C | C | 562 | 295 | 267 | 224 | 224 | 0 |
| novel | **s_S0521** | s_Mouse-Pool | G | G | 551 | 65 | 486 | 184 | 183 | 1 |
| novel | **s_S0521** | s_Mouse-Pool | - | - | 750 | 96 | 654 | 358 | 358 | 0 |
| novel | **s_S0521** | s_Mouse-Pool | C | C | 632 | 346 | 286 | 224 | 224 | 0 |
| novel | **s_Y0666 (S0200)** | s_Mouse-Pool | T | T | 423 | 234 | 189 | 145 | 145 | 0 |
| novel | **s_Y0666 (S0200)** | s_Mouse-Pool | - | - | 637 | 420 | 217 | 358 | 358 | 0 |
| novel | **s_Y0666 (S0200)** | s_Mouse-Pool | C | C | 523 | 289 | 234 | 224 | 224 | 0 |
| novel | **s_Y0670 (S0500)** | s_Mouse-Pool | - | - | 641 | 406 | 235 | 358 | 358 | 0 |
| novel | **s_Y0670 (S0500)** | s_Mouse-Pool | C | C | 525 | 274 | 251 | 224 | 224 | 0 |
| novel | **s_Y0670 (S0500)** | s_Mouse-Pool | C | C | 542 | 522 | 19 | 168 | 168 | 0 |
| novel | **s_Y0671 (S0200)** | s_Mouse-Pool | C | C | 407 | 393 | 14 | 135 | 135 | 0 |
| novel | **s_Y0671 (S0200)** | s_Mouse-Pool | T | T | 408 | 236 | 172 | 145 | 145 | 0 |
| novel | **s_Y0671 (S0200)** | s_Mouse-Pool | - | - | 702 | 422 | 280 | 358 | 358 | 0 |
| novel | **s_Y0671 (S0200)** | s_Mouse-Pool | C | C | 484 | 265 | 218 | 224 | 224 | 0 |
| novel | **s_Y0672 (S0165)** | s_Mouse-Pool | C | C | 402 | 203 | 198 | 136 | 136 | 0 |
| novel | **s_Y0672 (S0165)** | s_Mouse-Pool | T | T | 448 | 238 | 210 | 145 | 145 | 0 |
| novel | **s_Y0672 (S0165)** | s_Mouse-Pool | - | - | 621 | 405 | 216 | 358 | 358 | 0 |
| novel | **s_Y0672 (S0165)** | s_Mouse-Pool | C | C | 591 | 333 | 258 | 224 | 224 | 0 |
| novel | **s_Y0672 (S0165)** | s_Mouse-Pool | A | A | 490 | 476 | 13 | 168 | 168 | 0 |
| novel | **s_Y0672 (S0165)** | s_Mouse-Pool | C | C | 642 | 617 | 25 | 251 | 251 | 0 |
| novel | **s_Y0672 (S0165)** | s_Mouse-Pool | G | G | 574 | 340 | 234 | 218 | 218 | 0 |
| novel | **s_Y0674 (S0200)** | s_Mouse-Pool | T | T | 423 | 217 | 206 | 145 | 145 | 0 |
| novel | **s_Y0674 (S0200)** | s_Mouse-Pool | - | - | 628 | 405 | 223 | 358 | 358 | 0 |
| novel | **s_Y0674 (S0200)** | s_Mouse-Pool | C | C | 531 | 299 | 232 | 224 | 224 | 0 |
| novel | **s_Y0677 (S0437)** | s_Mouse-Pool | - | - | 630 | 392 | 238 | 358 | 358 | 0 |
| novel | **s_Y0677 (S0437)** | s_Mouse-Pool | C | C | 569 | 322 | 246 | 224 | 224 | 0 |
| novel | **s_Y0679 (S0283)** | s_Mouse-Pool | G | G | 587 | 383 | 204 | 196 | 196 | 0 |
| novel | **s_Y0679 (S0283)** | s_Mouse-Pool | C | C | 585 | 448 | 136 | 211 | 211 | 0 |
| novel | **s_Y0679 (S0283)** | s_Mouse-Pool | - | - | 693 | 449 | 244 | 358 | 358 | 0 |

| | | | | | | | | | | |
|---|---|---|---|---|---|---|---|---|---|---|
| novel | **s_Y0679 (S0283)** | s_Mouse-Pool | C | C | 606 | 366 | 240 | 224 | 224 | 0 |
| novel | **s_Y0679 (S0283)** | s_Mouse-Pool | A | A | 930 | 536 | 393 | 305 | 305 | 0 |
| novel | **s_Y0887 (S0165)** | s_Mouse-Pool | C | C | 400 | 225 | 175 | 136 | 136 | 0 |
| novel | **s_Y0887 (S0165)** | s_Mouse-Pool | T | T | 395 | 246 | 149 | 145 | 145 | 0 |
| novel | **s_Y0887 (S0165)** | s_Mouse-Pool | - | - | 699 | 481 | 218 | 358 | 358 | 0 |
| novel | **s_Y0887 (S0165)** | s_Mouse-Pool | C | C | 561 | 333 | 227 | 224 | 224 | 0 |
| novel | **s_Y0887 (S0165)** | s_Mouse-Pool | G | G | 565 | 335 | 229 | 218 | 218 | 0 |
| novel | **s_Y0888 (S0165)** | s_Mouse-Pool | C | C | 324 | 205 | 119 | 136 | 136 | 0 |
| novel | **s_Y0888 (S0165)** | s_Mouse-Pool | T | T | 365 | 227 | 138 | 145 | 145 | 0 |
| novel | **s_Y0888 (S0165)** | s_Mouse-Pool | TGCT | TG( | 681 | 619 | 62 | 246 | 246 | 0 |
| novel | **s_Y0888 (S0165)** | s_Mouse-Pool | - | - | 622 | 453 | 169 | 358 | 358 | 0 |
| novel | **s_Y0888 (S0165)** | s_Mouse-Pool | C | C | 563 | 363 | 200 | 224 | 224 | 0 |
| novel | **s_Y0888 (S0165)** | s_Mouse-Pool | G | G | 551 | 361 | 190 | 218 | 218 | 0 |
| novel | **s_Y0888 (S0165)** | s_Mouse-Pool | T | T | 321 | 304 | 17 | 101 | 101 | 0 |
| novel | **s_Y0909 (S0521)** | s_Mouse-Pool | TGCT | TG( | 839 | 742 | 97 | 246 | 246 | 0 |
| novel | **s_Y0909 (S0521)** | s_Mouse-Pool | G | G | 504 | 108 | 396 | 184 | 183 | 1 |
| novel | **s_Y0909 (S0521)** | s_Mouse-Pool | - | - | 764 | 264 | 500 | 358 | 358 | 0 |
| novel | **s_Y0909 (S0521)** | s_Mouse-Pool | C | C | 639 | 416 | 222 | 224 | 224 | 0 |

| t_var_freq | n_var_freq | FILTER | Cohort.N | Cohort.PCT | HGVSc | HGVSp |
|---|---|---|---|---|---|---|
| **0.51485149** | 0 | PASS | 4 | 0.17391304 | c.6868G>C | p.Val2290Leu |
| **0.02682927** | 0 | PASS | 1 | 0.04347826 | c.1105T>A | p.Tyr369Asn |
| **0.07556675** | 0 | PASS | 9 | 0.39130435 | c.2525A>G | p.Asp842Gly |
| **0.41223833** | 0 | PASS | 16 | 0.69565217 | c.854_857dup | p.Trp286Cysfs |
| **0.5009311** | 0 | PASS | 16 | 0.69565217 | c.226G>A | p.Glu76Lys |
| **0.27739726** | 0 | PASS | 4 | 0.17391304 | c.318C>A | p.Ala106= |
| **0.02046036** | 0 | PASS | 9 | 0.39130435 | c.2525A>G | p.Asp842Gly |
| **0.32150313** | 0 | PASS | 1 | 0.04347826 | c.4345C>A | p.Gln1449Lys |
| **0.35062241** | 0 | PASS | 1 | 0.04347826 | c.4432C>T | p.Arg1478Ter |
| **0.40087464** | 0 | PASS | 16 | 0.69565217 | c.854_857dup | p.Trp286Cysfs |
| **0.475382** | 0 | PASS | 16 | 0.69565217 | c.226G>A | p.Glu76Lys |
| **0.08823529** | 0 | PASS | 1 | 0.04347826 | c.4964_4973d | p.Arg1655Gln1 |
| **0.3939759** | 0 | PASS | 16 | 0.69565217 | c.854_857dup | p.Trp286Cysfs |
| **0.47649919** | 0 | PASS | 16 | 0.69565217 | c.226G>A | p.Glu76Lys |
| **0.49054622** | 0 | PASS | 2 | 0.08695652 | c.992T>C | p.Phe331Ser |
| **0.37269373** | 0 | PASS | 16 | 0.69565217 | c.854_857dup | p.Trp286Cysfs |
| **0.50815217** | 0 | PASS | 16 | 0.69565217 | c.226G>A | p.Glu76Lys |
| **0.15531335** | 0 | PASS | 9 | 0.39130435 | c.2525A>G | p.Asp842Gly |
| **0.0440613** | 0 | PASS | 1 | 0.04347826 | c.6811C>T | p.Arg2271Ter |
| **0.41316527** | 0 | PASS | 16 | 0.69565217 | c.854_857dup | p.Trp286Cysfs |
| **0.47508897** | 0 | PASS | 16 | 0.69565217 | c.226G>A | p.Glu76Lys |
| **0.88203267** | 0.00543478 | PASS | 2 | 0.08695652 | c.3718G>T | p.Glu1240Ter |
| **0.872** | 0 | PASS | 16 | 0.69565217 | c.854_857dup | p.Trp286Cysfs |
| **0.45253165** | 0 | PASS | 16 | 0.69565217 | c.226G>A | p.Glu76Lys |
| **0.44680851** | 0 | PASS | 9 | 0.39130435 | c.2525A>G | p.Asp842Gly |
| **0.34065934** | 0 | PASS | 16 | 0.69565217 | c.854_857dup | p.Trp286Cysfs |
| **0.44741874** | 0 | PASS | 16 | 0.69565217 | c.226G>A | p.Glu76Lys |
| **0.36661466** | 0 | PASS | 16 | 0.69565217 | c.854_857dup | p.Trp286Cysfs |
| **0.47809524** | 0 | PASS | 16 | 0.69565217 | c.226G>A | p.Glu76Lys |
| **0.03505535** | 0 | PASS | 1 | 0.04347826 | c.733G>A | p.Glu245Lys |
| **0.03439803** | 0 | PASS | 1 | 0.04347826 | c.744G>T | p.Thr248= |
| **0.42156863** | 0 | PASS | 9 | 0.39130435 | c.2525A>G | p.Asp842Gly |
| **0.3988604** | 0 | PASS | 16 | 0.69565217 | c.854_857dup | p.Trp286Cysfs |
| **0.45041322** | 0 | PASS | 16 | 0.69565217 | c.226G>A | p.Glu76Lys |
| **0.49253731** | 0 | PASS | 4 | 0.17391304 | c.6868G>C | p.Val2290Leu |
| **0.46875** | 0 | PASS | 9 | 0.39130435 | c.2525A>G | p.Asp842Gly |
| **0.34782609** | 0 | PASS | 16 | 0.69565217 | c.854_857dup | p.Trp286Cysfs |
| **0.43654822** | 0 | PASS | 16 | 0.69565217 | c.226G>A | p.Glu76Lys |
| **0.02653061** | 0 | PASS | 1 | 0.04347826 | c.1494A>C | p.Arg498= |
| **0.03894081** | 0 | PASS | 1 | 0.04347826 | c.4195G>T | p.Ala1399Ser |
| **0.40766551** | 0 | PASS | 4 | 0.17391304 | c.318C>A | p.Ala106= |
| **0.48699764** | 0 | PASS | 9 | 0.39130435 | c.2525A>G | p.Asp842Gly |
| **0.35509554** | 0 | PASS | 16 | 0.69565217 | c.854_857dup | p.Trp286Cysfs |
| **0.43691149** | 0 | PASS | 16 | 0.69565217 | c.226G>A | p.Glu76Lys |
| **0.37777778** | 0 | PASS | 16 | 0.69565217 | c.854_857dup | p.Trp286Cysfs |
| **0.43233743** | 0 | PASS | 16 | 0.69565217 | c.226G>A | p.Glu76Lys |
| **0.34752981** | 0 | PASS | 1 | 0.04347826 | c.2452G>T | p.Asp818Tyr |
| **0.23247863** | 0 | PASS | 1 | 0.04347826 | c.2316G>A | p.Thr772= |
| **0.35209235** | 0 | PASS | 16 | 0.69565217 | c.854_857dup | p.Trp286Cysfs |

| | | | | | | |
|---|---|---|---|---|---|---|
| **0.3960396** | 0 | PASS | 16 | 0.69565217 | c.226G>A | p.Glu76Lys |
| **0.42258065** | 0 | PASS | 2 | 0.08695652 | c.992T>C | p.Phe331Ser |
| **0.4375** | 0 | PASS | 4 | 0.17391304 | c.6868G>C | p.Val2290Leu |
| **0.37721519** | 0 | PASS | 9 | 0.39130435 | c.2525A>G | p.Asp842Gly |
| **0.31187411** | 0 | PASS | 16 | 0.69565217 | c.854_857dup | p.Trp286Cysfs |
| **0.40463458** | 0 | PASS | 16 | 0.69565217 | c.226G>A | p.Glu76Lys |
| **0.40530973** | 0 | PASS | 4 | 0.17391304 | c.318C>A | p.Ala106= |
| **0.36728395** | 0 | PASS | 4 | 0.17391304 | c.6868G>C | p.Val2290Leu |
| **0.37808219** | 0 | PASS | 9 | 0.39130435 | c.2525A>G | p.Asp842Gly |
| **0.09104258** | 0 | PASS | 2 | 0.08695652 | c.8340_8375d | p.Gln2786_Gl |
| **0.27170418** | 0 | PASS | 16 | 0.69565217 | c.854_857dup | p.Trp286Cysfs |
| **0.35523979** | 0 | PASS | 16 | 0.69565217 | c.226G>A | p.Glu76Lys |
| **0.34482759** | 0 | PASS | 4 | 0.17391304 | c.318C>A | p.Ala106= |
| **0.0529595** | 0 | PASS | 1 | 0.04347826 | c.228A>G | p.Gly76= |
| **0.11561383** | 0 | PASS | 2 | 0.08695652 | c.8340_8375d | p.Gln2786_Gl |
| **0.78571429** | 0.00543478 | PASS | 2 | 0.08695652 | c.3718G>T | p.Glu1240Ter |
| **0.65445026** | 0 | PASS | 16 | 0.69565217 | c.854_857dup | p.Trp286Cysfs |
| **0.34741784** | 0 | PASS | 16 | 0.69565217 | c.226G>A | p.Glu76Lys |

| HGVSp_Short | Transcript_ID | Exon_Number | all_effects | Allele | Gene | Feature |
|---|---|---|---|---|---|---|
| p.V2290L | ENSMUST000( | 9/13 | Ankrd11,misse | G | ENSMUSG000 | ENSMUST000( |
| p.Y369N | ENSMUST000( | 8/16 | Cbl,missense_ | T | ENSMUSG000 | ENSMUST000( |
| p.D842G | ENSMUST000( | 20/24 | Flt3,missense_ | C | ENSMUSG000 | ENSMUST000( |
| p.W286Cfs*27 | ENSMUST000( | 11/11 | Npm1,framesh | CAGA | ENSMUSG000 | ENSMUST000( |
| p.E76K | ENSMUST000( | 3/16 | Ptpn11,misser | T | ENSMUSG000 | ENSMUST000( |
| p.A106= | ENSMUST000( | 2/16 | Samhd1,synor | T | ENSMUSG000 | ENSMUST000( |
| p.D842G | ENSMUST000( | 20/24 | Flt3,missense_ | C | ENSMUSG000 | ENSMUST000( |
| p.Q1449K | ENSMUST000( | 33/58 | Nf1,missense_ | A | ENSMUSG000 | ENSMUST000( |
| p.R1478* | ENSMUST000( | 33/58 | Nf1,stop_gain | T | ENSMUSG000 | ENSMUST000( |
| p.W286Cfs*27 | ENSMUST000( | 11/11 | Npm1,framesh | CAGA | ENSMUSG000 | ENSMUST000( |
| p.E76K | ENSMUST000( | 3/16 | Ptpn11,misser | T | ENSMUSG000 | ENSMUST000( |
| p.R1655Qfs*4 | ENSMUST000( | 37/58 | Nf1,frameshift | - | ENSMUSG000 | ENSMUST000( |
| p.W286Cfs*27 | ENSMUST000( | 11/11 | Npm1,framesh | CAGA | ENSMUSG000 | ENSMUST000( |
| p.E76K | ENSMUST000( | 3/16 | Ptpn11,misser | T | ENSMUSG000 | ENSMUST000( |
| p.F331S | ENSMUST000( | 8/8 | Serpinb3b,mis | G | ENSMUSG000 | ENSMUST000( |
| p.W286Cfs*27 | ENSMUST000( | 11/11 | Npm1,framesh | CAGA | ENSMUSG000 | ENSMUST000( |
| p.E76K | ENSMUST000( | 3/16 | Ptpn11,misser | T | ENSMUSG000 | ENSMUST000( |
| p.D842G | ENSMUST000( | 20/24 | Flt3,missense_ | C | ENSMUSG000 | ENSMUST000( |
| p.R2271* | ENSMUST000( | 45/58 | Nf1,stop_gain | T | ENSMUSG000 | ENSMUST000( |
| p.W286Cfs*27 | ENSMUST000( | 11/11 | Npm1,framesh | CAGA | ENSMUSG000 | ENSMUST000( |
| p.E76K | ENSMUST000( | 3/16 | Ptpn11,misser | T | ENSMUSG000 | ENSMUST000( |
| p.E1240* | ENSMUST000( | 28/58 | Nf1,stop_gain | T | ENSMUSG000 | ENSMUST000( |
| p.W286Cfs*27 | ENSMUST000( | 11/11 | Npm1,framesh | CAGA | ENSMUSG000 | ENSMUST000( |
| p.E76K | ENSMUST000( | 3/16 | Ptpn11,misser | T | ENSMUSG000 | ENSMUST000( |
| p.D842G | ENSMUST000( | 20/24 | Flt3,missense_ | C | ENSMUSG000 | ENSMUST000( |
| p.W286Cfs*27 | ENSMUST000( | 11/11 | Npm1,framesh | CAGA | ENSMUSG000 | ENSMUST000( |
| p.E76K | ENSMUST000( | 3/16 | Ptpn11,misser | T | ENSMUSG000 | ENSMUST000( |
| p.W286Cfs*27 | ENSMUST000( | 11/11 | Npm1,framesh | CAGA | ENSMUSG000 | ENSMUST000( |
| p.E76K | ENSMUST000( | 3/16 | Ptpn11,misser | T | ENSMUSG000 | ENSMUST000( |
| p.E245K | ENSMUST000( | 3/8 | Vegfa,missens | T | ENSMUSG000 | ENSMUST000( |
| p.T248= | ENSMUST000( | 7/11 | Erg,synonymo | A | ENSMUSG000 | ENSMUST000( |
| p.D842G | ENSMUST000( | 20/24 | Flt3,missense_ | C | ENSMUSG000 | ENSMUST000( |
| p.W286Cfs*27 | ENSMUST000( | 11/11 | Npm1,framesh | CAGA | ENSMUSG000 | ENSMUST000( |
| p.E76K | ENSMUST000( | 3/16 | Ptpn11,misser | T | ENSMUSG000 | ENSMUST000( |
| p.V2290L | ENSMUST000( | 9/13 | Ankrd11,misse | G | ENSMUSG000 | ENSMUST000( |
| p.D842G | ENSMUST000( | 20/24 | Flt3,missense_ | C | ENSMUSG000 | ENSMUST000( |
| p.W286Cfs*27 | ENSMUST000( | 11/11 | Npm1,framesh | CAGA | ENSMUSG000 | ENSMUST000( |
| p.E76K | ENSMUST000( | 3/16 | Ptpn11,misser | T | ENSMUSG000 | ENSMUST000( |
| p.R498= | ENSMUST000( | 9/10 | Rnf43,synonyr | C | ENSMUSG000 | ENSMUST000( |
| p.A1399S | ENSMUST000( | 27/44 | Ros1,missense | A | ENSMUSG000 | ENSMUST000( |
| p.A106= | ENSMUST000( | 2/16 | Samhd1,synor | T | ENSMUSG000 | ENSMUST000( |
| p.D842G | ENSMUST000( | 20/24 | Flt3,missense_ | C | ENSMUSG000 | ENSMUST000( |
| p.W286Cfs*27 | ENSMUST000( | 11/11 | Npm1,framesh | CAGA | ENSMUSG000 | ENSMUST000( |
| p.E76K | ENSMUST000( | 3/16 | Ptpn11,misser | T | ENSMUSG000 | ENSMUST000( |
| p.W286Cfs*27 | ENSMUST000( | 11/11 | Npm1,framesh | CAGA | ENSMUSG000 | ENSMUST000( |
| p.E76K | ENSMUST000( | 3/16 | Ptpn11,misser | T | ENSMUSG000 | ENSMUST000( |
| p.D818Y | ENSMUST000( | 17/21 | Kit,missense_v | T | ENSMUSG000 | ENSMUST000( |
| p.T772= | ENSMUST000( | 14/91 | Lrp1b,synonyr | T | ENSMUSG000 | ENSMUST000( |
| p.W286Cfs*27 | ENSMUST000( | 11/11 | Npm1,framesh | CAGA | ENSMUSG000 | ENSMUST000( |

| | | | | | | |
|---|---|---|---|---|---|---|
| p.E76K | ENSMUST000( | 3/16 | Ptpn11,misser | T | ENSMUSG000 | ENSMUST000( |
| p.F331S | ENSMUST000( | 8/8 | Serpinb3b,mis | G | ENSMUSG000 | ENSMUST000( |
| p.V2290L | ENSMUST000( | 9/13 | Ankrd11,misse | G | ENSMUSG000 | ENSMUST000( |
| p.D842G | ENSMUST000( | 20/24 | Flt3,missense_ | C | ENSMUSG000 | ENSMUST000( |
| p.W286Cfs*27 | ENSMUST000( | 11/11 | Npm1,framesh | CAGA | ENSMUSG000 | ENSMUST000( |
| p.E76K | ENSMUST000( | 3/16 | Ptpn11,misser | T | ENSMUSG000 | ENSMUST000( |
| p.A106= | ENSMUST000( | 2/16 | Samhd1,synor | T | ENSMUSG000 | ENSMUST000( |
| p.V2290L | ENSMUST000( | 9/13 | Ankrd11,misse | G | ENSMUSG000 | ENSMUST000( |
| p.D842G | ENSMUST000( | 20/24 | Flt3,missense_ | C | ENSMUSG000 | ENSMUST000( |
| p.Q2786_Q27 | ENSMUST000( | 35/55 | Kmt2d,inframe | - | ENSMUSG000 | ENSMUST000( |
| p.W286Cfs*27 | ENSMUST000( | 11/11 | Npm1,framesh | CAGA | ENSMUSG000 | ENSMUST000( |
| p.E76K | ENSMUST000( | 3/16 | Ptpn11,misser | T | ENSMUSG000 | ENSMUST000( |
| p.A106= | ENSMUST000( | 2/16 | Samhd1,synor | T | ENSMUSG000 | ENSMUST000( |
| p.G76= | ENSMUST000( | 2/25 | Zmym3,synon | C | ENSMUSG000 | ENSMUST000( |
| p.Q2786_Q27 | ENSMUST000( | 35/55 | Kmt2d,inframe | - | ENSMUSG000 | ENSMUST000( |
| p.E1240* | ENSMUST000( | 28/58 | Nf1,stop_gain | T | ENSMUSG000 | ENSMUST000( |
| p.W286Cfs*27 | ENSMUST000( | 11/11 | Npm1,framesh | CAGA | ENSMUSG000 | ENSMUST000( |
| p.E76K | ENSMUST000( | 3/16 | Ptpn11,misser | T | ENSMUSG000 | ENSMUST000( |

| Feature_type | Consequence | cDNA_position | CDS_position | Protein_positi | Amino_acids | Codons |
|---|---|---|---|---|---|---|
| Transcript | missense_vari | 7062/8455 | 6868/7932 | 2290/2643 | V/L | Gtt/Ctt |
| Transcript | missense_vari | 1266/11372 | 1105/2742 | 369/913 | Y/N | Tac/Aac |
| Transcript | missense_vari | 2749/3657 | 2525/3003 | 842/1000 | D/G | gAc/gGc |
| Transcript | frameshift_va | 1084-1085/16 | 857-858/879 | 286/292 | W/CLX | tgg/tgTCTGg |
| Transcript | missense_vari | 340/5535 | 226/1794 | 76/597 | E/K | Gaa/Aaa |
| Transcript | synonymous_ | 401/3926 | 318/1977 | 106/658 | A | gcC/gcA |
| Transcript | missense_vari | 2749/3657 | 2525/3003 | 842/1000 | D/G | gAc/gGc |
| Transcript | missense_vari | 4520/11917 | 4345/8526 | 1449/2841 | Q/K | Cag/Aag |
| Transcript | stop_gained | 4607/11917 | 4432/8526 | 1478/2841 | R/* | Cga/Tga |
| Transcript | frameshift_va | 1084-1085/16 | 857-858/879 | 286/292 | W/CLX | tgg/tgTCTGg |
| Transcript | missense_vari | 340/5535 | 226/1794 | 76/597 | E/K | Gaa/Aaa |
| Transcript | frameshift_va | 5138-5147/11 | 4963-4972/85 | 1655-1658/28 | RFKT/X | CGCTTTAAAA |
| Transcript | frameshift_va | 1084-1085/16 | 857-858/879 | 286/292 | W/CLX | tgg/tgTCTGg |
| Transcript | missense_vari | 340/5535 | 226/1794 | 76/597 | E/K | Gaa/Aaa |
| Transcript | missense_vari | 1052/1653 | 992/1164 | 331/387 | F/S | tTt/tCt |
| Transcript | frameshift_va | 1084-1085/16 | 857-858/879 | 286/292 | W/CLX | tgg/tgTCTGg |
| Transcript | missense_vari | 340/5535 | 226/1794 | 76/597 | E/K | Gaa/Aaa |
| Transcript | missense_vari | 2749/3657 | 2525/3003 | 842/1000 | D/G | gAc/gGc |
| Transcript | stop_gained | 6986/11917 | 6811/8526 | 2271/2841 | R/* | Cga/Tga |
| Transcript | frameshift_va | 1084-1085/16 | 857-858/879 | 286/292 | W/CLX | tgg/tgTCTGg |
| Transcript | missense_vari | 340/5535 | 226/1794 | 76/597 | E/K | Gaa/Aaa |
| Transcript | stop_gained | 3893/11917 | 3718/8526 | 1240/2841 | E/* | Gag/Tag |
| Transcript | frameshift_va | 1084-1085/16 | 857-858/879 | 286/292 | W/CLX | tgg/tgTCTGg |
| Transcript | missense_vari | 340/5535 | 226/1794 | 76/597 | E/K | Gaa/Aaa |
| Transcript | missense_vari | 2749/3657 | 2525/3003 | 842/1000 | D/G | gAc/gGc |
| Transcript | frameshift_va | 1084-1085/16 | 857-858/879 | 286/292 | W/CLX | tgg/tgTCTGg |
| Transcript | missense_vari | 340/5535 | 226/1794 | 76/597 | E/K | Gaa/Aaa |
| Transcript | frameshift_va | 1084-1085/16 | 857-858/879 | 286/292 | W/CLX | tgg/tgTCTGg |
| Transcript | missense_vari | 340/5535 | 226/1794 | 76/597 | E/K | Gaa/Aaa |
| Transcript | missense_vari | 1212/1973 | 733/1179 | 245/392 | E/K | Gag/Aag |
| Transcript | synonymous_ | 895/3249 | 744/1461 | 248/486 | T | acG/acT |
| Transcript | missense_vari | 2749/3657 | 2525/3003 | 842/1000 | D/G | gAc/gGc |
| Transcript | frameshift_va | 1084-1085/16 | 857-858/879 | 286/292 | W/CLX | tgg/tgTCTGg |
| Transcript | missense_vari | 340/5535 | 226/1794 | 76/597 | E/K | Gaa/Aaa |
| Transcript | missense_vari | 7062/8455 | 6868/7932 | 2290/2643 | V/L | Gtt/Ctt |
| Transcript | missense_vari | 2749/3657 | 2525/3003 | 842/1000 | D/G | gAc/gGc |
| Transcript | frameshift_va | 1084-1085/16 | 857-858/879 | 286/292 | W/CLX | tgg/tgTCTGg |
| Transcript | missense_vari | 340/5535 | 226/1794 | 76/597 | E/K | Gaa/Aaa |
| Transcript | synonymous_ | 2110/4310 | 1494/2355 | 498/784 | R | cgA/cgC |
| Transcript | missense_vari | 4483/7401 | 4195/7023 | 1399/2340 | A/S | Gcc/Tcc |
| Transcript | synonymous_ | 401/3926 | 318/1977 | 106/658 | A | gcC/gcA |
| Transcript | missense_vari | 2749/3657 | 2525/3003 | 842/1000 | D/G | gAc/gGc |
| Transcript | frameshift_va | 1084-1085/16 | 857-858/879 | 286/292 | W/CLX | tgg/tgTCTGg |
| Transcript | missense_vari | 340/5535 | 226/1794 | 76/597 | E/K | Gaa/Aaa |
| Transcript | frameshift_va | 1084-1085/16 | 857-858/879 | 286/292 | W/CLX | tgg/tgTCTGg |
| Transcript | missense_vari | 340/5535 | 226/1794 | 76/597 | E/K | Gaa/Aaa |
| Transcript | missense_vari | 2549/5214 | 2452/2940 | 818/979 | D/Y | Gac/Tac |
| Transcript | synonymous_ | 3124/16294 | 2316/13800 | 772/4599 | T | acG/acA |
| Transcript | frameshift_va | 1084-1085/16 | 857-858/879 | 286/292 | W/CLX | tgg/tgTCTGg |

| | | | | | | |
|---|---|---|---|---|---|---|
| Transcript | missense_vari | 340/5535 | 226/1794 | 76/597 | E/K | Gaa/Aaa |
| Transcript | missense_vari | 1052/1653 | 992/1164 | 331/387 | F/S | tTt/tCt |
| Transcript | missense_vari | 7062/8455 | 6868/7932 | 2290/2643 | V/L | Gtt/Ctt |
| Transcript | missense_vari | 2749/3657 | 2525/3003 | 842/1000 | D/G | gAc/gGc |
| Transcript | frameshift_va | 1084-1085/16 | 857-858/879 | 286/292 | W/CLX | tgg/tgTCTGg |
| Transcript | missense_vari | 340/5535 | 226/1794 | 76/597 | E/K | Gaa/Aaa |
| Transcript | synonymous_ | 401/3926 | 318/1977 | 106/658 | A | gcC/gcA |
| Transcript | missense_vari | 7062/8455 | 6868/7932 | 2290/2643 | V/L | Gtt/Ctt |
| Transcript | missense_vari | 2749/3657 | 2525/3003 | 842/1000 | D/G | gAc/gGc |
| Transcript | inframe_delet | 8568-8603/19 | 8340-8375/16 | 2780-2792/55 | QQQQQQQQ | caACAACAGC |
| Transcript | frameshift_va | 1084-1085/16 | 857-858/879 | 286/292 | W/CLX | tgg/tgTCTGg |
| Transcript | missense_vari | 340/5535 | 226/1794 | 76/597 | E/K | Gaa/Aaa |
| Transcript | synonymous_ | 401/3926 | 318/1977 | 106/658 | A | gcC/gcA |
| Transcript | synonymous_ | 945/6057 | 228/4113 | 76/1370 | G | ggA/ggG |
| Transcript | inframe_delet | 8568-8603/19 | 8340-8375/16 | 2780-2792/55 | QQQQQQQQ | caACAACAGC |
| Transcript | stop_gained | 3893/11917 | 3718/8526 | 1240/2841 | E/* | Gag/Tag |
| Transcript | frameshift_va | 1084-1085/16 | 857-858/879 | 286/292 | W/CLX | tgg/tgTCTGg |
| Transcript | missense_vari | 340/5535 | 226/1794 | 76/597 | E/K | Gaa/Aaa |

| STRAND_VEP | SYMBOL | SYMBOL_SOU | BIOTYPE | CANONICAL | CCDS | ENSP |
|---|---|---|---|---|---|---|
| -1 | Ankrd11 | MGI | protein_codin | YES | CCDS40507.2 | ENSMUSP0000 |
| -1 | Cbl | MGI | protein_codin | YES | CCDS40598.1 | ENSMUSP0000 |
| -1 | Flt3 | MGI | protein_codin | YES | CCDS39400.1 | ENSMUSP0000 |
| -1 | Npm1 | MGI | protein_codin | YES | CCDS24532.1 | ENSMUSP0000 |
| -1 | Ptpn11 | MGI | protein_codin | YES | CCDS39247.1 | ENSMUSP0000 |
| -1 | Samhd1 | MGI | protein_codin | YES | CCDS16973.2 | ENSMUSP0000 |
| -1 | Flt3 | MGI | protein_codin | YES | CCDS39400.1 | ENSMUSP0000 |
| 1 | Nf1 | MGI | protein_codin | YES | CCDS25119.1 | ENSMUSP0000 |
| 1 | Nf1 | MGI | protein_codin | YES | CCDS25119.1 | ENSMUSP0000 |
| -1 | Npm1 | MGI | protein_codin | YES | CCDS24532.1 | ENSMUSP0000 |
| -1 | Ptpn11 | MGI | protein_codin | YES | CCDS39247.1 | ENSMUSP0000 |
| 1 | Nf1 | MGI | protein_codin | YES | CCDS25119.1 | ENSMUSP0000 |
| -1 | Npm1 | MGI | protein_codin | YES | CCDS24532.1 | ENSMUSP0000 |
| -1 | Ptpn11 | MGI | protein_codin | YES | CCDS39247.1 | ENSMUSP0000 |
| -1 | Serpinb3b | MGI | protein_codin | YES | CCDS15216.1 | ENSMUSP0000 |
| -1 | Npm1 | MGI | protein_codin | YES | CCDS24532.1 | ENSMUSP0000 |
| -1 | Ptpn11 | MGI | protein_codin | YES | CCDS39247.1 | ENSMUSP0000 |
| -1 | Flt3 | MGI | protein_codin | YES | CCDS39400.1 | ENSMUSP0000 |
| 1 | Nf1 | MGI | protein_codin | YES | CCDS25119.1 | ENSMUSP0000 |
| -1 | Npm1 | MGI | protein_codin | YES | CCDS24532.1 | ENSMUSP0000 |
| -1 | Ptpn11 | MGI | protein_codin | YES | CCDS39247.1 | ENSMUSP0000 |
| 1 | Nf1 | MGI | protein_codin | YES | CCDS25119.1 | ENSMUSP0000 |
| -1 | Npm1 | MGI | protein_codin | YES | CCDS24532.1 | ENSMUSP0000 |
| -1 | Ptpn11 | MGI | protein_codin | YES | CCDS39247.1 | ENSMUSP0000 |
| -1 | Flt3 | MGI | protein_codin | YES | CCDS39400.1 | ENSMUSP0000 |
| -1 | Npm1 | MGI | protein_codin | YES | CCDS24532.1 | ENSMUSP0000 |
| -1 | Ptpn11 | MGI | protein_codin | YES | CCDS39247.1 | ENSMUSP0000 |
| -1 | Npm1 | MGI | protein_codin | YES | CCDS24532.1 | ENSMUSP0000 |
| -1 | Ptpn11 | MGI | protein_codin | YES | CCDS39247.1 | ENSMUSP0000 |
| -1 | Vegfa | MGI | protein_codin | YES | | ENSMUSP0000 |
| -1 | Erg | MGI | protein_codin | YES | CCDS37411.1 | ENSMUSP0000 |
| -1 | Flt3 | MGI | protein_codin | YES | CCDS39400.1 | ENSMUSP0000 |
| -1 | Npm1 | MGI | protein_codin | YES | CCDS24532.1 | ENSMUSP0000 |
| -1 | Ptpn11 | MGI | protein_codin | YES | CCDS39247.1 | ENSMUSP0000 |
| -1 | Ankrd11 | MGI | protein_codin | YES | CCDS40507.2 | ENSMUSP0000 |
| -1 | Flt3 | MGI | protein_codin | YES | CCDS39400.1 | ENSMUSP0000 |
| -1 | Npm1 | MGI | protein_codin | YES | CCDS24532.1 | ENSMUSP0000 |
| -1 | Ptpn11 | MGI | protein_codin | YES | CCDS39247.1 | ENSMUSP0000 |
| 1 | Rnf43 | MGI | protein_codin | YES | CCDS25215.2 | ENSMUSP0000 |
| -1 | Ros1 | MGI | protein_codin | YES | CCDS23838.1 | ENSMUSP0000 |
| -1 | Samhd1 | MGI | protein_codin | YES | CCDS16973.2 | ENSMUSP0000 |
| -1 | Flt3 | MGI | protein_codin | YES | CCDS39400.1 | ENSMUSP0000 |
| -1 | Npm1 | MGI | protein_codin | YES | CCDS24532.1 | ENSMUSP0000 |
| -1 | Ptpn11 | MGI | protein_codin | YES | CCDS39247.1 | ENSMUSP0000 |
| -1 | Npm1 | MGI | protein_codin | YES | CCDS24532.1 | ENSMUSP0000 |
| -1 | Ptpn11 | MGI | protein_codin | YES | CCDS39247.1 | ENSMUSP0000 |
| 1 | Kit | MGI | protein_codin | YES | CCDS51525.1 | ENSMUSP0000 |
| -1 | Lrp1b | MGI | protein_codin | YES | CCDS84520.1 | ENSMUSP0000 |
| -1 | Npm1 | MGI | protein_codin | YES | CCDS24532.1 | ENSMUSP0000 |

| | | | | | | |
|---|---|---|---|---|---|---|
| -1 | Ptpn11 | MGI | protein_codin | YES | CCDS39247.1 | ENSMUSP0000 |
| -1 | Serpinb3b | MGI | protein_codin | YES | CCDS15216.1 | ENSMUSP0000 |
| -1 | Ankrd11 | MGI | protein_codin | YES | CCDS40507.2 | ENSMUSP0000 |
| -1 | Flt3 | MGI | protein_codin | YES | CCDS39400.1 | ENSMUSP0000 |
| -1 | Npm1 | MGI | protein_codin | YES | CCDS24532.1 | ENSMUSP0000 |
| -1 | Ptpn11 | MGI | protein_codin | YES | CCDS39247.1 | ENSMUSP0000 |
| -1 | Samhd1 | MGI | protein_codin | YES | CCDS16973.2 | ENSMUSP0000 |
| -1 | Ankrd11 | MGI | protein_codin | YES | CCDS40507.2 | ENSMUSP0000 |
| -1 | Flt3 | MGI | protein_codin | YES | CCDS39400.1 | ENSMUSP0000 |
| -1 | Kmt2d | MGI | protein_codin | YES | CCDS49725.2 | ENSMUSP0000 |
| -1 | Npm1 | MGI | protein_codin | YES | CCDS24532.1 | ENSMUSP0000 |
| -1 | Ptpn11 | MGI | protein_codin | YES | CCDS39247.1 | ENSMUSP0000 |
| -1 | Samhd1 | MGI | protein_codin | YES | CCDS16973.2 | ENSMUSP0000 |
| -1 | Zmym3 | MGI | protein_codin | YES | CCDS30315.1 | ENSMUSP0000 |
| -1 | Kmt2d | MGI | protein_codin | YES | CCDS49725.2 | ENSMUSP0000 |
| 1 | Nf1 | MGI | protein_codin | YES | CCDS25119.1 | ENSMUSP0000 |
| -1 | Npm1 | MGI | protein_codin | YES | CCDS24532.1 | ENSMUSP0000 |
| -1 | Ptpn11 | MGI | protein_codin | YES | CCDS39247.1 | ENSMUSP0000 |

| SWISSPROT | TREMBL | UNIPARC | RefSeq | SIFT | EXON | DOMAINS |
|---|---|---|---|---|---|---|
| E9Q4F7 | | UPI0000605E5 | NM_0010813 | tolerated_low | 9/13 | PANTHER:PTH |
| P22682 | | UPI00003580 | NM_007619.2 | deleterious(0) | 8/16 | Gene3D:3.30. |
| | Q3UEW6 | UPI0000356E | NM_010229.2 | deleterious(0) | 20/24 | Gene3D:1.10. |
| Q61937 | Q5SQB7 | UPI0000003F | NM_0012522 | | 11/11 | Gene3D:1.10. |
| P35235 | | UPI00000E5F1 | NM_011202.3 | deleterious(0. | 3/16 | CDD:cd10340, |
| Q60710 | | UPI00015DF5 | NM_018851.4 | | 2/16 | PROSITE_profi |
| | Q3UEW6 | UPI0000356E | NM_010229.2 | deleterious(0) | 20/24 | Gene3D:1.10. |
| Q04690 | | UPI00000271 | NM_010897.2 | deleterious(0) | 33/58 | Gene3D:1.10. |
| Q04690 | | UPI00000271 | NM_010897.2 | | 33/58 | Gene3D:1.10. |
| Q61937 | Q5SQB7 | UPI0000003F | NM_0012522 | | 11/11 | Gene3D:1.10. |
| P35235 | | UPI00000E5F1 | NM_011202.3 | deleterious(0. | 3/16 | CDD:cd10340, |
| Q04690 | | UPI00000271 | NM_010897.2 | | 37/58 | Gene3D:3.40. |
| Q61937 | Q5SQB7 | UPI0000003F | NM_0012522 | | 11/11 | Gene3D:1.10. |
| P35235 | | UPI00000E5F1 | NM_011202.3 | deleterious(0. | 3/16 | CDD:cd10340, |
| | Q9D1Q5 | UPI00000220 | NM_198680.2 | deleterious(0. | 8/8 | Gene3D:2.30. |
| Q61937 | Q5SQB7 | UPI0000003F | NM_0012522 | | 11/11 | Gene3D:1.10. |
| P35235 | | UPI00000E5F1 | NM_011202.3 | deleterious(0. | 3/16 | CDD:cd10340, |
| | Q3UEW6 | UPI0000356E | NM_010229.2 | deleterious(0) | 20/24 | Gene3D:1.10. |
| Q04690 | | UPI00000271 | NM_010897.2 | | 45/58 | PANTHER:PTH |
| Q61937 | Q5SQB7 | UPI0000003F | NM_0012522 | | 11/11 | Gene3D:1.10. |
| P35235 | | UPI00000E5F1 | NM_011202.3 | deleterious(0. | 3/16 | CDD:cd10340, |
| Q04690 | | UPI00000271 | NM_010897.2 | | 28/58 | Gene3D:1.10. |
| Q61937 | Q5SQB7 | UPI0000003F | NM_0012522 | | 11/11 | Gene3D:1.10. |
| P35235 | | UPI00000E5F1 | NM_011202.3 | deleterious(0. | 3/16 | CDD:cd10340, |
| | Q3UEW6 | UPI0000356E | NM_010229.2 | deleterious(0) | 20/24 | Gene3D:1.10. |
| Q61937 | Q5SQB7 | UPI0000003F | NM_0012522 | | 11/11 | Gene3D:1.10. |
| P35235 | | UPI00000E5F1 | NM_011202.3 | deleterious(0. | 3/16 | CDD:cd10340, |
| Q61937 | Q5SQB7 | UPI0000003F | NM_0012522 | | 11/11 | Gene3D:1.10. |
| P35235 | | UPI00000E5F1 | NM_011202.3 | deleterious(0. | 3/16 | CDD:cd10340, |
| | F8WH81 | UPI0001F793 | NM_0010252 | tolerated(0.07 | 3/8 | Gene3D:2.10. |
| P81270 | | UPI00000285 | NM_133659.3 | | 7/11 | PANTHER:PTH |
| | Q3UEW6 | UPI0000356E | NM_010229.2 | deleterious(0) | 20/24 | Gene3D:1.10. |
| Q61937 | Q5SQB7 | UPI0000003F | NM_0012522 | | 11/11 | Gene3D:1.10. |
| P35235 | | UPI00000E5F1 | NM_011202.3 | deleterious(0. | 3/16 | CDD:cd10340, |
| E9Q4F7 | | UPI0000605E5 | NM_0010813 | tolerated_low | 9/13 | PANTHER:PTH |
| | Q3UEW6 | UPI0000356E | NM_010229.2 | deleterious(0) | 20/24 | Gene3D:1.10. |
| Q61937 | Q5SQB7 | UPI0000003F | NM_0012522 | | 11/11 | Gene3D:1.10. |
| P35235 | | UPI00000E5F1 | NM_011202.3 | deleterious(0. | 3/16 | CDD:cd10340, |
| Q5NCP0 | | UPI000049C6 | NM_172448.4 | | 9/10 | PANTHER:PTH |
| Q78DX7 | | UPI00000217 | NM_011282.2 | tolerated(0.16 | 27/44 | Gene3D:2.120 |
| Q60710 | | UPI00015DF5 | NM_018851.4 | | 2/16 | PROSITE_profi |
| | Q3UEW6 | UPI0000356E | NM_010229.2 | deleterious(0) | 20/24 | Gene3D:1.10. |
| Q61937 | Q5SQB7 | UPI0000003F | NM_0012522 | | 11/11 | Gene3D:1.10. |
| P35235 | | UPI00000E5F1 | NM_011202.3 | deleterious(0. | 3/16 | CDD:cd10340, |
| Q61937 | Q5SQB7 | UPI0000003F | NM_0012522 | | 11/11 | Gene3D:1.10. |
| P35235 | | UPI00000E5F1 | NM_011202.3 | deleterious(0. | 3/16 | CDD:cd10340, |
| P05532 | | UPI00000E98 | NM_0011227 | deleterious(0) | 17/21 | PROSITE_profi |
| | A2API5 | UPI00015968 | NM_053011.2 | | 14/91 | Gene3D:2.120 |
| Q61937 | Q5SQB7 | UPI0000003F | NM_0012522 | | 11/11 | Gene3D:1.10. |

| | | | | | | |
|---|---|---|---|---|---|---|
| P35235 | | UPI00000E5F1 | NM_011202.3 | deleterious(0.( | 3/16 | CDD:cd10340, |
| | Q9D1Q5 | UPI00000220[ | NM_198680.2 | deleterious(0.( | 8/8 | Gene3D:2.30.3 |
| E9Q4F7 | | UPI0000605E5 | NM_0010813: | tolerated_low | 9/13 | PANTHER:PTH |
| | Q3UEW6 | UPI0000356EE | NM_010229.2 | deleterious(0) | 20/24 | Gene3D:1.10.5 |
| Q61937 | Q5SQB7 | UPI0000003F6 | NM_0012522( | | 11/11 | Gene3D:1.10.1 |
| P35235 | | UPI00000E5F1 | NM_011202.3 | deleterious(0.( | 3/16 | CDD:cd10340, |
| Q60710 | | UPI00015DF5I | NM_018851.4 | | 2/16 | PROSITE_profi |
| E9Q4F7 | | UPI0000605E5 | NM_0010813: | tolerated_low | 9/13 | PANTHER:PTH |
| | Q3UEW6 | UPI0000356EE | NM_010229.2 | deleterious(0) | 20/24 | Gene3D:1.10.5 |
| | A0A0A0MQ73 | UPI00023B36/ | NM_0010332: | | 35/55 | Coiled-coils_(N |
| Q61937 | Q5SQB7 | UPI0000003F6 | NM_0012522( | | 11/11 | Gene3D:1.10.1 |
| P35235 | | UPI00000E5F1 | NM_011202.3 | deleterious(0.( | 3/16 | CDD:cd10340, |
| Q60710 | | UPI00015DF5I | NM_018851.4 | | 2/16 | PROSITE_profi |
| Q9JLM4 | B1AXS5 | UPI00000237: | NM_019831.3 | | 2/25 | PANTHER:PTH |
| | A0A0A0MQ73 | UPI00023B36/ | NM_0010332: | | 35/55 | Coiled-coils_(N |
| Q04690 | | UPI00000271( | NM_010897.2 | | 28/58 | Gene3D:1.10.5 |
| Q61937 | Q5SQB7 | UPI0000003F6 | NM_0012522( | | 11/11 | Gene3D:1.10.1 |
| P35235 | | UPI00000E5F1 | NM_011202.3 | deleterious(0.( | 3/16 | CDD:cd10340, |

| IMPACT | PICK | VARIANT_CLA | TSL | HGVS_OFFSET | GENE_PHENO | flanking_bps |
|---|---|---|---|---|---|---|
| MODERATE | | SNV | 5 | | 1 | ACG |
| MODERATE | 1 | SNV | 1 | | 1 | TAT |
| MODERATE | 1 | SNV | 1 | | 1 | GTC |
| HIGH | 1 | insertion | 1 | | 1 | GCC |
| MODERATE | 1 | SNV | 1 | | 1 | TCA |
| LOW | 1 | SNV | 1 | | 1 | CGG |
| MODERATE | 1 | SNV | 1 | | 1 | GTC |
| MODERATE | 1 | SNV | 2 | | 1 | TCA |
| HIGH | 1 | SNV | 2 | | 1 | ACG |
| HIGH | 1 | insertion | 1 | | 1 | GCC |
| MODERATE | 1 | SNV | 1 | | 1 | TCA |
| HIGH | 1 | deletion | 2 | 1 | 1 | ATCGCTTTAA[illegible] |
| HIGH | 1 | insertion | 1 | | 1 | GCC |
| MODERATE | 1 | SNV | 1 | | 1 | TCA |
| MODERATE | 1 | SNV | 1 | | | AAA |
| HIGH | 1 | insertion | 1 | | 1 | GCC |
| MODERATE | 1 | SNV | 1 | | 1 | TCA |
| MODERATE | 1 | SNV | 1 | | 1 | GTC |
| HIGH | 1 | SNV | 2 | | 1 | CCG |
| HIGH | 1 | insertion | 1 | | 1 | GCC |
| MODERATE | 1 | SNV | 1 | | 1 | TCA |
| HIGH | 1 | SNV | 2 | | 1 | TGA |
| HIGH | 1 | insertion | 1 | | 1 | GCC |
| MODERATE | 1 | SNV | 1 | | 1 | TCA |
| MODERATE | 1 | SNV | 1 | | 1 | GTC |
| HIGH | 1 | insertion | 1 | | 1 | GCC |
| MODERATE | 1 | SNV | 1 | | 1 | TCA |
| HIGH | 1 | insertion | 1 | | 1 | GCC |
| MODERATE | 1 | SNV | 1 | | 1 | TCA |
| MODERATE | 1 | SNV | 1 | | 1 | TCG |
| LOW | 1 | SNV | | | 1 | GCG |
| MODERATE | 1 | SNV | 1 | | 1 | GTC |
| HIGH | 1 | insertion | 1 | | 1 | GCC |
| MODERATE | 1 | SNV | 1 | | 1 | TCA |
| MODERATE | | SNV | 5 | | 1 | ACG |
| MODERATE | 1 | SNV | 1 | | 1 | GTC |
| HIGH | 1 | insertion | 1 | | 1 | GCC |
| MODERATE | 1 | SNV | 1 | | 1 | TCA |
| LOW | 1 | SNV | 5 | | | GAA |
| MODERATE | 1 | SNV | 1 | | 1 | GCA |
| LOW | 1 | SNV | 1 | | 1 | CGG |
| MODERATE | 1 | SNV | 1 | | 1 | GTC |
| HIGH | 1 | insertion | 1 | | 1 | GCC |
| MODERATE | 1 | SNV | 1 | | 1 | TCA |
| HIGH | 1 | insertion | 1 | | 1 | GCC |
| MODERATE | 1 | SNV | 1 | | 1 | TCA |
| MODERATE | 1 | SNV | 1 | | 1 | AGA |
| LOW | 1 | SNV | 1 | | 1 | ACG |
| HIGH | 1 | insertion | 1 | | 1 | GCC |

| | | | | | |
|---|---|---|---|---|---|
| MODERATE | 1 | SNV | 1 | 1 | TCA |
| MODERATE | 1 | SNV | 1 | | AAA |
| MODERATE | | SNV | 5 | 1 | ACG |
| MODERATE | 1 | SNV | 1 | 1 | GTC |
| HIGH | 1 | insertion | 1 | 1 | GCC |
| MODERATE | 1 | SNV | 1 | 1 | TCA |
| LOW | 1 | SNV | 1 | 1 | CGG |
| MODERATE | | SNV | 5 | 1 | ACG |
| MODERATE | 1 | SNV | 1 | 1 | GTC |
| MODERATE | 1 | deletion | 5 | 1 | GCTGCTGCTG |
| HIGH | 1 | insertion | 1 | 1 | GCC |
| MODERATE | 1 | SNV | 1 | 1 | TCA |
| LOW | 1 | SNV | 1 | 1 | CGG |
| LOW | 1 | SNV | 1 | | CTC |
| MODERATE | 1 | deletion | 5 | 1 | GCTGCTGCTG |
| HIGH | 1 | SNV | 2 | 1 | TGA |
| HIGH | 1 | insertion | 1 | 1 | GCC |
| MODERATE | 1 | SNV | 1 | 1 | TCA |

| vcf_id | vcf_qual | vcf_pos | Caller | POS | TAG | LABEL |
|---|---|---|---|---|---|---|
| . | . | 122890181 | mutect | chr8:1228901 | chr8:1228901 | chr8:1228901 |
| . | . | 44164242 | mutect.Rescu | chr9:4416424 | chr9:4416424 | chr9:4416424 |
| . | . | 147341238 | mutect | chr5:1473412 | chr5:1473412 | chr5:1473412 |
| . | 122585 | 33152835 | HaplotypeCall | chr11:331528 | chr11:331528 | chr11:331528 |
| . | . | 121167955 | mutect | chr5:1211679 | chr5:1211679 | chr5:1211679 |
| rs586562802 | . | 157129903 | mutect | chr2:1571299 | chr2:1571299 | chr2:1571299 |
| . | . | 147341238 | mutect.Rescu | chr5:1473412 | chr5:1473412 | chr5:1473412 |
| . | . | 79469779 | mutect | chr11:794697 | chr11:794697 | chr11:794697 |
| . | . | 79469866 | mutect | chr11:794698 | chr11:794698 | chr11:794698 |
| . | 122585 | 33152835 | HaplotypeCall | chr11:331528 | chr11:331528 | chr11:331528 |
| . | . | 121167955 | mutect | chr5:1211679 | chr5:1211679 | chr5:1211679 |
| . | 791.66 | 79535723 | HaplotypeCall | chr11:795357 | chr11:795357 | chr11:795357 |
| . | 122585 | 33152835 | HaplotypeCall | chr11:331528 | chr11:331528 | chr11:331528 |
| . | . | 121167955 | mutect | chr5:1211679 | chr5:1211679 | chr5:1211679 |
| . | . | 107154541 | mutect | chr1:1071545 | chr1:1071545 | chr1:1071545 |
| . | 122585 | 33152835 | HaplotypeCall | chr11:331528 | chr11:331528 | chr11:331528 |
| . | . | 121167955 | mutect | chr5:1211679 | chr5:1211679 | chr5:1211679 |
| . | . | 147341238 | mutect | chr5:1473412 | chr5:1473412 | chr5:1473412 |
| . | . | 79548774 | mutect | chr11:795487 | chr11:795487 | chr11:795487 |
| . | 122585 | 33152835 | HaplotypeCall | chr11:331528 | chr11:331528 | chr11:331528 |
| . | . | 121167955 | mutect | chr5:1211679 | chr5:1211679 | chr5:1211679 |
| . | . | 79453806 | mutect | chr11:794538 | chr11:794538 | chr11:794538 |
| . | 122585 | 33152835 | HaplotypeCall | chr11:331528 | chr11:331528 | chr11:331528 |
| . | . | 121167955 | mutect | chr5:1211679 | chr5:1211679 | chr5:1211679 |
| . | . | 147341238 | mutect | chr5:1473412 | chr5:1473412 | chr5:1473412 |
| . | 122585 | 33152835 | HaplotypeCall | chr11:331528 | chr11:331528 | chr11:331528 |
| . | . | 121167955 | mutect | chr5:1211679 | chr5:1211679 | chr5:1211679 |
| . | 122585 | 33152835 | HaplotypeCall | chr11:331528 | chr11:331528 | chr11:331528 |
| . | . | 121167955 | mutect | chr5:1211679 | chr5:1211679 | chr5:1211679 |
| . | . | 46025460 | mutect | chr17:460254 | chr17:460254 | chr17:460254 |
| . | . | 95377323 | mutect | chr16:953773 | chr16:953773 | chr16:953773 |
| . | . | 147341238 | mutect | chr5:1473412 | chr5:1473412 | chr5:1473412 |
| . | 122585 | 33152835 | HaplotypeCall | chr11:331528 | chr11:331528 | chr11:331528 |
| . | . | 121167955 | mutect | chr5:1211679 | chr5:1211679 | chr5:1211679 |
| . | . | 122890181 | mutect | chr8:1228901 | chr8:1228901 | chr8:1228901 |
| . | . | 147341238 | mutect | chr5:1473412 | chr5:1473412 | chr5:1473412 |
| . | 122585 | 33152835 | HaplotypeCall | chr11:331528 | chr11:331528 | chr11:331528 |
| . | . | 121167955 | mutect | chr5:1211679 | chr5:1211679 | chr5:1211679 |
| . | . | 87731568 | mutect | chr11:877315 | chr11:877315 | chr11:877315 |
| . | . | 52115994 | mutect | chr10:521159 | chr10:521159 | chr10:521159 |
| rs586562802 | . | 157129903 | mutect | chr2:1571299 | chr2:1571299 | chr2:1571299 |
| . | . | 147341238 | mutect | chr5:1473412 | chr5:1473412 | chr5:1473412 |
| . | 122585 | 33152835 | HaplotypeCall | chr11:331528 | chr11:331528 | chr11:331528 |
| . | . | 121167955 | mutect | chr5:1211679 | chr5:1211679 | chr5:1211679 |
| . | 122585 | 33152835 | HaplotypeCall | chr11:331528 | chr11:331528 | chr11:331528 |
| . | . | 121167955 | mutect | chr5:1211679 | chr5:1211679 | chr5:1211679 |
| . | . | 75649631 | mutect | chr5:7564963 | chr5:7564963 | chr5:7564963 |
| . | . | 41468886 | mutect | chr2:4146888 | chr2:4146888 | chr2:4146888 |
| . | 122585 | 33152835 | HaplotypeCall | chr11:331528 | chr11:331528 | chr11:331528 |

| | | | | | | |
|---|---|---|---|---|---|---|
| . | . | 121167955 | mutect | chr5:1211679 | chr5:1211679 | chr5:1211679 |
| . | . | 107154541 | mutect | chr1:1071545 | chr1:1071545 | chr1:1071545 |
| . | . | 122890181 | mutect | chr8:1228901 | chr8:1228901 | chr8:1228901 |
| . | . | 147341238 | mutect | chr5:1473412 | chr5:1473412 | chr5:1473412 |
| . | 122585 | 33152835 | HaplotypeCall | chr11:331528 | chr11:331528 | chr11:331528 |
| . | . | 121167955 | mutect | chr5:1211679 | chr5:1211679 | chr5:1211679 |
| rs586562802 | . | 157129903 | mutect | chr2:1571299 | chr2:1571299 | chr2:1571299 |
| . | . | 122890181 | mutect | chr8:1228901 | chr8:1228901 | chr8:1228901 |
| . | . | 147341238 | mutect | chr5:1473412 | chr5:1473412 | chr5:1473412 |
| . | 775.96 | 98851066 | HaplotypeCall | chr15:988510 | chr15:988510 | chr15:988510 |
| . | 122585 | 33152835 | HaplotypeCall | chr11:331528 | chr11:331528 | chr11:331528 |
| . | . | 121167955 | mutect | chr5:1211679 | chr5:1211679 | chr5:1211679 |
| rs586562802 | . | 157129903 | mutect | chr2:1571299 | chr2:1571299 | chr2:1571299 |
| . | . | 101418712 | mutect | chrX:1014187 | chrX:1014187 | chrX:1014187 |
| . | 775.96 | 98851066 | HaplotypeCall | chr15:988510 | chr15:988510 | chr15:988510 |
| . | . | 79453806 | mutect | chr11:794538 | chr11:794538 | chr11:794538 |
| . | 122585 | 33152835 | HaplotypeCall | chr11:331528 | chr11:331528 | chr11:331528 |
| . | . | 121167955 | mutect | chr5:1211679 | chr5:1211679 | chr5:1211679 |

| TriNuc | ETAG | TargetPanel | Normal_CTRL | Normal_CTRL | Normal_CTRL | Normal_CTRL |
|---|---|---|---|---|---|---|
| ACG | chr8:1228901 | M-IMPACT_v2 | 0 | 0 | 2609 | 0 |
| TAT | chr9:4416424 | M-IMPACT_v2 | 0.010989 | 3 | 4197 | 0.0007148 |
| GTC | chr5:1473412 | M-IMPACT_v2 | 0 | 0 | 3961 | 0 |
|  | chr11:331528 | M-IMPACT_v2 | 0 | 0 | 9470 | 0 |
| TCA | chr5:1211679 | M-IMPACT_v2 | 0.003663 | 1 | 5850 | 0.00017094 |
| CGG | chr2:1571299 | M-IMPACT_v2 | 0.00303951 | 4 | 7188 | 0.00055648 |
| GTC | chr5:1473412 | M-IMPACT_v2 | 0 | 0 | 3961 | 0 |
| TCA | chr11:794697 | M-IMPACT_v2 | 0.00234742 | 2 | 6270 | 0.00031898 |
| ACG | chr11:794698 | M-IMPACT_v2 | 0.0027027 | 1 | 5892 | 0.00016972 |
|  | chr11:331528 | M-IMPACT_v2 | 0 | 0 | 9470 | 0 |
| TCA | chr5:1211679 | M-IMPACT_v2 | 0.003663 | 1 | 5850 | 0.00017094 |
|  | chr11:795357 | M-IMPACT_v2 | 0 | 0 | 6962 | 0 |
|  | chr11:331528 | M-IMPACT_v2 | 0 | 0 | 9470 | 0 |
| TCA | chr5:1211679 | M-IMPACT_v2 | 0.003663 | 1 | 5850 | 0.00017094 |
| AAA | chr1:1071545 | M-IMPACT_v2 | 0.00263852 | 6 | 8341 | 0.00071934 |
|  | chr11:331528 | M-IMPACT_v2 | 0 | 0 | 9470 | 0 |
| TCA | chr5:1211679 | M-IMPACT_v2 | 0.003663 | 1 | 5850 | 0.00017094 |
| GTC | chr5:1473412 | M-IMPACT_v2 | 0 | 0 | 3961 | 0 |
| CCG | chr11:795487 | M-IMPACT_v2 | 0 | 0 | 5404 | 0 |
|  | chr11:331528 | M-IMPACT_v2 | 0 | 0 | 9470 | 0 |
| TCA | chr5:1211679 | M-IMPACT_v2 | 0.003663 | 1 | 5850 | 0.00017094 |
| TGA | chr11:794538 | M-IMPACT_v2 | 0.00273973 | 3 | 5086 | 0.00058985 |
|  | chr11:331528 | M-IMPACT_v2 | 0 | 0 | 9470 | 0 |
| TCA | chr5:1211679 | M-IMPACT_v2 | 0.003663 | 1 | 5850 | 0.00017094 |
| GTC | chr5:1473412 | M-IMPACT_v2 | 0 | 0 | 3961 | 0 |
|  | chr11:331528 | M-IMPACT_v2 | 0 | 0 | 9470 | 0 |
| TCA | chr5:1211679 | M-IMPACT_v2 | 0.003663 | 1 | 5850 | 0.00017094 |
|  | chr11:331528 | M-IMPACT_v2 | 0 | 0 | 9470 | 0 |
| TCA | chr5:1211679 | M-IMPACT_v2 | 0.003663 | 1 | 5850 | 0.00017094 |
| TCG | chr17:460254 | M-IMPACT_v2 | 0 | 0 | 4530 | 0 |
| GCG | chr16:953773 | M-IMPACT_v2 | 0 | 0 | 4393 | 0 |
| GTC | chr5:1473412 | M-IMPACT_v2 | 0 | 0 | 3961 | 0 |
|  | chr11:331528 | M-IMPACT_v2 | 0 | 0 | 9470 | 0 |
| TCA | chr5:1211679 | M-IMPACT_v2 | 0.003663 | 1 | 5850 | 0.00017094 |
| ACG | chr8:1228901 | M-IMPACT_v2 | 0 | 0 | 2609 | 0 |
| GTC | chr5:1473412 | M-IMPACT_v2 | 0 | 0 | 3961 | 0 |
|  | chr11:331528 | M-IMPACT_v2 | 0 | 0 | 9470 | 0 |
| TCA | chr5:1211679 | M-IMPACT_v2 | 0.003663 | 1 | 5850 | 0.00017094 |
| GAA | chr11:877315 | M-IMPACT_v2 | 0.00518135 | 3 | 4373 | 0.00068603 |
| GCA | chr10:521159 | M-IMPACT_v2 | 0.00701754 | 7 | 7245 | 0.00096618 |
| CGG | chr2:1571299 | M-IMPACT_v2 | 0.00303951 | 4 | 7188 | 0.00055648 |
| GTC | chr5:1473412 | M-IMPACT_v2 | 0 | 0 | 3961 | 0 |
|  | chr11:331528 | M-IMPACT_v2 | 0 | 0 | 9470 | 0 |
| TCA | chr5:1211679 | M-IMPACT_v2 | 0.003663 | 1 | 5850 | 0.00017094 |
|  | chr11:331528 | M-IMPACT_v2 | 0 | 0 | 9470 | 0 |
| TCA | chr5:1211679 | M-IMPACT_v2 | 0.003663 | 1 | 5850 | 0.00017094 |
| AGA | chr5:7564963 | M-IMPACT_v2 | 0.00512821 | 3 | 6092 | 0.00049245 |
| ACG | chr2:4146888 | M-IMPACT_v2 | 0.00425532 | 3 | 6322 | 0.00047453 |
|  | chr11:331528 | M-IMPACT_v2 | 0 | 0 | 9470 | 0 |

| | | | | | | |
|---|---|---|---|---|---|---|
| TCA | chr5:1211679[illegible] | M-IMPACT_v2 | 0.003663 | 1 | 5850 | 0.00017094 |
| AAA | chr1:1071545[illegible] | M-IMPACT_v2 | 0.00263852 | 6 | 8341 | 0.00071934 |
| ACG | chr8:1228901[illegible] | M-IMPACT_v2 | 0 | 0 | 2609 | 0 |
| GTC | chr5:1473412[illegible] | M-IMPACT_v2 | 0 | 0 | 3961 | 0 |
| | chr11:331528[illegible] | M-IMPACT_v2 | 0 | 0 | 9470 | 0 |
| TCA | chr5:1211679[illegible] | M-IMPACT_v2 | 0.003663 | 1 | 5850 | 0.00017094 |
| CGG | chr2:1571299[illegible] | M-IMPACT_v2 | 0.00303951 | 4 | 7188 | 0.00055648 |
| ACG | chr8:1228901[illegible] | M-IMPACT_v2 | 0 | 0 | 2609 | 0 |
| GTC | chr5:1473412[illegible] | M-IMPACT_v2 | 0 | 0 | 3961 | 0 |
| | chr15:988510[illegible] | M-IMPACT_v2 | 0 | 0 | 4632 | 0 |
| | chr11:331528[illegible] | M-IMPACT_v2 | 0 | 0 | 9470 | 0 |
| TCA | chr5:1211679[illegible] | M-IMPACT_v2 | 0.003663 | 1 | 5850 | 0.00017094 |
| CGG | chr2:1571299[illegible] | M-IMPACT_v2 | 0.00303951 | 4 | 7188 | 0.00055648 |
| CTC | chrX:1014187 | M-IMPACT_v2 | 0 | 0 | 2602 | 0 |
| | chr15:988510[illegible] | M-IMPACT_v2 | 0 | 0 | 4632 | 0 |
| TGA | chr11:794538[illegible] | M-IMPACT_v2 | 0.00273973 | 3 | 5086 | 0.00058985 |
| | chr11:331528[illegible] | M-IMPACT_v2 | 0 | 0 | 9470 | 0 |
| TCA | chr5:1211679[illegible] | M-IMPACT_v2 | 0.003663 | 1 | 5850 | 0.00017094 |

| POOL_maxVF | POOL_tAD | POOL_tDP | POOL_tVAF | AllNormal_tDI | AllNormal_tAI | AllNormal_tFr |
|---|---|---|---|---|---|---|
| 0 | 0 | 174 | 0 | 2783 | 0 | 0.00035907 |
| 0.00377358 | 1 | 265 | 0.00377358 | 4462 | 4 | 0.00112007 |
| 0 | 0 | 224 | 0 | 4185 | 0 | 0.00023883 |
| 0 | 0 | 477 | 0 | 9947 | 0 | 0.00010051 |
| 0 | 0 | 334 | 0 | 6184 | 1 | 0.00032331 |
| 0 | 0 | 331 | 0 | 7519 | 4 | 0.00066481 |
| 0 | 0 | 224 | 0 | 4185 | 0 | 0.00023883 |
| 0 | 0 | 340 | 0 | 6610 | 2 | 0.00045372 |
| 0 | 0 | 320 | 0 | 6212 | 1 | 0.00032185 |
| 0 | 0 | 477 | 0 | 9947 | 0 | 0.00010051 |
| 0 | 0 | 334 | 0 | 6184 | 1 | 0.00032331 |
| 0 | 0 | 389 | 0 | 7351 | 0 | 0.000136 |
| 0 | 0 | 477 | 0 | 9947 | 0 | 0.00010051 |
| 0 | 0 | 334 | 0 | 6184 | 1 | 0.00032331 |
| 0 | 0 | 480 | 0 | 8821 | 6 | 0.00079338 |
| 0 | 0 | 477 | 0 | 9947 | 0 | 0.00010051 |
| 0 | 0 | 334 | 0 | 6184 | 1 | 0.00032331 |
| 0 | 0 | 224 | 0 | 4185 | 0 | 0.00023883 |
| 0 | 0 | 264 | 0 | 5668 | 0 | 0.00017637 |
| 0 | 0 | 477 | 0 | 9947 | 0 | 0.00010051 |
| 0 | 0 | 334 | 0 | 6184 | 1 | 0.00032331 |
| 0 | 0 | 271 | 0 | 5357 | 3 | 0.00074641 |
| 0 | 0 | 477 | 0 | 9947 | 0 | 0.00010051 |
| 0 | 0 | 334 | 0 | 6184 | 1 | 0.00032331 |
| 0 | 0 | 224 | 0 | 4185 | 0 | 0.00023883 |
| 0 | 0 | 477 | 0 | 9947 | 0 | 0.00010051 |
| 0 | 0 | 334 | 0 | 6184 | 1 | 0.00032331 |
| 0 | 0 | 477 | 0 | 9947 | 0 | 0.00010051 |
| 0 | 0 | 334 | 0 | 6184 | 1 | 0.00032331 |
| 0 | 0 | 261 | 0 | 4791 | 0 | 0.00020864 |
| 0 | 0 | 255 | 0 | 4648 | 0 | 0.00021505 |
| 0 | 0 | 224 | 0 | 4185 | 0 | 0.00023883 |
| 0 | 0 | 477 | 0 | 9947 | 0 | 0.00010051 |
| 0 | 0 | 334 | 0 | 6184 | 1 | 0.00032331 |
| 0 | 0 | 174 | 0 | 2783 | 0 | 0.00035907 |
| 0 | 0 | 224 | 0 | 4185 | 0 | 0.00023883 |
| 0 | 0 | 477 | 0 | 9947 | 0 | 0.00010051 |
| 0 | 0 | 334 | 0 | 6184 | 1 | 0.00032331 |
| 0 | 0 | 292 | 0 | 4665 | 3 | 0.00085708 |
| 0 | 0 | 403 | 0 | 7648 | 7 | 0.00104575 |
| 0 | 0 | 331 | 0 | 7519 | 4 | 0.00066481 |
| 0 | 0 | 224 | 0 | 4185 | 0 | 0.00023883 |
| 0 | 0 | 477 | 0 | 9947 | 0 | 0.00010051 |
| 0 | 0 | 334 | 0 | 6184 | 1 | 0.00032331 |
| 0 | 0 | 477 | 0 | 9947 | 0 | 0.00010051 |
| 0 | 0 | 334 | 0 | 6184 | 1 | 0.00032331 |
| 0 | 0 | 348 | 0 | 6440 | 3 | 0.00062093 |
| 0 | 0 | 399 | 0 | 6721 | 3 | 0.00059497 |
| 0 | 0 | 477 | 0 | 9947 | 0 | 0.00010051 |

| | | | | | | |
|---|---|---|---|---|---|---|
| 0 | 0 | 334 | 0 | 6184 | 1 | 0.00032331 |
| 0 | 0 | 480 | 0 | 8821 | 6 | 0.00079338 |
| 0 | 0 | 174 | 0 | 2783 | 0 | 0.00035907 |
| 0 | 0 | 224 | 0 | 4185 | 0 | 0.00023883 |
| 0 | 0 | 477 | 0 | 9947 | 0 | 0.00010051 |
| 0 | 0 | 334 | 0 | 6184 | 1 | 0.00032331 |
| 0 | 0 | 331 | 0 | 7519 | 4 | 0.00066481 |
| 0 | 0 | 174 | 0 | 2783 | 0 | 0.00035907 |
| 0 | 0 | 224 | 0 | 4185 | 0 | 0.00023883 |
| 0 | 0 | 314 | 0 | 4946 | 0 | 0.0002021 |
| 0 | 0 | 477 | 0 | 9947 | 0 | 0.00010051 |
| 0 | 0 | 334 | 0 | 6184 | 1 | 0.00032331 |
| 0 | 0 | 331 | 0 | 7519 | 4 | 0.00066481 |
| 0 | 0 | 179 | 0 | 2781 | 0 | 0.00035932 |
| 0 | 0 | 314 | 0 | 4946 | 0 | 0.0002021 |
| 0 | 0 | 271 | 0 | 5357 | 3 | 0.00074641 |
| 0 | 0 | 477 | 0 | 9947 | 0 | 0.00010051 |
| 0 | 0 | 334 | 0 | 6184 | 1 | 0.00032331 |

| tVafFreqP | BINOM.pv | SNR.lor | BINOM.fdr |
|---|---|---|---|
| 0.5147541 | 0.00071826 | 3.47030717 | 0.00193758 |
| 0.02912621 | 0.00223939 | 1.42738872 | 0.00406288 |
| 0.07769424 | 0.00047773 | 2.5473131 | 0.00182271 |
| 0.41252006 | 0.00020104 | 3.84418781 | 0.00182271 |
| 0.50092764 | 0.00064662 | 3.49185097 | 0.00186468 |
| 0.278157 | 0.00132935 | 2.762864 | 0.00281776 |
| 0.02290076 | 0.00047773 | 1.99171051 | 0.00182271 |
| 0.32224532 | 0.00090737 | 3.0201286 | 0.00236872 |
| 0.35123967 | 0.00064371 | 3.22572072 | 0.00186468 |
| 0.40116279 | 0.00020104 | 3.82374764 | 0.00182271 |
| 0.47546531 | 0.00064662 | 3.44758411 | 0.00186468 |
| 0.08913043 | 0.00027202 | 2.85697533 | 0.00182271 |
| 0.39423077 | 0.00020104 | 3.81117908 | 0.00182271 |
| 0.47657512 | 0.00064662 | 3.44951649 | 0.00186468 |
| 0.49056604 | 0.00158631 | 3.08378313 | 0.00314724 |
| 0.37300613 | 0.00020104 | 3.77218846 | 0.00182271 |
| 0.50813008 | 0.00064662 | 3.50436413 | 0.00186468 |
| 0.15718157 | 0.00047773 | 2.89246683 | 0.00182271 |
| 0.04580153 | 0.00035276 | 2.43474769 | 0.00182271 |
| 0.41340782 | 0.00020104 | 3.8457782 | 0.00182271 |
| 0.4751773 | 0.00064662 | 3.44708257 | 0.00186468 |
| 0.88065099 | 0.00149254 | 3.99468451 | 0.00303401 |
| 0.87101064 | 0.00020104 | 4.82720534 | 0.00182271 |
| 0.45268139 | 0.00064662 | 3.40779191 | 0.00186468 |
| 0.44705882 | 0.00047773 | 3.52948496 | 0.00182271 |
| 0.34115806 | 0.00020104 | 3.71191017 | 0.00182271 |
| 0.44761905 | 0.00064662 | 3.39890935 | 0.00186468 |
| 0.36702955 | 0.00020104 | 3.76105337 | 0.00182271 |
| 0.47817837 | 0.00064662 | 3.45230733 | 0.00186468 |
| 0.03676471 | 0.00041732 | 2.26221552 | 0.00182271 |
| 0.03667482 | 0.00043015 | 2.24795458 | 0.00182271 |
| 0.42195122 | 0.00047773 | 3.48509698 | 0.00182271 |
| 0.39914773 | 0.00020104 | 3.82010173 | 0.00182271 |
| 0.45061728 | 0.00064662 | 3.40417234 | 0.00186468 |
| 0.49257426 | 0.00071826 | 3.43176845 | 0.00193758 |
| 0.46888889 | 0.00047773 | 3.56768378 | 0.00182271 |
| 0.34831461 | 0.00020104 | 3.72566948 | 0.00182271 |
| 0.43676223 | 0.00064662 | 3.37979278 | 0.00186468 |
| 0.02845528 | 0.0017138 | 1.53330556 | 0.00337319 |
| 0.04037267 | 0.00209068 | 1.60410192 | 0.00386932 |
| 0.40798611 | 0.00132935 | 3.01533025 | 0.00281776 |
| 0.48705882 | 0.00047773 | 3.59931308 | 0.00182271 |
| 0.35555556 | 0.00020104 | 3.73945776 | 0.00182271 |
| 0.43714822 | 0.00064662 | 3.38047415 | 0.00186468 |
| 0.37816456 | 0.00020104 | 3.78174113 | 0.00182271 |
| 0.43257443 | 0.00064662 | 3.37239143 | 0.00186468 |
| 0.34804754 | 0.00124166 | 2.93411362 | 0.00277326 |
| 0.23339012 | 0.00118977 | 2.7087527 | 0.00275759 |
| 0.35251799 | 0.00020104 | 3.73368935 | 0.00182271 |

| | | | |
|---|---|---|---|
| 0.39638158 | 0.00064662 | 3.30759046 | 0.00186468 |
| 0.42274678 | 0.00158631 | 2.96488749 | 0.00314724 |
| 0.43781095 | 0.00071826 | 3.33607346 | 0.00193758 |
| 0.37783375 | 0.00047773 | 3.40519353 | 0.00182271 |
| 0.31241084 | 0.00020104 | 3.65513285 | 0.00182271 |
| 0.40497336 | 0.00064662 | 3.32312953 | 0.00186468 |
| 0.40564374 | 0.00132935 | 3.0111147 | 0.00281776 |
| 0.36809816 | 0.00071826 | 3.20998326 | 0.00193758 |
| 0.37874659 | 0.00047773 | 3.40687918 | 0.00182271 |
| 0.09224012 | 0.00040424 | 2.70129077 | 0.00182271 |
| 0.2724359 | 0.00020104 | 3.57112885 | 0.00182271 |
| 0.35575221 | 0.00064662 | 3.23233416 | 0.00186468 |
| 0.34538879 | 0.00132935 | 2.89934156 | 0.00281776 |
| 0.05572755 | 0.00071878 | 2.21532979 | 0.00193758 |
| 0.11652794 | 0.00040424 | 2.81457917 | 0.00182271 |
| 0.78458498 | 0.00149254 | 3.68806349 | 0.00303401 |
| 0.654047 | 0.00020104 | 4.27432763 | 0.00182271 |
| 0.34789392 | 0.00064662 | 3.21736807 | 0.00186468 |

| Hugo_Symbol | Entrez_Gene_ | Center | NCBI_Build | Chromosome | Start_Position | End_Position |
|---|---|---|---|---|---|---|
| Etv4 | 0 | . | mm10 | chr11 | 101774169 | 101774169 |
| Npm1 | 0 | . | mm10 | chr11 | 33152835 | 33152836 |
| Myo18a | 0 | . | mm10 | chr11 | 77843108 | 77843108 |
| Smyd3 | 0 | . | mm10 | chr1 | 179092925 | 179092925 |
| Sptbn5 | 0 | . | mm10 | chr2 | 120047526 | 120047528 |
| Samhd1 | 0 | . | mm10 | chr2 | 157129903 | 157129903 |
| Ivl | 0 | . | mm10 | chr3 | 92572302 | 92572304 |
| Ptpn11 | 0 | . | mm10 | chr5 | 121167955 | 121167955 |
| Gigyf1 | 0 | . | mm10 | chr5 | 137525323 | 137525325 |
| Flt3 | 0 | . | mm10 | chr5 | 147341238 | 147341238 |
| Pik3c2g | 0 | . | mm10 | chr6 | 139843901 | 139843901 |
| Maz | 0 | . | mm10 | chr7 | 127023125 | 127023125 |
| Maz | 0 | . | mm10 | chr7 | 127023125 | 127023126 |
| Setd1a | 0 | . | mm10 | chr7 | 127785317 | 127785317 |
| Scgb1b30 | 0 | . | mm10 | chr7 | 34095574 | 34095574 |
| Rsf1 | 0 | . | mm10 | chr7 | 97579910 | 97579910 |
| Ankrd11 | 0 | . | mm10 | chr8 | 122890181 | 122890181 |
| Cbl | 0 | . | mm10 | chr9 | 44164242 | 44164242 |
| Rbm10 | 0 | . | mm10 | chrX | 20637584 | 20637584 |
| Egr2 | 0 | . | mm10 | chr10 | 67540518 | 67540518 |
| Npm1 | 0 | . | mm10 | chr11 | 33152835 | 33152836 |
| Nf1 | 0 | . | mm10 | chr11 | 79469779 | 79469779 |
| Nf1 | 0 | . | mm10 | chr11 | 79469866 | 79469866 |
| Vgll3 | 0 | . | mm10 | chr16 | 65827954 | 65827956 |
| Klhl14 | 0 | . | mm10 | chr18 | 21652101 | 21652103 |
| Nt5c2 | 0 | . | mm10 | chr19 | 46924325 | 46924325 |
| Arid1a | 0 | . | mm10 | chr4 | 133752569 | 133752571 |
| Prdm13 | 0 | . | mm10 | chr4 | 21678845 | 21678845 |
| Ptpn11 | 0 | . | mm10 | chr5 | 121167955 | 121167955 |
| Flt3 | 0 | . | mm10 | chr5 | 147341238 | 147341238 |
| Maz | 0 | . | mm10 | chr7 | 127023125 | 127023125 |
| Maz | 0 | . | mm10 | chr7 | 127023125 | 127023126 |
| Npm1 | 0 | . | mm10 | chr11 | 33152835 | 33152836 |
| Nf1 | 0 | . | mm10 | chr11 | 79535724 | 79535733 |
| Rb1 | 0 | . | mm10 | chr14 | 73325591 | 73325591 |
| Serpinb3b | 0 | . | mm10 | chr1 | 107154541 | 107154541 |
| Sptbn5 | 0 | . | mm10 | chr2 | 120047526 | 120047528 |
| Ivl | 0 | . | mm10 | chr3 | 92572302 | 92572304 |
| Ptpn11 | 0 | . | mm10 | chr5 | 121167955 | 121167955 |
| Gigyf1 | 0 | . | mm10 | chr5 | 137525323 | 137525325 |
| D6Ertd527e | 0 | . | mm10 | chr6 | 87112125 | 87112125 |
| Plk1 | 0 | . | mm10 | chr7 | 122167650 | 122167650 |
| Maz | 0 | . | mm10 | chr7 | 127023125 | 127023125 |
| Maz | 0 | . | mm10 | chr7 | 127023125 | 127023126 |
| Rps19 | 0 | . | mm10 | chr7 | 24888290 | 24888290 |
| Cdh16 | 0 | . | mm10 | chr8 | 104615215 | 104615215 |
| Ammecr1 | 0 | . | mm10 | chrX | 142966444 | 142966446 |
| Npm1 | 0 | . | mm10 | chr11 | 33152835 | 33152836 |
| Zswim6 | 0 | . | mm10 | chr13 | 107889970 | 107889972 |

| | | | | | | |
|---|---|---|---|---|---|---|
| Hcn1 | 0 | . | mm10 | chr13 | 117975767 | 117975769 |
| Ranbp9 | 0 | . | mm10 | chr13 | 43480914 | 43480914 |
| Fam170b | 0 | . | mm10 | chr14 | 32835939 | 32835939 |
| Dach1 | 0 | . | mm10 | chr14 | 98168875 | 98168875 |
| Kmt2d | 0 | . | mm10 | chr15 | 98849588 | 98849590 |
| Vgll3 | 0 | . | mm10 | chr16 | 65827954 | 65827956 |
| Sptbn5 | 0 | . | mm10 | chr2 | 120047526 | 120047528 |
| Maml3 | 0 | . | mm10 | chr3 | 51856069 | 51856069 |
| Ivl | 0 | . | mm10 | chr3 | 92572302 | 92572304 |
| Ptpn11 | 0 | . | mm10 | chr5 | 121167955 | 121167955 |
| Rbm33 | 0 | . | mm10 | chr5 | 28394148 | 28394150 |
| Cckar | 0 | . | mm10 | chr5 | 53700295 | 53700295 |
| Maz | 0 | . | mm10 | chr7 | 127023125 | 127023125 |
| Maz | 0 | . | mm10 | chr7 | 127023125 | 127023126 |
| Srcap | 0 | . | mm10 | chr7 | 127558307 | 127558307 |
| Cdh16 | 0 | . | mm10 | chr8 | 104615215 | 104615215 |
| Zfp703 | 0 | . | mm10 | chr8 | 26977426 | 26977426 |
| Ammecr1 | 0 | . | mm10 | chrX | 142966444 | 142966446 |
| Rbm10 | 0 | . | mm10 | chrX | 20637584 | 20637584 |
| Sry | 0 | . | mm10 | chrY | 2662945 | 2662947 |
| Npm1 | 0 | . | mm10 | chr11 | 33152835 | 33152836 |
| Nf1 | 0 | . | mm10 | chr11 | 79548774 | 79548774 |
| Zswim6 | 0 | . | mm10 | chr13 | 107889970 | 107889972 |
| Zfp503 | 0 | . | mm10 | chr14 | 21987453 | 21987455 |
| Vgll3 | 0 | . | mm10 | chr16 | 65827954 | 65827956 |
| Sptbn5 | 0 | . | mm10 | chr2 | 120047526 | 120047528 |
| Dennd4b | 0 | . | mm10 | chr3 | 90275569 | 90275571 |
| Ptpn11 | 0 | . | mm10 | chr5 | 121167955 | 121167955 |
| Gigyf1 | 0 | . | mm10 | chr5 | 137525323 | 137525325 |
| Flt3 | 0 | . | mm10 | chr5 | 147341238 | 147341238 |
| Cckar | 0 | . | mm10 | chr5 | 53700295 | 53700295 |
| Ammecr1 | 0 | . | mm10 | chrX | 142966444 | 142966446 |
| Mamld1 | 0 | . | mm10 | chrX | 71118851 | 71118851 |
| Amer1 | 0 | . | mm10 | chrX | 95427365 | 95427365 |
| Npm1 | 0 | . | mm10 | chr11 | 33152835 | 33152836 |
| Adamts2 | 0 | . | mm10 | chr11 | 50602214 | 50602214 |
| Nf1 | 0 | . | mm10 | chr11 | 79453806 | 79453806 |
| Hcn1 | 0 | . | mm10 | chr13 | 117975767 | 117975769 |
| Kmt2d | 0 | . | mm10 | chr15 | 98845132 | 98845134 |
| Sptbn5 | 0 | . | mm10 | chr2 | 120047526 | 120047528 |
| Ep400 | 0 | . | mm10 | chr5 | 110673217 | 110673217 |
| Ptpn11 | 0 | . | mm10 | chr5 | 121167955 | 121167955 |
| Rbm33 | 0 | . | mm10 | chr5 | 28394148 | 28394150 |
| D6Ertd527e | 0 | . | mm10 | chr6 | 87112125 | 87112125 |
| Maz | 0 | . | mm10 | chr7 | 127023125 | 127023125 |
| Maz | 0 | . | mm10 | chr7 | 127023125 | 127023126 |
| Rsf1 | 0 | . | mm10 | chr7 | 97579910 | 97579910 |
| Rsf1 | 0 | . | mm10 | chr7 | 97579936 | 97579936 |
| Zfp703 | 0 | . | mm10 | chr8 | 26977426 | 26977426 |
| Gab1 | 0 | . | mm10 | chr8 | 80791601 | 80791601 |

| | | | | | | |
|---|---|---|---|---|---|---|
| Zfc3h1 | 0 | . | mm10 | chr10 | 115385327 | 115385327 |
| Actg1 | 0 | . | mm10 | chr11 | 120347330 | 120347330 |
| Npm1 | 0 | . | mm10 | chr11 | 33152835 | 33152836 |
| Dsp | 0 | . | mm10 | chr13 | 38151597 | 38151597 |
| Mdc1 | 0 | . | mm10 | chr17 | 35847894 | 35847896 |
| Ankhd1 | 0 | . | mm10 | chr18 | 36560930 | 36560930 |
| Elf3 | 0 | . | mm10 | chr1 | 135257018 | 135257018 |
| Elf3 | 0 | . | mm10 | chr1 | 135257117 | 135257117 |
| Wt1 | 0 | . | mm10 | chr2 | 105127069 | 105127069 |
| Sptbn5 | 0 | . | mm10 | chr2 | 120047526 | 120047528 |
| Gm10800 | 0 | . | mm10 | chr2 | 98666667 | 98666667 |
| Arid1a | 0 | . | mm10 | chr4 | 133752569 | 133752571 |
| Mllt3 | 0 | . | mm10 | chr4 | 87841291 | 87841291 |
| Ptpn11 | 0 | . | mm10 | chr5 | 121167955 | 121167955 |
| Hectd4 | 0 | . | mm10 | chr5 | 121220505 | 121220507 |
| Setd1b | 0 | . | mm10 | chr5 | 123158559 | 123158559 |
| Gigyf1 | 0 | . | mm10 | chr5 | 137525323 | 137525325 |
| Flt3 | 0 | . | mm10 | chr5 | 147341238 | 147341238 |
| Kit | 0 | . | mm10 | chr5 | 75648432 | 75648432 |
| Maz | 0 | . | mm10 | chr7 | 127023125 | 127023125 |
| Maz | 0 | . | mm10 | chr7 | 127023125 | 127023126 |
| Setd1a | 0 | . | mm10 | chr7 | 127785317 | 127785317 |
| Fat1 | 0 | . | mm10 | chr8 | 45033500 | 45033500 |
| Ammecr1 | 0 | . | mm10 | chrX | 142966444 | 142966446 |
| Npm1 | 0 | . | mm10 | chr11 | 33152835 | 33152836 |
| Rpa1 | 0 | . | mm10 | chr11 | 75318489 | 75318491 |
| Vgll3 | 0 | . | mm10 | chr16 | 65827954 | 65827956 |
| Vegfa | 0 | . | mm10 | chr17 | 46025460 | 46025460 |
| Chka | 0 | . | mm10 | chr19 | 3852243 | 3852245 |
| Sptbn5 | 0 | . | mm10 | chr2 | 120047526 | 120047528 |
| Arid1a | 0 | . | mm10 | chr4 | 133752569 | 133752571 |
| Mllt3 | 0 | . | mm10 | chr4 | 87841342 | 87841342 |
| Ptpn11 | 0 | . | mm10 | chr5 | 121167955 | 121167955 |
| Maz | 0 | . | mm10 | chr7 | 127023125 | 127023125 |
| Maz | 0 | . | mm10 | chr7 | 127023125 | 127023126 |
| Srcap | 0 | . | mm10 | chr7 | 127538590 | 127538590 |
| Zfp703 | 0 | . | mm10 | chr8 | 26977426 | 26977426 |
| Fyn | 0 | . | mm10 | chr10 | 39563910 | 39563910 |
| Npm1 | 0 | . | mm10 | chr11 | 33152835 | 33152836 |
| Vgll3 | 0 | . | mm10 | chr16 | 65827954 | 65827956 |
| Erg | 0 | . | mm10 | chr16 | 95377323 | 95377323 |
| Sos1 | 0 | . | mm10 | chr17 | 80459934 | 80459934 |
| Olfr1463 | 0 | . | mm10 | chr19 | 13234889 | 13234889 |
| Olfr1463 | 0 | . | mm10 | chr19 | 13234890 | 13234890 |
| Elf3 | 0 | . | mm10 | chr1 | 135257018 | 135257018 |
| Elf3 | 0 | . | mm10 | chr1 | 135257117 | 135257117 |
| Sptbn5 | 0 | . | mm10 | chr2 | 120047526 | 120047528 |
| Prrc2b | 0 | . | mm10 | chr2 | 32206562 | 32206562 |
| Notch2 | 0 | . | mm10 | chr3 | 98146725 | 98146725 |
| Ptpn11 | 0 | . | mm10 | chr5 | 121167955 | 121167955 |

| | | | | | | |
|---|---|---|---|---|---|---|
| Gigyf1 | 0 | . | mm10 | chr5 | 137525323 | 137525325 |
| Flt3 | 0 | . | mm10 | chr5 | 147341238 | 147341238 |
| Maz | 0 | . | mm10 | chr7 | 127023125 | 127023125 |
| Maz | 0 | . | mm10 | chr7 | 127023125 | 127023126 |
| Grin2d | 0 | . | mm10 | chr7 | 45834480 | 45834480 |
| Ntrk3 | 0 | . | mm10 | chr7 | 78462871 | 78462871 |
| Zfp703 | 0 | . | mm10 | chr8 | 26977426 | 26977426 |
| Kmt2a | 0 | . | mm10 | chr9 | 44847876 | 44847876 |
| Ammecr1 | 0 | . | mm10 | chrX | 142966444 | 142966446 |
| Rbm10 | 0 | . | mm10 | chrX | 20637584 | 20637584 |
| Amer1 | 0 | . | mm10 | chrX | 95427365 | 95427365 |
| Ros1 | 0 | . | mm10 | chr10 | 52115994 | 52115994 |
| Npm1 | 0 | . | mm10 | chr11 | 33152835 | 33152836 |
| Rnf43 | 0 | . | mm10 | chr11 | 87731568 | 87731568 |
| Phlpp1 | 0 | . | mm10 | chr1 | 106393219 | 106393219 |
| Elf3 | 0 | . | mm10 | chr1 | 135257018 | 135257018 |
| Elf3 | 0 | . | mm10 | chr1 | 135257117 | 135257117 |
| Sptbn5 | 0 | . | mm10 | chr2 | 120047526 | 120047528 |
| Samhd1 | 0 | . | mm10 | chr2 | 157129903 | 157129903 |
| Mapkap1 | 0 | . | mm10 | chr2 | 34497744 | 34497744 |
| Arid1a | 0 | . | mm10 | chr4 | 133752569 | 133752571 |
| Ptpn11 | 0 | . | mm10 | chr5 | 121167955 | 121167955 |
| Gigyf1 | 0 | . | mm10 | chr5 | 137525323 | 137525325 |
| Flt3 | 0 | . | mm10 | chr5 | 147341238 | 147341238 |
| Rbm33 | 0 | . | mm10 | chr5 | 28394148 | 28394150 |
| Maz | 0 | . | mm10 | chr7 | 127023125 | 127023125 |
| Maz | 0 | . | mm10 | chr7 | 127023125 | 127023126 |
| Cdh16 | 0 | . | mm10 | chr8 | 104615215 | 104615215 |
| Ankrd11 | 0 | . | mm10 | chr8 | 122890181 | 122890181 |
| Ammecr1 | 0 | . | mm10 | chrX | 142966444 | 142966446 |
| Alg13 | 0 | . | mm10 | chrX | 144363899 | 144363901 |
| Npm1 | 0 | . | mm10 | chr11 | 33152835 | 33152836 |
| Tcf20 | 0 | . | mm10 | chr15 | 82856700 | 82856700 |
| Vgll3 | 0 | . | mm10 | chr16 | 65827954 | 65827956 |
| Ccnd3 | 0 | . | mm10 | chr17 | 47598127 | 47598127 |
| Olfr1463 | 0 | . | mm10 | chr19 | 13234889 | 13234889 |
| Olfr1463 | 0 | . | mm10 | chr19 | 13234890 | 13234890 |
| Chka | 0 | . | mm10 | chr19 | 3852243 | 3852245 |
| Elf3 | 0 | . | mm10 | chr1 | 135257018 | 135257018 |
| Elf3 | 0 | . | mm10 | chr1 | 135257117 | 135257117 |
| Sptbn5 | 0 | . | mm10 | chr2 | 120047526 | 120047528 |
| Ptpn11 | 0 | . | mm10 | chr5 | 121167955 | 121167955 |
| Gigyf1 | 0 | . | mm10 | chr5 | 137525323 | 137525325 |
| Flt3 | 0 | . | mm10 | chr5 | 147341238 | 147341238 |
| Maz | 0 | . | mm10 | chr7 | 127023125 | 127023125 |
| Maz | 0 | . | mm10 | chr7 | 127023125 | 127023126 |
| Zfhx3 | 0 | . | mm10 | chr8 | 108948478 | 108948478 |
| Tgfbr2 | 0 | . | mm10 | chr9 | 116110064 | 116110064 |
| Alg13 | 0 | . | mm10 | chrX | 144363899 | 144363901 |
| Npm1 | 0 | . | mm10 | chr11 | 33152835 | 33152836 |

| | | | | | | |
|---|---|---|---|---|---|---|
| Arid4b | 0 | . | mm10 | chr13 | 14187126 | 14187126 |
| Zfp503 | 0 | . | mm10 | chr14 | 21987453 | 21987455 |
| Rb1 | 0 | . | mm10 | chr14 | 73325591 | 73325591 |
| Ankhd1 | 0 | . | mm10 | chr18 | 36560930 | 36560930 |
| Elf3 | 0 | . | mm10 | chr1 | 135257018 | 135257018 |
| Elf3 | 0 | . | mm10 | chr1 | 135257117 | 135257117 |
| Itpkb | 0 | . | mm10 | chr1 | 180332597 | 180332597 |
| Sptbn5 | 0 | . | mm10 | chr2 | 120047526 | 120047528 |
| Ivl | 0 | . | mm10 | chr3 | 92572302 | 92572304 |
| Ptpn11 | 0 | . | mm10 | chr5 | 121167955 | 121167955 |
| Hectd4 | 0 | . | mm10 | chr5 | 121220505 | 121220507 |
| Gigyf1 | 0 | . | mm10 | chr5 | 137525323 | 137525325 |
| Cckar | 0 | . | mm10 | chr5 | 53700295 | 53700295 |
| Maz | 0 | . | mm10 | chr7 | 127023125 | 127023125 |
| Maz | 0 | . | mm10 | chr7 | 127023125 | 127023126 |
| Rsf1 | 0 | . | mm10 | chr7 | 97579936 | 97579936 |
| Cdh16 | 0 | . | mm10 | chr8 | 104615215 | 104615215 |
| Smarca4 | 0 | . | mm10 | chr9 | 21700957 | 21700957 |
| Ammecr1 | 0 | . | mm10 | chrX | 142966444 | 142966446 |
| Sry | 0 | . | mm10 | chrY | 2662864 | 2662864 |
| Sry | 0 | . | mm10 | chrY | 2663197 | 2663199 |
| Zfc3h1 | 0 | . | mm10 | chr10 | 115385309 | 115385311 |
| Lats1 | 0 | . | mm10 | chr10 | 7705538 | 7705538 |
| Npm1 | 0 | . | mm10 | chr11 | 33152835 | 33152836 |
| Hcn1 | 0 | . | mm10 | chr13 | 117975767 | 117975769 |
| Rb1 | 0 | . | mm10 | chr14 | 73325591 | 73325591 |
| Kmt2d | 0 | . | mm10 | chr15 | 98846475 | 98846475 |
| Vgll3 | 0 | . | mm10 | chr16 | 65827954 | 65827956 |
| Zfp318 | 0 | . | mm10 | chr17 | 46412515 | 46412517 |
| Serpinb3b | 0 | . | mm10 | chr1 | 107154541 | 107154541 |
| Elf3 | 0 | . | mm10 | chr1 | 135257018 | 135257018 |
| Elf3 | 0 | . | mm10 | chr1 | 135257117 | 135257117 |
| Sptbn5 | 0 | . | mm10 | chr2 | 120047526 | 120047528 |
| Lrp1b | 0 | . | mm10 | chr2 | 41468886 | 41468886 |
| Maml3 | 0 | . | mm10 | chr3 | 51856069 | 51856069 |
| Zfhx4 | 0 | . | mm10 | chr3 | 5400849 | 5400849 |
| Ptpn11 | 0 | . | mm10 | chr5 | 121167955 | 121167955 |
| Gigyf1 | 0 | . | mm10 | chr5 | 137525323 | 137525325 |
| Kit | 0 | . | mm10 | chr5 | 75649631 | 75649631 |
| Kras | 0 | . | mm10 | chr6 | 145246771 | 145246771 |
| Plk1 | 0 | . | mm10 | chr7 | 122167650 | 122167650 |
| Maz | 0 | . | mm10 | chr7 | 127023125 | 127023125 |
| Maz | 0 | . | mm10 | chr7 | 127023125 | 127023126 |
| Rsf1 | 0 | . | mm10 | chr7 | 97579936 | 97579936 |
| Mst1r | 0 | . | mm10 | chr9 | 107912134 | 107912134 |
| Npm1 | 0 | . | mm10 | chr11 | 33152835 | 33152836 |
| Lats2 | 0 | . | mm10 | chr14 | 57691464 | 57691464 |
| Vgll3 | 0 | . | mm10 | chr16 | 65827954 | 65827956 |
| Elf3 | 0 | . | mm10 | chr1 | 135257018 | 135257018 |
| Elf3 | 0 | . | mm10 | chr1 | 135257117 | 135257117 |

| | | | | | | |
|---|---|---|---|---|---|---|
| Sptbn5 | 0 | . | mm10 | chr2 | 120047526 | 120047528 |
| Samhd1 | 0 | . | mm10 | chr2 | 157129903 | 157129903 |
| Wdr63 | 0 | . | mm10 | chr3 | 146103165 | 146103165 |
| Ptpn11 | 0 | . | mm10 | chr5 | 121167955 | 121167955 |
| Gigyf1 | 0 | . | mm10 | chr5 | 137525323 | 137525325 |
| Flt3 | 0 | . | mm10 | chr5 | 147341238 | 147341238 |
| Maz | 0 | . | mm10 | chr7 | 127023125 | 127023125 |
| Maz | 0 | . | mm10 | chr7 | 127023125 | 127023126 |
| Srcap | 0 | . | mm10 | chr7 | 127538590 | 127538590 |
| Ntrk3 | 0 | . | mm10 | chr7 | 78356179 | 78356179 |
| Ankrd11 | 0 | . | mm10 | chr8 | 122890181 | 122890181 |
| Acta1 | 0 | . | mm10 | chr8 | 123893153 | 123893153 |
| Zfp703 | 0 | . | mm10 | chr8 | 26977426 | 26977426 |
| Zrsr2 | 0 | . | mm10 | chrX | 163936864 | 163936864 |
| Npm1 | 0 | . | mm10 | chr11 | 33152835 | 33152836 |
| Chd3 | 0 | . | mm10 | chr11 | 69369157 | 69369159 |
| Ep300 | 0 | . | mm10 | chr15 | 81613269 | 81613269 |
| Kmt2d | 0 | . | mm10 | chr15 | 98851067 | 98851102 |
| Robo1 | 0 | . | mm10 | chr16 | 73035926 | 73035926 |
| Sptbn5 | 0 | . | mm10 | chr2 | 120047526 | 120047528 |
| Samhd1 | 0 | . | mm10 | chr2 | 157129903 | 157129903 |
| Ivl | 0 | . | mm10 | chr3 | 92572302 | 92572304 |
| Ptpn11 | 0 | . | mm10 | chr5 | 121167955 | 121167955 |
| Gigyf1 | 0 | . | mm10 | chr5 | 137525323 | 137525325 |
| Flt3 | 0 | . | mm10 | chr5 | 147341238 | 147341238 |
| Rbm33 | 0 | . | mm10 | chr5 | 28394148 | 28394150 |
| Maz | 0 | . | mm10 | chr7 | 127023125 | 127023125 |
| Maz | 0 | . | mm10 | chr7 | 127023125 | 127023126 |
| Grin2d | 0 | . | mm10 | chr7 | 45834480 | 45834480 |
| Rsf1 | 0 | . | mm10 | chr7 | 97579910 | 97579910 |
| Ankrd11 | 0 | . | mm10 | chr8 | 122890181 | 122890181 |
| Zfp703 | 0 | . | mm10 | chr8 | 26977426 | 26977426 |
| Zmym3 | 0 | . | mm10 | chrX | 101418712 | 101418712 |
| Ammecr1 | 0 | . | mm10 | chrX | 142966444 | 142966446 |
| Rbm10 | 0 | . | mm10 | chrX | 20637584 | 20637584 |
| Amer1 | 0 | . | mm10 | chrX | 95427365 | 95427365 |
| Npm1 | 0 | . | mm10 | chr11 | 33152835 | 33152836 |
| Nf1 | 0 | . | mm10 | chr11 | 79453806 | 79453806 |
| Fam170b | 0 | . | mm10 | chr14 | 32835939 | 32835939 |
| Kmt2d | 0 | . | mm10 | chr15 | 98851067 | 98851102 |
| Arid1b | 0 | . | mm10 | chr17 | 5264048 | 5264048 |
| Sptbn5 | 0 | . | mm10 | chr2 | 120047526 | 120047528 |
| Ivl | 0 | . | mm10 | chr3 | 92572302 | 92572304 |
| Arid1a | 0 | . | mm10 | chr4 | 133752569 | 133752571 |
| Ptpn11 | 0 | . | mm10 | chr5 | 121167955 | 121167955 |
| Rbm33 | 0 | . | mm10 | chr5 | 28394148 | 28394150 |
| Kdm5a | 0 | . | mm10 | chr6 | 120427190 | 120427190 |
| D6Ertd527e | 0 | . | mm10 | chr6 | 87112125 | 87112125 |
| Maz | 0 | . | mm10 | chr7 | 127023125 | 127023125 |
| Maz | 0 | . | mm10 | chr7 | 127023125 | 127023126 |

| | | | | | | |
|---|---|---|---|---|---|---|
| Srcap | 0 | . | mm10 | chr7 | 127538590 | 127538590 |
| Grin2d | 0 | . | mm10 | chr7 | 45834480 | 45834480 |
| Kmt2a | 0 | . | mm10 | chr9 | 44819379 | 44819379 |
| Ammecr1 | 0 | . | mm10 | chrX | 142966444 | 142966446 |
| Alg13 | 0 | . | mm10 | chrX | 144363899 | 144363901 |
| Rbm10 | 0 | . | mm10 | chrX | 20637584 | 20637584 |

| Strand | Variant_Classi | Variant_Type | Reference_All | Tumor_Seq_A | Tumor_Seq_A | dbSNP_RS |
|---|---|---|---|---|---|---|
| + | Missense_Mu | SNP | A | A | G | novel |
| + | Frame_Shift_I | INS | - | - | CAGA | novel |
| + | Silent | SNP | A | A | G | novel |
| + | Missense_Mu | SNP | C | C | T | novel |
| + | In_Frame_Del | DEL | TGC | TGC | - | novel |
| + | Silent | SNP | G | G | T | novel |
| + | In_Frame_Del | DEL | TGC | TGC | - | novel |
| + | Missense_Mu | SNP | C | C | T | novel |
| + | In_Frame_Del | DEL | GCA | GCA | - | novel |
| + | Missense_Mu | SNP | T | T | C | novel |
| + | Missense_Mu | SNP | G | G | A | novel |
| + | Silent | SNP | C | C | T | novel |
| + | In_Frame_Ins | INS | - | - | GCT | novel |
| + | Missense_Mu | SNP | A | A | G | novel |
| + | Missense_Mu | SNP | G | G | A | novel |
| + | Missense_Mu | SNP | G | G | A | rs230103531 |
| + | Missense_Mu | SNP | C | C | G | novel |
| + | Missense_Mu | SNP | A | A | T | novel |
| + | Missense_Mu | SNP | G | G | T | novel |
| + | Missense_Mu | SNP | C | C | T | novel |
| + | Frame_Shift_I | INS | - | - | CAGA | novel |
| + | Missense_Mu | SNP | C | C | A | novel |
| + | Nonsense_Mu | SNP | C | C | T | novel |
| + | In_Frame_Del | DEL | GAG | GAG | - | novel |
| + | In_Frame_Del | DEL | GCT | GCT | - | novel |
| + | Missense_Mu | SNP | T | T | G | novel |
| + | In_Frame_Del | DEL | GCC | GCC | - | novel |
| + | Silent | SNP | C | C | T | novel |
| + | Missense_Mu | SNP | C | C | T | novel |
| + | Missense_Mu | SNP | T | T | C | novel |
| + | Silent | SNP | C | C | T | novel |
| + | In_Frame_Ins | INS | - | - | GCT | novel |
| + | Frame_Shift_I | INS | - | - | CAGA | novel |
| + | Frame_Shift_D | DEL | CGCTTTAAAA | CGCTTTAAAA | - | novel |
| + | Silent | SNP | C | C | T | novel |
| + | Missense_Mu | SNP | A | A | G | novel |
| + | In_Frame_Del | DEL | TGC | TGC | - | novel |
| + | In_Frame_Del | DEL | TGC | TGC | - | novel |
| + | Missense_Mu | SNP | C | C | T | novel |
| + | In_Frame_Del | DEL | GCA | GCA | - | novel |
| + | Silent | SNP | C | C | T | novel |
| + | Missense_Mu | SNP | C | C | T | rs239855953 |
| + | Silent | SNP | C | C | T | novel |
| + | In_Frame_Ins | INS | - | - | GCT | novel |
| + | Missense_Mu | SNP | G | G | A | novel |
| + | Missense_Mu | SNP | T | T | G | novel |
| + | In_Frame_Del | DEL | GCC | GCC | - | novel |
| + | Frame_Shift_I | INS | - | - | CAGA | novel |
| + | In_Frame_Del | DEL | CCG | CCG | - | novel |

| | | | | | | |
|---|---|---|---|---|---|---|
| + | In_Frame_Del | DEL | CAG | CAG | - | novel |
| + | Silent | SNP | C | C | T | novel |
| + | Missense_Mu | SNP | A | A | C | novel |
| + | Silent | SNP | G | G | A | novel |
| + | In_Frame_Del | DEL | TGC | TGC | - | novel |
| + | In_Frame_Del | DEL | GAG | GAG | - | novel |
| + | In_Frame_Del | DEL | TGC | TGC | - | novel |
| + | Silent | SNP | C | C | T | rs582628252 |
| + | In_Frame_Del | DEL | TGC | TGC | - | novel |
| + | Missense_Mu | SNP | C | C | T | novel |
| + | In_Frame_Del | DEL | GCA | GCA | - | novel |
| + | Missense_Mu | SNP | C | C | T | rs217659485 |
| + | Silent | SNP | C | C | T | novel |
| + | In_Frame_Ins | INS | - | - | GCT | novel |
| + | Missense_Mu | SNP | C | C | G | novel |
| + | Missense_Mu | SNP | T | T | G | novel |
| + | Missense_Mu | SNP | G | G | A | novel |
| + | In_Frame_Del | DEL | GCC | GCC | - | novel |
| + | Missense_Mu | SNP | G | G | T | novel |
| + | In_Frame_Del | DEL | CTG | CTG | - | novel |
| + | Frame_Shift_I | INS | - | - | CAGA | novel |
| + | Nonsense_Mu | SNP | C | C | T | novel |
| + | In_Frame_Del | DEL | CCG | CCG | - | novel |
| + | In_Frame_Del | DEL | GCC | GCC | - | novel |
| + | In_Frame_Del | DEL | GAG | GAG | - | novel |
| + | In_Frame_Del | DEL | TGC | TGC | - | novel |
| + | In_Frame_Del | DEL | CAG | CAG | - | novel |
| + | Missense_Mu | SNP | C | C | T | novel |
| + | In_Frame_Del | DEL | GCA | GCA | - | novel |
| + | Missense_Mu | SNP | T | T | C | novel |
| + | Missense_Mu | SNP | C | C | T | rs217659485 |
| + | In_Frame_Del | DEL | GCC | GCC | - | novel |
| + | Silent | SNP | G | G | A | rs578925286 |
| + | Missense_Mu | SNP | T | T | A | novel |
| + | Frame_Shift_I | INS | - | - | CAGA | novel |
| + | Silent | SNP | G | G | A | novel |
| + | Nonsense_Mu | SNP | G | G | T | novel |
| + | In_Frame_Del | DEL | CAG | CAG | - | novel |
| + | In_Frame_Del | DEL | TGC | TGC | - | novel |
| + | In_Frame_Del | DEL | TGC | TGC | - | novel |
| + | Silent | SNP | C | C | T | novel |
| + | Missense_Mu | SNP | C | C | T | novel |
| + | In_Frame_Del | DEL | GCA | GCA | - | novel |
| + | Silent | SNP | C | C | T | novel |
| + | Silent | SNP | C | C | T | novel |
| + | In_Frame_Ins | INS | - | - | GCT | novel |
| + | Missense_Mu | SNP | G | G | A | rs230103531 |
| + | Silent | SNP | A | A | G | novel |
| + | Missense_Mu | SNP | G | G | A | novel |
| + | Silent | SNP | C | C | T | novel |

| | | | | | | |
|---|---|---|---|---|---|---|
| + | Missense_Mu | SNP | A | A | G | novel |
| + | Missense_Mu | SNP | T | T | G | novel |
| + | Frame_Shift_I | INS | - | - | CAGA | novel |
| + | Nonsense_Mu | SNP | C | C | A | novel |
| + | In_Frame_Del | DEL | GAG | GAG | - | novel |
| + | Missense_Mu | SNP | G | G | A | novel |
| + | Silent | SNP | C | C | T | rs39394853 |
| + | Silent | SNP | G | G | A | rs49378976 |
| + | Silent | SNP | C | C | T | novel |
| + | In_Frame_Del | DEL | TGC | TGC | - | novel |
| + | Missense_Mu | SNP | G | G | C | novel |
| + | In_Frame_Del | DEL | GCC | GCC | - | novel |
| + | Silent | SNP | G | G | A | novel |
| + | Missense_Mu | SNP | C | C | T | novel |
| + | In_Frame_Del | DEL | GCG | GCG | - | novel |
| + | Silent | SNP | G | G | A | novel |
| + | In_Frame_Del | DEL | GCA | GCA | - | novel |
| + | Missense_Mu | SNP | T | T | C | novel |
| + | Missense_Mu | SNP | A | A | G | novel |
| + | Silent | SNP | C | C | T | novel |
| + | In_Frame_Ins | INS | - | - | GCT | novel |
| + | Missense_Mu | SNP | A | A | G | novel |
| + | Missense_Mu | SNP | A | A | G | novel |
| + | In_Frame_Del | DEL | GCC | GCC | - | novel |
| + | Frame_Shift_I | INS | - | - | CAGA | novel |
| + | In_Frame_Del | DEL | TGC | TGC | - | novel |
| + | In_Frame_Del | DEL | GAG | GAG | - | novel |
| + | Missense_Mu | SNP | C | C | T | novel |
| + | In_Frame_Del | DEL | GCC | GCC | - | novel |
| + | In_Frame_Del | DEL | TGC | TGC | - | novel |
| + | In_Frame_Del | DEL | GCC | GCC | - | novel |
| + | Silent | SNP | G | G | A | novel |
| + | Missense_Mu | SNP | C | C | T | novel |
| + | Silent | SNP | C | C | T | novel |
| + | In_Frame_Ins | INS | - | - | GCT | novel |
| + | Silent | SNP | T | T | C | novel |
| + | Missense_Mu | SNP | G | G | A | novel |
| + | Missense_Mu | SNP | G | G | A | novel |
| + | Frame_Shift_I | INS | - | - | CAGA | novel |
| + | In_Frame_Del | DEL | GAG | GAG | - | novel |
| + | Silent | SNP | C | C | A | novel |
| + | Silent | SNP | A | A | G | novel |
| + | Missense_Mu | SNP | T | T | A | novel |
| + | Silent | SNP | A | A | T | novel |
| + | Silent | SNP | C | C | T | rs39394853 |
| + | Silent | SNP | G | G | A | rs49378976 |
| + | In_Frame_Del | DEL | TGC | TGC | - | novel |
| + | Silent | SNP | G | G | A | novel |
| + | Missense_Mu | SNP | G | G | T | novel |
| + | Missense_Mu | SNP | C | C | T | novel |

| | | | | | | |
|---|---|---|---|---|---|---|
| + | In_Frame_Del | DEL | GCA | GCA | - | novel |
| + | Missense_Mu | SNP | T | T | C | novel |
| + | Silent | SNP | C | C | T | novel |
| + | In_Frame_Ins | INS | - | - | GCT | novel |
| + | Missense_Mu | SNP | A | A | C | novel |
| + | Missense_Mu | SNP | T | T | C | novel |
| + | Missense_Mu | SNP | G | G | A | novel |
| + | Missense_Mu | SNP | T | T | C | novel |
| + | In_Frame_Del | DEL | GCC | GCC | - | novel |
| + | Missense_Mu | SNP | G | G | T | novel |
| + | Missense_Mu | SNP | T | T | A | novel |
| + | Missense_Mu | SNP | C | C | A | novel |
| + | Frame_Shift_I | INS | - | - | CAGA | novel |
| + | Silent | SNP | A | A | C | novel |
| + | Missense_Mu | SNP | A | A | T | novel |
| + | Silent | SNP | C | C | T | rs39394853 |
| + | Silent | SNP | G | G | A | rs49378976 |
| + | In_Frame_Del | DEL | TGC | TGC | - | novel |
| + | Silent | SNP | G | G | T | novel |
| + | Missense_Mu | SNP | T | T | C | novel |
| + | In_Frame_Del | DEL | GCC | GCC | - | novel |
| + | Missense_Mu | SNP | C | C | T | novel |
| + | In_Frame_Del | DEL | GCA | GCA | - | novel |
| + | Missense_Mu | SNP | T | T | C | novel |
| + | In_Frame_Del | DEL | GCA | GCA | - | novel |
| + | Silent | SNP | C | C | T | novel |
| + | In_Frame_Ins | INS | - | - | GCT | novel |
| + | Missense_Mu | SNP | T | T | G | novel |
| + | Missense_Mu | SNP | C | C | G | novel |
| + | In_Frame_Del | DEL | GCC | GCC | - | novel |
| + | In_Frame_Del | DEL | AGC | AGC | - | novel |
| + | Frame_Shift_I | INS | - | - | CAGA | novel |
| + | Silent | SNP | C | C | T | rs1135367000 |
| + | In_Frame_Del | DEL | GAG | GAG | - | novel |
| + | Silent | SNP | A | A | G | novel |
| + | Missense_Mu | SNP | T | T | A | novel |
| + | Silent | SNP | A | A | T | novel |
| + | In_Frame_Del | DEL | GCC | GCC | - | novel |
| + | Silent | SNP | C | C | T | rs39394853 |
| + | Silent | SNP | G | G | A | rs49378976 |
| + | In_Frame_Del | DEL | TGC | TGC | - | novel |
| + | Missense_Mu | SNP | C | C | T | novel |
| + | In_Frame_Del | DEL | GCA | GCA | - | novel |
| + | Missense_Mu | SNP | T | T | C | novel |
| + | Silent | SNP | C | C | T | novel |
| + | In_Frame_Ins | INS | - | - | GCT | novel |
| + | Silent | SNP | T | T | C | novel |
| + | Missense_Mu | SNP | C | C | T | novel |
| + | In_Frame_Del | DEL | AGC | AGC | - | novel |
| + | Frame_Shift_I | INS | - | - | CAGA | novel |

| | | | | | | |
|---|---|---|---|---|---|---|
| + | Missense_Mu | SNP | C | C | A | novel |
| + | In_Frame_Del | DEL | GCC | GCC | - | novel |
| + | Silent | SNP | C | C | T | novel |
| + | Missense_Mu | SNP | G | G | A | novel |
| + | Silent | SNP | C | C | T | rs39394853 |
| + | Silent | SNP | G | G | A | rs49378976 |
| + | Missense_Mu | SNP | G | G | A | novel |
| + | In_Frame_Del | DEL | TGC | TGC | - | novel |
| + | In_Frame_Del | DEL | TGC | TGC | - | novel |
| + | Missense_Mu | SNP | C | C | T | novel |
| + | In_Frame_Del | DEL | GCG | GCG | - | novel |
| + | In_Frame_Del | DEL | GCA | GCA | - | novel |
| + | Missense_Mu | SNP | C | C | T | rs217659485 |
| + | Silent | SNP | C | C | T | novel |
| + | In_Frame_Ins | INS | - | - | GCT | novel |
| + | Silent | SNP | A | A | G | novel |
| + | Missense_Mu | SNP | T | T | G | novel |
| + | Missense_Mu | SNP | A | A | G | novel |
| + | In_Frame_Del | DEL | GCC | GCC | - | novel |
| + | Missense_Mu | SNP | C | C | G | novel |
| + | In_Frame_Del | DEL | CTG | CTG | - | novel |
| + | In_Frame_Del | DEL | AGC | AGC | - | novel |
| + | Missense_Mu | SNP | A | A | G | novel |
| + | Frame_Shift_I | INS | - | - | CAGA | novel |
| + | In_Frame_Del | DEL | CAG | CAG | - | novel |
| + | Silent | SNP | C | C | T | novel |
| + | Silent | SNP | C | C | T | novel |
| + | In_Frame_Del | DEL | GAG | GAG | - | novel |
| + | In_Frame_Del | DEL | GAA | GAA | - | novel |
| + | Missense_Mu | SNP | A | A | G | novel |
| + | Silent | SNP | C | C | T | rs39394853 |
| + | Silent | SNP | G | G | A | rs49378976 |
| + | In_Frame_Del | DEL | TGC | TGC | - | novel |
| + | Silent | SNP | C | C | T | novel |
| + | Silent | SNP | C | C | T | rs582628252 |
| + | Silent | SNP | T | T | A | novel |
| + | Missense_Mu | SNP | C | C | T | novel |
| + | In_Frame_Del | DEL | GCA | GCA | - | novel |
| + | Missense_Mu | SNP | G | G | T | novel |
| + | Missense_Mu | SNP | C | C | T | novel |
| + | Missense_Mu | SNP | C | C | T | rs239855953 |
| + | Silent | SNP | C | C | T | novel |
| + | In_Frame_Ins | INS | - | - | GCT | novel |
| + | Silent | SNP | A | A | G | novel |
| + | Silent | SNP | T | T | G | novel |
| + | Frame_Shift_I | INS | - | - | CAGA | novel |
| + | Missense_Mu | SNP | A | A | G | novel |
| + | In_Frame_Del | DEL | GAG | GAG | - | novel |
| + | Silent | SNP | C | C | T | rs39394853 |
| + | Silent | SNP | G | G | A | rs49378976 |

| | | | | | | |
|---|---|---|---|---|---|---|
| + | In_Frame_Del | DEL | TGC | TGC | - | novel |
| + | Silent | SNP | G | G | T | novel |
| + | Missense_Mu | SNP | T | T | C | novel |
| + | Missense_Mu | SNP | C | C | T | novel |
| + | In_Frame_Del | DEL | GCA | GCA | - | novel |
| + | Missense_Mu | SNP | T | T | C | novel |
| + | Silent | SNP | C | C | T | novel |
| + | In_Frame_Ins | INS | - | - | GCT | novel |
| + | Silent | SNP | T | T | C | novel |
| + | Missense_Mu | SNP | G | G | A | novel |
| + | Missense_Mu | SNP | C | C | G | novel |
| + | Silent | SNP | A | A | C | novel |
| + | Missense_Mu | SNP | G | G | A | novel |
| + | Missense_Mu | SNP | A | A | C | novel |
| + | Frame_Shift_I | INS | - | - | CAGA | novel |
| + | In_Frame_Del | DEL | GCG | GCG | - | novel |
| + | Missense_Mu | SNP | C | C | A | novel |
| + | In_Frame_Del | DEL | TGCTGCTGCT( | TGCTGCTGCT( | - | novel |
| + | Missense_Mu | SNP | C | C | T | novel |
| + | In_Frame_Del | DEL | TGC | TGC | - | novel |
| + | Silent | SNP | G | G | T | novel |
| + | In_Frame_Del | DEL | TGC | TGC | - | novel |
| + | Missense_Mu | SNP | C | C | T | novel |
| + | In_Frame_Del | DEL | GCA | GCA | - | novel |
| + | Missense_Mu | SNP | T | T | C | novel |
| + | In_Frame_Del | DEL | GCA | GCA | - | novel |
| + | Silent | SNP | C | C | T | novel |
| + | In_Frame_Ins | INS | - | - | GCT | novel |
| + | Missense_Mu | SNP | A | A | C | novel |
| + | Missense_Mu | SNP | G | G | A | rs230103531 |
| + | Missense_Mu | SNP | C | C | G | novel |
| + | Missense_Mu | SNP | G | G | A | novel |
| + | Silent | SNP | T | T | C | novel |
| + | In_Frame_Del | DEL | GCC | GCC | - | novel |
| + | Missense_Mu | SNP | G | G | T | novel |
| + | Missense_Mu | SNP | T | T | A | novel |
| + | Frame_Shift_I | INS | - | - | CAGA | novel |
| + | Nonsense_Mu | SNP | G | G | T | novel |
| + | Missense_Mu | SNP | A | A | C | novel |
| + | In_Frame_Del | DEL | TGCTGCTGCT( | TGCTGCTGCT( | - | novel |
| + | Silent | SNP | G | G | T | novel |
| + | In_Frame_Del | DEL | TGC | TGC | - | novel |
| + | In_Frame_Del | DEL | TGC | TGC | - | novel |
| + | In_Frame_Del | DEL | GCC | GCC | - | novel |
| + | Missense_Mu | SNP | C | C | T | novel |
| + | In_Frame_Del | DEL | GCA | GCA | - | novel |
| + | Missense_Mu | SNP | G | G | A | novel |
| + | Silent | SNP | C | C | T | novel |
| + | Silent | SNP | C | C | T | novel |
| + | In_Frame_Ins | INS | - | - | GCT | novel |

| | | | | | | |
|---|---|---|---|---|---|---|
| + | Silent | SNP | T | T | C | novel |
| + | Missense_Mu | SNP | A | A | C | novel |
| + | Missense_Mu | SNP | G | G | T | novel |
| + | In_Frame_Del | DEL | GCC | GCC | - | novel |
| + | In_Frame_Del | DEL | AGC | AGC | - | novel |
| + | Missense_Mu | SNP | G | G | T | novel |

| dbSNP_Val_St | Tumor_Sampl | Matched_Nor | Match_Norm_ | Match_Norm_ | Tumor_Valida | Tumor_Valida |
|---|---|---|---|---|---|---|
| | s_S0165 | s_Mouse-Pool | A | A | | |
| | s_S0165 | s_Mouse-Pool | - | - | | |
| | s_S0165 | s_Mouse-Pool | A | A | | |
| | s_S0165 | s_Mouse-Pool | C | C | | |
| | s_S0165 | s_Mouse-Pool | TGC | TGC | | |
| | s_S0165 | s_Mouse-Pool | G | G | | |
| | s_S0165 | s_Mouse-Pool | TGC | TGC | | |
| | s_S0165 | s_Mouse-Pool | C | C | | |
| | s_S0165 | s_Mouse-Pool | GCA | GCA | | |
| | s_S0165 | s_Mouse-Pool | T | T | | |
| | s_S0165 | s_Mouse-Pool | G | G | | |
| | s_S0165 | s_Mouse-Pool | C | C | | |
| | s_S0165 | s_Mouse-Pool | - | - | | |
| | s_S0165 | s_Mouse-Pool | A | A | | |
| | s_S0165 | s_Mouse-Pool | G | G | | |
| | s_S0165 | s_Mouse-Pool | G | G | | |
| | s_S0165 | s_Mouse-Pool | C | C | | |
| | s_S0165 | s_Mouse-Pool | A | A | | |
| | s_S0165 | s_Mouse-Pool | G | G | | |
| | s_S0200 | s_Mouse-Pool | C | C | | |
| | s_S0200 | s_Mouse-Pool | - | - | | |
| | s_S0200 | s_Mouse-Pool | C | C | | |
| | s_S0200 | s_Mouse-Pool | C | C | | |
| | s_S0200 | s_Mouse-Pool | GAG | GAG | | |
| | s_S0200 | s_Mouse-Pool | GCT | GCT | | |
| | s_S0200 | s_Mouse-Pool | T | T | | |
| | s_S0200 | s_Mouse-Pool | GCC | GCC | | |
| | s_S0200 | s_Mouse-Pool | C | C | | |
| | s_S0200 | s_Mouse-Pool | C | C | | |
| | s_S0200 | s_Mouse-Pool | T | T | | |
| | s_S0200 | s_Mouse-Pool | C | C | | |
| | s_S0200 | s_Mouse-Pool | - | - | | |
| | s_S0283 | s_Mouse-Pool | - | - | | |
| | s_S0283 | s_Mouse-Pool | CGCTTTAAAA | CGCTTTAAAA | | |
| | s_S0283 | s_Mouse-Pool | C | C | | |
| | s_S0283 | s_Mouse-Pool | A | A | | |
| | s_S0283 | s_Mouse-Pool | TGC | TGC | | |
| | s_S0283 | s_Mouse-Pool | TGC | TGC | | |
| | s_S0283 | s_Mouse-Pool | C | C | | |
| | s_S0283 | s_Mouse-Pool | GCA | GCA | | |
| | s_S0283 | s_Mouse-Pool | C | C | | |
| | s_S0283 | s_Mouse-Pool | C | C | | |
| | s_S0283 | s_Mouse-Pool | C | C | | |
| | s_S0283 | s_Mouse-Pool | - | - | | |
| | s_S0283 | s_Mouse-Pool | G | G | | |
| | s_S0283 | s_Mouse-Pool | T | T | | |
| | s_S0283 | s_Mouse-Pool | GCC | GCC | | |
| | s_S0437 | s_Mouse-Pool | - | - | | |
| | s_S0437 | s_Mouse-Pool | CCG | CCG | | |

| | | | |
|---|---|---|---|
| s_S0437 | s_Mouse-Pool | CAG | CAG |
| s_S0437 | s_Mouse-Pool | C | C |
| s_S0437 | s_Mouse-Pool | A | A |
| s_S0437 | s_Mouse-Pool | G | G |
| s_S0437 | s_Mouse-Pool | TGC | TGC |
| s_S0437 | s_Mouse-Pool | GAG | GAG |
| s_S0437 | s_Mouse-Pool | TGC | TGC |
| s_S0437 | s_Mouse-Pool | C | C |
| s_S0437 | s_Mouse-Pool | TGC | TGC |
| s_S0437 | s_Mouse-Pool | C | C |
| s_S0437 | s_Mouse-Pool | GCA | GCA |
| s_S0437 | s_Mouse-Pool | C | C |
| s_S0437 | s_Mouse-Pool | C | C |
| s_S0437 | s_Mouse-Pool | - | - |
| s_S0437 | s_Mouse-Pool | C | C |
| s_S0437 | s_Mouse-Pool | T | T |
| s_S0437 | s_Mouse-Pool | G | G |
| s_S0437 | s_Mouse-Pool | GCC | GCC |
| s_S0437 | s_Mouse-Pool | G | G |
| s_S0437 | s_Mouse-Pool | CTG | CTG |
| s_S0518 | s_Mouse-Pool | - | - |
| s_S0518 | s_Mouse-Pool | C | C |
| s_S0518 | s_Mouse-Pool | CCG | CCG |
| s_S0518 | s_Mouse-Pool | GCC | GCC |
| s_S0518 | s_Mouse-Pool | GAG | GAG |
| s_S0518 | s_Mouse-Pool | TGC | TGC |
| s_S0518 | s_Mouse-Pool | CAG | CAG |
| s_S0518 | s_Mouse-Pool | C | C |
| s_S0518 | s_Mouse-Pool | GCA | GCA |
| s_S0518 | s_Mouse-Pool | T | T |
| s_S0518 | s_Mouse-Pool | C | C |
| s_S0518 | s_Mouse-Pool | GCC | GCC |
| s_S0518 | s_Mouse-Pool | G | G |
| s_S0518 | s_Mouse-Pool | T | T |
| s_S0521 | s_Mouse-Pool | - | - |
| s_S0521 | s_Mouse-Pool | G | G |
| s_S0521 | s_Mouse-Pool | G | G |
| s_S0521 | s_Mouse-Pool | CAG | CAG |
| s_S0521 | s_Mouse-Pool | TGC | TGC |
| s_S0521 | s_Mouse-Pool | TGC | TGC |
| s_S0521 | s_Mouse-Pool | C | C |
| s_S0521 | s_Mouse-Pool | C | C |
| s_S0521 | s_Mouse-Pool | GCA | GCA |
| s_S0521 | s_Mouse-Pool | C | C |
| s_S0521 | s_Mouse-Pool | C | C |
| s_S0521 | s_Mouse-Pool | - | - |
| s_S0521 | s_Mouse-Pool | G | G |
| s_S0521 | s_Mouse-Pool | A | A |
| s_S0521 | s_Mouse-Pool | G | G |
| s_S0521 | s_Mouse-Pool | C | C |

| | | | |
|---|---|---|---|
| s_Y0666 | s_Mouse-Pool | A | A |
| s_Y0666 | s_Mouse-Pool | T | T |
| s_Y0666 | s_Mouse-Pool | - | - |
| s_Y0666 | s_Mouse-Pool | C | C |
| s_Y0666 | s_Mouse-Pool | GAG | GAG |
| s_Y0666 | s_Mouse-Pool | G | G |
| s_Y0666 | s_Mouse-Pool | C | C |
| s_Y0666 | s_Mouse-Pool | G | G |
| s_Y0666 | s_Mouse-Pool | C | C |
| s_Y0666 | s_Mouse-Pool | TGC | TGC |
| s_Y0666 | s_Mouse-Pool | G | G |
| s_Y0666 | s_Mouse-Pool | GCC | GCC |
| s_Y0666 | s_Mouse-Pool | G | G |
| s_Y0666 | s_Mouse-Pool | C | C |
| s_Y0666 | s_Mouse-Pool | GCG | GCG |
| s_Y0666 | s_Mouse-Pool | G | G |
| s_Y0666 | s_Mouse-Pool | GCA | GCA |
| s_Y0666 | s_Mouse-Pool | T | T |
| s_Y0666 | s_Mouse-Pool | A | A |
| s_Y0666 | s_Mouse-Pool | C | C |
| s_Y0666 | s_Mouse-Pool | - | - |
| s_Y0666 | s_Mouse-Pool | A | A |
| s_Y0666 | s_Mouse-Pool | A | A |
| s_Y0666 | s_Mouse-Pool | GCC | GCC |
| s_Y0670 | s_Mouse-Pool | - | - |
| s_Y0670 | s_Mouse-Pool | TGC | TGC |
| s_Y0670 | s_Mouse-Pool | GAG | GAG |
| s_Y0670 | s_Mouse-Pool | C | C |
| s_Y0670 | s_Mouse-Pool | GCC | GCC |
| s_Y0670 | s_Mouse-Pool | TGC | TGC |
| s_Y0670 | s_Mouse-Pool | GCC | GCC |
| s_Y0670 | s_Mouse-Pool | G | G |
| s_Y0670 | s_Mouse-Pool | C | C |
| s_Y0670 | s_Mouse-Pool | C | C |
| s_Y0670 | s_Mouse-Pool | - | - |
| s_Y0670 | s_Mouse-Pool | T | T |
| s_Y0670 | s_Mouse-Pool | G | G |
| s_Y0671 | s_Mouse-Pool | G | G |
| s_Y0671 | s_Mouse-Pool | - | - |
| s_Y0671 | s_Mouse-Pool | GAG | GAG |
| s_Y0671 | s_Mouse-Pool | C | C |
| s_Y0671 | s_Mouse-Pool | A | A |
| s_Y0671 | s_Mouse-Pool | T | T |
| s_Y0671 | s_Mouse-Pool | A | A |
| s_Y0671 | s_Mouse-Pool | C | C |
| s_Y0671 | s_Mouse-Pool | G | G |
| s_Y0671 | s_Mouse-Pool | TGC | TGC |
| s_Y0671 | s_Mouse-Pool | G | G |
| s_Y0671 | s_Mouse-Pool | G | G |
| s_Y0671 | s_Mouse-Pool | C | C |

| | | | |
|---|---|---|---|
| s_Y0671 | s_Mouse-Pool | GCA | GCA |
| s_Y0671 | s_Mouse-Pool | T | T |
| s_Y0671 | s_Mouse-Pool | C | C |
| s_Y0671 | s_Mouse-Pool | - | - |
| s_Y0671 | s_Mouse-Pool | A | A |
| s_Y0671 | s_Mouse-Pool | T | T |
| s_Y0671 | s_Mouse-Pool | G | G |
| s_Y0671 | s_Mouse-Pool | T | T |
| s_Y0671 | s_Mouse-Pool | GCC | GCC |
| s_Y0671 | s_Mouse-Pool | G | G |
| s_Y0671 | s_Mouse-Pool | T | T |
| s_Y0672 | s_Mouse-Pool | C | C |
| s_Y0672 | s_Mouse-Pool | - | - |
| s_Y0672 | s_Mouse-Pool | A | A |
| s_Y0672 | s_Mouse-Pool | A | A |
| s_Y0672 | s_Mouse-Pool | C | C |
| s_Y0672 | s_Mouse-Pool | G | G |
| s_Y0672 | s_Mouse-Pool | TGC | TGC |
| s_Y0672 | s_Mouse-Pool | G | G |
| s_Y0672 | s_Mouse-Pool | T | T |
| s_Y0672 | s_Mouse-Pool | GCC | GCC |
| s_Y0672 | s_Mouse-Pool | C | C |
| s_Y0672 | s_Mouse-Pool | GCA | GCA |
| s_Y0672 | s_Mouse-Pool | T | T |
| s_Y0672 | s_Mouse-Pool | GCA | GCA |
| s_Y0672 | s_Mouse-Pool | C | C |
| s_Y0672 | s_Mouse-Pool | - | - |
| s_Y0672 | s_Mouse-Pool | T | T |
| s_Y0672 | s_Mouse-Pool | C | C |
| s_Y0672 | s_Mouse-Pool | GCC | GCC |
| s_Y0672 | s_Mouse-Pool | AGC | AGC |
| s_Y0674 | s_Mouse-Pool | - | - |
| s_Y0674 | s_Mouse-Pool | C | C |
| s_Y0674 | s_Mouse-Pool | GAG | GAG |
| s_Y0674 | s_Mouse-Pool | A | A |
| s_Y0674 | s_Mouse-Pool | T | T |
| s_Y0674 | s_Mouse-Pool | A | A |
| s_Y0674 | s_Mouse-Pool | GCC | GCC |
| s_Y0674 | s_Mouse-Pool | C | C |
| s_Y0674 | s_Mouse-Pool | G | G |
| s_Y0674 | s_Mouse-Pool | TGC | TGC |
| s_Y0674 | s_Mouse-Pool | C | C |
| s_Y0674 | s_Mouse-Pool | GCA | GCA |
| s_Y0674 | s_Mouse-Pool | T | T |
| s_Y0674 | s_Mouse-Pool | C | C |
| s_Y0674 | s_Mouse-Pool | - | - |
| s_Y0674 | s_Mouse-Pool | T | T |
| s_Y0674 | s_Mouse-Pool | C | C |
| s_Y0674 | s_Mouse-Pool | AGC | AGC |
| s_Y0677 | s_Mouse-Pool | - | - |

| | | |
|---|---|---|
| s_Y0677 | s_Mouse-Pool C | C |
| s_Y0677 | s_Mouse-Pool GCC | GCC |
| s_Y0677 | s_Mouse-Pool C | C |
| s_Y0677 | s_Mouse-Pool G | G |
| s_Y0677 | s_Mouse-Pool C | C |
| s_Y0677 | s_Mouse-Pool G | G |
| s_Y0677 | s_Mouse-Pool G | G |
| s_Y0677 | s_Mouse-Pool TGC | TGC |
| s_Y0677 | s_Mouse-Pool TGC | TGC |
| s_Y0677 | s_Mouse-Pool C | C |
| s_Y0677 | s_Mouse-Pool GCG | GCG |
| s_Y0677 | s_Mouse-Pool GCA | GCA |
| s_Y0677 | s_Mouse-Pool C | C |
| s_Y0677 | s_Mouse-Pool C | C |
| s_Y0677 | s_Mouse-Pool - | - |
| s_Y0677 | s_Mouse-Pool A | A |
| s_Y0677 | s_Mouse-Pool T | T |
| s_Y0677 | s_Mouse-Pool A | A |
| s_Y0677 | s_Mouse-Pool GCC | GCC |
| s_Y0677 | s_Mouse-Pool C | C |
| s_Y0677 | s_Mouse-Pool CTG | CTG |
| s_Y0679 | s_Mouse-Pool AGC | AGC |
| s_Y0679 | s_Mouse-Pool A | A |
| s_Y0679 | s_Mouse-Pool - | - |
| s_Y0679 | s_Mouse-Pool CAG | CAG |
| s_Y0679 | s_Mouse-Pool C | C |
| s_Y0679 | s_Mouse-Pool C | C |
| s_Y0679 | s_Mouse-Pool GAG | GAG |
| s_Y0679 | s_Mouse-Pool GAA | GAA |
| s_Y0679 | s_Mouse-Pool A | A |
| s_Y0679 | s_Mouse-Pool C | C |
| s_Y0679 | s_Mouse-Pool G | G |
| s_Y0679 | s_Mouse-Pool TGC | TGC |
| s_Y0679 | s_Mouse-Pool C | C |
| s_Y0679 | s_Mouse-Pool C | C |
| s_Y0679 | s_Mouse-Pool T | T |
| s_Y0679 | s_Mouse-Pool C | C |
| s_Y0679 | s_Mouse-Pool GCA | GCA |
| s_Y0679 | s_Mouse-Pool G | G |
| s_Y0679 | s_Mouse-Pool C | C |
| s_Y0679 | s_Mouse-Pool C | C |
| s_Y0679 | s_Mouse-Pool C | C |
| s_Y0679 | s_Mouse-Pool - | - |
| s_Y0679 | s_Mouse-Pool A | A |
| s_Y0679 | s_Mouse-Pool T | T |
| s_Y0887 | s_Mouse-Pool - | - |
| s_Y0887 | s_Mouse-Pool A | A |
| s_Y0887 | s_Mouse-Pool GAG | GAG |
| s_Y0887 | s_Mouse-Pool C | C |
| s_Y0887 | s_Mouse-Pool G | G |

| | | | |
|---|---|---|---|
| s_Y0887 | s_Mouse-Pool | TGC | TGC |
| s_Y0887 | s_Mouse-Pool | G | G |
| s_Y0887 | s_Mouse-Pool | T | T |
| s_Y0887 | s_Mouse-Pool | C | C |
| s_Y0887 | s_Mouse-Pool | GCA | GCA |
| s_Y0887 | s_Mouse-Pool | T | T |
| s_Y0887 | s_Mouse-Pool | C | C |
| s_Y0887 | s_Mouse-Pool | - | - |
| s_Y0887 | s_Mouse-Pool | T | T |
| s_Y0887 | s_Mouse-Pool | G | G |
| s_Y0887 | s_Mouse-Pool | C | C |
| s_Y0887 | s_Mouse-Pool | A | A |
| s_Y0887 | s_Mouse-Pool | G | G |
| s_Y0887 | s_Mouse-Pool | A | A |
| s_Y0888 | s_Mouse-Pool | - | - |
| s_Y0888 | s_Mouse-Pool | GCG | GCG |
| s_Y0888 | s_Mouse-Pool | C | C |
| s_Y0888 | s_Mouse-Pool | TGCTGCTGCT( | TGCTGCTGCT( |
| s_Y0888 | s_Mouse-Pool | C | C |
| s_Y0888 | s_Mouse-Pool | TGC | TGC |
| s_Y0888 | s_Mouse-Pool | G | G |
| s_Y0888 | s_Mouse-Pool | TGC | TGC |
| s_Y0888 | s_Mouse-Pool | C | C |
| s_Y0888 | s_Mouse-Pool | GCA | GCA |
| s_Y0888 | s_Mouse-Pool | T | T |
| s_Y0888 | s_Mouse-Pool | GCA | GCA |
| s_Y0888 | s_Mouse-Pool | C | C |
| s_Y0888 | s_Mouse-Pool | - | - |
| s_Y0888 | s_Mouse-Pool | A | A |
| s_Y0888 | s_Mouse-Pool | G | G |
| s_Y0888 | s_Mouse-Pool | C | C |
| s_Y0888 | s_Mouse-Pool | G | G |
| s_Y0888 | s_Mouse-Pool | T | T |
| s_Y0888 | s_Mouse-Pool | GCC | GCC |
| s_Y0888 | s_Mouse-Pool | G | G |
| s_Y0888 | s_Mouse-Pool | T | T |
| s_Y0909 | s_Mouse-Pool | - | - |
| s_Y0909 | s_Mouse-Pool | G | G |
| s_Y0909 | s_Mouse-Pool | A | A |
| s_Y0909 | s_Mouse-Pool | TGCTGCTGCT( | TGCTGCTGCT( |
| s_Y0909 | s_Mouse-Pool | G | G |
| s_Y0909 | s_Mouse-Pool | TGC | TGC |
| s_Y0909 | s_Mouse-Pool | TGC | TGC |
| s_Y0909 | s_Mouse-Pool | GCC | GCC |
| s_Y0909 | s_Mouse-Pool | C | C |
| s_Y0909 | s_Mouse-Pool | GCA | GCA |
| s_Y0909 | s_Mouse-Pool | G | G |
| s_Y0909 | s_Mouse-Pool | C | C |
| s_Y0909 | s_Mouse-Pool | C | C |
| s_Y0909 | s_Mouse-Pool | - | - |

| | | | |
|---|---|---|---|
| s_Y0909 | s_Mouse-Pool | T | T |
| s_Y0909 | s_Mouse-Pool | A | A |
| s_Y0909 | s_Mouse-Pool | G | G |
| s_Y0909 | s_Mouse-Pool | GCC | GCC |
| s_Y0909 | s_Mouse-Pool | AGC | AGC |
| s_Y0909 | s_Mouse-Pool | G | G |

| Match_Norm_ | Match_Norm_ | Verification_S | Validation_Sta | Mutation_Stat | Sequencing_P | Sequence_Sou |
|---|---|---|---|---|---|---|

| Validation_Me | Score | BAM_File | Sequencer | Tumor_Sampl | Matched_Nor | HGVSc |
|---|---|---|---|---|---|---|
| | | | | | | c.460T>C |
| | | | | | | c.854_857dup |
| | | | | | | c.4239A>G |
| | | | | | | c.643G>A |
| | | | | | | c.9906_9908d |
| | | | | | | c.318C>A |
| | | | | | | c.453_455del |
| | | | | | | c.226G>A |
| | | | | | | c.2950_2952d |
| | | | | | | c.2525A>G |
| | | | | | | c.1123G>A |
| | | | | | | c.1323G>A |
| | | | | | | c.1320_1322d |
| | | | | | | c.1339A>G |
| | | | | | | c.61G>A |
| | | | | | | c.7G>A |
| | | | | | | c.6868G>C |
| | | | | | | c.1105T>A |
| | | | | | | c.360G>T |
| | | | | | | c.943C>T |
| | | | | | | c.854_857dup |
| | | | | | | c.4345C>A |
| | | | | | | c.4432C>T |
| | | | | | | c.210_212del |
| | | | | | | c.266_268del |
| | | | | | | c.190A>C |
| | | | | | | c.1041_1043d |
| | | | | | | c.1644G>A |
| | | | | | | c.226G>A |
| | | | | | | c.2525A>G |
| | | | | | | c.1323G>A |
| | | | | | | c.1320_1322d |
| | | | | | | c.854_857dup |
| | | | | | | c.4964_4973d |
| | | | | | | c.51G>A |
| | | | | | | c.992T>C |
| | | | | | | c.9906_9908d |
| | | | | | | c.453_455del |
| | | | | | | c.226G>A |
| | | | | | | c.2950_2952d |
| | | | | | | c.1269C>T |
| | | | | | | c.1090C>T |
| | | | | | | c.1323G>A |
| | | | | | | c.1320_1322d |
| | | | | | | c.181G>A |
| | | | | | | c.2317A>C |
| | | | | | | c.268_270del |
| | | | | | | c.854_857dup |
| | | | | | | c.78_80del |

c.2324_2326d
c.15G>A
c.730A>C
c.435C>T
c.9852_9854d
c.210_212del
c.9906_9908d
c.1473G>A
c.453_455del
c.226G>A
c.2667_2669d
c.772G>A
c.1323G>A
c.1320_1322d
c.7354C>G
c.2317A>C
c.73G>A
c.268_270del
c.360G>T
c.712_714del
c.854_857dup
c.6811C>T
c.78_80del
c.79_81del
c.210_212del
c.9906_9908d
c.2752_2754d
c.226G>A
c.2950_2952d
c.2525A>G
c.772G>A
c.268_270del
c.1467G>A
c.1146A>T
c.854_857dup
c.51G>A
c.3718G>T
c.2324_2326d
c.12144_1214
c.9906_9908d
c.7956G>A
c.226G>A
c.2667_2669d
c.1269C>T
c.1323G>A
c.1320_1322d
c.7G>A
c.33A>G
c.73G>A
c.381G>A

c.130A>G
c.383A>C
c.854_857dup
c.9C>A
c.1190_1192d
c.193G>A
c.429G>A
c.330C>T
c.273C>T
c.9906_9908d
c.540C>G
c.1041_1043d
c.519C>T
c.226G>A
c.53_55del
c.3582G>A
c.2950_2952d
c.2525A>G
c.2285A>G
c.1323G>A
c.1320_1322d
c.1339A>G
c.10021A>G
c.268_270del
c.854_857dup
c.465_467del
c.210_212del
c.733G>A
c.186_188del
c.9906_9908d
c.1041_1043d
c.468C>T
c.226G>A
c.1323G>A
c.1320_1322d
c.3111T>C
c.73G>A
c.1552G>A
c.854_857dup
c.210_212del
c.744G>T
c.201T>C
c.638T>A
c.639A>T
c.429G>A
c.330C>T
c.9906_9908d
c.1944G>A
c.6703G>T
c.226G>A

c.2950_2952d
c.2525A>G
c.1323G>A
c.1320_1322d
c.2433T>G
c.536A>G
c.73G>A
c.2675A>G
c.268_270del
c.360G>T
c.1146A>T
c.4195G>T
c.854_857dup
c.1494A>C
c.4943A>T
c.429G>A
c.330C>T
c.9906_9908d
c.318C>A
c.542T>C
c.1041_1043d
c.226G>A
c.2950_2952d
c.2525A>G
c.2667_2669d
c.1323G>A
c.1320_1322d
c.2317A>C
c.6868G>C
c.268_270del
c.2131_2133d
c.854_857dup
c.549G>A
c.210_212del
c.699A>G
c.638T>A
c.639A>T
c.186_188del
c.429G>A
c.330C>T
c.9906_9908d
c.226G>A
c.2950_2952d
c.2525A>G
c.1323G>A
c.1320_1322d
c.6159T>C
c.769G>A
c.2131_2133d
c.854_857dup

c.2435C>A
c.79_81del
c.51G>A
c.193G>A
c.429G>A
c.330C>T
c.287G>A
c.9906_9908d
c.453_455del
c.226G>A
c.53_55del
c.2950_2952d
c.772G>A
c.1323G>A
c.1320_1322d
c.33A>G
c.2317A>C
c.4649A>G
c.268_270del
c.795G>C
c.460_462del
c.130_132del
c.2086A>G
c.854_857dup
c.2324_2326d
c.51G>A
c.10803G>A
c.210_212del
c.5468_5470d
c.992T>C
c.429G>A
c.330C>T
c.9906_9908d
c.2316G>A
c.1473G>A
c.6066T>A
c.226G>A
c.2950_2952d
c.2452G>T
c.35G>A
c.1090C>T
c.1323G>A
c.1320_1322d
c.33A>G
c.1611T>G
c.854_857dup
c.2953T>C
c.210_212del
c.429G>A
c.330C>T

c.9906_9908d
c.318C>A
c.41A>G
c.226G>A
c.2950_2952d
c.2525A>G
c.1323G>A
c.1320_1322d
c.3111T>C
c.1433C>T
c.6868G>C
c.447T>G
c.73G>A
c.1051T>G
c.854_857dup
c.233_235del
c.1385C>A
c.8340_8375d
c.4586C>T
c.9906_9908d
c.318C>A
c.453_455del
c.226G>A
c.2950_2952d
c.2525A>G
c.2667_2669d
c.1323G>A
c.1320_1322d
c.2433T>G
c.7G>A
c.6868G>C
c.73G>A
c.228A>G
c.268_270del
c.360G>T
c.1146A>T
c.854_857dup
c.3718G>T
c.730A>C
c.8340_8375d
c.2124G>T
c.9906_9908d
c.453_455del
c.1041_1043d
c.226G>A
c.2667_2669d
c.3755G>A
c.1269C>T
c.1323G>A
c.1320_1322d

c.3111T>C
c.2433T>G
c.9631C>A
c.268_270del
c.2131_2133d
c.360G>T

| HGVSp | HGVSp_Short | Transcript_ID | Exon_Number | t_depth | t_ref_count | t_alt_count |
|---|---|---|---|---|---|---|
| p.Phe154Leu | p.F154L | ENSMUST000( | 7/13 | 370 | 365 | 5 |
| p.Trp286Cysfs | p.W286Cfs*27 | ENSMUST000( | 11/11 | 621 | 365 | 256 |
| p.Gly1413= | p.G1413= | ENSMUST000( | 27/41 | 373 | 368 | 5 |
| p.Val215Ile | p.V215I | ENSMUST000( | 7/12 | 598 | 592 | 6 |
| p.Gln3302del | p.Q3302del | ENSMUST000( | 58/66 | 92 | 76 | 16 |
| p.Ala106= | p.A106= | ENSMUST000( | 2/16 | 584 | 421 | 162 |
| p.Gln152del | p.Q152del | ENSMUST000( | 2/2 | 176 | 142 | 34 |
| p.Glu76Lys | p.E76K | ENSMUST000( | 3/16 | 537 | 268 | 269 |
| p.Gln984del | p.Q984del | ENSMUST000( | 22/24 | 134 | 123 | 11 |
| p.Asp842Gly | p.D842G | ENSMUST000( | 20/24 | 397 | 366 | 30 |
| p.Ala375Thr | p.A375T | ENSMUST000( | 10/24 | 431 | 425 | 5 |
| p.Ala441= | p.A441= | ENSMUST000( | 5/5 | 507 | 459 | 48 |
| p.Ala447dup | p.A447dup | ENSMUST000( | 5/5 | 584 | 152 | 432 |
| p.Ser447Gly | p.S447G | ENSMUST000( | 7/19 | 73 | 63 | 5 |
| p.Asp21Asn | p.D21N | ENSMUST000( | 1/3 | 111 | 105 | 5 |
| p.Ala3Thr | p.A3T | ENSMUST000( | 1/16 | 82 | 74 | 8 |
| p.Val2290Leu | p.V2290L | ENSMUST000( | 9/13 | 303 | 147 | 156 |
| p.Tyr369Asn | p.Y369N | ENSMUST000( | 8/16 | 410 | 399 | 11 |
| p.Glu120Asp | p.E120D | ENSMUST000( | 4/24 | 377 | 370 | 7 |
| p.Arg315Trp | p.R315W | ENSMUST000( | 2/2 | 372 | 367 | 5 |
| p.Trp286Cysfs | p.W286Cfs*27 | ENSMUST000( | 11/11 | 686 | 411 | 275 |
| p.Gln1449Lys | p.Q1449K | ENSMUST000( | 33/58 | 479 | 325 | 154 |
| p.Arg1478Ter | p.R1478* | ENSMUST000( | 33/58 | 482 | 313 | 169 |
| p.Glu70del | p.E70del | ENSMUST000( | 2/4 | 61 | 51 | 10 |
| p.Gln89del | p.Q89del | ENSMUST000( | 2/9 | 84 | 78 | 6 |
| p.Asn64His | p.N64H | ENSMUST000( | 3/18 | 581 | 571 | 9 |
| p.Ala350del | p.A350del | ENSMUST000( | 1/20 | 740 | 685 | 55 |
| p.Ala548= | p.A548= | ENSMUST000( | 4/4 | 54 | 51 | 3 |
| p.Glu76Lys | p.E76K | ENSMUST000( | 3/16 | 589 | 309 | 280 |
| p.Asp842Gly | p.D842G | ENSMUST000( | 20/24 | 391 | 383 | 8 |
| p.Ala441= | p.A441= | ENSMUST000( | 5/5 | 568 | 508 | 60 |
| p.Ala447dup | p.A447dup | ENSMUST000( | 5/5 | 664 | 143 | 521 |
| p.Trp286Cysfs | p.W286Cfs*27 | ENSMUST000( | 11/11 | 830 | 503 | 327 |
| p.Arg1655Gln | p.R1655Qfs*4 | ENSMUST000( | 37/58 | 918 | 837 | 81 |
| p.Pro17= | p.P17= | ENSMUST000( | 1/27 | 447 | 440 | 7 |
| p.Phe331Ser | p.F331S | ENSMUST000( | 8/8 | 952 | 484 | 467 |
| p.Gln3302del | p.Q3302del | ENSMUST000( | 58/66 | 124 | 94 | 30 |
| p.Gln152del | p.Q152del | ENSMUST000( | 2/2 | 218 | 177 | 41 |
| p.Glu76Lys | p.E76K | ENSMUST000( | 3/16 | 617 | 323 | 294 |
| p.Gln984del | p.Q984del | ENSMUST000( | 22/24 | 194 | 175 | 19 |
| p.Ser423= | p.S423= | ENSMUST000( | 2/2 | 459 | 453 | 6 |
| p.Arg364Trp | p.R364W | ENSMUST000( | 6/10 | 787 | 530 | 257 |
| p.Ala441= | p.A441= | ENSMUST000( | 5/5 | 565 | 528 | 37 |
| p.Ala447dup | p.A447dup | ENSMUST000( | 5/5 | 644 | 314 | 330 |
| p.Ala61Thr | p.A61T | ENSMUST000( | 4/6 | 551 | 545 | 6 |
| p.Met773Leu | p.M773L | ENSMUST000( | 17/18 | 174 | 156 | 18 |
| p.Gly90del | p.G90del | ENSMUST000( | 1/6 | 133 | 118 | 15 |
| p.Trp286Cysfs | p.W286Cfs*27 | ENSMUST000( | 11/11 | 813 | 510 | 303 |
| p.Gly30del | p.G30del | ENSMUST000( | 1/15 | 143 | 131 | 12 |

| | | | | | | |
|---|---|---|---|---|---|---|
| p.Gln775del | p.Q775del | ENSMUST000( | 8/8 | 353 | 306 | 47 |
| p.Pro5= | p.P5= | ENSMUST000( | 1/14 | 204 | 199 | 4 |
| p.Thr244Pro | p.T244P | ENSMUST000( | 2/2 | 108 | 95 | 13 |
| p.Ser145= | p.S145= | ENSMUST000( | 1/12 | 366 | 360 | 6 |
| p.Gln3284del | p.Q3284del | ENSMUST000( | 35/55 | 411 | 354 | 57 |
| p.Glu70del | p.E70del | ENSMUST000( | 2/4 | 90 | 80 | 10 |
| p.Gln3302del | p.Q3302del | ENSMUST000( | 58/66 | 140 | 116 | 24 |
| p.Gln491= | p.Q491= | ENSMUST000( | 2/5 | 513 | 507 | 6 |
| p.Gln152del | p.Q152del | ENSMUST000( | 2/2 | 271 | 228 | 43 |
| p.Glu76Lys | p.E76K | ENSMUST000( | 3/16 | 736 | 362 | 374 |
| p.Gln889del | p.Q889del | ENSMUST000( | 15/19 | 351 | 317 | 34 |
| p.Gly258Ser | p.G258S | ENSMUST000( | 5/5 | 270 | 265 | 5 |
| p.Ala441= | p.A441= | ENSMUST000( | 5/5 | 810 | 708 | 102 |
| p.Ala447dup | p.A447dup | ENSMUST000( | 5/5 | 949 | 191 | 758 |
| p.Pro2452Ala | p.P2452A | ENSMUST000( | 34/34 | 798 | 790 | 8 |
| p.Met773Leu | p.M773L | ENSMUST000( | 17/18 | 212 | 195 | 17 |
| p.Gly25Ser | p.G25S | ENSMUST000( | 1/2 | 156 | 149 | 7 |
| p.Gly90del | p.G90del | ENSMUST000( | 1/6 | 157 | 141 | 16 |
| p.Glu120Asp | p.E120D | ENSMUST000( | 4/24 | 262 | 257 | 5 |
| p.Gln238del | p.Q238del | ENSMUST000( | 1/1 | 117 | 108 | 9 |
| p.Trp286Cysfs | p.W286Cfs*27 | ENSMUST000( | 11/11 | 714 | 419 | 295 |
| p.Arg2271Ter | p.R2271* | ENSMUST000( | 45/58 | 522 | 498 | 23 |
| p.Gly30del | p.G30del | ENSMUST000( | 1/15 | 56 | 49 | 7 |
| p.Gly27del | p.G27del | ENSMUST000( | 2/3 | 68 | 56 | 12 |
| p.Glu70del | p.E70del | ENSMUST000( | 2/4 | 76 | 64 | 12 |
| p.Gln3302del | p.Q3302del | ENSMUST000( | 58/66 | 105 | 93 | 12 |
| p.Gln918del | p.Q918del | ENSMUST000( | 18/28 | 254 | 225 | 29 |
| p.Glu76Lys | p.E76K | ENSMUST000( | 3/16 | 562 | 295 | 267 |
| p.Gln984del | p.Q984del | ENSMUST000( | 22/24 | 174 | 157 | 17 |
| p.Asp842Gly | p.D842G | ENSMUST000( | 20/24 | 367 | 310 | 57 |
| p.Gly258Ser | p.G258S | ENSMUST000( | 5/5 | 187 | 181 | 6 |
| p.Gly90del | p.G90del | ENSMUST000( | 1/6 | 92 | 85 | 7 |
| p.Gln489= | p.Q489= | ENSMUST000( | 3/7 | 399 | 394 | 5 |
| p.Glu382Asp | p.E382D | ENSMUST000( | 2/2 | 198 | 191 | 7 |
| p.Trp286Cysfs | p.W286Cfs*27 | ENSMUST000( | 11/11 | 750 | 96 | 654 |
| p.Leu17= | p.L17= | ENSMUST000( | 1/22 | 322 | 316 | 6 |
| p.Glu1240Ter | p.E1240* | ENSMUST000( | 28/58 | 551 | 65 | 486 |
| p.Gln775del | p.Q775del | ENSMUST000( | 8/8 | 331 | 286 | 45 |
| p.Gln4049del | p.Q4049del | ENSMUST000( | 40/55 | 580 | 510 | 70 |
| p.Gln3302del | p.Q3302del | ENSMUST000( | 58/66 | 123 | 101 | 22 |
| p.Gln2652= | p.Q2652= | ENSMUST000( | 46/52 | 591 | 585 | 6 |
| p.Glu76Lys | p.E76K | ENSMUST000( | 3/16 | 632 | 346 | 286 |
| p.Gln889del | p.Q889del | ENSMUST000( | 15/19 | 264 | 241 | 23 |
| p.Ser423= | p.S423= | ENSMUST000( | 2/2 | 440 | 434 | 5 |
| p.Ala441= | p.A441= | ENSMUST000( | 5/5 | 527 | 503 | 24 |
| p.Ala447dup | p.A447dup | ENSMUST000( | 5/5 | 596 | 278 | 318 |
| p.Ala3Thr | p.A3T | ENSMUST000( | 1/16 | 99 | 92 | 7 |
| p.Ala11= | p.A11= | ENSMUST000( | 1/16 | 117 | 113 | 4 |
| p.Gly25Ser | p.G25S | ENSMUST000( | 1/2 | 112 | 107 | 5 |
| p.Pro127= | p.P127= | ENSMUST000( | 3/11 | 460 | 454 | 5 |

| | | | | | | |
|---|---|---|---|---|---|---|
| p.Ser44Gly | p.S44G | ENSMUST000( | 1/35 | 50 | 47 | 3 |
| p.Asn128Thr | p.N128T | ENSMUST000( | 4/6 | 295 | 291 | 4 |
| p.Trp286Cysfs | p.W286Cfs*27 | ENSMUST000( | 11/11 | 637 | 420 | 217 |
| p.Cys3Ter | p.C3* | ENSMUST000( | 1/24 | 45 | 42 | 3 |
| p.Glu397del | p.E397del | ENSMUST000( | 5/16 | 377 | 346 | 31 |
| p.Gly65Ser | p.G65S | ENSMUST000( | 1/34 | 411 | 405 | 6 |
| p.Gln143= | p.Q143= | ENSMUST000( | 3/9 | 397 | 383 | 14 |
| p.Asp110= | p.D110= | ENSMUST000( | 3/9 | 520 | 502 | 18 |
| p.Gly91= | p.G91= | ENSMUST000( | 1/10 | 490 | 485 | 5 |
| p.Gln3302del | p.Q3302del | ENSMUST000( | 58/66 | 94 | 73 | 21 |
| p.Phe180Leu | p.F180L | ENSMUST000( | 5/5 | 1296 | 1274 | 15 |
| p.Ala350del | p.A350del | ENSMUST000( | 1/20 | 897 | 831 | 66 |
| p.Ser173= | p.S173= | ENSMUST000( | 5/11 | 273 | 267 | 5 |
| p.Glu76Lys | p.E76K | ENSMUST000( | 3/16 | 523 | 289 | 234 |
| p.Ala18del | p.A18del | ENSMUST000( | 1/76 | 61 | 54 | 7 |
| p.Glu1194= | p.E1194= | ENSMUST000( | 11/17 | 559 | 549 | 7 |
| p.Gln984del | p.Q984del | ENSMUST000( | 22/24 | 176 | 158 | 18 |
| p.Asp842Gly | p.D842G | ENSMUST000( | 20/24 | 423 | 234 | 189 |
| p.Asp762Gly | p.D762G | ENSMUST000( | 16/21 | 491 | 486 | 5 |
| p.Ala441= | p.A441= | ENSMUST000( | 5/5 | 656 | 600 | 56 |
| p.Ala447dup | p.A447dup | ENSMUST000( | 5/5 | 771 | 155 | 616 |
| p.Ser447Gly | p.S447G | ENSMUST000( | 7/19 | 80 | 73 | 3 |
| p.Thr3341Ala | p.T3341A | ENSMUST000( | 15/27 | 392 | 386 | 4 |
| p.Gly90del | p.G90del | ENSMUST000( | 1/6 | 97 | 90 | 7 |
| p.Trp286Cysfs | p.W286Cfs*27 | ENSMUST000( | 11/11 | 641 | 406 | 235 |
| p.Gln156del | p.Q156del | ENSMUST000( | 7/18 | 176 | 159 | 17 |
| p.Glu70del | p.E70del | ENSMUST000( | 2/4 | 69 | 60 | 9 |
| p.Glu245Lys | p.E245K | ENSMUST000( | 3/8 | 542 | 522 | 19 |
| p.Pro63del | p.P63del | ENSMUST000( | 1/12 | 55 | 46 | 9 |
| p.Gln3302del | p.Q3302del | ENSMUST000( | 58/66 | 108 | 87 | 21 |
| p.Ala350del | p.A350del | ENSMUST000( | 1/20 | 959 | 895 | 64 |
| p.Ser156= | p.S156= | ENSMUST000( | 5/11 | 214 | 209 | 4 |
| p.Glu76Lys | p.E76K | ENSMUST000( | 3/16 | 525 | 274 | 251 |
| p.Ala441= | p.A441= | ENSMUST000( | 5/5 | 492 | 460 | 31 |
| p.Ala447dup | p.A447dup | ENSMUST000( | 5/5 | 549 | 250 | 299 |
| p.Arg1037= | p.R1037= | ENSMUST000( | 20/34 | 978 | 965 | 12 |
| p.Gly25Ser | p.G25S | ENSMUST000( | 1/2 | 121 | 115 | 6 |
| p.Gly518Ser | p.G518S | ENSMUST000( | 14/14 | 447 | 441 | 6 |
| p.Trp286Cysfs | p.W286Cfs*27 | ENSMUST000( | 11/11 | 702 | 422 | 280 |
| p.Glu70del | p.E70del | ENSMUST000( | 2/4 | 58 | 52 | 6 |
| p.Thr248= | p.T248= | ENSMUST000( | 7/11 | 407 | 393 | 14 |
| p.Ala67= | p.A67= | ENSMUST000( | 2/22 | 407 | 402 | 5 |
| p.Ile213Lys | p.I213K | ENSMUST000( | 2/2 | 8 | 2 | 6 |
| p.Ile213= | p.I213= | ENSMUST000( | 2/2 | 8 | 2 | 6 |
| p.Gln143= | p.Q143= | ENSMUST000( | 3/9 | 388 | 355 | 33 |
| p.Asp110= | p.D110= | ENSMUST000( | 3/9 | 459 | 415 | 44 |
| p.Gln3302del | p.Q3302del | ENSMUST000( | 58/66 | 90 | 76 | 14 |
| p.Gln648= | p.Q648= | ENSMUST000( | 13/32 | 326 | 320 | 5 |
| p.Gly2235Cys | p.G2235C | ENSMUST000( | 34/34 | 348 | 342 | 5 |
| p.Glu76Lys | p.E76K | ENSMUST000( | 3/16 | 484 | 265 | 218 |

| | | | | | | |
|---|---|---|---|---|---|---|
| p.Gln984del | p.Q984del | ENSMUST000( | 22/24 | 186 | 167 | 19 |
| p.Asp842Gly | p.D842G | ENSMUST000( | 20/24 | 408 | 236 | 172 |
| p.Ala441= | p.A441= | ENSMUST000( | 5/5 | 511 | 460 | 50 |
| p.Ala447dup | p.A447dup | ENSMUST000( | 5/5 | 603 | 109 | 494 |
| p.Asp811Glu | p.D811E | ENSMUST000( | 11/12 | 107 | 92 | 15 |
| p.Glu179Gly | p.E179G | ENSMUST000( | 5/17 | 599 | 593 | 6 |
| p.Gly25Ser | p.G25S | ENSMUST000( | 1/2 | 99 | 94 | 5 |
| p.Lys892Arg | p.K892R | ENSMUST000( | 3/36 | 418 | 413 | 5 |
| p.Gly90del | p.G90del | ENSMUST000( | 1/6 | 82 | 71 | 11 |
| p.Glu120Asp | p.E120D | ENSMUST000( | 4/24 | 356 | 351 | 5 |
| p.Glu382Asp | p.E382D | ENSMUST000( | 2/2 | 160 | 152 | 8 |
| p.Ala1399Ser | p.A1399S | ENSMUST000( | 27/44 | 642 | 617 | 25 |
| p.Trp286Cysfs | p.W286Cfs*27 | ENSMUST000( | 11/11 | 621 | 405 | 216 |
| p.Arg498= | p.R498= | ENSMUST000( | 9/10 | 490 | 476 | 13 |
| p.Gln1648Leu | p.Q1648L | ENSMUST000( | 17/17 | 55 | 52 | 3 |
| p.Gln143= | p.Q143= | ENSMUST000( | 3/9 | 476 | 453 | 22 |
| p.Asp110= | p.D110= | ENSMUST000( | 3/9 | 548 | 519 | 29 |
| p.Gln3302del | p.Q3302del | ENSMUST000( | 58/66 | 115 | 96 | 19 |
| p.Ala106= | p.A106= | ENSMUST000( | 2/16 | 574 | 340 | 234 |
| p.Leu181Pro | p.L181P | ENSMUST000( | 5/12 | 417 | 411 | 5 |
| p.Ala350del | p.A350del | ENSMUST000( | 1/20 | 850 | 796 | 54 |
| p.Glu76Lys | p.E76K | ENSMUST000( | 3/16 | 591 | 333 | 258 |
| p.Gln984del | p.Q984del | ENSMUST000( | 22/24 | 190 | 171 | 19 |
| p.Asp842Gly | p.D842G | ENSMUST000( | 20/24 | 448 | 238 | 210 |
| p.Gln889del | p.Q889del | ENSMUST000( | 15/19 | 248 | 227 | 21 |
| p.Ala441= | p.A441= | ENSMUST000( | 5/5 | 638 | 569 | 69 |
| p.Ala447dup | p.A447dup | ENSMUST000( | 5/5 | 749 | 154 | 595 |
| p.Met773Leu | p.M773L | ENSMUST000( | 17/18 | 181 | 165 | 16 |
| p.Val2290Leu | p.V2290L | ENSMUST000( | 9/13 | 402 | 203 | 198 |
| p.Gly90del | p.G90del | ENSMUST000( | 1/6 | 117 | 104 | 13 |
| p.Ala711del | p.A711del | ENSMUST000( | 24/27 | 124 | 114 | 10 |
| p.Trp286Cysfs | p.W286Cfs*27 | ENSMUST000( | 11/11 | 628 | 405 | 223 |
| p.Gln183= | p.Q183= | ENSMUST000( | 2/6 | 426 | 421 | 5 |
| p.Glu70del | p.E70del | ENSMUST000( | 2/4 | 59 | 51 | 8 |
| p.Thr233= | p.T233= | ENSMUST000( | 6/7 | 357 | 349 | 5 |
| p.Ile213Lys | p.I213K | ENSMUST000( | 2/2 | 9 | 3 | 6 |
| p.Ile213= | p.I213= | ENSMUST000( | 2/2 | 9 | 3 | 6 |
| p.Pro63del | p.P63del | ENSMUST000( | 1/12 | 69 | 63 | 6 |
| p.Gln143= | p.Q143= | ENSMUST000( | 3/9 | 382 | 363 | 19 |
| p.Asp110= | p.D110= | ENSMUST000( | 3/9 | 461 | 430 | 31 |
| p.Gln3302del | p.Q3302del | ENSMUST000( | 58/66 | 107 | 87 | 20 |
| p.Glu76Lys | p.E76K | ENSMUST000( | 3/16 | 531 | 299 | 232 |
| p.Gln984del | p.Q984del | ENSMUST000( | 22/24 | 155 | 141 | 14 |
| p.Asp842Gly | p.D842G | ENSMUST000( | 20/24 | 423 | 217 | 206 |
| p.Ala441= | p.A441= | ENSMUST000( | 5/5 | 547 | 499 | 48 |
| p.Ala447dup | p.A447dup | ENSMUST000( | 5/5 | 657 | 117 | 540 |
| p.Pro2053= | p.P2053= | ENSMUST000( | 15/16 | 350 | 346 | 4 |
| p.Ala257Thr | p.A257T | ENSMUST000( | 5/8 | 455 | 450 | 5 |
| p.Ala711del | p.A711del | ENSMUST000( | 24/27 | 113 | 105 | 8 |
| p.Trp286Cysfs | p.W286Cfs*27 | ENSMUST000( | 11/11 | 630 | 392 | 238 |

| | | | | | | |
|---|---|---|---|---|---|---|
| p.Thr812Lys | p.T812K | ENSMUST000( | 20/24 | 557 | 550 | 6 |
| p.Gly27del | p.G27del | ENSMUST000( | 2/3 | 96 | 87 | 9 |
| p.Pro17= | p.P17= | ENSMUST000( | 1/27 | 486 | 479 | 7 |
| p.Gly65Ser | p.G65S | ENSMUST000( | 1/34 | 425 | 420 | 5 |
| p.Gln143= | p.Q143= | ENSMUST000( | 3/9 | 434 | 418 | 16 |
| p.Asp110= | p.D110= | ENSMUST000( | 3/9 | 527 | 500 | 27 |
| p.Ser96Asn | p.S96N | ENSMUST000( | 2/8 | 541 | 535 | 6 |
| p.Gln3302del | p.Q3302del | ENSMUST000( | 58/66 | 134 | 110 | 24 |
| p.Gln152del | p.Q152del | ENSMUST000( | 2/2 | 219 | 183 | 36 |
| p.Glu76Lys | p.E76K | ENSMUST000( | 3/16 | 569 | 322 | 246 |
| p.Ala18del | p.A18del | ENSMUST000( | 1/76 | 61 | 53 | 8 |
| p.Gln984del | p.Q984del | ENSMUST000( | 22/24 | 217 | 196 | 21 |
| p.Gly258Ser | p.G258S | ENSMUST000( | 5/5 | 255 | 249 | 6 |
| p.Ala441= | p.A441= | ENSMUST000( | 5/5 | 697 | 635 | 62 |
| p.Ala447dup | p.A447dup | ENSMUST000( | 5/5 | 865 | 175 | 690 |
| p.Ala11= | p.A11= | ENSMUST000( | 1/16 | 176 | 168 | 8 |
| p.Met773Leu | p.M773L | ENSMUST000( | 17/18 | 150 | 138 | 12 |
| p.Glu1550Gly | p.E1550G | ENSMUST000( | 32/34 | 747 | 738 | 9 |
| p.Gly90del | p.G90del | ENSMUST000( | 1/6 | 114 | 104 | 10 |
| p.Gln265His | p.Q265H | ENSMUST000( | 1/1 | 87 | 81 | 5 |
| p.Gln154del | p.Q154del | ENSMUST000( | 1/1 | 100 | 88 | 12 |
| p.Ser44del | p.S44del | ENSMUST000( | 1/35 | 39 | 32 | 7 |
| p.Arg696Gly | p.R696G | ENSMUST000( | 5/8 | 639 | 631 | 7 |
| p.Trp286Cysfs | p.W286Cfs*27 | ENSMUST000( | 11/11 | 693 | 449 | 244 |
| p.Gln775del | p.Q775del | ENSMUST000( | 8/8 | 257 | 221 | 36 |
| p.Pro17= | p.P17= | ENSMUST000( | 1/27 | 356 | 352 | 4 |
| p.Gln3601= | p.Q3601= | ENSMUST000( | 40/55 | 667 | 660 | 7 |
| p.Glu70del | p.E70del | ENSMUST000( | 2/4 | 62 | 56 | 6 |
| p.Glu1823del | p.E1823del | ENSMUST000( | 10/10 | 629 | 571 | 58 |
| p.Phe331Ser | p.F331S | ENSMUST000( | 8/8 | 930 | 536 | 393 |
| p.Gln143= | p.Q143= | ENSMUST000( | 3/9 | 459 | 432 | 26 |
| p.Asp110= | p.D110= | ENSMUST000( | 3/9 | 536 | 494 | 42 |
| p.Gln3302del | p.Q3302del | ENSMUST000( | 58/66 | 109 | 86 | 23 |
| p.Thr772= | p.T772= | ENSMUST000( | 14/91 | 585 | 448 | 136 |
| p.Gln491= | p.Q491= | ENSMUST000( | 2/5 | 376 | 368 | 6 |
| p.Pro2022= | p.P2022= | ENSMUST000( | 10/11 | 351 | 334 | 13 |
| p.Glu76Lys | p.E76K | ENSMUST000( | 3/16 | 606 | 366 | 240 |
| p.Gln984del | p.Q984del | ENSMUST000( | 22/24 | 166 | 149 | 17 |
| p.Asp818Tyr | p.D818Y | ENSMUST000( | 17/21 | 587 | 383 | 204 |
| p.Gly12Asp | p.G12D | ENSMUST000( | 2/5 | 674 | 667 | 7 |
| p.Arg364Trp | p.R364W | ENSMUST000( | 6/10 | 634 | 459 | 173 |
| p.Ala441= | p.A441= | ENSMUST000( | 5/5 | 418 | 383 | 35 |
| p.Ala447dup | p.A447dup | ENSMUST000( | 5/5 | 439 | 200 | 239 |
| p.Ala11= | p.A11= | ENSMUST000( | 1/16 | 113 | 106 | 6 |
| p.Arg537= | p.R537= | ENSMUST000( | 4/19 | 541 | 532 | 9 |
| p.Trp286Cysfs | p.W286Cfs*27 | ENSMUST000( | 11/11 | 699 | 481 | 218 |
| p.Ser985Pro | p.S985P | ENSMUST000( | 8/8 | 425 | 420 | 5 |
| p.Glu70del | p.E70del | ENSMUST000( | 2/4 | 53 | 45 | 8 |
| p.Gln143= | p.Q143= | ENSMUST000( | 3/9 | 388 | 363 | 25 |
| p.Asp110= | p.D110= | ENSMUST000( | 3/9 | 455 | 421 | 34 |

| | | | | | | |
|---|---|---|---|---|---|---|
| p.Gln3302del | p.Q3302del | ENSMUST000( | 58/66 | 117 | 91 | 26 |
| p.Ala106= | p.A106= | ENSMUST000( | 2/16 | 565 | 335 | 229 |
| p.Lys14Arg | p.K14R | ENSMUST000( | 2/23 | 3 | 1 | 2 |
| p.Glu76Lys | p.E76K | ENSMUST000( | 3/16 | 561 | 333 | 227 |
| p.Gln984del | p.Q984del | ENSMUST000( | 22/24 | 152 | 138 | 14 |
| p.Asp842Gly | p.D842G | ENSMUST000( | 20/24 | 395 | 246 | 149 |
| p.Ala441= | p.A441= | ENSMUST000( | 5/5 | 511 | 453 | 56 |
| p.Ala447dup | p.A447dup | ENSMUST000( | 5/5 | 597 | 133 | 464 |
| p.Arg1037= | p.R1037= | ENSMUST000( | 20/34 | 875 | 865 | 9 |
| p.Ala478Val | p.A478V | ENSMUST000( | 12/17 | 481 | 475 | 5 |
| p.Val2290Leu | p.V2290L | ENSMUST000( | 9/13 | 400 | 225 | 175 |
| p.Arg149= | p.R149= | ENSMUST000( | 3/7 | 86 | 76 | 10 |
| p.Gly25Ser | p.G25S | ENSMUST000( | 1/2 | 86 | 79 | 6 |
| p.Tyr351Asp | p.Y351D | ENSMUST000( | 11/11 | 55 | 47 | 8 |
| p.Trp286Cysfs | p.W286Cfs*27 | ENSMUST000( | 11/11 | 622 | 453 | 169 |
| p.Pro78del | p.P78del | ENSMUST000( | 1/40 | 34 | 28 | 6 |
| p.Pro462His | p.P462H | ENSMUST000( | 6/31 | 329 | 323 | 6 |
| p.Gln2786_Gl | p.Q2786_Q27 | ENSMUST000( | 35/55 | 681 | 619 | 62 |
| p.Ala1529Val | p.A1529V | ENSMUST000( | 27/29 | 414 | 409 | 5 |
| p.Gln3302del | p.Q3302del | ENSMUST000( | 58/66 | 112 | 97 | 15 |
| p.Ala106= | p.A106= | ENSMUST000( | 2/16 | 551 | 361 | 190 |
| p.Gln152del | p.Q152del | ENSMUST000( | 2/2 | 163 | 126 | 37 |
| p.Glu76Lys | p.E76K | ENSMUST000( | 3/16 | 563 | 363 | 200 |
| p.Gln984del | p.Q984del | ENSMUST000( | 22/24 | 139 | 126 | 13 |
| p.Asp842Gly | p.D842G | ENSMUST000( | 20/24 | 365 | 227 | 138 |
| p.Gln889del | p.Q889del | ENSMUST000( | 15/19 | 209 | 194 | 15 |
| p.Ala441= | p.A441= | ENSMUST000( | 5/5 | 436 | 404 | 31 |
| p.Ala447dup | p.A447dup | ENSMUST000( | 5/5 | 510 | 127 | 383 |
| p.Asp811Glu | p.D811E | ENSMUST000( | 11/12 | 106 | 87 | 19 |
| p.Ala3Thr | p.A3T | ENSMUST000( | 1/16 | 77 | 70 | 7 |
| p.Val2290Leu | p.V2290L | ENSMUST000( | 9/13 | 324 | 205 | 119 |
| p.Gly25Ser | p.G25S | ENSMUST000( | 1/2 | 96 | 92 | 4 |
| p.Gly76= | p.G76= | ENSMUST000( | 2/25 | 321 | 304 | 17 |
| p.Gly90del | p.G90del | ENSMUST000( | 1/6 | 68 | 57 | 11 |
| p.Glu120Asp | p.E120D | ENSMUST000( | 4/24 | 300 | 290 | 10 |
| p.Glu382Asp | p.E382D | ENSMUST000( | 2/2 | 142 | 136 | 6 |
| p.Trp286Cysfs | p.W286Cfs*27 | ENSMUST000( | 11/11 | 764 | 264 | 500 |
| p.Glu1240Ter | p.E1240* | ENSMUST000( | 28/58 | 504 | 108 | 396 |
| p.Thr244Pro | p.T244P | ENSMUST000( | 2/2 | 83 | 72 | 11 |
| p.Gln2786_Gl | p.Q2786_Q27 | ENSMUST000( | 35/55 | 839 | 742 | 97 |
| p.Pro708= | p.P708= | ENSMUST000( | 6/20 | 571 | 561 | 8 |
| p.Gln3302del | p.Q3302del | ENSMUST000( | 58/66 | 103 | 81 | 22 |
| p.Gln152del | p.Q152del | ENSMUST000( | 2/2 | 194 | 150 | 44 |
| p.Ala350del | p.A350del | ENSMUST000( | 1/20 | 756 | 704 | 52 |
| p.Glu76Lys | p.E76K | ENSMUST000( | 3/16 | 639 | 416 | 222 |
| p.Gln889del | p.Q889del | ENSMUST000( | 15/19 | 245 | 223 | 22 |
| p.Arg1252His | p.R1252H | ENSMUST000( | 23/28 | 488 | 483 | 5 |
| p.Ser423= | p.S423= | ENSMUST000( | 2/2 | 440 | 435 | 5 |
| p.Ala441= | p.A441= | ENSMUST000( | 5/5 | 475 | 441 | 34 |
| p.Ala447dup | p.A447dup | ENSMUST000( | 5/5 | 521 | 280 | 241 |

| | | | | | | |
|---|---|---|---|---|---|---|
| p.Arg1037= | p.R1037= | ENSMUST000( | 20/34 | 1004 | 993 | 11 |
| p.Asp811Glu | p.D811E | ENSMUST000( | 11/12 | 108 | 97 | 11 |
| p.Leu3211Ile | p.L3211I | ENSMUST000( | 27/36 | 409 | 404 | 5 |
| p.Gly90del | p.G90del | ENSMUST000( | 1/6 | 94 | 87 | 7 |
| p.Ala711del | p.A711del | ENSMUST000( | 24/27 | 84 | 78 | 6 |
| p.Glu120Asp | p.E120D | ENSMUST000( | 4/24 | 209 | 205 | 4 |

| n_depth | n_ref_count | n_alt_count | all_effects | Allele | Gene | Feature |
|---|---|---|---|---|---|---|
| 142 | 142 | 0 | Etv4,missense | G | ENSMUSG000 | ENSMUST000( |
| 358 | 358 | 0 | Npm1,framesl | CAGA | ENSMUSG000 | ENSMUST000( |
| 131 | 130 | 0 | Myo18a,synor | G | ENSMUSG000 | ENSMUST000( |
| 251 | 251 | 0 | Smyd3,missen | T | ENSMUSG000 | ENSMUST000( |
| 9 | 8 | 1 | Sptbn5,infram | - | ENSMUSG000 | ENSMUST000( |
| 218 | 218 | 0 | Samhd1,synor | T | ENSMUSG000 | ENSMUST000( |
| 41 | 35 | 6 | Ivl,inframe_de | - | ENSMUSG000 | ENSMUST000( |
| 224 | 224 | 0 | Ptpn11,misser | T | ENSMUSG000 | ENSMUST000( |
| 84 | 82 | 2 | Gigyf1,inframe | - | ENSMUSG000 | ENSMUST000( |
| 145 | 145 | 0 | Flt3,missense_ | C | ENSMUSG000 | ENSMUST000( |
| 174 | 174 | 0 | Pik3c2g,misse | A | ENSMUSG000 | ENSMUST000( |
| 138 | 137 | 1 | Maz,synonym | T | ENSMUSG000 | ENSMUST000( |
| 145 | 145 | 0 | Maz,inframe_i | GCT | ENSMUSG000 | ENSMUST000( |
| 13 | 12 | 0 | Setd1a,misser | G | ENSMUSG000 | ENSMUST000( |
| 29 | 29 | 0 | Scgb1b30,mis | A | ENSMUSG000 | ENSMUST000( |
| 34 | 33 | 1 | Rsf1,missense | A | ENSMUSG000 | ENSMUST000( |
| 136 | 136 | 0 | Ankrd11,miss | G | ENSMUSG000 | ENSMUST000( |
| 135 | 135 | 0 | Cbl,missense_ | T | ENSMUSG000 | ENSMUST000( |
| 110 | 110 | 0 | Rbm10,misser | T | ENSMUSG000 | ENSMUST000( |
| 134 | 134 | 0 | Egr2,missense | T | ENSMUSG000 | ENSMUST000( |
| 358 | 358 | 0 | Npm1,framesl | CAGA | ENSMUSG000 | ENSMUST000( |
| 225 | 225 | 0 | Nf1,missense_ | A | ENSMUSG000 | ENSMUST000( |
| 213 | 213 | 0 | Nf1,stop_gain | T | ENSMUSG000 | ENSMUST000( |
| 22 | 22 | 0 | Vgll3,inframe_ | - | ENSMUSG000 | ENSMUST000( |
| 69 | 68 | 1 | Klhl14,inframe | - | ENSMUSG000 | ENSMUST000( |
| 255 | 255 | 0 | Nt5c2,missens | G | ENSMUSG000 | ENSMUST000( |
| 243 | 233 | 10 | Arid1a,infram | - | ENSMUSG000 | ENSMUST000( |
| 57 | 57 | 0 | Prdm13,synon | T | ENSMUSG000 | ENSMUST000( |
| 224 | 224 | 0 | Ptpn11,misser | T | ENSMUSG000 | ENSMUST000( |
| 145 | 145 | 0 | Flt3,missense_ | C | ENSMUSG000 | ENSMUST000( |
| 138 | 137 | 1 | Maz,synonym | T | ENSMUSG000 | ENSMUST000( |
| 145 | 145 | 0 | Maz,inframe_i | GCT | ENSMUSG000 | ENSMUST000( |
| 358 | 358 | 0 | Npm1,framesl | CAGA | ENSMUSG000 | ENSMUST000( |
| 266 | 266 | 0 | Nf1,frameshif | - | ENSMUSG000 | ENSMUST000( |
| 115 | 115 | 0 | Rb1,synonym | T | ENSMUSG000 | ENSMUST000( |
| 305 | 305 | 0 | Serpinb3b,mis | G | ENSMUSG000 | ENSMUST000( |
| 9 | 8 | 1 | Sptbn5,infram | - | ENSMUSG000 | ENSMUST000( |
| 41 | 35 | 6 | Ivl,inframe_de | - | ENSMUSG000 | ENSMUST000( |
| 224 | 224 | 0 | Ptpn11,misser | T | ENSMUSG000 | ENSMUST000( |
| 84 | 82 | 2 | Gigyf1,inframe | - | ENSMUSG000 | ENSMUST000( |
| 201 | 201 | 0 | D6Ertd527e,sy | T | ENSMUSG000 | ENSMUST000( |
| 147 | 147 | 0 | Plk1,missense | T | ENSMUSG000 | ENSMUST000( |
| 138 | 137 | 1 | Maz,synonym | T | ENSMUSG000 | ENSMUST000( |
| 145 | 145 | 0 | Maz,inframe_i | GCT | ENSMUSG000 | ENSMUST000( |
| 109 | 109 | 0 | Rps19,missens | A | ENSMUSG000 | ENSMUST000( |
| 76 | 75 | 1 | Cdh16,missen | G | ENSMUSG000 | ENSMUST000( |
| 30 | 29 | 1 | Ammecr1,infr | - | ENSMUSG000 | ENSMUST000( |
| 358 | 358 | 0 | Npm1,framesl | CAGA | ENSMUSG000 | ENSMUST000( |
| 21 | 21 | 0 | Zswim6,infran | - | ENSMUSG000 | ENSMUST000( |

| | | | | | | |
|---|---|---|---|---|---|---|
| 40 | 36 | 4 | Hcn1,inframe_ | - | ENSMUSG000 | ENSMUST000( |
| 36 | 36 | 0 | Ranbp9,synon | T | ENSMUSG000 | ENSMUST000( |
| 41 | 40 | 1 | Fam170b,miss | C | ENSMUSG000 | ENSMUST000( |
| 130 | 130 | 0 | Dach1,synony | A | ENSMUSG000 | ENSMUST000( |
| 91 | 80 | 11 | Kmt2d,inframe | - | ENSMUSG000 | ENSMUST000( |
| 22 | 22 | 0 | Vgll3,inframe_ | - | ENSMUSG000 | ENSMUST000( |
| 9 | 8 | 1 | Sptbn5,infram | - | ENSMUSG000 | ENSMUST000( |
| 166 | 165 | 0 | Maml3,synony | T | ENSMUSG000 | ENSMUST000( |
| 41 | 35 | 6 | Ivl,inframe_de | - | ENSMUSG000 | ENSMUST000( |
| 224 | 224 | 0 | Ptpn11,misser | T | ENSMUSG000 | ENSMUST000( |
| 32 | 31 | 1 | Rbm33,infram | - | ENSMUSG000 | ENSMUST000( |
| 33 | 33 | 0 | Cckar,missens | T | ENSMUSG000 | ENSMUST000( |
| 138 | 137 | 1 | Maz,synonym | T | ENSMUSG000 | ENSMUST000( |
| 145 | 145 | 0 | Maz,inframe_i | GCT | ENSMUSG000 | ENSMUST000( |
| 168 | 168 | 0 | Srcap,missens | G | ENSMUSG000 | ENSMUST000( |
| 76 | 75 | 1 | Cdh16,missen | G | ENSMUSG000 | ENSMUST000( |
| 40 | 39 | 1 | Zfp703,misser | A | ENSMUSG000 | ENSMUST000( |
| 30 | 29 | 1 | Ammecr1,infra | - | ENSMUSG000 | ENSMUST000( |
| 110 | 110 | 0 | Rbm10,misser | T | ENSMUSG000 | ENSMUST000( |
| 16 | 16 | 0 | Sry,inframe_d | - | ENSMUSG000 | ENSMUST000( |
| 358 | 358 | 0 | Npm1,framesh | CAGA | ENSMUSG000 | ENSMUST000( |
| 189 | 189 | 0 | Nf1,stop_gain | T | ENSMUSG000 | ENSMUST000( |
| 21 | 21 | 0 | Zswim6,infram | - | ENSMUSG000 | ENSMUST000( |
| 14 | 14 | 0 | Zfp503,infram | - | ENSMUSG000 | ENSMUST000( |
| 22 | 22 | 0 | Vgll3,inframe_ | - | ENSMUSG000 | ENSMUST000( |
| 9 | 8 | 1 | Sptbn5,infram | - | ENSMUSG000 | ENSMUST000( |
| 97 | 92 | 5 | Dennd4b,infra | - | ENSMUSG000 | ENSMUST000( |
| 224 | 224 | 0 | Ptpn11,misser | T | ENSMUSG000 | ENSMUST000( |
| 84 | 82 | 2 | Gigyf1,inframe | - | ENSMUSG000 | ENSMUST000( |
| 145 | 145 | 0 | Flt3,missense_ | C | ENSMUSG000 | ENSMUST000( |
| 33 | 33 | 0 | Cckar,missens | T | ENSMUSG000 | ENSMUST000( |
| 30 | 29 | 1 | Ammecr1,infra | - | ENSMUSG000 | ENSMUST000( |
| 131 | 131 | 0 | Mamld1,synor | A | ENSMUSG000 | ENSMUST000( |
| 72 | 71 | 1 | Amer1,missen | A | ENSMUSG000 | ENSMUST000( |
| 358 | 358 | 0 | Npm1,framesh | CAGA | ENSMUSG000 | ENSMUST000( |
| 106 | 106 | 0 | Adamts2,syno | A | ENSMUSG000 | ENSMUST000( |
| 184 | 183 | 1 | Nf1,stop_gain | T | ENSMUSG000 | ENSMUST000( |
| 40 | 36 | 4 | Hcn1,inframe_ | - | ENSMUSG000 | ENSMUST000( |
| 192 | 179 | 13 | Kmt2d,inframe | - | ENSMUSG000 | ENSMUST000( |
| 9 | 8 | 1 | Sptbn5,infram | - | ENSMUSG000 | ENSMUST000( |
| 196 | 195 | 0 | Ep400,synony | T | ENSMUSG000 | ENSMUST000( |
| 224 | 224 | 0 | Ptpn11,misser | T | ENSMUSG000 | ENSMUST000( |
| 32 | 31 | 1 | Rbm33,infram | - | ENSMUSG000 | ENSMUST000( |
| 201 | 201 | 0 | D6Ertd527e,sy | T | ENSMUSG000 | ENSMUST000( |
| 138 | 137 | 1 | Maz,synonym | T | ENSMUSG000 | ENSMUST000( |
| 145 | 145 | 0 | Maz,inframe_i | GCT | ENSMUSG000 | ENSMUST000( |
| 34 | 33 | 1 | Rsf1,missense | A | ENSMUSG000 | ENSMUST000( |
| 38 | 38 | 0 | Rsf1,synonym | G | ENSMUSG000 | ENSMUST000( |
| 40 | 39 | 1 | Zfp703,misser | A | ENSMUSG000 | ENSMUST000( |
| 145 | 145 | 0 | Gab1,synonym | T | ENSMUSG000 | ENSMUST000( |

| | | | | | | |
|---|---|---|---|---|---|---|
| 43 | 43 | 0 | Zfc3h1,missen | G | ENSMUSG000 | ENSMUST000( |
| 183 | 183 | 0 | Actg1,missens | G | ENSMUSG000 | ENSMUST000( |
| 358 | 358 | 0 | Npm1,framesl | CAGA | ENSMUSG000 | ENSMUST000( |
| 10 | 10 | 0 | Dsp,stop_gain | A | ENSMUSG000 | ENSMUST000( |
| 142 | 136 | 6 | Mdc1,inframe | - | ENSMUSG000 | ENSMUST000( |
| 112 | 112 | 0 | Ankhd1,misse | A | ENSMUSG000 | ENSMUST000( |
| 137 | 137 | 0 | Elf3,synonymc | T | ENSMUSG000 | ENSMUST000( |
| 145 | 145 | 0 | Elf3,synonymc | A | ENSMUSG000 | ENSMUST000( |
| 171 | 170 | 0 | Wt1,synonym | T | ENSMUSG000 | ENSMUST000( |
| 9 | 8 | 1 | Sptbn5,infram | - | ENSMUSG000 | ENSMUST000( |
| 1263 | 1262 | 1 | Gm10800,mis | C | ENSMUSG000 | ENSMUST000( |
| 243 | 233 | 10 | Arid1a,infram | - | ENSMUSG000 | ENSMUST000( |
| 153 | 153 | 0 | Mllt3,synonyn | A | ENSMUSG000 | ENSMUST000( |
| 224 | 224 | 0 | Ptpn11,misser | T | ENSMUSG000 | ENSMUST000( |
| 24 | 24 | 0 | Hectd4,infram | - | ENSMUSG000 | ENSMUST000( |
| 201 | 201 | 0 | Setd1b,synony | A | ENSMUSG000 | ENSMUST000( |
| 84 | 82 | 2 | Gigyf1,inframe | - | ENSMUSG000 | ENSMUST000( |
| 145 | 145 | 0 | Flt3,missense_ | C | ENSMUSG000 | ENSMUST000( |
| 198 | 198 | 0 | Kit,missense_v | G | ENSMUSG000 | ENSMUST000( |
| 138 | 137 | 1 | Maz,synonym | T | ENSMUSG000 | ENSMUST000( |
| 145 | 145 | 0 | Maz,inframe_i | GCT | ENSMUSG000 | ENSMUST000( |
| 13 | 12 | 0 | Setd1a,misser | G | ENSMUSG000 | ENSMUST000( |
| 165 | 164 | 0 | Fat1,missense | G | ENSMUSG000 | ENSMUST000( |
| 30 | 29 | 1 | Ammecr1,infra | - | ENSMUSG000 | ENSMUST000( |
| 358 | 358 | 0 | Npm1,framesl | CAGA | ENSMUSG000 | ENSMUST000( |
| 40 | 37 | 3 | Rpa1,inframe_ | - | ENSMUSG000 | ENSMUST000( |
| 22 | 22 | 0 | Vgll3,inframe_ | - | ENSMUSG000 | ENSMUST000( |
| 168 | 168 | 0 | Vegfa,missens | T | ENSMUSG000 | ENSMUST000( |
| 39 | 39 | 0 | Chka,inframe_ | - | ENSMUSG000 | ENSMUST000( |
| 9 | 8 | 1 | Sptbn5,infram | - | ENSMUSG000 | ENSMUST000( |
| 243 | 233 | 10 | Arid1a,infram | - | ENSMUSG000 | ENSMUST000( |
| 132 | 132 | 0 | Mllt3,synonyn | A | ENSMUSG000 | ENSMUST000( |
| 224 | 224 | 0 | Ptpn11,misser | T | ENSMUSG000 | ENSMUST000( |
| 138 | 137 | 1 | Maz,synonym | T | ENSMUSG000 | ENSMUST000( |
| 145 | 145 | 0 | Maz,inframe_i | GCT | ENSMUSG000 | ENSMUST000( |
| 242 | 242 | 0 | Srcap,synonyn | C | ENSMUSG000 | ENSMUST000( |
| 40 | 39 | 1 | Zfp703,misser | A | ENSMUSG000 | ENSMUST000( |
| 192 | 192 | 0 | Fyn,missense_ | A | ENSMUSG000 | ENSMUST000( |
| 358 | 358 | 0 | Npm1,framesl | CAGA | ENSMUSG000 | ENSMUST000( |
| 22 | 22 | 0 | Vgll3,inframe_ | - | ENSMUSG000 | ENSMUST000( |
| 135 | 135 | 0 | Erg,synonymo | A | ENSMUSG000 | ENSMUST000( |
| 170 | 170 | 0 | Sos1,synonym | G | ENSMUSG000 | ENSMUST000( |
| 10 | 10 | 0 | Olfr1463,miss | A | ENSMUSG000 | ENSMUST000( |
| 10 | 10 | 0 | Olfr1463,syno | T | ENSMUSG000 | ENSMUST000( |
| 137 | 137 | 0 | Elf3,synonymc | T | ENSMUSG000 | ENSMUST000( |
| 145 | 145 | 0 | Elf3,synonymc | A | ENSMUSG000 | ENSMUST000( |
| 9 | 8 | 1 | Sptbn5,infram | - | ENSMUSG000 | ENSMUST000( |
| 168 | 168 | 0 | Prrc2b,synony | A | ENSMUSG000 | ENSMUST000( |
| 138 | 138 | 0 | Notch2,misser | T | ENSMUSG000 | ENSMUST000( |
| 224 | 224 | 0 | Ptpn11,misser | T | ENSMUSG000 | ENSMUST000( |

| | | | | | | |
|---|---|---|---|---|---|---|
| 84 | 82 | 2 | Gigyf1,inframe | - | ENSMUSG000 | ENSMUST000( |
| 145 | 145 | 0 | Flt3,missense_ | C | ENSMUSG000 | ENSMUST000( |
| 138 | 137 | 1 | Maz,synonym | T | ENSMUSG000 | ENSMUST000( |
| 145 | 145 | 0 | Maz,inframe_i | GCT | ENSMUSG000 | ENSMUST000( |
| 33 | 33 | 0 | Grin2d,missen | C | ENSMUSG000 | ENSMUST000( |
| 145 | 143 | 0 | Ntrk3,missens | C | ENSMUSG000 | ENSMUST000( |
| 40 | 39 | 1 | Zfp703,misser | A | ENSMUSG000 | ENSMUST000( |
| 192 | 192 | 0 | Kmt2a,missen | C | ENSMUSG000 | ENSMUST000( |
| 30 | 29 | 1 | Ammecr1,infr | - | ENSMUSG000 | ENSMUST000( |
| 110 | 110 | 0 | Rbm10,misser | T | ENSMUSG000 | ENSMUST000( |
| 72 | 71 | 1 | Amer1,missen | A | ENSMUSG000 | ENSMUST000( |
| 251 | 251 | 0 | Ros1,missense | A | ENSMUSG000 | ENSMUST000( |
| 358 | 358 | 0 | Npm1,framesl | CAGA | ENSMUSG000 | ENSMUST000( |
| 168 | 168 | 0 | Rnf43,synony | C | ENSMUSG000 | ENSMUST000( |
| 23 | 23 | 0 | Phlpp1,misser | T | ENSMUSG000 | ENSMUST000( |
| 137 | 137 | 0 | Elf3,synonymc | T | ENSMUSG000 | ENSMUST000( |
| 145 | 145 | 0 | Elf3,synonymc | A | ENSMUSG000 | ENSMUST000( |
| 9 | 8 | 1 | Sptbn5,infram | - | ENSMUSG000 | ENSMUST000( |
| 218 | 218 | 0 | Samhd1,synor | T | ENSMUSG000 | ENSMUST000( |
| 154 | 154 | 0 | Mapkap1,miss | C | ENSMUSG000 | ENSMUST000( |
| 243 | 233 | 10 | Arid1a,inframe | - | ENSMUSG000 | ENSMUST000( |
| 224 | 224 | 0 | Ptpn11,misser | T | ENSMUSG000 | ENSMUST000( |
| 84 | 82 | 2 | Gigyf1,inframe | - | ENSMUSG000 | ENSMUST000( |
| 145 | 145 | 0 | Flt3,missense_ | C | ENSMUSG000 | ENSMUST000( |
| 32 | 31 | 1 | Rbm33,infram | - | ENSMUSG000 | ENSMUST000( |
| 138 | 137 | 1 | Maz,synonym | T | ENSMUSG000 | ENSMUST000( |
| 145 | 145 | 0 | Maz,inframe_i | GCT | ENSMUSG000 | ENSMUST000( |
| 76 | 75 | 1 | Cdh16,missen | G | ENSMUSG000 | ENSMUST000( |
| 136 | 136 | 0 | Ankrd11,misse | G | ENSMUSG000 | ENSMUST000( |
| 30 | 29 | 1 | Ammecr1,infr | - | ENSMUSG000 | ENSMUST000( |
| 58 | 57 | 1 | Alg13,inframe | - | ENSMUSG000 | ENSMUST000( |
| 358 | 358 | 0 | Npm1,framesl | CAGA | ENSMUSG000 | ENSMUST000( |
| 193 | 193 | 0 | Tcf20,synonyn | T | ENSMUSG000 | ENSMUST000( |
| 22 | 22 | 0 | Vgll3,inframe_ | - | ENSMUSG000 | ENSMUST000( |
| 132 | 132 | 0 | Ccnd3,synony | G | ENSMUSG000 | ENSMUST000( |
| 10 | 10 | 0 | Olfr1463,miss | A | ENSMUSG000 | ENSMUST000( |
| 10 | 10 | 0 | Olfr1463,syno | T | ENSMUSG000 | ENSMUST000( |
| 39 | 39 | 0 | Chka,inframe_ | - | ENSMUSG000 | ENSMUST000( |
| 137 | 137 | 0 | Elf3,synonymc | T | ENSMUSG000 | ENSMUST000( |
| 145 | 145 | 0 | Elf3,synonymc | A | ENSMUSG000 | ENSMUST000( |
| 9 | 8 | 1 | Sptbn5,infram | - | ENSMUSG000 | ENSMUST000( |
| 224 | 224 | 0 | Ptpn11,misser | T | ENSMUSG000 | ENSMUST000( |
| 84 | 82 | 2 | Gigyf1,inframe | - | ENSMUSG000 | ENSMUST000( |
| 145 | 145 | 0 | Flt3,missense_ | C | ENSMUSG000 | ENSMUST000( |
| 138 | 137 | 1 | Maz,synonym | T | ENSMUSG000 | ENSMUST000( |
| 145 | 145 | 0 | Maz,inframe_i | GCT | ENSMUSG000 | ENSMUST000( |
| 161 | 160 | 0 | Zfhx3,synonyn | C | ENSMUSG000 | ENSMUST000( |
| 174 | 174 | 0 | Tgfbr2,missen | T | ENSMUSG000 | ENSMUST000( |
| 58 | 57 | 1 | Alg13,inframe | - | ENSMUSG000 | ENSMUST000( |
| 358 | 358 | 0 | Npm1,framesl | CAGA | ENSMUSG000 | ENSMUST000( |

| | | | | | | |
|---|---|---|---|---|---|---|
| 236 | 236 | 0 | Arid4b,missen | A | ENSMUSG000 | ENSMUST000( |
| 14 | 14 | 0 | Zfp503,infram | - | ENSMUSG000 | ENSMUST000( |
| 115 | 115 | 0 | Rb1,synonymc | T | ENSMUSG000 | ENSMUST000( |
| 112 | 112 | 0 | Ankhd1,misse | A | ENSMUSG000 | ENSMUST000( |
| 137 | 137 | 0 | Elf3,synonymc | T | ENSMUSG000 | ENSMUST000( |
| 145 | 145 | 0 | Elf3,synonymc | A | ENSMUSG000 | ENSMUST000( |
| 171 | 171 | 0 | Itpkb,missense | A | ENSMUSG000 | ENSMUST000( |
| 9 | 8 | 1 | Sptbn5,infram | - | ENSMUSG000 | ENSMUST000( |
| 41 | 35 | 6 | Ivl,inframe_de | - | ENSMUSG000 | ENSMUST000( |
| 224 | 224 | 0 | Ptpn11,misser | T | ENSMUSG000 | ENSMUST000( |
| 24 | 24 | 0 | Hectd4,infram | - | ENSMUSG000 | ENSMUST000( |
| 84 | 82 | 2 | Gigyf1,inframe | - | ENSMUSG000 | ENSMUST000( |
| 33 | 33 | 0 | Cckar,missens | T | ENSMUSG000 | ENSMUST000( |
| 138 | 137 | 1 | Maz,synonym | T | ENSMUSG000 | ENSMUST000( |
| 145 | 145 | 0 | Maz,inframe_i | GCT | ENSMUSG000 | ENSMUST000( |
| 38 | 38 | 0 | Rsf1,synonym | G | ENSMUSG000 | ENSMUST000( |
| 76 | 75 | 1 | Cdh16,missen | G | ENSMUSG000 | ENSMUST000( |
| 227 | 227 | 0 | Smarca4,misse | G | ENSMUSG000 | ENSMUST000( |
| 30 | 29 | 1 | Ammecr1,infra | - | ENSMUSG000 | ENSMUST000( |
| 12 | 12 | 0 | Sry,missense_ | G | ENSMUSG000 | ENSMUST000( |
| 17 | 16 | 1 | Sry,inframe_d | - | ENSMUSG000 | ENSMUST000( |
| 45 | 45 | 0 | Zfc3h1,infram | - | ENSMUSG000 | ENSMUST000( |
| 191 | 191 | 0 | Lats1,missense | G | ENSMUSG000 | ENSMUST000( |
| 358 | 358 | 0 | Npm1,framesh | CAGA | ENSMUSG000 | ENSMUST000( |
| 40 | 36 | 4 | Hcn1,inframe_ | - | ENSMUSG000 | ENSMUST000( |
| 115 | 115 | 0 | Rb1,synonymc | T | ENSMUSG000 | ENSMUST000( |
| 231 | 231 | 0 | Kmt2d,synony | T | ENSMUSG000 | ENSMUST000( |
| 22 | 22 | 0 | Vgll3,inframe_ | - | ENSMUSG000 | ENSMUST000( |
| 93 | 84 | 9 | Zfp318,infram | - | ENSMUSG000 | ENSMUST000( |
| 305 | 305 | 0 | Serpinb3b,mis | G | ENSMUSG000 | ENSMUST000( |
| 137 | 137 | 0 | Elf3,synonymc | T | ENSMUSG000 | ENSMUST000( |
| 145 | 145 | 0 | Elf3,synonymc | A | ENSMUSG000 | ENSMUST000( |
| 9 | 8 | 1 | Sptbn5,infram | - | ENSMUSG000 | ENSMUST000( |
| 211 | 211 | 0 | Lrp1b,synonyr | T | ENSMUSG000 | ENSMUST000( |
| 166 | 165 | 0 | Maml3,synony | T | ENSMUSG000 | ENSMUST000( |
| 151 | 150 | 1 | Zfhx4,synonyn | A | ENSMUSG000 | ENSMUST000( |
| 224 | 224 | 0 | Ptpn11,misser | T | ENSMUSG000 | ENSMUST000( |
| 84 | 82 | 2 | Gigyf1,inframe | - | ENSMUSG000 | ENSMUST000( |
| 196 | 196 | 0 | Kit,missense_ | T | ENSMUSG000 | ENSMUST000( |
| 257 | 257 | 0 | Kras,missense | T | ENSMUSG000 | ENSMUST000( |
| 147 | 147 | 0 | Plk1,missense | T | ENSMUSG000 | ENSMUST000( |
| 138 | 137 | 1 | Maz,synonym | T | ENSMUSG000 | ENSMUST000( |
| 145 | 145 | 0 | Maz,inframe_i | GCT | ENSMUSG000 | ENSMUST000( |
| 38 | 38 | 0 | Rsf1,synonym | G | ENSMUSG000 | ENSMUST000( |
| 165 | 165 | 0 | Mst1r,synonyr | G | ENSMUSG000 | ENSMUST000( |
| 358 | 358 | 0 | Npm1,framesh | CAGA | ENSMUSG000 | ENSMUST000( |
| 178 | 178 | 0 | Lats2,missense | G | ENSMUSG000 | ENSMUST000( |
| 22 | 22 | 0 | Vgll3,inframe_ | - | ENSMUSG000 | ENSMUST000( |
| 137 | 137 | 0 | Elf3,synonymc | T | ENSMUSG000 | ENSMUST000( |
| 145 | 145 | 0 | Elf3,synonymc | A | ENSMUSG000 | ENSMUST000( |

| | | | | | | |
|---|---|---|---|---|---|---|
| 9 | 8 | 1 | Sptbn5,infram | - | ENSMUSG000 | ENSMUST000( |
| 218 | 218 | 0 | Samhd1,synor | T | ENSMUSG000 | ENSMUST000( |
| 12 | 12 | 0 | Wdr63,missen | C | ENSMUSG000 | ENSMUST000( |
| 224 | 224 | 0 | Ptpn11,misser | T | ENSMUSG000 | ENSMUST000( |
| 84 | 82 | 2 | Gigyf1,inframe | - | ENSMUSG000 | ENSMUST000( |
| 145 | 145 | 0 | Flt3,missense_ | C | ENSMUSG000 | ENSMUST000( |
| 138 | 137 | 1 | Maz,synonym | T | ENSMUSG000 | ENSMUST000( |
| 145 | 145 | 0 | Maz,inframe_i | GCT | ENSMUSG000 | ENSMUST000( |
| 242 | 242 | 0 | Srcap,synonyn | C | ENSMUSG000 | ENSMUST000( |
| 120 | 120 | 0 | Ntrk3,missens | A | ENSMUSG000 | ENSMUST000( |
| 136 | 136 | 0 | Ankrd11,misse | G | ENSMUSG000 | ENSMUST000( |
| 22 | 22 | 0 | Acta1,synonyr | C | ENSMUSG000 | ENSMUST000( |
| 40 | 39 | 1 | Zfp703,misser | A | ENSMUSG000 | ENSMUST000( |
| 16 | 16 | 0 | Zrsr2,missense | C | ENSMUSG000 | ENSMUST000( |
| 358 | 358 | 0 | Npm1,framesl | CAGA | ENSMUSG000 | ENSMUST000( |
| 26 | 26 | 0 | Chd3,inframe_ | - | ENSMUSG000 | ENSMUST000( |
| 119 | 119 | 0 | Ep300,missen: | A | ENSMUSG000 | ENSMUST000( |
| 246 | 246 | 0 | Kmt2d,inframe | - | ENSMUSG000 | ENSMUST000( |
| 178 | 177 | 0 | Robo1,missen | T | ENSMUSG000 | ENSMUST000( |
| 9 | 8 | 1 | Sptbn5,infram | - | ENSMUSG000 | ENSMUST000( |
| 218 | 218 | 0 | Samhd1,synor | T | ENSMUSG000 | ENSMUST000( |
| 41 | 35 | 6 | Ivl,inframe_de | - | ENSMUSG000 | ENSMUST000( |
| 224 | 224 | 0 | Ptpn11,misser | T | ENSMUSG000 | ENSMUST000( |
| 84 | 82 | 2 | Gigyf1,inframe | - | ENSMUSG000 | ENSMUST000( |
| 145 | 145 | 0 | Flt3,missense_ | C | ENSMUSG000 | ENSMUST000( |
| 32 | 31 | 1 | Rbm33,infram | - | ENSMUSG000 | ENSMUST000( |
| 138 | 137 | 1 | Maz,synonym | T | ENSMUSG000 | ENSMUST000( |
| 145 | 145 | 0 | Maz,inframe_i | GCT | ENSMUSG000 | ENSMUST000( |
| 33 | 33 | 0 | Grin2d,missen | C | ENSMUSG000 | ENSMUST000( |
| 34 | 33 | 1 | Rsf1,missense | A | ENSMUSG000 | ENSMUST000( |
| 136 | 136 | 0 | Ankrd11,misse | G | ENSMUSG000 | ENSMUST000( |
| 40 | 39 | 1 | Zfp703,misser | A | ENSMUSG000 | ENSMUST000( |
| 101 | 101 | 0 | Zmym3,synon | C | ENSMUSG000 | ENSMUST000( |
| 30 | 29 | 1 | Ammecr1,infr | - | ENSMUSG000 | ENSMUST000( |
| 110 | 110 | 0 | Rbm10,misser | T | ENSMUSG000 | ENSMUST000( |
| 72 | 71 | 1 | Amer1,missen | A | ENSMUSG000 | ENSMUST000( |
| 358 | 358 | 0 | Npm1,framesl | CAGA | ENSMUSG000 | ENSMUST000( |
| 184 | 183 | 1 | Nf1,stop_gain | T | ENSMUSG000 | ENSMUST000( |
| 41 | 40 | 1 | Fam170b,miss | C | ENSMUSG000 | ENSMUST000( |
| 246 | 246 | 0 | Kmt2d,inframe | - | ENSMUSG000 | ENSMUST000( |
| 156 | 156 | 0 | Arid1b,synony | T | ENSMUSG000 | ENSMUST000( |
| 9 | 8 | 1 | Sptbn5,infram | - | ENSMUSG000 | ENSMUST000( |
| 41 | 35 | 6 | Ivl,inframe_de | - | ENSMUSG000 | ENSMUST000( |
| 243 | 233 | 10 | Arid1a,inframe | - | ENSMUSG000 | ENSMUST000( |
| 224 | 224 | 0 | Ptpn11,misser | T | ENSMUSG000 | ENSMUST000( |
| 32 | 31 | 1 | Rbm33,infram | - | ENSMUSG000 | ENSMUST000( |
| 170 | 170 | 0 | Kdm5a,misser | A | ENSMUSG000 | ENSMUST000( |
| 201 | 201 | 0 | D6Ertd527e,sy | T | ENSMUSG000 | ENSMUST000( |
| 138 | 137 | 1 | Maz,synonym | T | ENSMUSG000 | ENSMUST000( |
| 145 | 145 | 0 | Maz,inframe_i | GCT | ENSMUSG000 | ENSMUST000( |

| | | | | | | |
|---|---|---|---|---|---|---|
| 242 | 242 | 0 | Srcap,synonyn | C | ENSMUSG000 | ENSMUST000( |
| 33 | 33 | 0 | Grin2d,missen | C | ENSMUSG000 | ENSMUST000( |
| 155 | 155 | 0 | Kmt2a,missen | T | ENSMUSG000 | ENSMUST000( |
| 30 | 29 | 1 | Ammecr1,infr | - | ENSMUSG000 | ENSMUST000( |
| 58 | 57 | 1 | Alg13,inframe | - | ENSMUSG000 | ENSMUST000( |
| 110 | 110 | 0 | Rbm10,misser | T | ENSMUSG000 | ENSMUST000( |

| Feature_type | Consequence | cDNA_positior | CDS_position | Protein_positi | Amino_acids | Codons |
|---|---|---|---|---|---|---|
| Transcript | missense_vari | 775/2395 | 460/1461 | 154/486 | F/L | Ttt/Ctt |
| Transcript | frameshift_va | 1084-1085/16 | 857-858/879 | 286/292 | W/CLX | tgg/tgTCTGg |
| Transcript | synonymous_ | 4262/6582 | 4239/6252 | 1413/2083 | G | ggA/ggG |
| Transcript | missense_vari | 726/6871 | 643/1287 | 215/428 | V/I | Gta/Ata |
| Transcript | inframe_delet | 9906-9908/11 | 9906-9908/10 | 3302-3303/36 | QH/H | caGCAt/cat |
| Transcript | synonymous_ | 401/3926 | 318/1977 | 106/658 | A | gcC/gcA |
| Transcript | inframe_delet | 520-522/1922 | 453-455/1407 | 151-152/468 | QQ/Q | caGCAa/caa |
| Transcript | missense_vari | 340/5535 | 226/1794 | 76/597 | E/K | Gaa/Aaa |
| Transcript | inframe_delet | 2990-2992/54 | 2916-2918/31 | 972-973/1044 | RQ/R | cgGCAg/cgg |
| Transcript | missense_vari | 2749/3657 | 2525/3003 | 842/1000 | D/G | gAc/gGc |
| Transcript | missense_vari | 1123/4407 | 1123/3066 | 375/1021 | A/T | Gcc/Acc |
| Transcript | synonymous_ | 1356/2347 | 1323/1434 | 441/477 | A | gcG/gcA |
| Transcript | inframe_inser | 1355-1356/23 | 1322-1323/14 | 441/477 | A/AA | gcg/gcAGCg |
| Transcript | missense_vari | 2032/6487 | 1339/5151 | 447/1716 | S/G | Agt/Ggt |
| Transcript | missense_vari | 144/491 | 61/279 | 21/92 | D/N | Gat/Aat |
| Transcript | missense_vari | 14/11124 | 7/4326 | 3/1441 | A/T | Gcg/Acg |
| Transcript | missense_vari | 7062/8455 | 6868/7932 | 2290/2643 | V/L | Gtt/Ctt |
| Transcript | missense_vari | 1266/11372 | 1105/2742 | 369/913 | Y/N | Tac/Aac |
| Transcript | missense_vari | 914/3564 | 360/2793 | 120/930 | E/D | gaG/gaT |
| Transcript | missense_vari | 1295/2965 | 943/1413 | 315/470 | R/W | Cgg/Tgg |
| Transcript | frameshift_va | 1084-1085/16 | 857-858/879 | 286/292 | W/CLX | tgg/tgTCTGg |
| Transcript | missense_vari | 4520/11917 | 4345/8526 | 1449/2841 | Q/K | Cag/Aag |
| Transcript | stop_gained | 4607/11917 | 4432/8526 | 1478/2841 | R/* | Cga/Tga |
| Transcript | inframe_delet | 563-565/7009 | 187-189/978 | 63/325 | E/- | GAG/- |
| Transcript | inframe_delet | 673-675/4423 | 266-268/1893 | 89-90/630 | QP/P | cAGCct/cct |
| Transcript | missense_vari | 587/3148 | 190/1761 | 64/586 | N/H | Aac/Cac |
| Transcript | inframe_delet | 1041-1043/81 | 1041-1043/68 | 347-348/2283 | AA/A | gcGGCa/gca |
| Transcript | synonymous_ | 1809/3173 | 1644/2265 | 548/754 | A | gcG/gcA |
| Transcript | missense_vari | 340/5535 | 226/1794 | 76/597 | E/K | Gaa/Aaa |
| Transcript | missense_vari | 2749/3657 | 2525/3003 | 842/1000 | D/G | gAc/gGc |
| Transcript | synonymous_ | 1356/2347 | 1323/1434 | 441/477 | A | gcG/gcA |
| Transcript | inframe_inser | 1355-1356/23 | 1322-1323/14 | 441/477 | A/AA | gcg/gcAGCg |
| Transcript | frameshift_va | 1084-1085/16 | 857-858/879 | 286/292 | W/CLX | tgg/tgTCTGg |
| Transcript | frameshift_va | 5138-5147/11 | 4963-4972/85 | 1655-1658/28 | RFKT/X | CGCTTTAAAA |
| Transcript | synonymous_ | 232/4656 | 51/2766 | 17/921 | P | ccG/ccA |
| Transcript | missense_vari | 1052/1653 | 992/1164 | 331/387 | F/S | tTt/tCt |
| Transcript | inframe_delet | 9906-9908/11 | 9906-9908/10 | 3302-3303/36 | QH/H | caGCAt/cat |
| Transcript | inframe_delet | 520-522/1922 | 453-455/1407 | 151-152/468 | QQ/Q | caGCAa/caa |
| Transcript | missense_vari | 340/5535 | 226/1794 | 76/597 | E/K | Gaa/Aaa |
| Transcript | inframe_delet | 2990-2992/54 | 2916-2918/31 | 972-973/1044 | RQ/R | cgGCAg/cgg |
| Transcript | synonymous_ | 1347/2218 | 1269/1395 | 423/464 | S | agC/agT |
| Transcript | missense_vari | 1194/2199 | 1090/1812 | 364/603 | R/W | Cgg/Tgg |
| Transcript | synonymous_ | 1356/2347 | 1323/1434 | 441/477 | A | gcG/gcA |
| Transcript | inframe_inser | 1355-1356/23 | 1322-1323/14 | 441/477 | A/AA | gcg/gcAGCg |
| Transcript | missense_vari | 478/774 | 181/438 | 61/145 | A/T | Gca/Aca |
| Transcript | missense_vari | 2446/2874 | 2317/2493 | 773/830 | M/L | Atg/Ctg |
| Transcript | inframe_delet | 283-285/5437 | 268-270/1035 | 90/344 | G/- | GGC/- |
| Transcript | frameshift_va | 1084-1085/16 | 857-858/879 | 286/292 | W/CLX | tgg/tgTCTGg |
| Transcript | inframe_delet | 93-95/5456 | 78-80/3624 | 26-27/1207 | GG/G | ggCGGt/ggt |

| | | | | | | |
|---|---|---|---|---|---|---|
| Transcript | inframe_delet | 2490-2492/14 | 2266-2268/27 | 756/910 | Q/- | CAG/- |
| Transcript | synonymous_\ | 369/3371 | 15/2133 | 5/710 | P | ccG/ccA |
| Transcript | missense_vari | 844/1694 | 730/1137 | 244/378 | T/P | Acc/Ccc |
| Transcript | synonymous_\ | 669/5219 | 435/2256 | 145/751 | S | agC/agT |
| Transcript | inframe_delet | 10080-10082/ | 9852-9854/16 | 3284-3285/55 | QH/H | caGCAt/cat |
| Transcript | inframe_delet | 563-565/7009 | 187-189/978 | 63/325 | E/- | GAG/- |
| Transcript | inframe_delet | 9906-9908/11 | 9906-9908/10 | 3302-3303/36 | QH/H | caGCAt/cat |
| Transcript | synonymous_\ | 2406/8288 | 1473/3408 | 491/1135 | Q | caG/caA |
| Transcript | inframe_delet | 520-522/1922 | 453-455/1407 | 151-152/468 | QQ/Q | caGCAa/caa |
| Transcript | missense_vari | 340/5535 | 226/1794 | 76/597 | E/K | Gaa/Aaa |
| Transcript | inframe_delet | 2828-2830/95 | 2637-2639/36 | 879-880/1231 | AQ/A | gcGCAg/gcg |
| Transcript | missense_vari | 1120/2930 | 772/1311 | 258/436 | G/S | Ggc/Agc |
| Transcript | synonymous_\ | 1356/2347 | 1323/1434 | 441/477 | A | gcG/gcA |
| Transcript | inframe_inser | 1355-1356/23 | 1322-1323/14 | 441/477 | A/AA | gcg/gcAGCg |
| Transcript | missense_vari | 7742/10654 | 7354/9816 | 2452/3271 | P/A | Cct/Gct |
| Transcript | missense_vari | 2446/2874 | 2317/2493 | 773/830 | M/L | Atg/Ctg |
| Transcript | missense_vari | 73/3152 | 73/1785 | 25/594 | G/S | Ggc/Agc |
| Transcript | inframe_delet | 283-285/5437 | 268-270/1035 | 90/344 | G/- | GGC/- |
| Transcript | missense_vari | 914/3564 | 360/2793 | 120/930 | E/D | gaG/gaT |
| Transcript | inframe_delet | 712-714/1188 | 712-714/1188 | 238/395 | Q/- | CAG/- |
| Transcript | frameshift_va | 1084-1085/16 | 857-858/879 | 286/292 | W/CLX | tgg/tgTCTGg |
| Transcript | stop_gained | 6986/11917 | 6811/8526 | 2271/2841 | R/* | Cga/Tga |
| Transcript | inframe_delet | 93-95/5456 | 78-80/3624 | 26-27/1207 | GG/G | ggCGGt/ggt |
| Transcript | inframe_delet | 1407-1409/42 | 79-81/1959 | 27/652 | G/- | GGC/- |
| Transcript | inframe_delet | 563-565/7009 | 187-189/978 | 63/325 | E/- | GAG/- |
| Transcript | inframe_delet | 9906-9908/11 | 9906-9908/10 | 3302-3303/36 | QH/H | caGCAt/cat |
| Transcript | inframe_delet | 2868-2870/54 | 2725-2727/45 | 909/1510 | Q/- | CAG/- |
| Transcript | missense_vari | 340/5535 | 226/1794 | 76/597 | E/K | Gaa/Aaa |
| Transcript | inframe_delet | 2990-2992/54 | 2916-2918/31 | 972-973/1044 | RQ/R | cgGCAg/cgg |
| Transcript | missense_vari | 2749/3657 | 2525/3003 | 842/1000 | D/G | gAc/gGc |
| Transcript | missense_vari | 1120/2930 | 772/1311 | 258/436 | G/S | Ggc/Agc |
| Transcript | inframe_delet | 283-285/5437 | 268-270/1035 | 90/344 | G/- | GGC/- |
| Transcript | synonymous_\ | 1624/4796 | 1467/2412 | 489/803 | Q | caG/caA |
| Transcript | missense_vari | 1449/8496 | 1146/3399 | 382/1132 | E/D | gaA/gaT |
| Transcript | frameshift_va | 1084-1085/16 | 857-858/879 | 286/292 | W/CLX | tgg/tgTCTGg |
| Transcript | synonymous_\ | 131/7266 | 51/3642 | 17/1213 | L | ctG/ctA |
| Transcript | stop_gained | 3893/11917 | 3718/8526 | 1240/2841 | E/* | Gag/Tag |
| Transcript | inframe_delet | 2490-2492/14 | 2266-2268/27 | 756/910 | Q/- | CAG/- |
| Transcript | inframe_delet | 12372-12374/ | 12144-12146/ | 4048-4049/55 | QQ/Q | caGCAa/caa |
| Transcript | inframe_delet | 9906-9908/11 | 9906-9908/10 | 3302-3303/36 | QH/H | caGCAt/cat |
| Transcript | synonymous_\ | 8027/10608 | 7956/9108 | 2652/3035 | Q | caG/caA |
| Transcript | missense_vari | 340/5535 | 226/1794 | 76/597 | E/K | Gaa/Aaa |
| Transcript | inframe_delet | 2828-2830/95 | 2637-2639/36 | 879-880/1231 | AQ/A | gcGCAg/gcg |
| Transcript | synonymous_\ | 1347/2218 | 1269/1395 | 423/464 | S | agC/agT |
| Transcript | synonymous_\ | 1356/2347 | 1323/1434 | 441/477 | A | gcG/gcA |
| Transcript | inframe_inser | 1355-1356/23 | 1322-1323/14 | 441/477 | A/AA | gcg/gcAGCg |
| Transcript | missense_vari | 14/11124 | 7/4326 | 3/1441 | A/T | Gcg/Acg |
| Transcript | synonymous_\ | 40/11124 | 33/4326 | 11/1441 | A | gcA/gcG |
| Transcript | missense_vari | 73/3152 | 73/1785 | 25/594 | G/S | Ggc/Agc |
| Transcript | synonymous_\ | 1297/4985 | 381/2178 | 127/725 | P | ccG/ccA |

| | | | | | | |
|---|---|---|---|---|---|---|
| Transcript | missense_vari | 369/6957 | 130/5979 | 44/1992 | S/G | Agc/Ggc |
| Transcript | missense_vari | 507/1952 | 383/1128 | 128/375 | N/T | aAt/aCt |
| Transcript | frameshift_va | 1084-1085/16 | 857-858/879 | 286/292 | W/CLX | tgg/tgTCTGg |
| Transcript | stop_gained | 304/9592 | 9/8652 | 3/2883 | C/* | tgC/tgA |
| Transcript | inframe_delet | 1333-1335/73 | 1165-1167/51 | 389/1708 | E/- | GAG/- |
| Transcript | missense_vari | 320/8223 | 193/7647 | 65/2548 | G/S | Ggc/Agc |
| Transcript | synonymous_v | 657/2095 | 429/1176 | 143/391 | Q | caG/caA |
| Transcript | synonymous_v | 558/2095 | 330/1176 | 110/391 | D | gaC/gaT |
| Transcript | synonymous_v | 541/3090 | 273/1554 | 91/517 | G | ggC/ggT |
| Transcript | inframe_delet | 9906-9908/11 | 9906-9908/10 | 3302-3303/36 | QH/H | caGCAt/cat |
| Transcript | missense_vari | 540/660 | 540/660 | 180/220 | F/L | ttC/ttG |
| Transcript | inframe_delet | 1041-1043/81 | 1041-1043/68 | 347-348/2283 | AA/A | gcGGCa/gca |
| Transcript | synonymous_v | 788/6069 | 519/1710 | 173/569 | S | agC/agT |
| Transcript | missense_vari | 340/5535 | 226/1794 | 76/597 | E/K | Gaa/Aaa |
| Transcript | inframe_delet | 287-289/1548 | 34-36/13257 | 12/4418 | A/- | GCG/- |
| Transcript | synonymous_v | 3858/8857 | 3582/5958 | 1194/1985 | E | gaG/gaA |
| Transcript | inframe_delet | 2990-2992/54 | 2916-2918/31 | 972-973/1044 | RQ/R | cgGCAg/cgg |
| Transcript | missense_vari | 2749/3657 | 2525/3003 | 842/1000 | D/G | gAc/gGc |
| Transcript | missense_vari | 2382/5214 | 2285/2940 | 762/979 | D/G | gAc/gGc |
| Transcript | synonymous_v | 1356/2347 | 1323/1434 | 441/477 | A | gcG/gcA |
| Transcript | inframe_inser | 1355-1356/23 | 1322-1323/14 | 441/477 | A/AA | gcg/gcAGCg |
| Transcript | missense_vari | 2032/6487 | 1339/5151 | 447/1716 | S/G | Agt/Ggt |
| Transcript | missense_vari | 10214/14814 | 10021/13773 | 3341/4590 | T/A | Acc/Gcc |
| Transcript | inframe_delet | 283-285/5437 | 268-270/1035 | 90/344 | G/- | GGC/- |
| Transcript | frameshift_va | 1084-1085/16 | 857-858/879 | 286/292 | W/CLX | tgg/tgTCTGg |
| Transcript | inframe_delet | 517-519/2942 | 465-467/1935 | 155-156/644 | QQ/Q | caGCAa/caa |
| Transcript | inframe_delet | 563-565/7009 | 187-189/978 | 63/325 | E/- | GAG/- |
| Transcript | missense_vari | 1212/1973 | 733/1179 | 245/392 | E/K | Gag/Aag |
| Transcript | inframe_delet | 168-170/2319 | 168-170/1362 | 56-57/453 | LP/L | ctGCCg/ctg |
| Transcript | inframe_delet | 9906-9908/11 | 9906-9908/10 | 3302-3303/36 | QH/H | caGCAt/cat |
| Transcript | inframe_delet | 1041-1043/81 | 1041-1043/68 | 347-348/2283 | AA/A | gcGGCa/gca |
| Transcript | synonymous_v | 737/6069 | 468/1710 | 156/569 | S | agC/agT |
| Transcript | missense_vari | 340/5535 | 226/1794 | 76/597 | E/K | Gaa/Aaa |
| Transcript | synonymous_v | 1356/2347 | 1323/1434 | 441/477 | A | gcG/gcA |
| Transcript | inframe_inser | 1355-1356/23 | 1322-1323/14 | 441/477 | A/AA | gcg/gcAGCg |
| Transcript | synonymous_v | 3499/10654 | 3111/9816 | 1037/3271 | R | cgT/cgC |
| Transcript | missense_vari | 73/3152 | 73/1785 | 25/594 | G/S | Ggc/Agc |
| Transcript | missense_vari | 2066/3528 | 1552/1614 | 518/537 | G/S | Ggc/Agc |
| Transcript | frameshift_va | 1084-1085/16 | 857-858/879 | 286/292 | W/CLX | tgg/tgTCTGg |
| Transcript | inframe_delet | 563-565/7009 | 187-189/978 | 63/325 | E/- | GAG/- |
| Transcript | synonymous_v | 895/3249 | 744/1461 | 248/486 | T | acG/acT |
| Transcript | synonymous_v | 247/8436 | 201/3960 | 67/1319 | A | gcT/gcC |
| Transcript | missense_vari | 804/2315 | 638/933 | 213/310 | I/K | aTa/aAa |
| Transcript | synonymous_v | 805/2315 | 639/933 | 213/310 | I | atA/atT |
| Transcript | synonymous_v | 657/2095 | 429/1176 | 143/391 | Q | caG/caA |
| Transcript | synonymous_v | 558/2095 | 330/1176 | 110/391 | D | gaC/gaT |
| Transcript | inframe_delet | 9906-9908/11 | 9906-9908/10 | 3302-3303/36 | QH/H | caGCAt/cat |
| Transcript | synonymous_v | 2142/10671 | 1944/6693 | 648/2230 | Q | caG/caA |
| Transcript | missense_vari | 6864/10500 | 6703/7422 | 2235/2473 | G/C | Ggt/Tgt |
| Transcript | missense_vari | 340/5535 | 226/1794 | 76/597 | E/K | Gaa/Aaa |

| | | | | | | |
|---|---|---|---|---|---|---|
| Transcript | inframe_delet | 2990-2992/54 | 2916-2918/31 | 972-973/1044 | RQ/R | cgGCAg/cgg |
| Transcript | missense_vari | 2749/3657 | 2525/3003 | 842/1000 | D/G | gAc/gGc |
| Transcript | synonymous_ | 1356/2347 | 1323/1434 | 441/477 | A | gcG/gcA |
| Transcript | inframe_inser | 1355-1356/23 | 1322-1323/14 | 441/477 | A/AA | gcg/gcAGCg |
| Transcript | missense_vari | 2995/5431 | 2433/3972 | 811/1323 | D/E | gaT/gaG |
| Transcript | missense_vari | 1161/19703 | 536/2478 | 179/825 | E/G | gAg/gGg |
| Transcript | missense_vari | 73/3152 | 73/1785 | 25/594 | G/S | Ggc/Agc |
| Transcript | missense_vari | 2697/16461 | 2675/11892 | 892/3963 | K/R | aAg/aGg |
| Transcript | inframe_delet | 283-285/5437 | 268-270/1035 | 90/344 | G/- | GGC/- |
| Transcript | missense_vari | 914/3564 | 360/2793 | 120/930 | E/D | gaG/gaT |
| Transcript | missense_vari | 1449/8496 | 1146/3399 | 382/1132 | E/D | gaA/gaT |
| Transcript | missense_vari | 4483/7401 | 4195/7023 | 1399/2340 | A/S | Gcc/Tcc |
| Transcript | frameshift_va | 1084-1085/16 | 857-858/879 | 286/292 | W/CLX | tgg/tgTCTGg |
| Transcript | synonymous_ | 2110/4310 | 1494/2355 | 498/784 | R | cgA/cgC |
| Transcript | missense_vari | 5195/6226 | 4943/5064 | 1648/1687 | Q/L | cAg/cTg |
| Transcript | synonymous_ | 657/2095 | 429/1176 | 143/391 | Q | caG/caA |
| Transcript | synonymous_ | 558/2095 | 330/1176 | 110/391 | D | gaC/gaT |
| Transcript | inframe_delet | 9906-9908/11 | 9906-9908/10 | 3302-3303/36 | QH/H | caGCAt/cat |
| Transcript | synonymous_ | 401/3926 | 318/1977 | 106/658 | A | gcC/gcA |
| Transcript | missense_vari | 683/3111 | 542/1569 | 181/522 | L/P | cTc/cCc |
| Transcript | inframe_delet | 1041-1043/81 | 1041-1043/68 | 347-348/2283 | AA/A | gcGGCa/gca |
| Transcript | missense_vari | 340/5535 | 226/1794 | 76/597 | E/K | Gaa/Aaa |
| Transcript | inframe_delet | 2990-2992/54 | 2916-2918/31 | 972-973/1044 | RQ/R | cgGCAg/cgg |
| Transcript | missense_vari | 2749/3657 | 2525/3003 | 842/1000 | D/G | gAc/gGc |
| Transcript | inframe_delet | 2828-2830/95 | 2637-2639/36 | 879-880/1231 | AQ/A | gcGCAg/gcg |
| Transcript | synonymous_ | 1356/2347 | 1323/1434 | 441/477 | A | gcG/gcA |
| Transcript | inframe_inser | 1355-1356/23 | 1322-1323/14 | 441/477 | A/AA | gcg/gcAGCg |
| Transcript | missense_vari | 2446/2874 | 2317/2493 | 773/830 | M/L | Atg/Ctg |
| Transcript | missense_vari | 7062/8455 | 6868/7932 | 2290/2643 | V/L | Gtt/Ctt |
| Transcript | inframe_delet | 283-285/5437 | 268-270/1035 | 90/344 | G/- | GGC/- |
| Transcript | inframe_delet | 2738-2740/41 | 2103-2105/29 | 701-702/969 | GA/G | ggAGCa/gga |
| Transcript | frameshift_va | 1084-1085/16 | 857-858/879 | 286/292 | W/CLX | tgg/tgTCTGg |
| Transcript | synonymous_ | 707/7323 | 549/5964 | 183/1987 | Q | caG/caA |
| Transcript | inframe_delet | 563-565/7009 | 187-189/978 | 63/325 | E/- | GAG/- |
| Transcript | synonymous_ | 1026/2113 | 699/879 | 233/292 | T | acA/acG |
| Transcript | missense_vari | 804/2315 | 638/933 | 213/310 | I/K | aTa/aAa |
| Transcript | synonymous_ | 805/2315 | 639/933 | 213/310 | I | atA/atT |
| Transcript | inframe_delet | 168-170/2319 | 168-170/1362 | 56-57/453 | LP/L | ctGCCg/ctg |
| Transcript | synonymous_ | 657/2095 | 429/1176 | 143/391 | Q | caG/caA |
| Transcript | synonymous_ | 558/2095 | 330/1176 | 110/391 | D | gaC/gaT |
| Transcript | inframe_delet | 9906-9908/11 | 9906-9908/10 | 3302-3303/36 | QH/H | caGCAt/cat |
| Transcript | missense_vari | 340/5535 | 226/1794 | 76/597 | E/K | Gaa/Aaa |
| Transcript | inframe_delet | 2990-2992/54 | 2916-2918/31 | 972-973/1044 | RQ/R | cgGCAg/cgg |
| Transcript | missense_vari | 2749/3657 | 2525/3003 | 842/1000 | D/G | gAc/gGc |
| Transcript | synonymous_ | 1356/2347 | 1323/1434 | 441/477 | A | gcG/gcA |
| Transcript | inframe_inser | 1355-1356/23 | 1322-1323/14 | 441/477 | A/AA | gcg/gcAGCg |
| Transcript | synonymous_ | 7611/17152 | 6159/11172 | 2053/3723 | P | ccT/ccC |
| Transcript | missense_vari | 997/8092 | 769/1779 | 257/592 | A/T | Gcc/Acc |
| Transcript | inframe_delet | 2738-2740/41 | 2103-2105/29 | 701-702/969 | GA/G | ggAGCa/gga |
| Transcript | frameshift_va | 1084-1085/16 | 857-858/879 | 286/292 | W/CLX | tgg/tgTCTGg |

| | | | | | | |
|---|---|---|---|---|---|---|
| Transcript | missense_vari | 2725/5864 | 2435/3945 | 812/1314 | T/K | aCa/aAa |
| Transcript | inframe_delet | 1407-1409/42 | 79-81/1959 | 27/652 | G/- | GGC/- |
| Transcript | synonymous_\ | 232/4656 | 51/2766 | 17/921 | P | ccG/ccA |
| Transcript | missense_vari | 320/8223 | 193/7647 | 65/2548 | G/S | Ggc/Agc |
| Transcript | synonymous_\ | 657/2095 | 429/1176 | 143/391 | Q | caG/caA |
| Transcript | synonymous_\ | 558/2095 | 330/1176 | 110/391 | D | gaC/gaT |
| Transcript | missense_vari | 781/6235 | 287/2829 | 96/942 | S/N | aGc/aAc |
| Transcript | inframe_delet | 9906-9908/11 | 9906-9908/10 | 3302-3303/36 | QH/H | caGCAt/cat |
| Transcript | inframe_delet | 520-522/1922 | 453-455/1407 | 151-152/468 | QQ/Q | caGCAa/caa |
| Transcript | missense_vari | 340/5535 | 226/1794 | 76/597 | E/K | Gaa/Aaa |
| Transcript | inframe_delet | 287-289/1548 | 34-36/13257 | 12/4418 | A/- | GCG/- |
| Transcript | inframe_delet | 2990-2992/54 | 2916-2918/31 | 972-973/1044 | RQ/R | cgGCAg/cgg |
| Transcript | missense_vari | 1120/2930 | 772/1311 | 258/436 | G/S | Ggc/Agc |
| Transcript | synonymous_\ | 1356/2347 | 1323/1434 | 441/477 | A | gcG/gcA |
| Transcript | inframe_inser | 1355-1356/23 | 1322-1323/14 | 441/477 | A/AA | gcg/gcAGCg |
| Transcript | synonymous_\ | 40/11124 | 33/4326 | 11/1441 | A | gcA/gcG |
| Transcript | missense_vari | 2446/2874 | 2317/2493 | 773/830 | M/L | Atg/Ctg |
| Transcript | missense_vari | 4909/6376 | 4649/4854 | 1550/1617 | E/G | gAg/gGg |
| Transcript | inframe_delet | 283-285/5437 | 268-270/1035 | 90/344 | G/- | GGC/- |
| Transcript | missense_vari | 795/1188 | 795/1188 | 265/395 | Q/H | caG/caC |
| Transcript | inframe_delet | 460-462/1188 | 460-462/1188 | 154/395 | Q/- | CAG/- |
| Transcript | inframe_delet | 351-353/6957 | 112-114/5979 | 38/1992 | S/- | AGC/- |
| Transcript | missense_vari | 2455/7209 | 2086/3390 | 696/1129 | R/G | Agg/Ggg |
| Transcript | frameshift_va | 1084-1085/16 | 857-858/879 | 286/292 | W/CLX | tgg/tgTCTGg |
| Transcript | inframe_delet | 2490-2492/14 | 2266-2268/27 | 756/910 | Q/- | CAG/- |
| Transcript | synonymous_\ | 232/4656 | 51/2766 | 17/921 | P | ccG/ccA |
| Transcript | synonymous_\ | 11031/19823 | 10803/16767 | 3601/5588 | Q | caG/caA |
| Transcript | inframe_delet | 563-565/7009 | 187-189/978 | 63/325 | E/- | GAG/- |
| Transcript | inframe_delet | 5551-5553/13 | 5443-5445/67 | 1815/2237 | E/- | GAA/- |
| Transcript | missense_vari | 1052/1653 | 992/1164 | 331/387 | F/S | tTt/tCt |
| Transcript | synonymous_\ | 657/2095 | 429/1176 | 143/391 | Q | caG/caA |
| Transcript | synonymous_\ | 558/2095 | 330/1176 | 110/391 | D | gaC/gaT |
| Transcript | inframe_delet | 9906-9908/11 | 9906-9908/10 | 3302-3303/36 | QH/H | caGCAt/cat |
| Transcript | synonymous_\ | 3124/16294 | 2316/13800 | 772/4599 | T | acG/acA |
| Transcript | synonymous_\ | 2406/8288 | 1473/3408 | 491/1135 | Q | caG/caA |
| Transcript | synonymous_\ | 6486/13874 | 6066/10746 | 2022/3581 | P | ccT/ccA |
| Transcript | missense_vari | 340/5535 | 226/1794 | 76/597 | E/K | Gaa/Aaa |
| Transcript | inframe_delet | 2990-2992/54 | 2916-2918/31 | 972-973/1044 | RQ/R | cgGCAg/cgg |
| Transcript | missense_vari | 2549/5214 | 2452/2940 | 818/979 | D/Y | Gac/Tac |
| Transcript | missense_vari | 237/4678 | 35/567 | 12/188 | G/D | gGt/gAt |
| Transcript | missense_vari | 1194/2199 | 1090/1812 | 364/603 | R/W | Cgg/Tgg |
| Transcript | synonymous_\ | 1356/2347 | 1323/1434 | 441/477 | A | gcG/gcA |
| Transcript | inframe_inser | 1355-1356/23 | 1322-1323/14 | 441/477 | A/AA | gcg/gcAGCg |
| Transcript | synonymous_\ | 40/11124 | 33/4326 | 11/1441 | A | gcA/gcG |
| Transcript | synonymous_\ | 1883/4722 | 1611/4137 | 537/1378 | R | cgT/cgG |
| Transcript | frameshift_va | 1084-1085/16 | 857-858/879 | 286/292 | W/CLX | tgg/tgTCTGg |
| Transcript | missense_vari | 3389/5191 | 2953/3129 | 985/1042 | S/P | Tcc/Ccc |
| Transcript | inframe_delet | 563-565/7009 | 187-189/978 | 63/325 | E/- | GAG/- |
| Transcript | synonymous_\ | 657/2095 | 429/1176 | 143/391 | Q | caG/caA |
| Transcript | synonymous_\ | 558/2095 | 330/1176 | 110/391 | D | gaC/gaT |

| | | | | | | | |
|---|---|---|---|---|---|---|---|
| Transcript | inframe_delet | 9906-9908/11 | 9906-9908/10 | 3302-3303/36 | QH/H | | caGCAt/cat |
| Transcript | synonymous_ | 401/3926 | 318/1977 | 106/658 | A | | gcC/gcA |
| Transcript | missense_vari | 273/3176 | 41/2772 | 14/923 | K/R | | aAg/aGg |
| Transcript | missense_vari | 340/5535 | 226/1794 | 76/597 | E/K | | Gaa/Aaa |
| Transcript | inframe_delet | 2990-2992/54 | 2916-2918/31 | 972-973/1044 | RQ/R | | cgGCAg/cgg |
| Transcript | missense_vari | 2749/3657 | 2525/3003 | 842/1000 | D/G | | gAc/gGc |
| Transcript | synonymous_ | 1356/2347 | 1323/1434 | 441/477 | A | | gcG/gcA |
| Transcript | inframe_inser | 1355-1356/23 | 1322-1323/14 | 441/477 | A/AA | | gcg/gcAGCg |
| Transcript | synonymous_ | 3499/10654 | 3111/9816 | 1037/3271 | R | | cgT/cgC |
| Transcript | missense_vari | 2058/19703 | 1433/2478 | 478/825 | A/V | | gCc/gTc |
| Transcript | missense_vari | 7062/8455 | 6868/7932 | 2290/2643 | V/L | | Gtt/Ctt |
| Transcript | synonymous_ | 519/1446 | 447/1134 | 149/377 | R | | cgT/cgG |
| Transcript | missense_vari | 73/3152 | 73/1785 | 25/594 | G/S | | Ggc/Agc |
| Transcript | missense_vari | 1314/2735 | 1051/1626 | 351/541 | Y/D | | Tac/Gac |
| Transcript | frameshift_va | 1084-1085/16 | 857-858/879 | 286/292 | W/CLX | | tgg/tgTCTGg |
| Transcript | inframe_delet | 248-250/7261 | 233-235/6168 | 78-79/2055 | PL/L | | cCGCtg/ctg |
| Transcript | missense_vari | 2617/9566 | 1385/7239 | 462/2412 | P/H | | cCc/cAc |
| Transcript | inframe_delet | 8568-8603/19 | 8340-8375/16 | 2780-2792/55 | QQQQQQQQ | | caACAACAGC |
| Transcript | missense_vari | 5577/7563 | 4586/4839 | 1529/1612 | A/V | | gCc/gTc |
| Transcript | inframe_delet | 9906-9908/11 | 9906-9908/10 | 3302-3303/36 | QH/H | | caGCAt/cat |
| Transcript | synonymous_ | 401/3926 | 318/1977 | 106/658 | A | | gcC/gcA |
| Transcript | inframe_delet | 520-522/1922 | 453-455/1407 | 151-152/468 | QQ/Q | | caGCAa/caa |
| Transcript | missense_vari | 340/5535 | 226/1794 | 76/597 | E/K | | Gaa/Aaa |
| Transcript | inframe_delet | 2990-2992/54 | 2916-2918/31 | 972-973/1044 | RQ/R | | cgGCAg/cgg |
| Transcript | missense_vari | 2749/3657 | 2525/3003 | 842/1000 | D/G | | gAc/gGc |
| Transcript | inframe_delet | 2828-2830/95 | 2637-2639/36 | 879-880/1231 | AQ/A | | gcGCAg/gcg |
| Transcript | synonymous_ | 1356/2347 | 1323/1434 | 441/477 | A | | gcG/gcA |
| Transcript | inframe_inser | 1355-1356/23 | 1322-1323/14 | 441/477 | A/AA | | gcg/gcAGCg |
| Transcript | missense_vari | 2995/5431 | 2433/3972 | 811/1323 | D/E | | gaT/gaG |
| Transcript | missense_vari | 14/11124 | 7/4326 | 3/1441 | A/T | | Gcg/Acg |
| Transcript | missense_vari | 7062/8455 | 6868/7932 | 2290/2643 | V/L | | Gtt/Ctt |
| Transcript | missense_vari | 73/3152 | 73/1785 | 25/594 | G/S | | Ggc/Agc |
| Transcript | synonymous_ | 945/6057 | 228/4113 | 76/1370 | G | | ggA/ggG |
| Transcript | inframe_delet | 283-285/5437 | 268-270/1035 | 90/344 | G/- | | GGC/- |
| Transcript | missense_vari | 914/3564 | 360/2793 | 120/930 | E/D | | gaG/gaT |
| Transcript | missense_vari | 1449/8496 | 1146/3399 | 382/1132 | E/D | | gaA/gaT |
| Transcript | frameshift_va | 1084-1085/16 | 857-858/879 | 286/292 | W/CLX | | tgg/tgTCTGg |
| Transcript | stop_gained | 3893/11917 | 3718/8526 | 1240/2841 | E/* | | Gag/Tag |
| Transcript | missense_vari | 844/1694 | 730/1137 | 244/378 | T/P | | Acc/Ccc |
| Transcript | inframe_delet | 8568-8603/19 | 8340-8375/16 | 2780-2792/55 | QQQQQQQQ | | caACAACAGC |
| Transcript | synonymous_ | 2145/11325 | 2124/6735 | 708/2244 | P | | ccG/ccT |
| Transcript | inframe_delet | 9906-9908/11 | 9906-9908/10 | 3302-3303/36 | QH/H | | caGCAt/cat |
| Transcript | inframe_delet | 520-522/1922 | 453-455/1407 | 151-152/468 | QQ/Q | | caGCAa/caa |
| Transcript | inframe_delet | 1041-1043/81 | 1041-1043/68 | 347-348/2283 | AA/A | | gcGGCa/gca |
| Transcript | missense_vari | 340/5535 | 226/1794 | 76/597 | E/K | | Gaa/Aaa |
| Transcript | inframe_delet | 2828-2830/95 | 2637-2639/36 | 879-880/1231 | AQ/A | | gcGCAg/gcg |
| Transcript | missense_vari | 4245/10944 | 3755/5073 | 1252/1690 | R/H | | cGt/cAt |
| Transcript | synonymous_ | 1347/2218 | 1269/1395 | 423/464 | S | | agC/agT |
| Transcript | synonymous_ | 1356/2347 | 1323/1434 | 441/477 | A | | gcG/gcA |
| Transcript | inframe_inser | 1355-1356/23 | 1322-1323/14 | 441/477 | A/AA | | gcg/gcAGCg |

| | | | | | | |
|---|---|---|---|---|---|---|
| Transcript | synonymous_ | 3499/10654 | 3111/9816 | 1037/3271 | R | cgT/cgC |
| Transcript | missense_vari | 2995/5431 | 2433/3972 | 811/1323 | D/E | gaT/gaG |
| Transcript | missense_vari | 9653/16461 | 9631/11892 | 3211/3963 | L/I | Ctc/Atc |
| Transcript | inframe_delet | 283-285/5437 | 268-270/1035 | 90/344 | G/- | GGC/- |
| Transcript | inframe_delet | 2738-2740/41 | 2103-2105/29 | 701-702/969 | GA/G | ggAGCa/gga |
| Transcript | missense_vari | 914/3564 | 360/2793 | 120/930 | E/D | gaG/gaT |

| Existing_variat | ALLELE_NUM | DISTANCE | STRAND_VEP | SYMBOL | SYMBOL_SOU | BIOTYPE |
|---|---|---|---|---|---|---|
| | 1 | | -1 | Etv4 | MGI | protein_codin |
| | 1 | | -1 | Npm1 | MGI | protein_codin |
| | 1 | | 1 | Myo18a | MGI | protein_codin |
| | 1 | | -1 | Smyd3 | MGI | protein_codin |
| | 1 | | -1 | Sptbn5 | MGI | protein_codin |
| | 1 | | -1 | Samhd1 | MGI | protein_codin |
| | 1 | | -1 | Ivl | MGI | protein_codin |
| | 1 | | -1 | Ptpn11 | MGI | protein_codin |
| | 1 | | 1 | Gigyf1 | MGI | protein_codin |
| | 1 | | -1 | Flt3 | MGI | protein_codin |
| | 1 | | 1 | Pik3c2g | MGI | protein_codin |
| | 1 | | -1 | Maz | MGI | protein_codin |
| | 1 | | -1 | Maz | MGI | protein_codin |
| | 1 | | 1 | Setd1a | MGI | protein_codin |
| | 1 | | 1 | Scgb1b30 | MGI | protein_codin |
| rs230103531 | 1 | | 1 | Rsf1 | MGI | protein_codin |
| | 1 | | -1 | Ankrd11 | MGI | protein_codin |
| | 1 | | -1 | Cbl | MGI | protein_codin |
| | 1 | | 1 | Rbm10 | MGI | protein_codin |
| | 1 | | 1 | Egr2 | MGI | protein_codin |
| | 1 | | -1 | Npm1 | MGI | protein_codin |
| | 1 | | 1 | Nf1 | MGI | protein_codin |
| | 1 | | 1 | Nf1 | MGI | protein_codin |
| | 1 | | 1 | Vgll3 | MGI | protein_codin |
| | 1 | | -1 | Klhl14 | MGI | protein_codin |
| | 1 | | -1 | Nt5c2 | MGI | protein_codin |
| | 1 | | -1 | Arid1a | MGI | protein_codin |
| | 1 | | -1 | Prdm13 | MGI | protein_codin |
| | 1 | | -1 | Ptpn11 | MGI | protein_codin |
| | 1 | | -1 | Flt3 | MGI | protein_codin |
| | 1 | | -1 | Maz | MGI | protein_codin |
| | 1 | | -1 | Maz | MGI | protein_codin |
| | 1 | | -1 | Npm1 | MGI | protein_codin |
| | 1 | | 1 | Nf1 | MGI | protein_codin |
| | 1 | | -1 | Rb1 | MGI | protein_codin |
| | 1 | | -1 | Serpinb3b | MGI | protein_codin |
| | 1 | | -1 | Sptbn5 | MGI | protein_codin |
| | 1 | | -1 | Ivl | MGI | protein_codin |
| | 1 | | -1 | Ptpn11 | MGI | protein_codin |
| | 1 | | 1 | Gigyf1 | MGI | protein_codin |
| | 1 | | 1 | D6Ertd527e | MGI | protein_codin |
| rs239855953 | 1 | | 1 | Plk1 | MGI | protein_codin |
| | 1 | | -1 | Maz | MGI | protein_codin |
| | 1 | | -1 | Maz | MGI | protein_codin |
| | 1 | | 1 | Rps19 | MGI | protein_codin |
| | 1 | | -1 | Cdh16 | MGI | protein_codin |
| | 1 | | -1 | Ammecr1 | MGI | protein_codin |
| | 1 | | -1 | Npm1 | MGI | protein_codin |
| | 1 | | -1 | Zswim6 | MGI | protein_codin |

| | | | | | |
|---|---|---|---|---|---|
| | 1 | 1 | Hcn1 | MGI | protein_codin |
| | 1 | -1 | Ranbp9 | MGI | protein_codin |
| | 1 | 1 | Fam170b | MGI | protein_codin |
| | 1 | -1 | Dach1 | MGI | protein_codin |
| | 1 | -1 | Kmt2d | MGI | protein_codin |
| | 1 | 1 | Vgll3 | MGI | protein_codin |
| | 1 | -1 | Sptbn5 | MGI | protein_codin |
| rs582628252 | 1 | -1 | Maml3 | MGI | protein_codin |
| | 1 | -1 | Ivl | MGI | protein_codin |
| | 1 | -1 | Ptpn11 | MGI | protein_codin |
| | 1 | 1 | Rbm33 | MGI | protein_codin |
| rs217659485 | 1 | -1 | Cckar | MGI | protein_codin |
| | 1 | -1 | Maz | MGI | protein_codin |
| | 1 | -1 | Maz | MGI | protein_codin |
| | 1 | 1 | Srcap | MGI | protein_codin |
| | 1 | -1 | Cdh16 | MGI | protein_codin |
| | 1 | 1 | Zfp703 | MGI | protein_codin |
| | 1 | -1 | Ammecr1 | MGI | protein_codin |
| | 1 | 1 | Rbm10 | MGI | protein_codin |
| | 1 | -1 | Sry | MGI | protein_codin |
| | 1 | -1 | Npm1 | MGI | protein_codin |
| | 1 | 1 | Nf1 | MGI | protein_codin |
| | 1 | -1 | Zswim6 | MGI | protein_codin |
| | 1 | -1 | Zfp503 | MGI | protein_codin |
| | 1 | 1 | Vgll3 | MGI | protein_codin |
| | 1 | -1 | Sptbn5 | MGI | protein_codin |
| | 1 | 1 | Dennd4b | MGI | protein_codin |
| | 1 | -1 | Ptpn11 | MGI | protein_codin |
| | 1 | 1 | Gigyf1 | MGI | protein_codin |
| | 1 | -1 | Flt3 | MGI | protein_codin |
| rs217659485 | 1 | -1 | Cckar | MGI | protein_codin |
| | 1 | -1 | Ammecr1 | MGI | protein_codin |
| rs578925286 | 1 | 1 | Mamld1 | MGI | protein_codin |
| | 1 | -1 | Amer1 | MGI | protein_codin |
| | 1 | -1 | Npm1 | MGI | protein_codin |
| | 1 | 1 | Adamts2 | MGI | protein_codin |
| | 1 | 1 | Nf1 | MGI | protein_codin |
| | 1 | 1 | Hcn1 | MGI | protein_codin |
| | 1 | -1 | Kmt2d | MGI | protein_codin |
| | 1 | -1 | Sptbn5 | MGI | protein_codin |
| | 1 | -1 | Ep400 | MGI | protein_codin |
| | 1 | -1 | Ptpn11 | MGI | protein_codin |
| | 1 | 1 | Rbm33 | MGI | protein_codin |
| | 1 | 1 | D6Ertd527e | MGI | protein_codin |
| | 1 | -1 | Maz | MGI | protein_codin |
| | 1 | -1 | Maz | MGI | protein_codin |
| rs230103531 | 1 | 1 | Rsf1 | MGI | protein_codin |
| | 1 | 1 | Rsf1 | MGI | protein_codin |
| | 1 | 1 | Zfp703 | MGI | protein_codin |
| | 1 | -1 | Gab1 | MGI | protein_codin |

| | | | | | |
|---|---|---|---|---|---|
| | 1 | 1 | Zfc3h1 | MGI | protein_codin |
| | 1 | -1 | Actg1 | MGI | protein_codin |
| | 1 | -1 | Npm1 | MGI | protein_codin |
| | 1 | 1 | Dsp | MGI | protein_codin |
| | 1 | 1 | Mdc1 | MGI | protein_codin |
| | 1 | 1 | Ankhd1 | MGI | protein_codin |
| rs39394853 | 1 | -1 | Elf3 | MGI | protein_codin |
| rs49378976 | 1 | -1 | Elf3 | MGI | protein_codin |
| | 1 | 1 | Wt1 | MGI | protein_codin |
| | 1 | -1 | Sptbn5 | MGI | protein_codin |
| | 1 | -1 | Gm10800 | MGI | protein_codin |
| | 1 | -1 | Arid1a | MGI | protein_codin |
| | 1 | -1 | Mllt3 | MGI | protein_codin |
| | 1 | -1 | Ptpn11 | MGI | protein_codin |
| | 1 | 1 | Hectd4 | MGI | protein_codin |
| | 1 | 1 | Setd1b | MGI | protein_codin |
| | 1 | 1 | Gigyf1 | MGI | protein_codin |
| | 1 | -1 | Flt3 | MGI | protein_codin |
| | 1 | 1 | Kit | MGI | protein_codin |
| | 1 | -1 | Maz | MGI | protein_codin |
| | 1 | -1 | Maz | MGI | protein_codin |
| | 1 | 1 | Setd1a | MGI | protein_codin |
| | 1 | 1 | Fat1 | MGI | protein_codin |
| | 1 | -1 | Ammecr1 | MGI | protein_codin |
| | 1 | -1 | Npm1 | MGI | protein_codin |
| | 1 | -1 | Rpa1 | MGI | protein_codin |
| | 1 | 1 | Vgll3 | MGI | protein_codin |
| | 1 | -1 | Vegfa | MGI | protein_codin |
| | 1 | 1 | Chka | MGI | protein_codin |
| | 1 | -1 | Sptbn5 | MGI | protein_codin |
| | 1 | -1 | Arid1a | MGI | protein_codin |
| | 1 | -1 | Mllt3 | MGI | protein_codin |
| | 1 | -1 | Ptpn11 | MGI | protein_codin |
| | 1 | -1 | Maz | MGI | protein_codin |
| | 1 | -1 | Maz | MGI | protein_codin |
| | 1 | 1 | Srcap | MGI | protein_codin |
| | 1 | 1 | Zfp703 | MGI | protein_codin |
| | 1 | 1 | Fyn | MGI | protein_codin |
| | 1 | -1 | Npm1 | MGI | protein_codin |
| | 1 | 1 | Vgll3 | MGI | protein_codin |
| | 1 | -1 | Erg | MGI | protein_codin |
| | 1 | -1 | Sos1 | MGI | protein_codin |
| | 1 | 1 | Olfr1463 | MGI | protein_codin |
| | 1 | 1 | Olfr1463 | MGI | protein_codin |
| rs39394853 | 1 | -1 | Elf3 | MGI | protein_codin |
| rs49378976 | 1 | -1 | Elf3 | MGI | protein_codin |
| | 1 | -1 | Sptbn5 | MGI | protein_codin |
| | 1 | 1 | Prrc2b | MGI | protein_codin |
| | 1 | 1 | Notch2 | MGI | protein_codin |
| | 1 | -1 | Ptpn11 | MGI | protein_codin |

| | | | | | |
|---|---|---|---|---|---|
| | 1 | 1 | Gigyf1 | MGI | protein_codin |
| | 1 | -1 | Flt3 | MGI | protein_codin |
| | 1 | -1 | Maz | MGI | protein_codin |
| | 1 | -1 | Maz | MGI | protein_codin |
| | 1 | -1 | Grin2d | MGI | protein_codin |
| | 1 | -1 | Ntrk3 | MGI | protein_codin |
| | 1 | 1 | Zfp703 | MGI | protein_codin |
| | 1 | -1 | Kmt2a | MGI | protein_codin |
| | 1 | -1 | Ammecr1 | MGI | protein_codin |
| | 1 | 1 | Rbm10 | MGI | protein_codin |
| | 1 | -1 | Amer1 | MGI | protein_codin |
| | 1 | -1 | Ros1 | MGI | protein_codin |
| | 1 | -1 | Npm1 | MGI | protein_codin |
| | 1 | 1 | Rnf43 | MGI | protein_codin |
| | 1 | 1 | Phlpp1 | MGI | protein_codin |
| rs39394853 | 1 | -1 | Elf3 | MGI | protein_codin |
| rs49378976 | 1 | -1 | Elf3 | MGI | protein_codin |
| | 1 | -1 | Sptbn5 | MGI | protein_codin |
| | 1 | -1 | Samhd1 | MGI | protein_codin |
| | 1 | 1 | Mapkap1 | MGI | protein_codin |
| | 1 | -1 | Arid1a | MGI | protein_codin |
| | 1 | -1 | Ptpn11 | MGI | protein_codin |
| | 1 | 1 | Gigyf1 | MGI | protein_codin |
| | 1 | -1 | Flt3 | MGI | protein_codin |
| | 1 | 1 | Rbm33 | MGI | protein_codin |
| | 1 | -1 | Maz | MGI | protein_codin |
| | 1 | -1 | Maz | MGI | protein_codin |
| | 1 | -1 | Cdh16 | MGI | protein_codin |
| | 1 | -1 | Ankrd11 | MGI | protein_codin |
| | 1 | -1 | Ammecr1 | MGI | protein_codin |
| | 1 | 1 | Alg13 | MGI | protein_codin |
| | 1 | -1 | Npm1 | MGI | protein_codin |
| rs1135367000 | 1 | -1 | Tcf20 | MGI | protein_codin |
| | 1 | 1 | Vgll3 | MGI | protein_codin |
| | 1 | 1 | Ccnd3 | MGI | protein_codin |
| | 1 | 1 | Olfr1463 | MGI | protein_codin |
| | 1 | 1 | Olfr1463 | MGI | protein_codin |
| | 1 | 1 | Chka | MGI | protein_codin |
| rs39394853 | 1 | -1 | Elf3 | MGI | protein_codin |
| rs49378976 | 1 | -1 | Elf3 | MGI | protein_codin |
| | 1 | -1 | Sptbn5 | MGI | protein_codin |
| | 1 | -1 | Ptpn11 | MGI | protein_codin |
| | 1 | 1 | Gigyf1 | MGI | protein_codin |
| | 1 | -1 | Flt3 | MGI | protein_codin |
| | 1 | -1 | Maz | MGI | protein_codin |
| | 1 | -1 | Maz | MGI | protein_codin |
| | 1 | 1 | Zfhx3 | MGI | protein_codin |
| | 1 | -1 | Tgfbr2 | MGI | protein_codin |
| | 1 | 1 | Alg13 | MGI | protein_codin |
| | 1 | -1 | Npm1 | MGI | protein_codin |

| | | | | | |
|---|---|---|---|---|---|
| | 1 | 1 | Arid4b | MGI | protein_codin |
| | 1 | -1 | Zfp503 | MGI | protein_codin |
| | 1 | -1 | Rb1 | MGI | protein_codin |
| | 1 | 1 | Ankhd1 | MGI | protein_codin |
| rs39394853 | 1 | -1 | Elf3 | MGI | protein_codin |
| rs49378976 | 1 | -1 | Elf3 | MGI | protein_codin |
| | 1 | 1 | Itpkb | MGI | protein_codin |
| | 1 | -1 | Sptbn5 | MGI | protein_codin |
| | 1 | -1 | Ivl | MGI | protein_codin |
| | 1 | -1 | Ptpn11 | MGI | protein_codin |
| | 1 | 1 | Hectd4 | MGI | protein_codin |
| | 1 | 1 | Gigyf1 | MGI | protein_codin |
| rs217659485 | 1 | -1 | Cckar | MGI | protein_codin |
| | 1 | -1 | Maz | MGI | protein_codin |
| | 1 | -1 | Maz | MGI | protein_codin |
| | 1 | 1 | Rsf1 | MGI | protein_codin |
| | 1 | -1 | Cdh16 | MGI | protein_codin |
| | 1 | 1 | Smarca4 | MGI | protein_codin |
| | 1 | -1 | Ammecr1 | MGI | protein_codin |
| | 1 | -1 | Sry | MGI | protein_codin |
| | 1 | -1 | Sry | MGI | protein_codin |
| | 1 | 1 | Zfc3h1 | MGI | protein_codin |
| | 1 | 1 | Lats1 | MGI | protein_codin |
| | 1 | -1 | Npm1 | MGI | protein_codin |
| | 1 | 1 | Hcn1 | MGI | protein_codin |
| | 1 | -1 | Rb1 | MGI | protein_codin |
| | 1 | -1 | Kmt2d | MGI | protein_codin |
| | 1 | 1 | Vgll3 | MGI | protein_codin |
| | 1 | 1 | Zfp318 | MGI | protein_codin |
| | 1 | -1 | Serpinb3b | MGI | protein_codin |
| rs39394853 | 1 | -1 | Elf3 | MGI | protein_codin |
| rs49378976 | 1 | -1 | Elf3 | MGI | protein_codin |
| | 1 | -1 | Sptbn5 | MGI | protein_codin |
| | 1 | -1 | Lrp1b | MGI | protein_codin |
| rs582628252 | 1 | -1 | Maml3 | MGI | protein_codin |
| | 1 | 1 | Zfhx4 | MGI | protein_codin |
| | 1 | -1 | Ptpn11 | MGI | protein_codin |
| | 1 | 1 | Gigyf1 | MGI | protein_codin |
| | 1 | 1 | Kit | MGI | protein_codin |
| | 1 | -1 | Kras | MGI | protein_codin |
| rs239855953 | 1 | 1 | Plk1 | MGI | protein_codin |
| | 1 | -1 | Maz | MGI | protein_codin |
| | 1 | -1 | Maz | MGI | protein_codin |
| | 1 | 1 | Rsf1 | MGI | protein_codin |
| | 1 | 1 | Mst1r | MGI | protein_codin |
| | 1 | -1 | Npm1 | MGI | protein_codin |
| | 1 | -1 | Lats2 | MGI | protein_codin |
| | 1 | 1 | Vgll3 | MGI | protein_codin |
| rs39394853 | 1 | -1 | Elf3 | MGI | protein_codin |
| rs49378976 | 1 | -1 | Elf3 | MGI | protein_codin |

| | | | | | |
|---|---|---|---|---|---|
| | 1 | -1 | Sptbn5 | MGI | protein_codin |
| | 1 | -1 | Samhd1 | MGI | protein_codin |
| | 1 | -1 | Wdr63 | MGI | protein_codin |
| | 1 | -1 | Ptpn11 | MGI | protein_codin |
| | 1 | 1 | Gigyf1 | MGI | protein_codin |
| | 1 | -1 | Flt3 | MGI | protein_codin |
| | 1 | -1 | Maz | MGI | protein_codin |
| | 1 | -1 | Maz | MGI | protein_codin |
| | 1 | 1 | Srcap | MGI | protein_codin |
| | 1 | -1 | Ntrk3 | MGI | protein_codin |
| | 1 | -1 | Ankrd11 | MGI | protein_codin |
| | 1 | -1 | Acta1 | MGI | protein_codin |
| | 1 | 1 | Zfp703 | MGI | protein_codin |
| | 1 | -1 | Zrsr2 | MGI | protein_codin |
| | 1 | -1 | Npm1 | MGI | protein_codin |
| | 1 | -1 | Chd3 | MGI | protein_codin |
| | 1 | 1 | Ep300 | MGI | protein_codin |
| | 1 | -1 | Kmt2d | MGI | protein_codin |
| | 1 | 1 | Robo1 | MGI | protein_codin |
| | 1 | -1 | Sptbn5 | MGI | protein_codin |
| | 1 | -1 | Samhd1 | MGI | protein_codin |
| | 1 | -1 | Ivl | MGI | protein_codin |
| | 1 | -1 | Ptpn11 | MGI | protein_codin |
| | 1 | 1 | Gigyf1 | MGI | protein_codin |
| | 1 | -1 | Flt3 | MGI | protein_codin |
| | 1 | 1 | Rbm33 | MGI | protein_codin |
| | 1 | -1 | Maz | MGI | protein_codin |
| | 1 | -1 | Maz | MGI | protein_codin |
| | 1 | -1 | Grin2d | MGI | protein_codin |
| rs230103531 | 1 | 1 | Rsf1 | MGI | protein_codin |
| | 1 | -1 | Ankrd11 | MGI | protein_codin |
| | 1 | 1 | Zfp703 | MGI | protein_codin |
| | 1 | -1 | Zmym3 | MGI | protein_codin |
| | 1 | -1 | Ammecr1 | MGI | protein_codin |
| | 1 | 1 | Rbm10 | MGI | protein_codin |
| | 1 | -1 | Amer1 | MGI | protein_codin |
| | 1 | -1 | Npm1 | MGI | protein_codin |
| | 1 | 1 | Nf1 | MGI | protein_codin |
| | 1 | 1 | Fam170b | MGI | protein_codin |
| | 1 | -1 | Kmt2d | MGI | protein_codin |
| | 1 | 1 | Arid1b | MGI | protein_codin |
| | 1 | -1 | Sptbn5 | MGI | protein_codin |
| | 1 | -1 | Ivl | MGI | protein_codin |
| | 1 | -1 | Arid1a | MGI | protein_codin |
| | 1 | -1 | Ptpn11 | MGI | protein_codin |
| | 1 | 1 | Rbm33 | MGI | protein_codin |
| | 1 | 1 | Kdm5a | MGI | protein_codin |
| | 1 | 1 | D6Ertd527e | MGI | protein_codin |
| | 1 | -1 | Maz | MGI | protein_codin |
| | 1 | -1 | Maz | MGI | protein_codin |

| | | | | |
|---|---|---|---|---|
| 1 | 1 | Srcap | MGI | protein_codin |
| 1 | -1 | Grin2d | MGI | protein_codin |
| 1 | -1 | Kmt2a | MGI | protein_codin |
| 1 | -1 | Ammecr1 | MGI | protein_codin |
| 1 | 1 | Alg13 | MGI | protein_codin |
| 1 | 1 | Rbm10 | MGI | protein_codin |

| CANONICAL | CCDS | ENSP | SWISSPROT | TREMBL | UNIPARC | RefSeq |
|---|---|---|---|---|---|---|
| YES | CCDS83903.1 | ENSMUSP0000( | | A2A5C3 | UPI00002146: | NM_00131636 |
| YES | CCDS24532.1 | ENSMUSP0000( | Q61937 | Q5SQB7 | UPI0000003F6 | NM_00125226 |
| YES | CCDS70253.1 | ENSMUSP0000( | | E9QAX2 | UPI0001F7894 | NM_00129121 |
| YES | CCDS15560.1 | ENSMUSP0000( | Q9CWR2 | | UPI00000250 | NM_027188.4 |
| YES | | ENSMUSP0000( | | A0A571BF02 | UPI00114BC67 | NM_00137093 |
| YES | CCDS16973.2 | ENSMUSP0000( | Q60710 | | UPI00015DF5I | NM_018851.4 |
| YES | CCDS17561.1 | ENSMUSP0000( | | G3X9D9 | UPI0000022D: | NM_008412.3 |
| YES | CCDS39247.1 | ENSMUSP0000( | P35235 | | UPI00000E5F1 | NM_011202.3 |
| YES | CCDS51673.1 | ENSMUSP0000( | Q99MR1 | | UPI00005643I | NM_031408.2 |
| YES | CCDS39400.1 | ENSMUSP0000( | | Q3UEW6 | UPI0000356EI | NM_010229.2 |
| YES | | ENSMUSP0000( | | E9QQ35 | UPI00004793: | NM_207683.3 |
| YES | CCDS40137.1 | ENSMUSP0000( | | F8VPK3 | UPI000017C8 | NM_010772.1 |
| YES | CCDS40137.1 | ENSMUSP0000( | | F8VPK3 | UPI000017C8 | NM_010772.1 |
| YES | CCDS40144.1 | ENSMUSP0000( | E9PYH6 | | UPI0000605A: | |
| YES | CCDS52201.1 | ENSMUSP0000( | | A2BH64 | UPI0000D77B | NM_00109933 |
| YES | CCDS40025.1 | ENSMUSP0000( | | E9PWW9 | UPI00006059 | NM_00108126 |
| YES | CCDS40507.2 | ENSMUSP0000( | E9Q4F7 | | UPI0000605E5 | NM_00108137 |
| YES | CCDS40598.1 | ENSMUSP0000( | P22682 | | UPI00003580 | NM_007619.2 |
| YES | CCDS40885.1 | ENSMUSP0000( | Q99KG3 | | UPI0000027AI | NM_145627.3 |
| YES | CCDS35927.2 | ENSMUSP0000( | P08152 | Q3U207 | UPI00000018( | NM_010118.3 |
| YES | CCDS24532.1 | ENSMUSP0000( | Q61937 | Q5SQB7 | UPI0000003F6 | NM_00125226 |
| YES | CCDS25119.1 | ENSMUSP0000( | Q04690 | | UPI00000271( | NM_010897.2 |
| YES | CCDS25119.1 | ENSMUSP0000( | Q04690 | | UPI00000271( | NM_010897.2 |
| YES | CCDS49884.1 | ENSMUSP0000( | | E9Q1Y1 | UPI00002159I | NM_00136876 |
| YES | CCDS37748.1 | ENSMUSP0000( | Q69ZK5 | | UPI000058AC | NM_00108140 |
| YES | CCDS50462.1 | ENSMUSP0000( | | E9Q9M1 | UPI0001B256( | NM_00116436 |
| YES | CCDS38908.1 | ENSMUSP0000( | A2BH40 | | UPI000059E1 | NM_00108081 |
| YES | CCDS38699.1 | ENSMUSP0000( | E9PZZ1 | | UPI00001E35I | NM_00108077 |
| YES | CCDS39247.1 | ENSMUSP0000( | P35235 | | UPI00000E5F1 | NM_011202.3 |
| YES | CCDS39400.1 | ENSMUSP0000( | | Q3UEW6 | UPI0000356EI | NM_010229.2 |
| YES | CCDS40137.1 | ENSMUSP0000( | | F8VPK3 | UPI000017C8 | NM_010772.1 |
| YES | CCDS40137.1 | ENSMUSP0000( | | F8VPK3 | UPI000017C8 | NM_010772.1 |
| YES | CCDS24532.1 | ENSMUSP0000( | Q61937 | Q5SQB7 | UPI0000003F6 | NM_00125226 |
| YES | CCDS25119.1 | ENSMUSP0000( | Q04690 | | UPI00000271( | NM_010897.2 |
| YES | CCDS27267.1 | ENSMUSP0000( | P13405 | | UPI0000157BI | NM_009029.3 |
| YES | CCDS15216.1 | ENSMUSP0000( | | Q9D1Q5 | UPI00000220I | NM_198680.2 |
| YES | | ENSMUSP0000( | | A0A571BF02 | UPI00114BC67 | NM_00137093 |
| YES | CCDS17561.1 | ENSMUSP0000( | | G3X9D9 | UPI0000022D: | NM_008412.3 |
| YES | CCDS39247.1 | ENSMUSP0000( | P35235 | | UPI00000E5F1 | NM_011202.3 |
| YES | CCDS51673.1 | ENSMUSP0000( | Q99MR1 | | UPI00005643I | NM_031408.2 |
| YES | CCDS51837.1 | ENSMUSP0000( | | A0A0N4SWI3 | UPI0001BE80 | NM_00116793 |
| YES | CCDS21812.1 | ENSMUSP0000( | Q07832 | Q3TPZ2 | UPI00000018 | NM_011121.4 |
| YES | CCDS40137.1 | ENSMUSP0000( | | F8VPK3 | UPI000017C8 | NM_010772.1 |
| YES | CCDS40137.1 | ENSMUSP0000( | | F8VPK3 | UPI000017C8 | NM_010772.1 |
| YES | CCDS20966.1 | ENSMUSP0000( | Q9CZX8 | | UPI00000081 | NM_00136011 |
| YES | CCDS22583.1 | ENSMUSP0000( | O88338 | Q546A8 | UPI00000238I | NM_007663.3 |
| YES | CCDS30452.1 | ENSMUSP0000( | Q9JHT5 | | UPI00000040( | NM_019496.4 |
| YES | CCDS24532.1 | ENSMUSP0000( | Q61937 | Q5SQB7 | UPI0000003F6 | NM_00125226 |
| YES | | ENSMUSP0000( | Q80TB7 | | UPI0001837E | NM_145456.3 |

| | | | | | | |
|---|---|---|---|---|---|---|
| YES | CCDS26793.1 | ENSMUSP0000 | O88704 | | UPI0000020F | NM_010408.3 |
| YES | CCDS49252.1 | ENSMUSP0000 | | E9Q5D6 | UPI00016024 | NM_019930.2 |
| YES | CCDS49439.1 | ENSMUSP0000 | E9PXT9 | | UPI0001B25B | NM_0011644 |
| YES | CCDS36994.1 | ENSMUSP0000 | Q9QYB2 | | UPI0000029D | NM_007826.3 |
| YES | CCDS49725.2 | ENSMUSP0000 | | A0A0A0MQ73 | UPI00023B36 | NM_0010332 |
| YES | CCDS49884.1 | ENSMUSP0000 | | E9Q1Y1 | UPI00002159 | NM_0013687 |
| YES | | ENSMUSP0000 | | A0A571BF02 | UPI00114BC6 | NM_0013709 |
| YES | CCDS17342.2 | ENSMUSP0000 | | D4QGC2 | UPI00016116 | NM_0010041 |
| YES | CCDS17561.1 | ENSMUSP0000 | | G3X9D9 | UPI0000022D | NM_008412.3 |
| YES | CCDS39247.1 | ENSMUSP0000 | P35235 | | UPI00000E5F1 | NM_011202.3 |
| YES | CCDS51448.1 | ENSMUSP0000 | Q9CXK9 | | UPI0000D661 | NM_028234.1 |
| YES | CCDS19293.1 | ENSMUSP0000 | O08786 | | UPI00000231 | NM_009827.2 |
| YES | CCDS40137.1 | ENSMUSP0000 | | F8VPK3 | UPI000017C8 | NM_010772.1 |
| YES | CCDS40137.1 | ENSMUSP0000 | | F8VPK3 | UPI000017C8 | NM_010772.1 |
| YES | CCDS80806.1 | ENSMUSP0000 | | A0A087WQ44 | UPI0003D70E | NM_0013042 |
| YES | CCDS22583.1 | ENSMUSP0000 | O88338 | Q546A8 | UPI00000238 | NM_007663.3 |
| YES | CCDS52532.1 | ENSMUSP0000 | P0CL69 | F6M3K1 | UPI000154B5 | NM_0011015 |
| YES | CCDS30452.1 | ENSMUSP0000 | Q9JHT5 | | UPI00000040 | NM_019496.4 |
| YES | CCDS40885.1 | ENSMUSP0000 | Q99KG3 | | UPI0000027A | NM_145627.3 |
| YES | CCDS30545.1 | ENSMUSP0000 | Q05738 | Q53WV7 | UPI00000005 | NM_011564.1 |
| YES | CCDS24532.1 | ENSMUSP0000 | Q61937 | Q5SQB7 | UPI0000003F | NM_0012522 |
| YES | CCDS25119.1 | ENSMUSP0000 | Q04690 | | UPI00000271 | NM_010897.2 |
| YES | | ENSMUSP0000 | Q80TB7 | | UPI0001837E | NM_145456.3 |
| YES | CCDS36822.1 | ENSMUSP0000 | Q7TMA2 | | UPI00001B06 | NM_145459.3 |
| YES | CCDS49884.1 | ENSMUSP0000 | | E9Q1Y1 | UPI00002159 | NM_0013687 |
| YES | | ENSMUSP0000 | | A0A571BF02 | UPI00114BC6 | NM_0013709 |
| YES | CCDS38502.2 | ENSMUSP0000 | | A0A0R4J172 | UPI00005ACF | NM_201407.4 |
| YES | CCDS39247.1 | ENSMUSP0000 | P35235 | | UPI00000E5F1 | NM_011202.3 |
| YES | CCDS51673.1 | ENSMUSP0000 | Q99MR1 | | UPI00005643 | NM_031408.2 |
| YES | CCDS39400.1 | ENSMUSP0000 | | Q3UEW6 | UPI0000356E | NM_010229.2 |
| YES | CCDS19293.1 | ENSMUSP0000 | O08786 | | UPI00000231 | NM_009827.2 |
| YES | CCDS30452.1 | ENSMUSP0000 | Q9JHT5 | | UPI00000040 | NM_019496.4 |
| YES | CCDS57759.1 | ENSMUSP0000 | P0C6A2 | | UPI00001F84 | NM_0010813 |
| YES | CCDS41067.1 | ENSMUSP0000 | Q7TS75 | | UPI0000EDA2 | NM_175179.4 |
| YES | CCDS24532.1 | ENSMUSP0000 | Q61937 | Q5SQB7 | UPI0000003F | NM_0012522 |
| YES | CCDS24636.1 | ENSMUSP0000 | Q8C9W3 | | UPI000000B9 | NM_175643.3 |
| YES | CCDS25119.1 | ENSMUSP0000 | Q04690 | | UPI00000271 | NM_010897.2 |
| YES | CCDS26793.1 | ENSMUSP0000 | O88704 | | UPI0000020F | NM_010408.3 |
| YES | CCDS49725.2 | ENSMUSP0000 | | A0A0A0MQ73 | UPI00023B36 | NM_0010332 |
| YES | | ENSMUSP0000 | | A0A571BF02 | UPI00114BC6 | NM_0013709 |
| YES | CCDS19529.1 | ENSMUSP0000 | Q8CHI8 | | UPI00001939 | NM_029337.2 |
| YES | CCDS39247.1 | ENSMUSP0000 | P35235 | | UPI00000E5F1 | NM_011202.3 |
| YES | CCDS51448.1 | ENSMUSP0000 | Q9CXK9 | | UPI0000D661 | NM_028234.1 |
| YES | CCDS51837.1 | ENSMUSP0000 | | A0A0N4SWI3 | UPI0001BE80 | NM_0011679 |
| YES | CCDS40137.1 | ENSMUSP0000 | | F8VPK3 | UPI000017C8 | NM_010772.1 |
| YES | CCDS40137.1 | ENSMUSP0000 | | F8VPK3 | UPI000017C8 | NM_010772.1 |
| YES | CCDS40025.1 | ENSMUSP0000 | | E9PWW9 | UPI00006059 | NM_0010812 |
| YES | CCDS40025.1 | ENSMUSP0000 | | E9PWW9 | UPI00006059 | NM_0010812 |
| YES | CCDS52532.1 | ENSMUSP0000 | P0CL69 | F6M3K1 | UPI000154B5 | NM_0011015 |
| YES | CCDS85569.1 | ENSMUSP0000 | | A0A1B0GS41 | UPI0003D704 | NM_0013012 |

| | | | | | | |
|---|---|---|---|---|---|---|
| YES | CCDS36062.1 | ENSMUSP0000( | | B2RT41 | UPI00006091⁄ | NM_00103326 |
| YES | CCDS25730.1 | ENSMUSP0000( | P63260 | Q4KL81 | UPI0000000C3 | NM_009609.3 |
| YES | CCDS24532.1 | ENSMUSP0000( | Q61937 | Q5SQB7 | UPI0000003F6 | NM_00125226 |
| YES | CCDS49239.1 | ENSMUSP0000( | E9Q557 | | UPI000042935 | NM_023842.2 |
| YES | CCDS37604.1 | ENSMUSP0000( | | E9QK89 | UPI0000D67D | NM_00101083 |
| YES | CCDS50256.1 | ENSMUSP0000( | | E9PUR0 | UPI00015E16A | NM_175375.3 |
| YES | CCDS48372.1 | ENSMUSP0000( | Q3UPW2 | | UPI00005AC1I | NM_00116313 |
| YES | CCDS48372.1 | ENSMUSP0000( | Q3UPW2 | | UPI00005AC1I | NM_00116313 |
| YES | | ENSMUSP0000( | | F6Y2T6 | UPI0001889B3 | NM_144783.2 |
| YES | | ENSMUSP0000( | | A0A571BF02 | UPI00114BC67 | NM_00137093 |
| YES | | ENSMUSP0000( | | D3Z496 | UPI0000D8B83 | |
| YES | CCDS38908.1 | ENSMUSP0000( | A2BH40 | | UPI000059E15 | NM_00108081 |
| YES | CCDS38796.1 | ENSMUSP0000( | A2AM29 | | UPI00001F0FA | NM_027326.3 |
| YES | CCDS39247.1 | ENSMUSP0000( | P35235 | | UPI00000E5F1 | NM_011202.3 |
| YES | CCDS57377.1 | ENSMUSP0000( | | E9Q2E4 | UPI00023B36A | NM_181421.4 |
| YES | CCDS59684.1 | ENSMUSP0000( | Q8CFT2 | | UPI000163AC7 | NM_00104039 |
| YES | CCDS51673.1 | ENSMUSP0000( | Q99MR1 | | UPI00005643I | NM_031408.2 |
| YES | CCDS39400.1 | ENSMUSP0000( | | Q3UEW6 | UPI0000356EE | NM_010229.2 |
| YES | CCDS51525.1 | ENSMUSP0000( | P05532 | | UPI00000E988 | NM_00112273 |
| YES | CCDS40137.1 | ENSMUSP0000( | | F8VPK3 | UPI000017C88 | NM_010772.1 |
| YES | CCDS40137.1 | ENSMUSP0000( | | F8VPK3 | UPI000017C88 | NM_010772.1 |
| YES | CCDS40144.1 | ENSMUSP0000( | E9PYH6 | | UPI0000605A7 | |
| YES | CCDS52547.1 | ENSMUSP0000( | | F2Z4A3 | UPI000050B5( | |
| YES | CCDS30452.1 | ENSMUSP0000( | Q9JHT5 | | UPI00000040( | NM_019496.4 |
| YES | CCDS24532.1 | ENSMUSP0000( | Q61937 | Q5SQB7 | UPI0000003F6 | NM_00125226 |
| YES | CCDS48847.1 | ENSMUSP0000( | | Q5SWN2 | UPI0000D8B6( | NM_00116422 |
| YES | CCDS49884.1 | ENSMUSP0000( | | E9Q1Y1 | UPI00002159I | NM_00136876 |
| YES | | ENSMUSP0000( | | F8WH81 | UPI0001F793( | NM_00102525 |
| YES | CCDS70913.1 | ENSMUSP0000( | O54804 | | UPI00001ECAI | NM_00127149 |
| YES | | ENSMUSP0000( | | A0A571BF02 | UPI00114BC67 | NM_00137093 |
| YES | CCDS38908.1 | ENSMUSP0000( | A2BH40 | | UPI000059E15 | NM_00108081 |
| YES | CCDS38796.1 | ENSMUSP0000( | A2AM29 | | UPI00001F0FA | NM_027326.3 |
| YES | CCDS39247.1 | ENSMUSP0000( | P35235 | | UPI00000E5F1 | NM_011202.3 |
| YES | CCDS40137.1 | ENSMUSP0000( | | F8VPK3 | UPI000017C88 | NM_010772.1 |
| YES | CCDS40137.1 | ENSMUSP0000( | | F8VPK3 | UPI000017C88 | NM_010772.1 |
| YES | CCDS80806.1 | ENSMUSP0000( | | A0A087WQ44 | UPI0003D70E9 | NM_00130426 |
| YES | CCDS52532.1 | ENSMUSP0000( | P0CL69 | F6M3K1 | UPI000154B57 | NM_00110150 |
| YES | CCDS48538.1 | ENSMUSP0000( | P39688 | | UPI000019652 | NM_00112289 |
| YES | CCDS24532.1 | ENSMUSP0000( | Q61937 | Q5SQB7 | UPI0000003F6 | NM_00125226 |
| YES | CCDS49884.1 | ENSMUSP0000( | | E9Q1Y1 | UPI00002159I | NM_00136876 |
| YES | CCDS37411.1 | ENSMUSP0000( | P81270 | | UPI000002855 | NM_133659.3 |
| YES | CCDS37704.1 | ENSMUSP0000( | Q62245 | | UPI0000028B8 | NM_009231.2 |
| YES | CCDS29654.1 | ENSMUSP0000( | | Q7TQR3 | UPI00001AA6I | |
| YES | CCDS29654.1 | ENSMUSP0000( | | Q7TQR3 | UPI00001AA6I | |
| YES | CCDS48372.1 | ENSMUSP0000( | Q3UPW2 | | UPI00005AC1I | NM_00116313 |
| YES | CCDS48372.1 | ENSMUSP0000( | Q3UPW2 | | UPI00005AC1I | NM_00116313 |
| YES | | ENSMUSP0000( | | A0A571BF02 | UPI00114BC67 | NM_00137093 |
| YES | CCDS50565.1 | ENSMUSP0000( | | F8WHT3 | UPI0000F4D4( | NM_00115963 |
| YES | CCDS51013.1 | ENSMUSP0000( | O35516 | | UPI0000022D7 | NM_010928.2 |
| YES | CCDS39247.1 | ENSMUSP0000( | P35235 | | UPI00000E5F1 | NM_011202.3 |

| | | | | | | |
|---|---|---|---|---|---|---|
| YES | CCDS51673.1 | ENSMUSP0000 | Q99MR1 | | UPI00005643 | NM_031408.2 |
| YES | CCDS39400.1 | ENSMUSP0000 | | Q3UEW6 | UPI0000356E | NM_010229.2 |
| YES | CCDS40137.1 | ENSMUSP0000 | | F8VPK3 | UPI000017C8 | NM_010772.1 |
| YES | CCDS40137.1 | ENSMUSP0000 | | F8VPK3 | UPI000017C8 | NM_010772.1 |
| YES | CCDS21267.1 | ENSMUSP0000 | Q03391 | | UPI0000D629 | NM_008172.2 |
| YES | CCDS21371.1 | ENSMUSP0000 | Q6VNS1 | | UPI000002B6 | NM_008746.5 |
| YES | CCDS52532.1 | ENSMUSP0000 | P0CL69 | F6M3K1 | UPI000154B5 | NM_0011015 |
| YES | CCDS40603.1 | ENSMUSP0000 | P55200 | | UPI00006061 | |
| YES | CCDS30452.1 | ENSMUSP0000 | Q9JHT5 | | UPI00000040 | NM_019496.4 |
| YES | CCDS40885.1 | ENSMUSP0000 | Q99KG3 | | UPI0000027A | NM_145627.3 |
| YES | CCDS41067.1 | ENSMUSP0000 | Q7TS75 | | UPI0000EDA2 | NM_175179.4 |
| YES | CCDS23838.1 | ENSMUSP0000 | Q78DX7 | | UPI00000217 | NM_011282.2 |
| YES | CCDS24532.1 | ENSMUSP0000 | Q61937 | Q5SQB7 | UPI0000003F | NM_0012522 |
| YES | CCDS25215.2 | ENSMUSP0000 | Q5NCP0 | | UPI000049C6 | NM_172448.4 |
| YES | CCDS48337.1 | ENSMUSP0000 | Q8CHE4 | | UPI000024ED | NM_133821.3 |
| YES | CCDS48372.1 | ENSMUSP0000 | Q3UPW2 | | UPI00005AC1 | NM_0011631 |
| YES | CCDS48372.1 | ENSMUSP0000 | Q3UPW2 | | UPI00005AC1 | NM_0011631 |
| YES | | ENSMUSP0000 | | A0A571BF02 | UPI00114BC6 | NM_0013709 |
| YES | CCDS16973.2 | ENSMUSP0000 | Q60710 | | UPI00015DF5 | NM_018851.4 |
| YES | CCDS15948.1 | ENSMUSP0000 | Q8BKH7 | | UPI00000EB5 | NM_0013629 |
| YES | CCDS38908.1 | ENSMUSP0000 | A2BH40 | | UPI000059E1 | NM_0010808 |
| YES | CCDS39247.1 | ENSMUSP0000 | P35235 | | UPI00000E5F | NM_011202.3 |
| YES | CCDS51673.1 | ENSMUSP0000 | Q99MR1 | | UPI00005643 | NM_031408.2 |
| YES | CCDS39400.1 | ENSMUSP0000 | | Q3UEW6 | UPI0000356E | NM_010229.2 |
| YES | CCDS51448.1 | ENSMUSP0000 | Q9CXK9 | | UPI0000D661 | NM_028234.1 |
| YES | CCDS40137.1 | ENSMUSP0000 | | F8VPK3 | UPI000017C8 | NM_010772.1 |
| YES | CCDS40137.1 | ENSMUSP0000 | | F8VPK3 | UPI000017C8 | NM_010772.1 |
| YES | CCDS22583.1 | ENSMUSP0000 | O88338 | Q546A8 | UPI00000238 | NM_007663.3 |
| YES | CCDS40507.2 | ENSMUSP0000 | E9Q4F7 | | UPI0000605E | NM_0010813 |
| YES | CCDS30452.1 | ENSMUSP0000 | Q9JHT5 | | UPI00000040 | NM_019496.4 |
| | | ENSMUSP0000 | | A0A5F8MPZ7 | UPI000047C1 | |
| YES | CCDS24532.1 | ENSMUSP0000 | Q61937 | Q5SQB7 | UPI0000003F | NM_0012522 |
| YES | CCDS49681.1 | ENSMUSP0000 | Q9EPQ8 | | UPI00001964 | NM_0011141 |
| YES | CCDS49884.1 | ENSMUSP0000 | | E9Q1Y1 | UPI00002159 | NM_0013687 |
| YES | CCDS28850.1 | ENSMUSP0000 | P30282 | Q3TSW4 | UPI00000188 | NM_0010816 |
| YES | CCDS29654.1 | ENSMUSP0000 | | Q7TQR3 | UPI00001AA6 | |
| YES | CCDS29654.1 | ENSMUSP0000 | | Q7TQR3 | UPI00001AA6 | |
| YES | CCDS70913.1 | ENSMUSP0000 | O54804 | | UPI00001ECA | NM_0012714 |
| YES | CCDS48372.1 | ENSMUSP0000 | Q3UPW2 | | UPI00005AC1 | NM_0011631 |
| YES | CCDS48372.1 | ENSMUSP0000 | Q3UPW2 | | UPI00005AC1 | NM_0011631 |
| YES | | ENSMUSP0000 | | A0A571BF02 | UPI00114BC6 | NM_0013709 |
| YES | CCDS39247.1 | ENSMUSP0000 | P35235 | | UPI00000E5F | NM_011202.3 |
| YES | CCDS51673.1 | ENSMUSP0000 | Q99MR1 | | UPI00005643 | NM_031408.2 |
| YES | CCDS39400.1 | ENSMUSP0000 | | Q3UEW6 | UPI0000356E | NM_010229.2 |
| YES | CCDS40137.1 | ENSMUSP0000 | | F8VPK3 | UPI000017C8 | NM_010772.1 |
| YES | CCDS40137.1 | ENSMUSP0000 | | F8VPK3 | UPI000017C8 | NM_010772.1 |
| YES | CCDS22651.1 | ENSMUSP0000 | | E9QMD3 | UPI000024E4 | |
| YES | CCDS23601.1 | ENSMUSP0000 | Q62312 | Q543C0 | UPI00000040 | NM_009371.3 |
| | | ENSMUSP0000 | | A0A5F8MPZ7 | UPI000047C1 | |
| YES | CCDS24532.1 | ENSMUSP0000 | Q61937 | Q5SQB7 | UPI0000003F | NM_0012522 |

| | | | | | | |
|---|---|---|---|---|---|---|
| YES | CCDS36599.1 | ENSMUSP000 | A2CG63 | | UPI00001D8A | NM_194262.2 |
| YES | CCDS36822.1 | ENSMUSP000 | Q7TMA2 | | UPI00001B06 | NM_145459.3 |
| YES | CCDS27267.1 | ENSMUSP000 | P13405 | | UPI0000157B | NM_009029.3 |
| YES | CCDS50256.1 | ENSMUSP000 | | E9PUR0 | UPI00015E16 | NM_175375.3 |
| YES | CCDS48372.1 | ENSMUSP000 | Q3UPW2 | | UPI00005AC1 | NM_0011631 |
| YES | CCDS48372.1 | ENSMUSP000 | Q3UPW2 | | UPI00005AC1 | NM_0011631 |
| YES | CCDS35808.1 | ENSMUSP000 | | B2RXC2 | UPI00001E2F | NM_0010811 |
| YES | | ENSMUSP000 | | A0A571BF02 | UPI00114BC6 | NM_0013709 |
| YES | CCDS17561.1 | ENSMUSP000 | | G3X9D9 | UPI0000022D | NM_008412.3 |
| YES | CCDS39247.1 | ENSMUSP000 | P35235 | | UPI00000E5F1 | NM_011202.3 |
| YES | CCDS57377.1 | ENSMUSP000 | | E9Q2E4 | UPI00023B36 | NM_181421.4 |
| YES | CCDS51673.1 | ENSMUSP000 | Q99MR1 | | UPI00005643 | NM_031408.2 |
| YES | CCDS19293.1 | ENSMUSP000 | O08786 | | UPI0000231 | NM_009827.2 |
| YES | CCDS40137.1 | ENSMUSP000 | | F8VPK3 | UPI000017C8 | NM_010772.1 |
| YES | CCDS40137.1 | ENSMUSP000 | | F8VPK3 | UPI000017C8 | NM_010772.1 |
| YES | CCDS40025.1 | ENSMUSP000 | | E9PWW9 | UPI0000605 | NM_0010812 |
| YES | CCDS22583.1 | ENSMUSP000 | O88338 | Q546A8 | UPI00000238 | NM_007663.3 |
| YES | CCDS52737.1 | ENSMUSP000 | | A0A0R4J170 | UPI00001C43 | NM_0011740 |
| YES | CCDS30452.1 | ENSMUSP000 | Q9JHT5 | | UPI0000004 | NM_019496.4 |
| YES | CCDS30545.1 | ENSMUSP000 | Q05738 | Q53WV7 | UPI0000005 | NM_011564.1 |
| YES | CCDS30545.1 | ENSMUSP000 | Q05738 | Q53WV7 | UPI0000005 | NM_011564.1 |
| YES | CCDS36062.1 | ENSMUSP000 | | B2RT41 | UPI0000609 | NM_0010332 |
| YES | CCDS48494.1 | ENSMUSP000 | Q8BYR2 | | UPI0000606 | NM_010690.1 |
| YES | CCDS24532.1 | ENSMUSP000 | Q61937 | Q5SQB7 | UPI0000003F | NM_0012522 |
| YES | CCDS26793.1 | ENSMUSP000 | O88704 | | UPI0000020F | NM_010408.3 |
| YES | CCDS27267.1 | ENSMUSP000 | P13405 | | UPI0000157B | NM_009029.3 |
| YES | CCDS49725.2 | ENSMUSP000 | | A0A0A0MQ73 | UPI00023B36 | NM_0010332 |
| YES | CCDS49884.1 | ENSMUSP000 | | E9Q1Y1 | UPI00002159 | NM_0013687 |
| YES | CCDS28827.2 | ENSMUSP000 | Q99PP2 | | UPI00015AA1 | NM_207671.4 |
| YES | CCDS15216.1 | ENSMUSP000 | | Q9D1Q5 | UPI0000220 | NM_198680.2 |
| YES | CCDS48372.1 | ENSMUSP000 | Q3UPW2 | | UPI00005AC1 | NM_0011631 |
| YES | CCDS48372.1 | ENSMUSP000 | Q3UPW2 | | UPI00005AC1 | NM_0011631 |
| YES | | ENSMUSP000 | | A0A571BF02 | UPI00114BC6 | NM_0013709 |
| YES | CCDS84520.1 | ENSMUSP000 | | A2API5 | UPI00015968 | NM_053011.2 |
| YES | CCDS17342.2 | ENSMUSP000 | | D4QGC2 | UPI00016116 | NM_0010041 |
| YES | CCDS38382.1 | ENSMUSP000 | | E9Q5A7 | UPI000024DC | NM_030708.2 |
| YES | CCDS39247.1 | ENSMUSP000 | P35235 | | UPI00000E5F1 | NM_011202.3 |
| YES | CCDS51673.1 | ENSMUSP000 | Q99MR1 | | UPI00005643 | NM_031408.2 |
| YES | CCDS51525.1 | ENSMUSP000 | P05532 | | UPI00000E98 | NM_0011227 |
| YES | CCDS20693.1 | ENSMUSP000 | P32883 | Q5J7N1 | UPI0000018 | NM_021284.6 |
| YES | CCDS21812.1 | ENSMUSP000 | Q07832 | Q3TPZ2 | UPI0000018 | NM_011121.4 |
| YES | CCDS40137.1 | ENSMUSP000 | | F8VPK3 | UPI000017C8 | NM_010772.1 |
| YES | CCDS40137.1 | ENSMUSP000 | | F8VPK3 | UPI000017C8 | NM_010772.1 |
| YES | CCDS40025.1 | ENSMUSP000 | | E9PWW9 | UPI0000605 | NM_0010812 |
| YES | CCDS23509.1 | ENSMUSP000 | Q62190 | | UPI0000193 | NM_009074.2 |
| YES | CCDS24532.1 | ENSMUSP000 | Q61937 | Q5SQB7 | UPI0000003F | NM_0012522 |
| YES | CCDS27158.1 | ENSMUSP000 | Q7TSJ6 | | UPI0000021D | NM_015771.2 |
| YES | CCDS49884.1 | ENSMUSP000 | | E9Q1Y1 | UPI00002159 | NM_0013687 |
| YES | CCDS48372.1 | ENSMUSP000 | Q3UPW2 | | UPI00005AC1 | NM_0011631 |
| YES | CCDS48372.1 | ENSMUSP000 | Q3UPW2 | | UPI00005AC1 | NM_0011631 |

| | | | | | | |
|---|---|---|---|---|---|---|
| YES | | ENSMUSP000( | | A0A571BF02 | UPI00114BC6: | NM_0013709: |
| YES | CCDS16973.2 | ENSMUSP000( | Q60710 | | UPI00015DF5I | NM_018851.4 |
| YES | CCDS17899.2 | ENSMUSP000( | | B2RY71 | UPI0001747B: | NM_172864.3 |
| YES | CCDS39247.1 | ENSMUSP000( | P35235 | | UPI00000E5F1 | NM_011202.3 |
| YES | CCDS51673.1 | ENSMUSP000( | Q99MR1 | | UPI00005643I | NM_031408.2 |
| YES | CCDS39400.1 | ENSMUSP000( | | Q3UEW6 | UPI0000356EI | NM_010229.2 |
| YES | CCDS40137.1 | ENSMUSP000( | | F8VPK3 | UPI000017C8I | NM_010772.1 |
| YES | CCDS40137.1 | ENSMUSP000( | | F8VPK3 | UPI000017C8I | NM_010772.1 |
| YES | CCDS80806.1 | ENSMUSP000( | | A0A087WQ44 | UPI0003D70E! | NM_0013042( |
| YES | CCDS21371.1 | ENSMUSP000( | Q6VNS1 | | UPI000002B6: | NM_008746.5 |
| YES | CCDS40507.2 | ENSMUSP000( | E9Q4F7 | | UPI0000605E! | NM_0010813: |
| YES | CCDS22764.1 | ENSMUSP000( | P68134 | | UPI00000008( | NM_009606.3 |
| YES | CCDS52532.1 | ENSMUSP000( | P0CL69 | F6M3K1 | UPI000154B5: | NM_0011015( |
| YES | CCDS41204.1 | ENSMUSP000( | | B1B0E8 | UPI0000ED93! | NM_009453.3 |
| YES | CCDS24532.1 | ENSMUSP000( | Q61937 | Q5SQB7 | UPI0000003F( | NM_0012522( |
| YES | CCDS48825.2 | ENSMUSP000( | | B1AR17 | UPI00001E5D/ | NM_146019.4 |
| YES | CCDS37149.1 | ENSMUSP000( | B2RWS6 | | UPI00004542I | NM_177821.6 |
| YES | CCDS49725.2 | ENSMUSP000( | | A0A0A0MQ73 | UPI00023B36/ | NM_0010332: |
| YES | CCDS37376.1 | ENSMUSP000( | | G5E843 | UPI0000354F/ | NM_019413.2 |
| YES | | ENSMUSP000( | | A0A571BF02 | UPI00114BC6: | NM_0013709: |
| YES | CCDS16973.2 | ENSMUSP000( | Q60710 | | UPI00015DF5I | NM_018851.4 |
| YES | CCDS17561.1 | ENSMUSP000( | | G3X9D9 | UPI0000022D: | NM_008412.3 |
| YES | CCDS39247.1 | ENSMUSP000( | P35235 | | UPI00000E5F1 | NM_011202.3 |
| YES | CCDS51673.1 | ENSMUSP000( | Q99MR1 | | UPI00005643I | NM_031408.2 |
| YES | CCDS39400.1 | ENSMUSP000( | | Q3UEW6 | UPI0000356EI | NM_010229.2 |
| YES | CCDS51448.1 | ENSMUSP000( | Q9CXK9 | | UPI0000D661! | NM_028234.1 |
| YES | CCDS40137.1 | ENSMUSP000( | | F8VPK3 | UPI000017C8I | NM_010772.1 |
| YES | CCDS40137.1 | ENSMUSP000( | | F8VPK3 | UPI000017C8I | NM_010772.1 |
| YES | CCDS21267.1 | ENSMUSP000( | Q03391 | | UPI0000D629I | NM_008172.2 |
| YES | CCDS40025.1 | ENSMUSP000( | | E9PWW9 | UPI00006059: | NM_0010812( |
| YES | CCDS40507.2 | ENSMUSP000( | E9Q4F7 | | UPI0000605E! | NM_0010813: |
| YES | CCDS52532.1 | ENSMUSP000( | P0CL69 | F6M3K1 | UPI000154B5: | NM_0011015( |
| YES | CCDS30315.1 | ENSMUSP000( | Q9JLM4 | B1AXS5 | UPI00000237: | NM_019831.3 |
| YES | CCDS30452.1 | ENSMUSP000( | Q9JHT5 | | UPI00000040( | NM_019496.4 |
| YES | CCDS40885.1 | ENSMUSP000( | Q99KG3 | | UPI0000027AI | NM_145627.3 |
| YES | CCDS41067.1 | ENSMUSP000( | Q7TS75 | | UPI0000EDA2I | NM_175179.4 |
| YES | CCDS24532.1 | ENSMUSP000( | Q61937 | Q5SQB7 | UPI0000003F( | NM_0012522( |
| YES | CCDS25119.1 | ENSMUSP000( | Q04690 | | UPI00000271( | NM_010897.2 |
| YES | CCDS49439.1 | ENSMUSP000( | E9PXT9 | | UPI0001B25B! | NM_0011644! |
| YES | CCDS49725.2 | ENSMUSP000( | | A0A0A0MQ73 | UPI00023B36/ | NM_0010332: |
| YES | CCDS49929.1 | ENSMUSP000( | E9Q4N7 | | UPI00006079( | NM_0010853! |
| YES | | ENSMUSP000( | | A0A571BF02 | UPI00114BC6: | NM_0013709: |
| YES | CCDS17561.1 | ENSMUSP000( | | G3X9D9 | UPI0000022D: | NM_008412.3 |
| YES | CCDS38908.1 | ENSMUSP000( | A2BH40 | | UPI000059E1! | NM_0010808: |
| YES | CCDS39247.1 | ENSMUSP000( | P35235 | | UPI00000E5F1 | NM_011202.3 |
| YES | CCDS51448.1 | ENSMUSP000( | Q9CXK9 | | UPI0000D661! | NM_028234.1 |
| YES | CCDS51889.1 | ENSMUSP000( | Q3UXZ9 | | UPI00006053! | NM_145997.2 |
| YES | CCDS51837.1 | ENSMUSP000( | | A0A0N4SWI3 | UPI0001BE80/ | NM_0011679: |
| YES | CCDS40137.1 | ENSMUSP000( | | F8VPK3 | UPI000017C8I | NM_010772.1 |
| YES | CCDS40137.1 | ENSMUSP000( | | F8VPK3 | UPI000017C8I | NM_010772.1 |

| | | | | | | |
|---|---|---|---|---|---|---|
| YES | CCDS80806.1 | ENSMUSP000 | | A0A087WQ44 | UPI0003D70E | NM_0013042 |
| YES | CCDS21267.1 | ENSMUSP000 | Q03391 | | UPI0000D629 | NM_008172.2 |
| YES | CCDS40603.1 | ENSMUSP000 | P55200 | | UPI00006061 | |
| YES | CCDS30452.1 | ENSMUSP000 | Q9JHT5 | | UPI00000040 | NM_019496.4 |
| | | ENSMUSP000 | | A0A5F8MPZ7 | UPI000047C1 | |
| YES | CCDS40885.1 | ENSMUSP000 | Q99KG3 | | UPI0000027A | NM_145627.3 |

| SIFT | EXON | INTRON | DOMAINS | PUBMED | IMPACT | PICK |
|---|---|---|---|---|---|---|
| tolerated(0.78 | 7/13 | | PANTHER:PTH | | MODERATE | 1 |
| | 11/11 | | Gene3D:1.10. | | HIGH | 1 |
| | 27/41 | | PANTHER:PTH | | LOW | 1 |
| tolerated(0.19 | 7/12 | | Gene3D:2.170 | | MODERATE | 1 |
| | 58/66 | | Low_complexi | | MODERATE | 1 |
| | 2/16 | | PROSITE_profi | | LOW | 1 |
| | 2/2 | | Coiled-coils_( | | MODERATE | 1 |
| deleterious(0. | 3/16 | | CDD:cd10340, | | MODERATE | 1 |
| | 22/24 | | Coiled-coils_( | | MODERATE | 1 |
| deleterious(0) | 20/24 | | Gene3D:1.10. | | MODERATE | 1 |
| tolerated(0.11 | 10/24 | | Gene3D:1.25. | | MODERATE | 1 |
| | 5/5 | | PANTHER:PTH | | LOW | 1 |
| | 5/5 | | PANTHER:PTH | | MODERATE | 1 |
| tolerated_low | 7/19 | | PANTHER:PTH | | MODERATE | |
| tolerated(1) | 1/3 | | PANTHER:PTH | | MODERATE | 1 |
| deleterious_lo | 1/16 | | Low_complexi | | MODERATE | |
| tolerated_low | 9/13 | | PANTHER:PTH | | MODERATE | |
| deleterious(0) | 8/16 | | Gene3D:3.30. | | MODERATE | 1 |
| tolerated_low | 4/24 | | Low_complexi | | MODERATE | 1 |
| deleterious(0) | 2/2 | | PANTHER:PTH | | MODERATE | 1 |
| | 11/11 | | Gene3D:1.10. | | HIGH | 1 |
| deleterious(0) | 33/58 | | Gene3D:1.10. | | MODERATE | 1 |
| | 33/58 | | Gene3D:1.10. | | HIGH | 1 |
| | 2/4 | | Low_complexi | | MODERATE | 1 |
| | 2/9 | | Low_complexi | | MODERATE | |
| deleterious(0. | 3/18 | | Gene3D:3.40. | | MODERATE | 1 |
| | 1/20 | | PANTHER:PTH | | MODERATE | 1 |
| | 4/4 | | PANTHER:PTH | | LOW | 1 |
| deleterious(0. | 3/16 | | CDD:cd10340, | | MODERATE | 1 |
| deleterious(0) | 20/24 | | Gene3D:1.10. | | MODERATE | 1 |
| | 5/5 | | PANTHER:PTH | | LOW | 1 |
| | 5/5 | | PANTHER:PTH | | MODERATE | 1 |
| | 11/11 | | Gene3D:1.10. | | HIGH | 1 |
| | 37/58 | | Gene3D:3.40. | | HIGH | 1 |
| | 1/27 | | Low_complexi | | LOW | 1 |
| deleterious(0. | 8/8 | | Gene3D:2.30. | | MODERATE | 1 |
| | 58/66 | | Low_complexi | | MODERATE | 1 |
| | 2/2 | | Coiled-coils_( | | MODERATE | 1 |
| deleterious(0. | 3/16 | | CDD:cd10340, | | MODERATE | 1 |
| | 22/24 | | Coiled-coils_( | | MODERATE | 1 |
| | 2/2 | | MobiDB_lite:n | | LOW | 1 |
| deleterious(0. | 6/10 | | PANTHER:PTH | | MODERATE | 1 |
| | 5/5 | | PANTHER:PTH | | LOW | 1 |
| | 5/5 | | PANTHER:PTH | | MODERATE | 1 |
| deleterious(0. | 4/6 | | Gene3D:1.10. | | MODERATE | 1 |
| tolerated(0.52 | 17/18 | | Gene3D:2.60. | | MODERATE | 1 |
| | 1/6 | | Low_complexi | | MODERATE | 1 |
| | 11/11 | | Gene3D:1.10. | | HIGH | 1 |
| | 1/15 | | Low_complexi | | MODERATE | 1 |

| | | | | |
|---|---|---|---|---|
| | 8/8 | Coiled-coils_(N | MODERATE | 1 |
| | 1/14 | MobiDB_lite:n | LOW | 1 |
| | 2/2 | PANTHER:PTH | MODERATE | 1 |
| | 1/12 | PANTHER:PTH | LOW | 1 |
| | 35/55 | PANTHER:PTH | MODERATE | 1 |
| | 2/4 | Low_complexi | MODERATE | 1 |
| | 58/66 | Low_complexi | MODERATE | 1 |
| | 2/5 | Low_complexi | LOW | 1 |
| | 2/2 | Coiled-coils_(N | MODERATE | 1 |
| deleterious(0.( | 3/16 | CDD:cd10340, | MODERATE | 1 |
| | 15/19 | Low_complexi | MODERATE | 1 |
| tolerated(0.45 | 5/5 | Pfam:PF00001 | MODERATE | 1 |
| | 5/5 | PANTHER:PTH | LOW | 1 |
| | 5/5 | PANTHER:PTH | MODERATE | 1 |
| tolerated_low | 34/34 | Low_complexi | MODERATE | |
| tolerated(0.52 | 17/18 | Gene3D:2.60.4 | MODERATE | 1 |
| tolerated(0.78 | 1/2 | PANTHER:PTH | MODERATE | 1 |
| | 1/6 | Low_complexi | MODERATE | 1 |
| tolerated_low | 4/24 | Low_complexi | MODERATE | 1 |
| | 1/1 | Low_complexi | MODERATE | 1 |
| | 11/11 | Gene3D:1.10.1 | HIGH | 1 |
| | 45/58 | PANTHER:PTH | HIGH | 1 |
| | 1/15 | Low_complexi | MODERATE | 1 |
| | 2/3 | Low_complexi | MODERATE | 1 |
| | 2/4 | Low_complexi | MODERATE | 1 |
| | 58/66 | Low_complexi | MODERATE | 1 |
| | 18/28 | Gene3D:1.25.4 | MODERATE | 1 |
| deleterious(0.( | 3/16 | CDD:cd10340, | MODERATE | 1 |
| | 22/24 | Coiled-coils_(N | MODERATE | 1 |
| deleterious(0) | 20/24 | Gene3D:1.10.5 | MODERATE | 1 |
| tolerated(0.45 | 5/5 | Pfam:PF00001 | MODERATE | 1 |
| | 1/6 | Low_complexi | MODERATE | 1 |
| | 3/7 | Low_complexi | LOW | 1 |
| tolerated_low | 2/2 | Coiled-coils_(N | MODERATE | 1 |
| | 11/11 | Gene3D:1.10.1 | HIGH | 1 |
| | 1/22 | Low_complexi | LOW | 1 |
| | 28/58 | Gene3D:1.10.5 | HIGH | 1 |
| | 8/8 | Coiled-coils_(N | MODERATE | 1 |
| | 40/55 | Coiled-coils_(N | MODERATE | 1 |
| | 58/66 | Low_complexi | MODERATE | 1 |
| | 46/52 | Low_complexi | LOW | 1 |
| deleterious(0.( | 3/16 | CDD:cd10340, | MODERATE | 1 |
| | 15/19 | Low_complexi | MODERATE | 1 |
| | 2/2 | MobiDB_lite:n | LOW | 1 |
| | 5/5 | PANTHER:PTH | LOW | 1 |
| | 5/5 | PANTHER:PTH | MODERATE | 1 |
| deleterious_lo | 1/16 | Low_complexi | MODERATE | |
| | 1/16 | Low_complexi | LOW | |
| tolerated(0.78 | 1/2 | PANTHER:PTH | MODERATE | 1 |
| | 3/11 | Gene3D:2.30.2 | LOW | 1 |

| | | | | |
|---|---|---|---|---|
| tolerated_low | 1/35 | Low_complexi | MODERATE | 1 |
| deleterious_lo | 4/6 | Gene3D:3.30. | MODERATE | 1 |
| | 11/11 | Gene3D:1.10. | HIGH | 1 |
| | 1/24 | MobiDB_lite:n | HIGH | 1 |
| | 5/16 | PANTHER:PTH | MODERATE | 1 |
| tolerated_low | 1/34 | PANTHER:PTH | MODERATE | 1 |
| | 3/9 | PROSITE_profi | LOW | 1 |
| | 3/9 | PROSITE_profi | LOW | 1 |
| | 1/10 | Low_complexi | LOW | 1 |
| | 58/66 | Low_complexi | MODERATE | 1 |
| deleterious_lo | 5/5 | PANTHER:PTH | MODERATE | 1 |
| | 1/20 | PANTHER:PTH | MODERATE | 1 |
| | 5/11 | PANTHER:PTH | LOW | 1 |
| deleterious(0. | 3/16 | CDD:cd10340, | MODERATE | 1 |
| | 1/76 | Low_complexi | MODERATE | 1 |
| | 11/17 | Coiled-coils_( | LOW | 1 |
| | 22/24 | Coiled-coils_( | MODERATE | 1 |
| deleterious(0) | 20/24 | Gene3D:1.10. | MODERATE | 1 |
| deleterious(0) | 16/21 | Low_complexi | MODERATE | 1 |
| | 5/5 | PANTHER:PTH | LOW | 1 |
| | 5/5 | PANTHER:PTH | MODERATE | 1 |
| tolerated_low | 7/19 | PANTHER:PTH | MODERATE | |
| tolerated(0.19 | 15/27 | Gene3D:2.60. | MODERATE | 1 |
| | 1/6 | Low_complexi | MODERATE | 1 |
| | 11/11 | Gene3D:1.10. | HIGH | 1 |
| | 7/18 | Low_complexi | MODERATE | 1 |
| | 2/4 | Low_complexi | MODERATE | 1 |
| tolerated(0.07 | 3/8 | Gene3D:2.10. | MODERATE | 1 |
| | 1/12 | PANTHER:PTH | MODERATE | |
| | 58/66 | Low_complexi | MODERATE | 1 |
| | 1/20 | PANTHER:PTH | MODERATE | 1 |
| | 5/11 | PANTHER:PTH | LOW | 1 |
| deleterious(0. | 3/16 | CDD:cd10340, | MODERATE | 1 |
| | 5/5 | PANTHER:PTH | LOW | 1 |
| | 5/5 | PANTHER:PTH | MODERATE | 1 |
| | 20/34 | Low_complexi | LOW | 1 |
| tolerated(0.78 | 1/2 | PANTHER:PTH | MODERATE | 1 |
| tolerated(1) | 14/14 | Gene3D:1.10. | MODERATE | 1 |
| | 11/11 | Gene3D:1.10. | HIGH | 1 |
| | 2/4 | Low_complexi | MODERATE | 1 |
| | 7/11 | PANTHER:PTH | LOW | 1 |
| | 2/22 | PANTHER:PTH | LOW | 1 |
| deleterious_lo | 2/2 | Gene3D:1.20. | MODERATE | 1 |
| | 2/2 | Gene3D:1.20. | LOW | 1 |
| | 3/9 | PROSITE_profi | LOW | 1 |
| | 3/9 | PROSITE_profi | LOW | 1 |
| | 58/66 | Low_complexi | MODERATE | 1 |
| | 13/32 | Low_complexi | LOW | 1 |
| deleterious(0. | 34/34 | PIRSF:PIRSF00 | MODERATE | 1 |
| deleterious(0. | 3/16 | CDD:cd10340, | MODERATE | 1 |

| | | | | |
|---|---|---|---|---|
| | 22/24 | Coiled-coils_(N | MODERATE | 1 |
| deleterious(0) | 20/24 | Gene3D:1.10.5 | MODERATE | 1 |
| | 5/5 | PANTHER:PTH | LOW | 1 |
| | 5/5 | PANTHER:PTH | MODERATE | 1 |
| deleterious(0.0 | 11/12 | CDD:cd13718, | MODERATE | 1 |
| tolerated(0.26 | 5/17 | Gene3D:3.80.1 | MODERATE | 1 |
| tolerated(0.78 | 1/2 | PANTHER:PTH | MODERATE | 1 |
| deleterious_lo | 3/36 | PIRSF:PIRSF01 | MODERATE | 1 |
| | 1/6 | Low_complexi | MODERATE | 1 |
| tolerated_low | 4/24 | Low_complexi | MODERATE | 1 |
| tolerated_low | 2/2 | Coiled-coils_(N | MODERATE | 1 |
| tolerated(0.16 | 27/44 | Gene3D:2.120 | MODERATE | 1 |
| | 11/11 | Gene3D:1.10.1 | HIGH | 1 |
| | 9/10 | PANTHER:PTH | LOW | 1 |
| tolerated_low | 17/17 | MobiDB_lite:m | MODERATE | 1 |
| | 3/9 | PROSITE_profi | LOW | 1 |
| | 3/9 | PROSITE_profi | LOW | 1 |
| | 58/66 | Low_complexi | MODERATE | 1 |
| | 2/16 | PROSITE_profi | LOW | 1 |
| deleterious(0) | 5/12 | Pfam:PF16978 | MODERATE | 1 |
| | 1/20 | PANTHER:PTH | MODERATE | 1 |
| deleterious(0.0 | 3/16 | CDD:cd10340, | MODERATE | 1 |
| | 22/24 | Coiled-coils_(N | MODERATE | 1 |
| deleterious(0) | 20/24 | Gene3D:1.10.5 | MODERATE | 1 |
| | 15/19 | Low_complexi | MODERATE | 1 |
| | 5/5 | PANTHER:PTH | LOW | 1 |
| | 5/5 | PANTHER:PTH | MODERATE | 1 |
| tolerated(0.52 | 17/18 | Gene3D:2.60.4 | MODERATE | 1 |
| tolerated_low | 9/13 | PANTHER:PTH | MODERATE | |
| | 1/6 | Low_complexi | MODERATE | 1 |
| | 24/27 | Low_complexi | MODERATE | |
| | 11/11 | Gene3D:1.10.1 | HIGH | 1 |
| | 2/6 | Coiled-coils_(N | LOW | 1 |
| | 2/4 | Low_complexi | MODERATE | 1 |
| | 6/7 | PANTHER:PTH | LOW | |
| deleterious_lo | 2/2 | Gene3D:1.20.1 | MODERATE | 1 |
| | 2/2 | Gene3D:1.20.1 | LOW | 1 |
| | 1/12 | PANTHER:PTH | MODERATE | |
| | 3/9 | PROSITE_profi | LOW | 1 |
| | 3/9 | PROSITE_profi | LOW | 1 |
| | 58/66 | Low_complexi | MODERATE | 1 |
| deleterious(0.0 | 3/16 | CDD:cd10340, | MODERATE | 1 |
| | 22/24 | Coiled-coils_(N | MODERATE | 1 |
| deleterious(0) | 20/24 | Gene3D:1.10.5 | MODERATE | 1 |
| | 5/5 | PANTHER:PTH | LOW | 1 |
| | 5/5 | PANTHER:PTH | MODERATE | 1 |
| | 15/16 | PANTHER:PTH | LOW | 1 |
| tolerated(0.07 | 5/8 | PIRSF:PIRSF03 | MODERATE | 1 |
| | 24/27 | Low_complexi | MODERATE | |
| | 11/11 | Gene3D:1.10.1 | HIGH | 1 |

| | | | | |
|---|---|---|---|---|
| tolerated_low | 20/24 | Low_complexi | MODERATE | 1 |
| | 2/3 | Low_complexi | MODERATE | 1 |
| | 1/27 | Low_complexi | LOW | 1 |
| tolerated_low | 1/34 | PANTHER:PTH | MODERATE | 1 |
| | 3/9 | PROSITE_profi | LOW | 1 |
| | 3/9 | PROSITE_profi | LOW | 1 |
| tolerated_low | 2/8 | MobiDB_lite:n | MODERATE | 1 |
| | 58/66 | Low_complexi | MODERATE | 1 |
| | 2/2 | Coiled-coils_(N | MODERATE | 1 |
| deleterious(0.( | 3/16 | CDD:cd10340, | MODERATE | 1 |
| | 1/76 | Low_complexi | MODERATE | 1 |
| | 22/24 | Coiled-coils_(N | MODERATE | 1 |
| tolerated(0.45 | 5/5 | Pfam:PF00001 | MODERATE | 1 |
| | 5/5 | PANTHER:PTH | LOW | 1 |
| | 5/5 | PANTHER:PTH | MODERATE | 1 |
| | 1/16 | Low_complexi | LOW | |
| tolerated(0.52 | 17/18 | Gene3D:2.60.4 | MODERATE | 1 |
| tolerated(0.1) | 32/34 | PANTHER:PTH | MODERATE | 1 |
| | 1/6 | Low_complexi | MODERATE | 1 |
| deleterious_lo | 1/1 | Low_complexi | MODERATE | 1 |
| | 1/1 | Low_complexi | MODERATE | 1 |
| | 1/35 | Low_complexi | MODERATE | 1 |
| deleterious(0) | 5/8 | PANTHER:PTH | MODERATE | 1 |
| | 11/11 | Gene3D:1.10.1 | HIGH | 1 |
| | 8/8 | Coiled-coils_(N | MODERATE | 1 |
| | 1/27 | Low_complexi | LOW | 1 |
| | 40/55 | Coiled-coils_(N | LOW | 1 |
| | 2/4 | Low_complexi | MODERATE | 1 |
| | 10/10 | Coiled-coils_(N | MODERATE | 1 |
| deleterious(0.( | 8/8 | Gene3D:2.30.3 | MODERATE | 1 |
| | 3/9 | PROSITE_profi | LOW | 1 |
| | 3/9 | PROSITE_profi | LOW | 1 |
| | 58/66 | Low_complexi | MODERATE | 1 |
| | 14/91 | Gene3D:2.120 | LOW | 1 |
| | 2/5 | Low_complexi | LOW | 1 |
| | 10/11 | PANTHER:PTH | LOW | 1 |
| deleterious(0.( | 3/16 | CDD:cd10340, | MODERATE | 1 |
| | 22/24 | Coiled-coils_(N | MODERATE | 1 |
| deleterious(0) | 17/21 | PROSITE_profi | MODERATE | 1 |
| deleterious(0) | 2/5 | Low_complexi | MODERATE | 1 |
| deleterious(0.( | 6/10 | PANTHER:PTH | MODERATE | 1 |
| | 5/5 | PANTHER:PTH | LOW | 1 |
| | 5/5 | PANTHER:PTH | MODERATE | 1 |
| | 1/16 | Low_complexi | LOW | |
| | 4/19 | Gene3D:3.30.1 | LOW | 1 |
| | 11/11 | Gene3D:1.10.1 | HIGH | 1 |
| tolerated(0.19 | 8/8 | PROSITE_profi | MODERATE | 1 |
| | 2/4 | Low_complexi | MODERATE | 1 |
| | 3/9 | PROSITE_profi | LOW | 1 |
| | 3/9 | PROSITE_profi | LOW | 1 |

| | | | | |
|---|---|---|---|---|
| | 58/66 | Low_complexi | MODERATE | 1 |
| | 2/16 | PROSITE_profi | LOW | 1 |
| tolerated(0.26 | 2/23 | MobiDB_lite:n | MODERATE | 1 |
| deleterious(0.( | 3/16 | CDD:cd10340, | MODERATE | 1 |
| | 22/24 | Coiled-coils_(N | MODERATE | 1 |
| deleterious(0) | 20/24 | Gene3D:1.10.! | MODERATE | 1 |
| | 5/5 | PANTHER:PTH | LOW | 1 |
| | 5/5 | PANTHER:PTH | MODERATE | 1 |
| | 20/34 | Low_complexi | LOW | 1 |
| deleterious(0.( | 12/17 | PANTHER:PTH | MODERATE | 1 |
| tolerated_low | 9/13 | PANTHER:PTH | MODERATE | |
| | 3/7 | Gene3D:3.30.4 | LOW | 1 |
| tolerated(0.78 | 1/2 | PANTHER:PTH | MODERATE | 1 |
| deleterious(0.( | 11/11 | PANTHER:PTH | MODERATE | 1 |
| | 11/11 | Gene3D:1.10.1 | HIGH | 1 |
| | 1/40 | PANTHER:PTH | MODERATE | 1 |
| deleterious(0) | 6/31 | PANTHER:PTH | MODERATE | 1 |
| | 35/55 | Coiled-coils_(N | MODERATE | 1 |
| tolerated(0.29 | 27/29 | MobiDB_lite:n | MODERATE | 1 |
| | 58/66 | Low_complexi | MODERATE | 1 |
| | 2/16 | PROSITE_profi | LOW | 1 |
| | 2/2 | Coiled-coils_(N | MODERATE | 1 |
| deleterious(0.( | 3/16 | CDD:cd10340, | MODERATE | 1 |
| | 22/24 | Coiled-coils_(N | MODERATE | 1 |
| deleterious(0) | 20/24 | Gene3D:1.10.! | MODERATE | 1 |
| | 15/19 | Low_complexi | MODERATE | 1 |
| | 5/5 | PANTHER:PTH | LOW | 1 |
| | 5/5 | PANTHER:PTH | MODERATE | 1 |
| deleterious(0.( | 11/12 | CDD:cd13718, | MODERATE | 1 |
| deleterious_lo | 1/16 | Low_complexi | MODERATE | |
| tolerated_low | 9/13 | PANTHER:PTH | MODERATE | |
| tolerated(0.78 | 1/2 | PANTHER:PTH | MODERATE | 1 |
| | 2/25 | PANTHER:PTH | LOW | 1 |
| | 1/6 | Low_complexi | MODERATE | 1 |
| tolerated_low | 4/24 | Low_complexi | MODERATE | 1 |
| tolerated_low | 2/2 | Coiled-coils_(N | MODERATE | 1 |
| | 11/11 | Gene3D:1.10.1 | HIGH | 1 |
| | 28/58 | Gene3D:1.10.! | HIGH | 1 |
| | 2/2 | PANTHER:PTH | MODERATE | 1 |
| | 35/55 | Coiled-coils_(N | MODERATE | 1 |
| | 6/20 | PANTHER:PTH | LOW | 1 |
| | 58/66 | Low_complexi | MODERATE | 1 |
| | 2/2 | Coiled-coils_(N | MODERATE | 1 |
| | 1/20 | PANTHER:PTH | MODERATE | 1 |
| deleterious(0.( | 3/16 | CDD:cd10340, | MODERATE | 1 |
| | 15/19 | Low_complexi | MODERATE | 1 |
| deleterious(0) | 23/28 | PANTHER:PTH | MODERATE | 1 |
| | 2/2 | MobiDB_lite:n | LOW | 1 |
| | 5/5 | PANTHER:PTH | LOW | 1 |
| | 5/5 | PANTHER:PTH | MODERATE | 1 |
