## Supplemental Data Statistics for "*PTPN11* Mutation Clonal Hierarchy in Acute Myeloid Leukemia"

| Report on 2024-03-xx by sheet in the excel data file |
|---|
| 1. For sheet Fig 2a, we fit Cox-PH model for Day and ANOVA model for log(Day) about Group using contr |
| 2. For other sheets, we fit ANOVA for log(Weight) or log(Count) about Group usinfgcontrast for each pair of group |

| Notes |
| --- |
| Estimate: coefficient estimate in ANOVA model |
| Pvalue: the p-value for test coefficient=0 |
| fdr_p: Adjusted FDR p-value |
| OverallP: p-value of Group |
| DIfM_LCL: lower limit of 95% CI for difference of 2 group means after logit or log transformation |
| DIfM_UCL: upper limit of 95% CI for difference of 2 group means after logit or log transformation |
| GMean: group mean calculated after logit or log transformation, then back to the original response value |
| GMean_LCL: lower limit of 95% CI of group mean after logit or log transformation, then back to the original respo[illegible] |
| GMean_UCL: upper limit of 95% CI of group mean after logit or log transformation, then back to the original respo[illegible] |
| Mean_O: Mean calculated from the original response value |
| Std_O: Standard deviation of original response value |
| Min_O: minmum value of original response value |
| Max_O: maximum value of original response value |

| |
|---|
| nse value |
| onse value |

| COX PH model | | | | | | | | |
|---|---|---|---|---|---|---|---|---|
| **Variables** | **Class** | **Est** | **pval** | **HazardRatio** | **HRLowerCL** | **HRUpperCL** | **Contrast** | **Estimate** |
| Group | G2 | -3.3373 | **0.0001** | 0.0355 | 0.0067 | 0.1897 | | |
| Group | G3 | -3.1927 | **0.0001** | 0.0411 | 0.0083 | 0.2031 | | |
| | | | | | | | G2 vs G1 | -3.3373 |
| | | | | | | | G3 vs G1 | -3.1927 |
| | | | | | | | G3 vs G2 | 0.1446 |

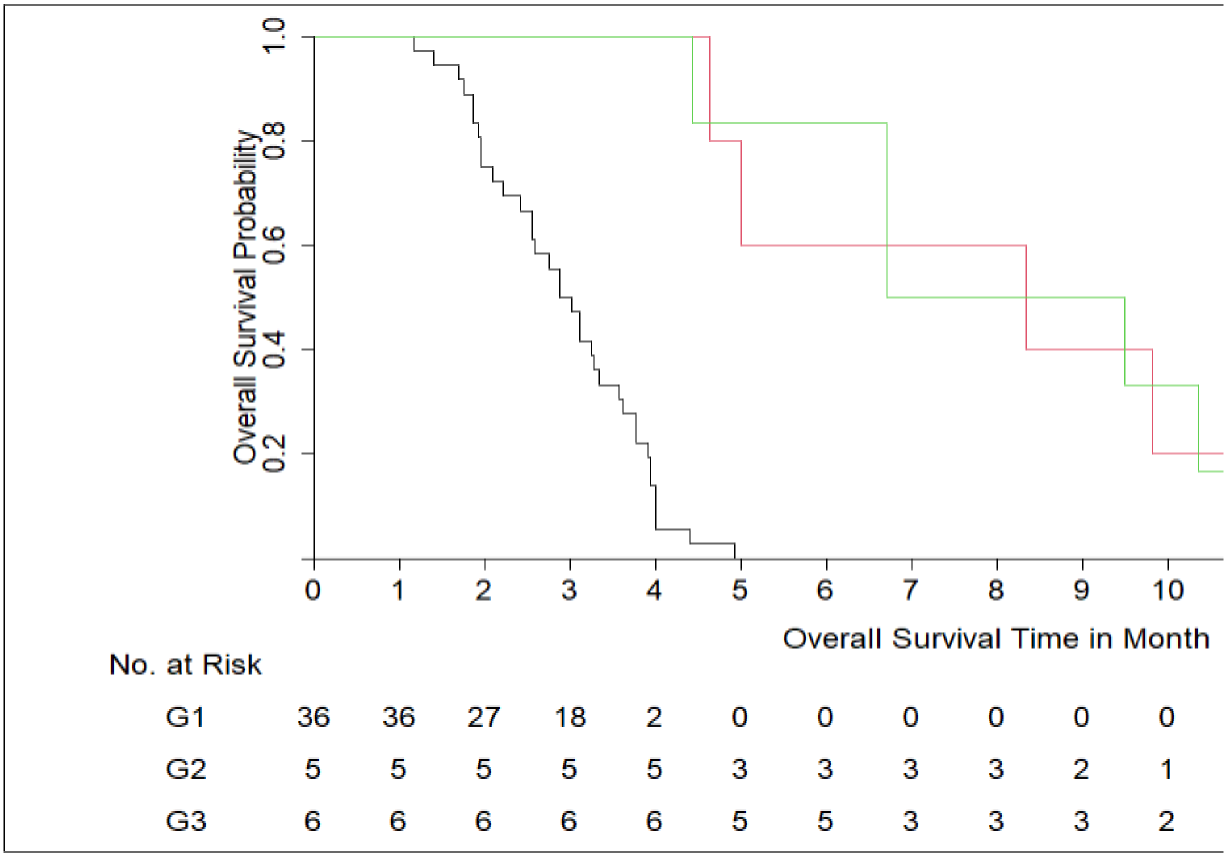

| | |
|---|---|
| **Group Names** | |
| Ptpn11$^{E76K}$/Npm1$^{cA}$ | G1 |
| Ptpn11$^{E76K}$ | G2 |
| Npm1$^{cA}$ | G3 |

| LowerLimit | UpperLimit | Pvalue | fdr_p |
|---|---|---|---|
| -5.0122 | -1.6624 | **0.0001** | 0.0001 |
| -4.7915 | -1.5939 | **0.0001** | 0.0001 |
| -1.1288 | 1.4181 | 0.8238 | 0.8238 |

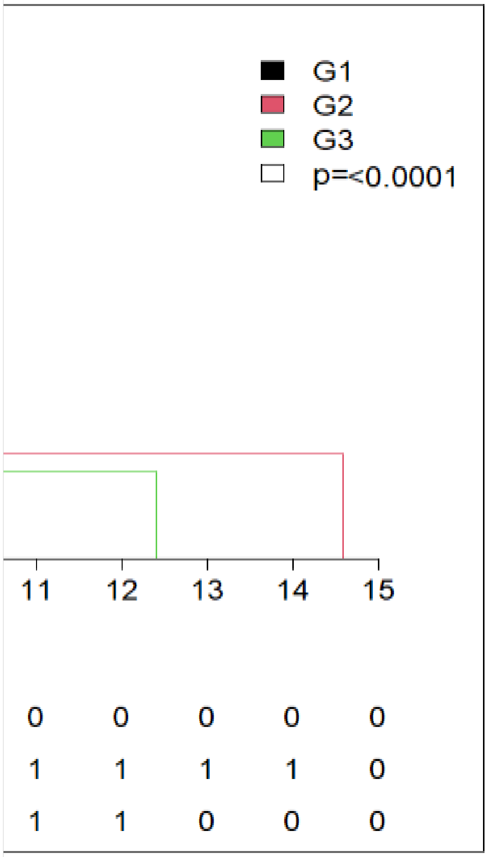

| Source | Parameter | Estimate | Pvalue | fdr_p |
|---|---|---|---|---|
| Intercept | Intercept | 4.4262 | 2.351E-47 | 1.411E-46 |
| Group | Group G2 | 1.0280 | **5.446E-07** | **6.536E-07** |
| Group | Group G3 | 1.0499 | **6.730E-08** | **1.346E-07** |
| Group | Group G1 | 0.0000 | | |
| Contrast | G2 vs G1 | 1.0280 | **5.446E-07** | **6.536E-07** |
| Contrast | G3 vs G1 | 1.0499 | **6.730E-08** | **1.346E-07** |
| Contrast | G3 vs G2 | 0.0219 | 9.220E-01 | 9.220E-01 |
| Mean_CI | G1 | | | |
| Mean_CI | G2 | | | |
| Mean_CI | G3 | | | |

| ANOVA Model for Log(Day) | | | | | | | | |
|---|---|---|---|---|---|---|---|---|
| **OverallP** | **DIfM_LCL** | **DiffM_UCL** | **GMean** | **GMean_LCL** | **GMean_UCL** | **Mean_O** | **Std_O** | **Min_O** |
| **1.327E-09** | | | | | | | | |
| | 0.6744 | 1.3817 | | | | | | |
| | 0.7232 | 1.3767 | | | | | | |
| | -0.4268 | 0.4706 | | | | | | |
| | | | 83.6144 | 73.9003 | 94.6053 | 88.4444 | 28.4358 | 35 |
| | | | 233.7438 | 167.8120 | 325.5797 | 256.2000 | 122.8849 | 140 |
| | | | 238.9236 | 176.5555 | 323.3232 | 252.5000 | 88.0903 | 134 |

| |
|---|
| Max_O |
| 149 |
| 441 |
| 375 |

| ANVOA Model for Log(We | | | | | | |
|---|---|---|---|---|---|---|
| **Source** | **Parameter** | **Estimate** | **Pvalue** | **fdr_p** | **OverallP** | **DIfM_LCL** |
| Intercept | Intercept | -2.6829 | 4.523E-24 | 4.523E-23 | | |
| Group | Group G2 | -0.0791 | 6.505E-01 | 6.505E-01 | **1.711E-21** | |
| Group | Group G3 | 1.6305 | **5.249E-11** | **8.748E-11** | | |
| Group | Group G4 | 2.8857 | **2.968E-21** | **9.892E-21** | | |
| Group | Group G1 | 0.0000 | | | | |
| Contrast | G2 vs G1 | -0.0791 | 6.505E-01 | 6.505E-01 | | -0.4308 |
| Contrast | G3 vs G1 | 1.6305 | **5.249E-11** | **8.748E-11** | | 1.2788 |
| Contrast | G4 vs G1 | 2.8857 | **2.968E-21** | **9.892E-21** | | 2.6106 |
| Contrast | G3 vs G2 | 1.7096 | **3.803E-10** | **5.433E-10** | | 1.3095 |
| Contrast | G4 vs G2 | 2.9648 | **5.928E-19** | **1.482E-18** | | 2.6301 |
| Contrast | G4 vs G3 | 1.2553 | **7.429E-09** | **9.287E-09** | | 0.9205 |
| Mean_CI | G1 | | | | | |
| Mean_CI | G2 | | | | | |
| Mean_CI | G3 | | | | | |
| Mean_CI | G4 | | | | | |
| **Group Names** | | | | | | |
| MxCre | G1 | | | | | |
| NPM1(fl/WT)/MxCre | G2 | | | | | |
| PTPN11/MxCre | G3 | | | | | |
| PTPN11/NPM1(fl/WT)/MxCre | G4 | | | | | |

| ight) | | | | | | | |
|---|---|---|---|---|---|---|---|
| DiffM_UCL | GMean | GMean_LCL | GMean_UCL | Mean_O | Std_O | Min_O | Max_O |
| 0.2726 | | | | | | | |
| 1.9822 | | | | | | | |
| 3.1608 | | | | | | | |
| 2.1096 | | | | | | | |
| 3.2996 | | | | | | | |
| 1.5900 | | | | | | | |
| | 0.0684 | 0.0555 | 0.0842 | 0.0700 | 0.016 | 0.05 | 0.1 |
| | 0.0632 | 0.0476 | 0.0838 | 0.0667 | 0.021 | 0.03 | 0.09 |
| | 0.3491 | 0.2631 | 0.4632 | 0.3883 | 0.213 | 0.21 | 0.78 |
| | 1.2248 | 1.0241 | 1.4648 | 1.2807 | 0.367 | 0.53 | 2.03 |

| ANVOA | | | | | | | | |
|---|---|---|---|---|---|---|---|---|
| | | | | | | ANOVA Model for Log(Count) for 18 S | | |
| Substance | Source | Parameter | Estimate | Pvalue | fdr_p | OverallP | DIfM_LCL | DiffM_UCL |
| WBC | Intercept | Intercept | 2.4066 | 4.276E-15 | 4.276E-14 | | | |
| | Group | Group G2 | -0.1804 | 5.637E-01 | 5.637E-01 | **1.386E-12** | | |
| | Group | Group G3 | 0.9683 | **1.477E-02** | **1.846E-02** | | | |
| | Group | Group G4 | 2.6595 | **1.677E-12** | **5.591E-12** | | | |
| | Group | Group G1 | 0.0000 | | | | | |
| | Contrast | G2 vs G1 | -0.1804 | 5.637E-01 | 5.637E-01 | | -0.8088 | 0.4480 |
| | Contrast | G3 vs G1 | 0.9683 | **1.477E-02** | **1.846E-02** | | 0.2019 | 1.7348 |
| | Contrast | G4 vs G1 | 2.6595 | **1.677E-12** | **5.591E-12** | | 2.1515 | 3.1674 |
| | Contrast | G3 vs G2 | 1.1487 | **8.784E-03** | **1.464E-02** | | 0.3086 | 1.9889 |
| | Contrast | G4 vs G2 | 2.8399 | **4.217E-11** | **1.054E-10** | | 2.2263 | 3.4535 |
| | Contrast | G4 vs G3 | 1.6911 | **6.170E-05** | **1.234E-04** | | 0.9368 | 2.4455 |
| | Mean_CI | G1 | | | | | | |
| | Mean_CI | G2 | | | | | | |
| | Mean_CI | G3 | | | | | | |
| | Mean_CI | G4 | | | | | | |
| PLT | Source | Parameter | Estimate | Pvalue | fdr_p | OverallP | DIfM_LCL | DiffM_UCL |
| | Intercept | Intercept | 7.0254 | 1.999E-29 | 1.999E-28 | | | |
| | Group | Group G2 | -0.6665 | **4.841E-02** | 6.915E-02 | **2.729E-10** | | |
| | Group | Group G3 | -0.3079 | 4.438E-01 | 4.438E-01 | | | |
| | Group | Group G4 | -2.4695 | **4.464E-11** | **1.488E-10** | | | |
| | Group | Group G1 | 0.0000 | | | | | |
| | Contrast | G2 vs G1 | -0.6665 | **4.841E-02** | 6.915E-02 | | -1.3281 | -0.0049 |
| | Contrast | G3 vs G1 | -0.3079 | 4.438E-01 | 4.438E-01 | | -1.1148 | 0.4990 |
| | Contrast | G4 vs G1 | -2.4695 | **4.464E-11** | **1.488E-10** | | -3.0043 | -1.9347 |
| | Contrast | G3 vs G2 | 0.3587 | 4.160E-01 | 4.438E-01 | | -0.5259 | 1.2432 |
| | Contrast | G4 vs G2 | -1.8030 | **2.127E-06** | **5.318E-06** | | -2.4489 | -1.1570 |
| | Contrast | G4 vs G3 | -2.1616 | **3.258E-06** | **6.516E-06** | | -2.9558 | -1.3675 |
| | Mean_CI | G1 | | | | | | |
| | Mean_CI | G2 | | | | | | |
| | Mean_CI | G3 | | | | | | |
| | Mean_CI | G4 | | | | | | |
| RBC | Source | Parameter | Estimate | Pvalue | fdr_p | OverallP | DIfM_LCL | DiffM_UCL |
| | Intercept | Intercept | 2.2587 | 2.082E-25 | 2.082E-24 | | | |
| | Group | Group G2 | -0.2484 | 7.982E-02 | 1.140E-01 | **7.191E-09** | | |
| | Group | Group G3 | -0.0273 | 8.716E-01 | 8.716E-01 | | | |
| | Group | Group G4 | -0.8989 | **1.652E-09** | **5.506E-09** | | | |
| | Group | Group G1 | 0.0000 | | | | | |
| | Contrast | G2 vs G1 | -0.2484 | 7.982E-02 | 1.140E-01 | | -0.5280 | 0.0311 |
| | Contrast | G3 vs G1 | -0.0273 | 8.716E-01 | 8.716E-01 | | -0.3683 | 0.3136 |
| | Contrast | G4 vs G1 | -0.8989 | **1.652E-09** | **5.506E-09** | | -1.1249 | -0.6730 |
| | Contrast | G3 vs G2 | 0.2211 | 2.379E-01 | 2.973E-01 | | -0.1527 | 0.5949 |
| | Contrast | G4 vs G2 | -0.6505 | **2.614E-05** | **5.227E-05** | | -0.9235 | -0.3775 |
| | Contrast | G4 vs G3 | -0.8716 | **7.020E-06** | **1.755E-05** | | -1.2072 | -0.5360 |

| | | | | | | | | |
|---|---|---|---|---|---|---|---|---|
| | Mean_CI | G1 | | | | | | |
| | Mean_CI | G2 | | | | | | |
| | Mean_CI | G3 | | | | | | |
| | Mean_CI | G4 | | | | | | |
| **Neu** | **Source** | **Parameter** | **Estimate** | **Pvalue** | **fdr_p** | **OverallP** | **DIfM_LCL** | **DiffM_UCL** |
| | Intercept | Intercept | 0.4392 | 5.325E-02 | 7.092E-02 | | | |
| | Group | Group G2 | 0.1428 | 7.028E-01 | 7.028E-01 | **2.265E-15** | | |
| | Group | Group G3 | 1.1204 | **1.829E-02** | **3.049E-02** | | | |
| | Group | Group G4 | 4.0994 | **1.348E-15** | **6.741E-15** | | | |
| | Group | Group G1 | 0.0000 | | | | | |
| | Contrast | G2 vs G1 | 0.1428 | 7.028E-01 | 7.028E-01 | | -0.6106 | 0.8962 |
| | Contrast | G3 vs G1 | 1.1204 | **1.829E-02** | **3.049E-02** | | 0.2016 | 2.0393 |
| | Contrast | G4 vs G1 | 4.0994 | **1.348E-15** | **6.741E-15** | | 3.4905 | 4.7084 |
| | Contrast | G3 vs G2 | 0.9777 | 5.674E-02 | 7.092E-02 | | -0.0296 | 1.9849 |
| | Contrast | G4 vs G2 | 3.9567 | **8.076E-13** | **2.692E-12** | | 3.2211 | 4.6923 |
| | Contrast | G4 vs G3 | 2.9790 | **9.707E-08** | **2.427E-07** | | 2.0747 | 3.8833 |
| | Mean_CI | G1 | | | | | | |
| | Mean_CI | G2 | | | | | | |
| | Mean_CI | G3 | | | | | | |
| | Mean_CI | G4 | | | | | | |
| **NeuPercent** | **Source** | **Parameter** | **Estimate** | **Pvalue** | **fdr_p** | **OverallP** | **DIfM_LCL** | **DiffM_UCL** |
| | Intercept | Intercept | 2.6389 | 5.240E-27 | 5.240E-26 | | | |
| | Group | Group G2 | 0.3216 | **3.228E-02** | **4.612E-02** | **1.071E-13** | | |
| | Group | Group G3 | 0.1543 | 3.863E-01 | 3.916E-01 | | | |
| | Group | Group G4 | 1.4394 | **2.594E-14** | **8.645E-14** | | | |
| | Group | Group G1 | 0.0000 | | | | | |
| | Contrast | G2 vs G1 | 0.3216 | **3.228E-02** | **4.612E-02** | | 0.0288 | 0.6143 |
| | Contrast | G3 vs G1 | 0.1543 | 3.863E-01 | 3.916E-01 | | -0.2028 | 0.5114 |
| | Contrast | G4 vs G1 | 1.4394 | **2.594E-14** | **8.645E-14** | | 1.2027 | 1.6760 |
| | Contrast | G3 vs G2 | -0.1672 | 3.916E-01 | 3.916E-01 | | -0.5587 | 0.2242 |
| | Contrast | G4 vs G2 | 1.1178 | **2.453E-09** | **6.132E-09** | | 0.8320 | 1.4037 |
| | Contrast | G4 vs G3 | 1.2851 | **1.095E-08** | **2.189E-08** | | 0.9336 | 1.6365 |
| | Mean_CI | G1 | | | | | | |
| | Mean_CI | G2 | | | | | | |
| | Mean_CI | G3 | | | | | | |
| | Mean_CI | G4 | | | | | | |
| **Bas** | **Source** | **Parameter** | **Estimate** | **Pvalue** | **fdr_p** | **OverallP** | **DIfM_LCL** | **DiffM_UCL** |
| | Intercept | Intercept | -2.6013 | 1.297E-03 | 1.297E-02 | | | |
| | Group | Group G2 | 0.1315 | 9.173E-01 | 9.173E-01 | 2.822E-01 | | |
| | Group | Group G3 | 1.4953 | 3.361E-01 | 5.602E-01 | | | |
| | Group | Group G4 | 1.8129 | 8.306E-02 | 2.769E-01 | | | |
| | Group | Group G1 | 0.0000 | | | | | |
| | Contrast | G2 vs G1 | 0.1315 | 9.173E-01 | 9.173E-01 | | -2.4206 | 2.6835 |
| | Contrast | G3 vs G1 | 1.4953 | 3.361E-01 | 5.602E-01 | | -1.6173 | 4.6078 |

| | | | | | | | | |
|---|---|---|---|---|---|---|---|---|
| | Contrast | G4 vs G1 | 1.8129 | 8.306E-02 | 2.769E-01 | | -0.2499 | 3.8757 |
| | Contrast | G3 vs G2 | 1.3638 | 4.226E-01 | 6.037E-01 | | -2.0482 | 4.7758 |
| | Contrast | G4 vs G2 | 1.6814 | 1.794E-01 | 4.486E-01 | | -0.8103 | 4.1732 |
| | Contrast | G4 vs G3 | 0.3177 | 8.345E-01 | 9.173E-01 | | -2.7457 | 3.3810 |
| | Mean_CI | G1 | | | | | | |
| | Mean_CI | G2 | | | | | | |
| | Mean_CI | G3 | | | | | | |
| | Mean_CI | G4 | | | | | | |
| **BasPercent** | **Source** | **Parameter** | **Estimate** | **Pvalue** | **fdr_p** | **OverallP** | **DIfM_LCL** | **DiffM_UCL** |
| | Intercept | Intercept | -0.4446 | 5.251E-01 | 8.803E-01 | | | |
| | Group | Group G2 | 0.3107 | 7.922E-01 | 8.803E-01 | 8.412E-01 | | |
| | Group | Group G3 | 0.5346 | 7.103E-01 | 8.803E-01 | | | |
| | Group | Group G4 | -0.4907 | 6.073E-01 | 8.803E-01 | | | |
| | Group | Group G1 | 0.0000 | | | | | |
| | Contrast | G2 vs G1 | 0.3107 | 7.922E-01 | 8.803E-01 | | -2.0658 | 2.6873 |
| | Contrast | G3 vs G1 | 0.5346 | 7.103E-01 | 8.803E-01 | | -2.3639 | 3.4331 |
| | Contrast | G4 vs G1 | -0.4907 | 6.073E-01 | 8.803E-01 | | -2.4116 | 1.4303 |
| | Contrast | G3 vs G2 | 0.2239 | 8.871E-01 | 8.871E-01 | | -2.9535 | 3.4013 |
| | Contrast | G4 vs G2 | -0.8014 | 4.878E-01 | 8.803E-01 | | -3.1218 | 1.5190 |
| | Contrast | G4 vs G3 | -1.0253 | 4.705E-01 | 8.803E-01 | | -3.8780 | 1.8274 |
| | Mean_CI | G1 | | | | | | |
| | Mean_CI | G2 | | | | | | |
| | Mean_CI | G3 | | | | | | |
| | Mean_CI | G4 | | | | | | |
| **Eos** | **Source** | **Parameter** | **Estimate** | **Pvalue** | **fdr_p** | **OverallP** | **DIfM_LCL** | **DiffM_UCL** |
| | Intercept | Intercept | -1.7395 | 2.901E-06 | 7.253E-06 | | | |
| | Group | Group G2 | 0.0398 | 9.404E-01 | 9.404E-01 | **1.417E-09** | | |
| | Group | Group G3 | 1.0765 | 1.038E-01 | 1.483E-01 | | | |
| | Group | Group G4 | 3.5831 | **6.800E-10** | **3.400E-09** | | | |
| | Group | Group G1 | 0.0000 | | | | | |
| | Contrast | G2 vs G1 | 0.0398 | 9.404E-01 | 9.404E-01 | | -1.0330 | 1.1126 |
| | Contrast | G3 vs G1 | 1.0765 | 1.038E-01 | 1.483E-01 | | -0.2320 | 2.3849 |
| | Contrast | G4 vs G1 | 3.5831 | **6.800E-10** | **3.400E-09** | | 2.7160 | 4.4502 |
| | Contrast | G3 vs G2 | 1.0367 | 1.512E-01 | 1.890E-01 | | -0.3976 | 2.4710 |
| | Contrast | G4 vs G2 | 3.5433 | **5.671E-08** | **1.890E-07** | | 2.4958 | 4.5908 |
| | Contrast | G4 vs G3 | 2.5066 | **3.589E-04** | **7.178E-04** | | 1.2189 | 3.7944 |
| | Mean_CI | G1 | | | | | | |
| | Mean_CI | G2 | | | | | | |
| | Mean_CI | G3 | | | | | | |
| | Mean_CI | G4 | | | | | | |
| **EosPercent** | **Source** | **Parameter** | **Estimate** | **Pvalue** | **fdr_p** | **OverallP** | **DIfM_LCL** | **DiffM_UCL** |
| | Intercept | Intercept | 0.4349 | 5.078E-02 | 1.245E-01 | | | |
| | Group | Group G2 | 0.2329 | 5.258E-01 | 7.511E-01 | **1.822E-02** | | |
| | Group | Group G3 | 0.0905 | 8.393E-01 | 8.393E-01 | | | |

| | | | | | | | | |
|---|---|---|---|---|---|---|---|---|
| | Group | Group G4 | 0.9306 | **3.176E-03** | **1.588E-02** | | | |
| | Group | Group G1 | 0.0000 | | | | | |
| | Contrast | G2 vs G1 | 0.2329 | 5.258E-01 | 7.511E-01 | | -0.5048 | 0.9706 |
| | Contrast | G3 vs G1 | 0.0905 | 8.393E-01 | 8.393E-01 | | -0.8092 | 0.9903 |
| | Contrast | G4 vs G1 | 0.9306 | **3.176E-03** | **1.588E-02** | | 0.3343 | 1.5269 |
| | Contrast | G3 vs G2 | -0.1424 | 7.712E-01 | 8.393E-01 | | -1.1287 | 0.8439 |
| | Contrast | G4 vs G2 | 0.6977 | 5.722E-02 | 1.245E-01 | | -0.0226 | 1.4180 |
| | Contrast | G4 vs G3 | 0.8401 | 6.227E-02 | 1.245E-01 | | -0.0454 | 1.7256 |
| | Mean_CI | G1 | | | | | | |
| | Mean_CI | G2 | | | | | | |
| | Mean_CI | G3 | | | | | | |
| | Mean_CI | G4 | | | | | | |
| **Mon** | **Source** | **Parameter** | **Estimate** | **Pvalue** | **fdr_p** | **OverallP** | **DIfM_LCL** | **DiffM_UCL** |
| | Intercept | Intercept | -0.9476 | 3.187E-03 | 6.374E-03 | | | |
| | Group | Group G2 | 0.0692 | 8.920E-01 | 8.920E-01 | **3.248E-12** | | |
| | Group | Group G3 | 1.5673 | **1.564E-02** | **2.235E-02** | | | |
| | Group | Group G4 | 4.3409 | **1.719E-12** | **8.595E-12** | | | |
| | Group | Group G1 | 0.0000 | | | | | |
| | Contrast | G2 vs G1 | 0.0692 | 8.920E-01 | 8.920E-01 | | -0.9575 | 1.0959 |
| | Contrast | G3 vs G1 | 1.5673 | **1.564E-02** | **2.235E-02** | | 0.3151 | 2.8194 |
| | Contrast | G4 vs G1 | 4.3409 | **1.719E-12** | **8.595E-12** | | 3.5110 | 5.1707 |
| | Contrast | G3 vs G2 | 1.4981 | **3.331E-02** | **4.164E-02** | | 0.1255 | 2.8707 |
| | Contrast | G4 vs G2 | 4.2717 | **3.256E-10** | **1.085E-09** | | 3.2693 | 5.2741 |
| | Contrast | G4 vs G3 | 2.7736 | **5.853E-05** | **1.463E-04** | | 1.5412 | 4.0060 |
| | Mean_CI | G1 | | | | | | |
| | Mean_CI | G2 | | | | | | |
| | Mean_CI | G3 | | | | | | |
| | Mean_CI | G4 | | | | | | |
| **MonPercent** | **Source** | **Parameter** | **Estimate** | **Pvalue** | **fdr_p** | **OverallP** | **DIfM_LCL** | **DiffM_UCL** |
| | Intercept | Intercept | 1.2551 | 7.100E-08 | 3.601E-07 | | | |
| | Group | Group G2 | 0.2517 | 4.259E-01 | 4.259E-01 | **7.114E-07** | | |
| | Group | Group G3 | 0.6012 | 1.235E-01 | 1.765E-01 | | | |
| | Group | Group G4 | 1.6794 | **1.080E-07** | **3.601E-07** | | | |
| | Group | Group G1 | 0.0000 | | | | | |
| | Contrast | G2 vs G1 | 0.2517 | 4.259E-01 | 4.259E-01 | | -0.3825 | 0.8858 |
| | Contrast | G3 vs G1 | 0.6012 | 1.235E-01 | 1.765E-01 | | -0.1722 | 1.3746 |
| | Contrast | G4 vs G1 | 1.6794 | **1.080E-07** | **3.601E-07** | | 1.1669 | 2.1920 |
| | Contrast | G3 vs G2 | 0.3496 | 4.083E-01 | 4.259E-01 | | -0.4983 | 1.1974 |
| | Contrast | G4 vs G2 | 1.4278 | **4.183E-05** | **1.046E-04** | | 0.8086 | 2.0469 |
| | Contrast | G4 vs G3 | 1.0782 | **6.819E-03** | **1.364E-02** | | 0.3170 | 1.8394 |
| | Mean_CI | G1 | | | | | | |
| | Mean_CI | G2 | | | | | | |
| | Mean_CI | G3 | | | | | | |
| | Mean_CI | G4 | | | | | | |

| Lym | Source | Parameter | Estimate | Pvalue | fdr_p | OverallP | DIfM_LCL | DiffM_UCL |
|---|---|---|---|---|---|---|---|---|
| | Intercept | Intercept | 2.1523 | 7.898E-13 | 7.898E-12 | | | |
| | Group | Group G2 | -0.4256 | 2.095E-01 | 2.328E-01 | **1.908E-02** | | |
| | Group | Group G3 | 0.8939 | **3.436E-02** | 6.873E-02 | | | |
| | Group | Group G4 | -0.3441 | 2.094E-01 | 2.328E-01 | | | |
| | Group | Group G1 | 0.0000 | | | | | |
| | Contrast | G2 vs G1 | -0.4256 | 2.095E-01 | 2.328E-01 | | -1.1014 | 0.2502 |
| | Contrast | G3 vs G1 | 0.8939 | **3.436E-02** | 6.873E-02 | | 0.0697 | 1.7182 |
| | Contrast | G4 vs G1 | -0.3441 | 2.094E-01 | 2.328E-01 | | -0.8903 | 0.2022 |
| | Contrast | G3 vs G2 | 1.3195 | **5.423E-03** | **1.808E-02** | | 0.4160 | 2.2231 |
| | Contrast | G4 vs G2 | 0.0815 | 8.035E-01 | 8.035E-01 | | -0.5783 | 0.7414 |
| | Contrast | G4 vs G3 | -1.2380 | **3.826E-03** | **1.808E-02** | | -2.0492 | -0.4268 |
| | Mean_CI | G1 | | | | | | |
| | Mean_CI | G2 | | | | | | |
| | Mean_CI | G3 | | | | | | |
| | Mean_CI | G4 | | | | | | |
| **LymPercent** | **Source** | **Parameter** | **Estimate** | **Pvalue** | **fdr_p** | **OverallP** | **DIfM_LCL** | **DiffM_UCL** |
| | Intercept | Intercept | 4.3507 | 8.231E-21 | 8.231E-20 | | | |
| | Group | Group G2 | -0.2451 | 5.072E-01 | 7.245E-01 | **6.233E-12** | | |
| | Group | Group G3 | -0.0740 | 8.692E-01 | 8.692E-01 | | | |
| | Group | Group G4 | -3.0198 | **4.834E-12** | **1.611E-11** | | | |
| | Group | Group G1 | 0.0000 | | | | | |
| | Contrast | G2 vs G1 | -0.2451 | 5.072E-01 | 7.245E-01 | | -0.9875 | 0.4974 |
| | Contrast | G3 vs G1 | -0.0740 | 8.692E-01 | 8.692E-01 | | -0.9795 | 0.8315 |
| | Contrast | G4 vs G1 | -3.0198 | **4.834E-12** | **1.611E-11** | | -3.6199 | -2.4197 |
| | Contrast | G3 vs G2 | 0.1711 | 7.285E-01 | 8.692E-01 | | -0.8215 | 1.1637 |
| | Contrast | G4 vs G2 | -2.7747 | **3.980E-09** | **9.950E-09** | | -3.4996 | -2.0498 |
| | Contrast | G4 vs G3 | -2.9458 | **9.056E-08** | **1.811E-07** | | -3.8370 | -2.0547 |
| | Mean_CI | G1 | | | | | | |
| | Mean_CI | G2 | | | | | | |
| | Mean_CI | G3 | | | | | | |
| | Mean_CI | G4 | | | | | | |
| **MCHC** | **Source** | **Parameter** | **Estimate** | **Pvalue** | **fdr_p** | **OverallP** | **DIfM_LCL** | **DiffM_UCL** |
| | Intercept | Intercept | 3.4800 | 1.830E-54 | 1.830E-53 | | | |
| | Group | Group G2 | 0.0086 | 7.821E-01 | 8.234E-01 | **2.442E-02** | | |
| | Group | Group G3 | -0.0084 | 8.234E-01 | 8.234E-01 | | | |
| | Group | Group G4 | 0.0708 | **7.275E-03** | **2.425E-02** | | | |
| | Group | Group G1 | 0.0000 | | | | | |
| | Contrast | G2 vs G1 | 0.0086 | 7.821E-01 | 8.234E-01 | | -0.0538 | 0.0709 |
| | Contrast | G3 vs G1 | -0.0084 | 8.234E-01 | 8.234E-01 | | -0.0845 | 0.0676 |
| | Contrast | G4 vs G1 | 0.0708 | **7.275E-03** | **2.425E-02** | | 0.0204 | 0.1211 |
| | Contrast | G3 vs G2 | -0.0170 | 6.817E-01 | 8.234E-01 | | -0.1003 | 0.0664 |
| | Contrast | G4 vs G2 | 0.0622 | **4.550E-02** | 9.099E-02 | | 0.0013 | 0.1231 |
| | Contrast | G4 vs G3 | 0.0792 | **3.874E-02** | 9.099E-02 | | 0.0043 | 0.1540 |
| | Mean_CI | G1 | | | | | | |

| | | | | | | | | |
|---|---|---|---|---|---|---|---|---|
| | Mean_CI | G2 | | | | | | |
| | Mean_CI | G3 | | | | | | |
| | Mean_CI | G4 | | | | | | |
| **MCH** | **Source** | **Parameter** | **Estimate** | **Pvalue** | **fdr_p** | **OverallP** | **DIfM_LCL** | **DiffM_UCL** |
| | Intercept | Intercept | 2.6930 | 1.888E-42 | 1.888E-41 | | | |
| | Group | Group G2 | 0.1297 | **1.849E-02** | **2.311E-02** | **4.123E-09** | | |
| | Group | Group G3 | -0.0720 | 2.684E-01 | 2.684E-01 | | | |
| | Group | Group G4 | 0.3330 | **3.157E-09** | **1.052E-08** | | | |
| | Group | Group G1 | 0.0000 | | | | | |
| | Contrast | G2 vs G1 | 0.1297 | **1.849E-02** | **2.311E-02** | | 0.0231 | 0.2362 |
| | Contrast | G3 vs G1 | -0.0720 | 2.684E-01 | 2.684E-01 | | -0.2019 | 0.0580 |
| | Contrast | G4 vs G1 | 0.3330 | **3.157E-09** | **1.052E-08** | | 0.2469 | 0.4191 |
| | Contrast | G3 vs G2 | -0.2017 | **6.846E-03** | **1.141E-02** | | -0.3441 | -0.0592 |
| | Contrast | G4 vs G2 | 0.2033 | **3.421E-04** | **6.841E-04** | | 0.0993 | 0.3074 |
| | Contrast | G4 vs G3 | 0.4050 | **2.109E-07** | **5.273E-07** | | 0.2771 | 0.5329 |
| | Mean_CI | G1 | | | | | | |
| | Mean_CI | G2 | | | | | | |
| | Mean_CI | G3 | | | | | | |
| | Mean_CI | G4 | | | | | | |
| **MCV** | **Source** | **Parameter** | **Estimate** | **Pvalue** | **fdr_p** | **OverallP** | **DIfM_LCL** | **DiffM_UCL** |
| | Intercept | Intercept | 3.8182 | 6.005E-55 | 6.005E-54 | | | |
| | Group | Group G2 | 0.1196 | **8.124E-04** | **1.016E-03** | **7.243E-12** | | |
| | Group | Group G3 | -0.0644 | 1.148E-01 | 1.148E-01 | | | |
| | Group | Group G4 | 0.2629 | **9.251E-12** | **3.084E-11** | | | |
| | Group | Group G1 | 0.0000 | | | | | |
| | Contrast | G2 vs G1 | 0.1196 | **8.124E-04** | **1.016E-03** | | 0.0534 | 0.1859 |
| | Contrast | G3 vs G1 | -0.0644 | 1.148E-01 | 1.148E-01 | | -0.1452 | 0.0164 |
| | Contrast | G4 vs G1 | 0.2629 | **9.251E-12** | **3.084E-11** | | 0.2094 | 0.3165 |
| | Contrast | G3 vs G2 | -0.1840 | **1.660E-04** | **2.766E-04** | | -0.2726 | -0.0954 |
| | Contrast | G4 vs G2 | 0.1433 | **7.254E-05** | **1.451E-04** | | 0.0786 | 0.2080 |
| | Contrast | G4 vs G3 | 0.3273 | **7.483E-10** | **1.871E-09** | | 0.2478 | 0.4069 |
| | Mean_CI | G1 | | | | | | |
| | Mean_CI | G2 | | | | | | |
| | Mean_CI | G3 | | | | | | |
| | Mean_CI | G4 | | | | | | |
| **HGB** | **Source** | **Parameter** | **Estimate** | **Pvalue** | **fdr_p** | **OverallP** | **DIfM_LCL** | **DiffM_UCL** |
| | Intercept | Intercept | 2.6492 | 2.192E-32 | 2.192E-31 | | | |
| | Group | Group G2 | -0.1192 | 2.445E-01 | 3.493E-01 | **2.842E-07** | | |
| | Group | Group G3 | -0.1018 | 4.127E-01 | 4.586E-01 | | | |
| | Group | Group G4 | -0.5654 | **4.502E-08** | **1.501E-07** | | | |
| | Group | Group G1 | 0.0000 | | | | | |
| | Contrast | G2 vs G1 | -0.1192 | 2.445E-01 | 3.493E-01 | | -0.3237 | 0.0853 |
| | Contrast | G3 vs G1 | -0.1018 | 4.127E-01 | 4.586E-01 | | -0.3513 | 0.1476 |
| | Contrast | G4 vs G1 | -0.5654 | **4.502E-08** | **1.501E-07** | | -0.7307 | -0.4001 |

| | | | | | | | | |
|---|---|---|---|---|---|---|---|---|
| | Contrast | G3 vs G2 | 0.0174 | 8.980E-01 | 8.980E-01 | | -0.2560 | 0.2908 |
| | Contrast | G4 vs G2 | -0.4462 | **6.444E-05** | **1.611E-04** | | -0.6459 | -0.2465 |
| | Contrast | G4 vs G3 | -0.4636 | **5.030E-04** | **1.006E-03** | | -0.7091 | -0.2181 |
| | Mean_CI | G1 | | | | | | |
| | Mean_CI | G2 | | | | | | |
| | Mean_CI | G3 | | | | | | |
| | Mean_CI | G4 | | | | | | |
| **MPV** | **Source** | **Parameter** | **Estimate** | **Pvalue** | **fdr_p** | **OverallP** | **DIfM_LCL** | **DiffM_UCL** |
| | Intercept | Intercept | 1.6572 | 9.667E-33 | 9.667E-32 | | | |
| | Group | Group G2 | 0.1288 | **4.363E-02** | 6.233E-02 | **5.953E-05** | | |
| | Group | Group G3 | 0.0954 | 2.120E-01 | 2.356E-01 | | | |
| | Group | Group G4 | 0.2719 | **3.903E-06** | **1.301E-05** | | | |
| | Group | Group G1 | 0.0000 | | | | | |
| | Contrast | G2 vs G1 | 0.1288 | **4.363E-02** | 6.233E-02 | | 0.0039 | 0.2537 |
| | Contrast | G3 vs G1 | 0.0954 | 2.120E-01 | 2.356E-01 | | -0.0569 | 0.2478 |
| | Contrast | G4 vs G1 | 0.2719 | **3.903E-06** | **1.301E-05** | | 0.1709 | 0.3728 |
| | Contrast | G3 vs G2 | -0.0334 | 6.872E-01 | 6.872E-01 | | -0.2004 | 0.1336 |
| | Contrast | G4 vs G2 | 0.1431 | **2.282E-02** | **4.564E-02** | | 0.0211 | 0.2650 |
| | Contrast | G4 vs G3 | 0.1765 | **2.240E-02** | **4.564E-02** | | 0.0265 | 0.3264 |
| | Mean_CI | G1 | | | | | | |
| | Mean_CI | G2 | | | | | | |
| | Mean_CI | G3 | | | | | | |
| | Mean_CI | G4 | | | | | | |

| ubstances | | | | | | | | Group Names |
|---|---|---|---|---|---|---|---|---|
| **GMean** | **GMean_LCL** | **GMean_UCL** | **Mean_O** | **Std_O** | **Min_O** | **Max_O** | | MxCre only |
| | | | | | | | | NPM1 |
| | | | | | | | | PTPN11 |
| | | | | | | | | PTPN11/NPM1 |
| 11.0957 | 7.6506 | 16.0923 | 12.2285 | 6.4820 | 5.35 | 31.48 | | |
| 9.2641 | 5.5817 | 15.3758 | 9.5914 | 2.5738 | 5.74 | 12.22 | | |
| 29.2209 | 14.9491 | 57.1179 | 31.1000 | 11.7732 | 16.65 | 43.52 | | |
| 158.5416 | 112.1582 | 224.1071 | 206.5667 | 128.1393 | 10.04 | 444.26 | | |
| **GMean** | **GMean_LCL** | **GMean_UCL** | **Mean_O** | **Std_O** | **Min_O** | **Max_O** | | |
| 1124.8214 | 760.4988 | 1663.6754 | 1161.0769 | 292.9185 | 702 | 1647 | | |
| 577.5838 | 338.8165 | 984.6130 | 756.1429 | 429.8742 | 78 | 1270 | | |
| 826.7594 | 408.2576 | 1674.2641 | 850.0000 | 237.4447 | 667 | 1169 | | |
| 95.1913 | 66.1226 | 137.0392 | 177.2667 | 363.7178 | 32 | 1488 | | |
| **GMean** | **GMean_LCL** | **GMean_UCL** | **Mean_O** | **Std_O** | **Min_O** | **Max_O** | | |

| | | | | | | |
|---|---|---|---|---|---|---|
| 9.5702 | 8.1113 | 11.2914 | 9.6038 | 0.8201 | 7.57 | 10.67 |
| 7.4649 | 5.9585 | 9.3521 | 7.5057 | 0.8603 | 6.57 | 8.7 |
| 9.3120 | 6.9112 | 12.5468 | 9.3625 | 1.1388 | 8.1 | 10.79 |
| 3.8950 | 3.3392 | 4.5434 | 4.2493 | 1.7862 | 1.65 | 8.3 |
| **GMean** | **GMean_LCL** | **GMean_UCL** | **Mean_O** | **Std_O** | **Min_O** | **Max_O** |
| 1.5515 | 0.9935 | 2.4228 | 1.8523 | 1.6079 | 0.87 | 7.05 |
| 1.7896 | 0.9749 | 3.2851 | 1.9300 | 0.7300 | 0.76 | 3 |
| 4.7571 | 2.1300 | 10.6245 | 4.8150 | 0.8770 | 4.09 | 5.86 |
| 93.5639 | 61.7881 | 141.6811 | 127.7707 | 79.9099 | 2.56 | 299.3 |
| **GMean** | **GMean_LCL** | **GMean_UCL** | **Mean_O** | **Std_O** | **Min_O** | **Max_O** |
| 13.9974 | 11.7713 | 16.6446 | 14.3615 | 3.5269 | 9.7 | 22.4 |
| 19.3063 | 15.2471 | 24.4462 | 20.6286 | 8.2819 | 10.8 | 36 |
| 16.3330 | 11.9524 | 22.3191 | 17.1250 | 6.0566 | 11.1 | 24.6 |
| 59.0424 | 50.2498 | 69.3734 | 61.3267 | 14.8326 | 25.5 | 80.3 |
| **GMean** | **GMean_LCL** | **GMean_UCL** | **Mean_O** | **Std_O** | **Min_O** | **Max_O** |

| | | | | | | |
|---|---|---|---|---|---|---|
| 0.0742 | 0.0164 | 0.3357 | 0.1500 | 0.2302 | 0.02 | 0.8 |
| 0.0846 | 0.0108 | 0.6621 | 0.0957 | 0.0529 | 0.03 | 0.19 |
| 0.3309 | 0.0218 | 5.0320 | 0.3650 | 0.1771 | 0.17 | 0.56 |
| 0.4546 | 0.1115 | 1.8537 | 5.8667 | 7.1935 | 0 | 24.75 |
| **GMean** | **GMean_LCL** | **GMean_UCL** | **Mean_O** | **Std_O** | **Min_O** | **Max_O** |
| 0.6411 | 0.1571 | 2.6153 | 1.1231 | 1.5520 | 0.1 | 5.8 |
| 0.8747 | 0.1287 | 5.9428 | 1.1000 | 0.8641 | 0.4 | 2.5 |
| 1.0942 | 0.0868 | 13.8005 | 1.1500 | 0.4203 | 0.7 | 1.7 |
| 0.3925 | 0.1060 | 1.4530 | 3.2733 | 3.3230 | 0 | 12.2 |
| **GMean** | **GMean_LCL** | **GMean_UCL** | **Mean_O** | **Std_O** | **Min_O** | **Max_O** |
| 0.1756 | 0.0931 | 0.3313 | 0.2592 | 0.2763 | 0.06 | 1.01 |
| 0.1827 | 0.0769 | 0.4340 | 0.2000 | 0.0891 | 0.09 | 0.33 |
| 0.5153 | 0.1641 | 1.6179 | 0.5825 | 0.2864 | 0.21 | 0.9 |
| 6.3193 | 3.5000 | 11.4096 | 16.7387 | 33.3506 | 0.13 | 134.98 |
| **GMean** | **GMean_LCL** | **GMean_UCL** | **Mean_O** | **Std_O** | **Min_O** | **Max_O** |

| | | | | | | |
|---|---|---|---|---|---|---|
| 1.5448 | 0.9984 | 2.3901 | 2.2769 | 3.0760 | 0.5 | 12.3 |
| 1.9499 | 1.0757 | 3.5344 | 2.1286 | 0.9050 | 0.9 | 3.3 |
| 1.6912 | 0.7700 | 3.7144 | 1.7250 | 0.3775 | 1.2 | 2 |
| 3.9176 | 2.6096 | 5.8814 | 6.1867 | 7.8127 | 0.9 | 32.3 |
| **GMean** | **GMean_LCL** | **GMean_UCL** | **Mean_O** | **Std_O** | **Min_O** | **Max_O** |
| 0.3877 | 0.2112 | 0.7116 | 0.6300 | 0.9968 | 0.15 | 3.9 |
| 0.4155 | 0.1816 | 0.9506 | 0.6329 | 0.8100 | 0.18 | 2.45 |
| 1.8584 | 0.6217 | 5.5550 | 2.1525 | 1.4196 | 0.95 | 4.21 |
| 29.7645 | 16.9095 | 52.3924 | 47.6640 | 36.3443 | 0.42 | 122.4 |
| **GMean** | **GMean_LCL** | **GMean_UCL** | **Mean_O** | **Std_O** | **Min_O** | **Max_O** |
| 3.5083 | 2.4109 | 5.1054 | 4.1385 | 2.8268 | 1.5 | 12.4 |
| 4.5123 | 2.7062 | 7.5237 | 8.9429 | 14.9255 | 1.5 | 42.7 |
| 6.4005 | 3.2545 | 12.5874 | 7.5000 | 5.4412 | 4.1 | 15.6 |
| 18.8137 | 13.2677 | 26.6780 | 20.6533 | 7.7471 | 4.2 | 35.4 |

| GMean | GMean_LCL | GMean_UCL | Mean_O | Std_O | Min_O | Max_O |
|---|---|---|---|---|---|---|
| 8.6046 | 5.7689 | 12.8342 | 9.3369 | 3.9519 | 4.2 | 19.1 |
| 5.6221 | 3.2604 | 9.6946 | 6.7329 | 2.8930 | 0.94 | 9.44 |
| 21.0360 | 10.2314 | 43.2508 | 23.1850 | 11.0345 | 11.23 | 34.45 |
| 6.0996 | 4.2039 | 8.8501 | 8.5267 | 7.4219 | 1.25 | 26.15 |
| GMean | GMean_LCL | GMean_UCL | Mean_O | Std_O | Min_O | Max_O |
| 77.5352 | 49.9740 | 120.2967 | 78.1000 | 9.1397 | 56.5 | 86.1 |
| 60.6826 | 33.3510 | 110.4126 | 67.2000 | 23.0635 | 16.4 | 84.5 |
| 72.0073 | 32.6204 | 158.9513 | 72.5000 | 9.6861 | 61.4 | 81.9 |
| 3.7845 | 2.5144 | 5.6963 | 8.5600 | 16.9969 | 0.7 | 68.7 |
| GMean | GMean_LCL | GMean_UCL | Mean_O | Std_O | Min_O | Max_O |
| 32.4600 | 31.2845 | 33.6796 | 32.4692 | 0.8420 | 30.4 | 33.8 |

| | | | | | | |
|---|---|---|---|---|---|---|
| 32.7391 | 31.1341 | 34.4268 | 32.7429 | 0.6024 | 31.9 | 33.5 |
| 32.1878 | 30.1170 | 34.4009 | 32.2000 | 1.0646 | 30.9 | 33.5 |
| 34.8399 | 33.6638 | 36.0570 | 35.0000 | 3.5313 | 29.6 | 43 |
| **GMean** | **GMean_LCL** | **GMean_UCL** | **Mean_O** | **Std_O** | **Min_O** | **Max_O** |
| 14.7764 | 13.8738 | 15.7377 | 14.7923 | 0.7065 | 12.6 | 15.3 |
| 16.8224 | 15.4377 | 18.3312 | 16.8286 | 0.5345 | 16.2 | 17.7 |
| 13.7502 | 12.2733 | 15.4048 | 13.7500 | 0.1732 | 13.6 | 14 |
| 20.6156 | 19.4408 | 21.8615 | 20.8933 | 3.5666 | 16.2 | 27.2 |
| **GMean** | **GMean_LCL** | **GMean_UCL** | **Mean_O** | **Std_O** | **Min_O** | **Max_O** |
| 45.5225 | 43.7725 | 47.3424 | 45.5462 | 1.5382 | 41.5 | 47.3 |
| 51.3074 | 48.6384 | 54.1229 | 51.3286 | 1.6408 | 49.5 | 54.2 |
| 42.6838 | 39.7714 | 45.8094 | 42.7000 | 1.3976 | 41 | 44.3 |
| 59.2135 | 57.0915 | 61.4143 | 59.5067 | 6.2520 | 50.8 | 73.2 |
| **GMean** | **GMean_LCL** | **GMean_UCL** | **Mean_O** | **Std_O** | **Min_O** | **Max_O** |

| | | | | | | |
|---|---|---|---|---|---|---|
| 14.1431 | 12.5315 | 15.9619 | 14.1846 | 1.1172 | 11.6 | 15.7 |
| 12.5534 | 10.6454 | 14.8035 | 12.6286 | 1.4852 | 10.6 | 14.4 |
| 12.7736 | 10.2706 | 15.8867 | 12.8500 | 1.6279 | 11 | 14.8 |
| 8.0348 | 7.1790 | 8.9927 | 8.3933 | 2.4592 | 4.1 | 13.4 |
| **GMean** | **GMean_LCL** | **GMean_UCL** | **Mean_O** | **Std_O** | **Min_O** | **Max_O** |
| 5.2448 | 4.8711 | 5.6470 | 5.2769 | 0.6340 | 4.6 | 6.5 |
| 5.9657 | 5.3942 | 6.5978 | 5.9714 | 0.3039 | 5.5 | 6.4 |
| 5.7698 | 5.0501 | 6.5920 | 5.8000 | 0.7118 | 5.2 | 6.8 |
| 6.8833 | 6.4257 | 7.3735 | 6.9733 | 1.1997 | 5.3 | 9.5 |

| |
|---|
| G1 |
| G2 |
| G3 |
| G4 |

| ANVOA | | | | | | | | |
|---|---|---|---|---|---|---|---|---|
| **ANOVA Model for Log(Cell)** | | | | | | | | |
| **Source** | **Parameter** | **Estimate** | **Pvalue** | **fdr_p** | **OverallP** | **DIfM_LCL** | **DiffM_UCL** | **GMean** |
| Intercept | Intercept | -1.8460 | 0.0019 | 0.0047 | | | | |
| Group | Group G2 | 1.0757 | 0.1416 | 0.1416 | **0.0000** | | | |
| Group | Group G3 | 2.2038 | **0.0069** | **0.0115** | | | | |
| Group | Group G4 | 3.8762 | **0.0000** | **0.0000** | | | | |
| Group | Group G1 | 0.0000 | | | | | | |
| Contrast | G2 vs G1 | 1.0757 | 0.1416 | 0.1416 | | -0.3889 | 2.5402 | |
| Contrast | G3 vs G1 | 2.2038 | **0.0069** | **0.0115** | | 0.6741 | 3.7335 | |
| Contrast | G4 vs G1 | 3.8762 | **0.0000** | **0.0000** | | 2.5271 | 5.2252 | |
| Contrast | G3 vs G2 | 1.1282 | 0.1241 | 0.1416 | | -0.3364 | 2.5928 | |
| Contrast | G4 vs G2 | 2.8005 | **0.0002** | **0.0006** | | 1.5257 | 4.0753 | |
| Contrast | G4 vs G3 | 1.6723 | **0.0175** | **0.0251** | | 0.3232 | 3.0214 | |
| Mean_CI | G1 | | | | | | | 0.1579 |
| Mean_CI | G2 | | | | | | | 0.4628 |
| Mean_CI | G3 | | | | | | | 1.4302 |
| Mean_CI | G4 | | | | | | | 7.6151 |
| **Group Names** | | | | | | | | |
| MxCre | G1 | | | | | | | |
| NPM1 | G2 | | | | | | | |
| PTPN11 | G3 | | | | | | | |
| NPM1/PTPN11 | G4 | | | | | | | |

| GMean_LCL | GMean_UCL | Mean_O | Std_O | Min_O | Max_O |
|---|---|---|---|---|---|
| 0.0535 | 0.4656 | 0.8100 | 1.540795249 | 0.02 | 3.56 |
| 0.1724 | 1.2424 | 0.7667 | 1.00205123 | 0.16 | 2.78 |
| 0.4849 | 4.2185 | 2.0620 | 1.751433698 | 0.37 | 4.18 |
| 3.4004 | 17.0537 | 8.4356 | 3.458204415 | 2.53 | 12.82 |

| ANOVA Model for Log(Cell) | | | | | | | | | |
|---|---|---|---|---|---|---|---|---|---|
| **Source** | **Parameter** | **Estimate** | **Pvalue** | **fdr_p** | **OverallP** | **DIfM_LCL** | **DiffM_UCL** | **GMean** | **GMean_LCL** |
| Intercept | Intercept | 1.4262 | 0.0000 | 0.0002 | | | | | |
| Group | Group G2 | 0.1057 | 0.7758 | 0.8273 | **0.0000** | | | | |
| Group | Group G3 | 0.1867 | 0.6308 | 0.8273 | | | | | |
| Group | Group G4 | 1.6931 | **0.0001** | **0.0002** | | | | | |
| Group | Group G1 | 0.0000 | | | | | | | |
| Contrast | G2 vs G1 | 0.1057 | 0.7758 | 0.8273 | | -0.6565 | 0.8680 | | |
| Contrast | G3 vs G1 | 0.1867 | 0.6308 | 0.8273 | | -0.6094 | 0.9829 | | |
| Contrast | G4 vs G1 | 1.6931 | **0.0001** | **0.0002** | | 0.9909 | 2.3952 | | |
| Contrast | G3 vs G2 | 0.0810 | 0.8273 | 0.8273 | | -0.6813 | 0.8433 | | |
| Contrast | G4 vs G2 | 1.5873 | **0.0001** | **0.0002** | | 0.9238 | 2.2508 | | |
| Contrast | G4 vs G3 | 1.5063 | **0.0002** | **0.0004** | | 0.8042 | 2.2085 | | |
| Mean_CI | G1 | | | | | | | 4.1628 | 2.3708 |
| Mean_CI | G2 | | | | | | | 4.6271 | 2.7677 |
| Mean_CI | G3 | | | | | | | 5.0174 | 2.8575 |
| Mean_CI | G4 | | | | | | | 22.6295 | 14.8743 |
| **Group Names** | | | | | | | | | |
| MxCre | G1 | | | | | | | | |
| NPM1 | G2 | | | | | | | | |
| PTPN11 | G3 | | | | | | | | |
| NPM1/PTP | G4 | | | | | | | | |

| GMean_UCL | Mean_O | Std_O | Min_O | Max_O |
|---|---|---|---|---|
| 7.3094 | 4.7520 | 2.8759 | 2.27 | 9.47 |
| 7.7358 | 6.6433 | 7.2420 | 2.07 | 20.99 |
| 8.8101 | 6.4140 | 5.1964 | 1.71 | 15.24 |
| 34.4279 | 22.9389 | 4.1255 | 18.58 | 31.03 |

| ANOVA Model for Log(Cell) | | | | | | | | | |
|---|---|---|---|---|---|---|---|---|---|
| **Source** | **Parameter** | **Estimate** | **Pvalue** | **fdr_p** | **OverallP** | **DIfM_LCL** | **DiffM_UCL** | **GMean** | **GMean_LCL** |
| Intercept | Intercept | -0.0451 | 0.9248 | 0.9401 | | | | | |
| Group | Group G2 | 0.0486 | 0.9401 | 0.9401 | 0.0854 | | | | |
| Group | Group G3 | 0.5030 | 0.4598 | 0.6932 | | | | | |
| Group | Group G4 | 1.3285 | **0.0349** | 0.1165 | | | | | |
| Group | Group G1 | 0.0000 | | | | | | | |
| Contrast | G2 vs G1 | 0.0486 | 0.9401 | 0.9401 | | -1.2816 | 1.3788 | | |
| Contrast | G3 vs G1 | 0.5030 | 0.4598 | 0.6932 | | -0.8863 | 1.8924 | | |
| Contrast | G4 vs G1 | 1.3285 | **0.0349** | 0.1165 | | 0.1032 | 2.5538 | | |
| Contrast | G3 vs G2 | 0.4544 | 0.4853 | 0.6932 | | -0.8758 | 1.7846 | | |
| Contrast | G4 vs G2 | 1.2799 | **0.0319** | 0.1165 | | 0.1221 | 2.4377 | | |
| Contrast | G4 vs G3 | 0.8255 | 0.1758 | 0.4395 | | -0.3998 | 2.0507 | | |
| Mean_CI | G1 | | | | | | | 0.9559 | 0.3579 |
| Mean_CI | G2 | | | | | | | 1.0035 | 0.4093 |
| Mean_CI | G3 | | | | | | | 1.5808 | 0.5919 |
| Mean_CI | G4 | | | | | | | 3.6089 | 1.7352 |
| **Group Names** | | | | | | | | | |
| MxCre | G1 | | | | | | | | |
| NPM1 | G2 | | | | | | | | |
| PTPN11 | G3 | | | | | | | | |
| NPM1/PTP | G4 | | | | | | | | |

| GMean_UCL | Mean_O | Std_O | Min_O | Max_O |
|---|---|---|---|---|
| 2.5531 | 1.1800 | 0.6835 | 0.24 | 2.06 |
| 2.4604 | 2.4467 | 4.2194 | 0.33 | 11.02 |
| 4.2221 | 1.7680 | 0.8903 | 0.85 | 2.92 |
| 7.5055 | 5.1389 | 3.0750 | 0.22 | 9.08 |

| ANOVA Model for Log(Cell) | | | | | | | | | |
|---|---|---|---|---|---|---|---|---|---|
| **Source** | **Parameter** | **Estimate** | **Pvalue** | **fdr_p** | **OverallP** | **DIfM_LCL** | **DiffM_UCL** | **GMean** | **GMean_LCL** |
| Intercept | Intercept | 0.1353 | 0.8127 | 0.8127 | 0.2737 | | | | |
| Group | Group G2 | 0.3917 | 0.6134 | 0.6816 | | | | | |
| Group | Group G3 | 1.5476 | 0.0659 | 0.3297 | | | | | |
| Group | Group G4 | 0.7589 | 0.2930 | 0.4883 | | | | | |
| Group | Group G1 | 0.0000 | | | | | | | |
| Contrast | G2 vs G1 | 0.3917 | 0.6134 | 0.6816 | | -1.1967 | 1.9800 | | |
| Contrast | G3 vs G1 | 1.5476 | 0.0659 | 0.3297 | | -0.1114 | 3.2066 | | |
| Contrast | G4 vs G1 | 0.7589 | 0.2930 | 0.4883 | | -0.7042 | 2.2220 | | |
| Contrast | G3 vs G2 | 1.1559 | 0.1451 | 0.4836 | | -0.4325 | 2.7443 | | |
| Contrast | G4 vs G2 | 0.3672 | 0.5866 | 0.6816 | | -1.0153 | 1.7497 | | |
| Contrast | G4 vs G3 | -0.7887 | 0.2749 | 0.4883 | | -2.2518 | 0.6744 | | |
| Mean_CI | G1 | | | | | | | 1.1449 | 0.3543 |
| Mean_CI | G2 | | | | | | | 1.6939 | 0.5805 |
| Mean_CI | G3 | | | | | | | 5.3813 | 1.6650 |
| Mean_CI | G4 | | | | | | | 2.4454 | 1.0200 |
| **Group Names** | | | | | | | | | |
| MxCre | G1 | | | | | | | | |
| NPM1 | G2 | | | | | | | | |
| PTPN11 | G3 | | | | | | | | |
| NPM1/PTP | G4 | | | | | | | | |

| | | | | |
|---|---|---|---|---|
| GMean_UCL | Mean_O | Std_O | Min_O | Max_O |
| 3.7003 | 2.2780 | 2.2238 | 0.22 | 4.85 |
| 4.9426 | 2.9400 | 3.5903 | 0.26 | 10.07 |
| 17.3921 | 7.7600 | 6.1779 | 0.85 | 17.76 |
| 5.8624 | 3.8889 | 2.7572 | 0.33 | 7.64 |

| ANOVA Model for Log(Cell) | | | | | | | | |
|---|---|---|---|---|---|---|---|---|
| **Source** | **Parameter** | **Estimate** | **Pvalue** | **fdr_p** | **OverallP** | **DIfM_LCL** | **DiffM_UCL** | **GMean** |
| Intercept | Intercept | -1.3884 | 0.0116 | 0.0232 | | | | |
| Group | Group G2 | -0.0621 | 0.9281 | 0.9281 | **0.0001** | | | |
| Group | Group G3 | 0.8231 | 0.2593 | 0.3242 | | | | |
| Group | Group G4 | 2.8991 | **0.0001** | **0.0005** | | | | |
| Group | Group G1 | 0.0000 | | | | | | |
| Contrast | G2 vs G1 | -0.0621 | 0.9281 | 0.9281 | | -1.4757 | 1.3516 | |
| Contrast | G3 vs G1 | 0.8231 | 0.2593 | 0.3242 | | -0.6534 | 2.2996 | |
| Contrast | G4 vs G1 | 2.8991 | **0.0001** | **0.0005** | | 1.5969 | 4.2012 | |
| Contrast | G3 vs G2 | 0.8852 | 0.2070 | 0.3242 | | -0.5285 | 2.2988 | |
| Contrast | G4 vs G2 | 2.9612 | **0.0001** | **0.0005** | | 1.7307 | 4.1916 | |
| Contrast | G4 vs G3 | 2.0760 | **0.0033** | **0.0082** | | 0.7738 | 3.3781 | |
| Mean_CI | G1 | | | | | | | 0.2495 |
| Mean_CI | G2 | | | | | | | 0.2345 |
| Mean_CI | G3 | | | | | | | 0.5682 |
| Mean_CI | G4 | | | | | | | 4.5299 |
| **Group Names** | | | | | | | | |
| MxCre | G1 | | | | | | | |
| NPM1 | G2 | | | | | | | |
| PTPN11 | G3 | | | | | | | |
| NPM1/PTP | G4 | | | | | | | |

| | | | | | |
|---|---|---|---|---|---|
| **GMean_LCL** | **GMean_UCL** | **Mean_O** | **Std_O** | **Min_O** | **Max_O** |
| 0.0878 | 0.7087 | 0.4920 | 0.5557 | 0.06 | 1.28 |
| 0.0904 | 0.6081 | 0.4400 | 0.4760 | 0.03 | 1.28 |
| 0.2000 | 1.6141 | 0.6560 | 0.4216 | 0.36 | 1.34 |
| 2.0803 | 9.8639 | 6.8444 | 6.2190 | 0.92 | 19.24 |

| | | | | | | | | |
|---|---|---|---|---|---|---|---|---|
| | | | | ANOVA Model for Log(Cell) for 3 Substa | | | | |
| **Substance** | **Source** | **Parameter** | **Estimate** | **Pvalue** | **fdr_p** | **OverallP** | **DIfM_LCL** | **DiffM_UCL** |
| **GMP** | Intercept | Intercept | 3.3104 | 0.0000 | 0.0000 | | | |
| | Group | Group G2 | -0.0765 | 0.7967 | 0.7987 | 0.3740 | | |
| | Group | Group G3 | -0.1583 | 0.6230 | 0.7987 | | | |
| | Group | Group G4 | -0.4642 | 0.1178 | 0.3928 | | | |
| | Group | Group G1 | 0.0000 | | | | | |
| | Contrast | G2 vs G1 | -0.0765 | 0.7967 | 0.7987 | | -0.7092 | 0.5562 |
| | Contrast | G3 vs G1 | -0.1583 | 0.6230 | 0.7987 | | -0.8417 | 0.5251 |
| | Contrast | G4 vs G1 | -0.4642 | 0.1178 | 0.3928 | | -1.0644 | 0.1361 |
| | Contrast | G3 vs G2 | -0.0818 | 0.7987 | 0.7987 | | -0.7652 | 0.6017 |
| | Contrast | G4 vs G2 | -0.3876 | 0.1848 | 0.4619 | | -0.9879 | 0.2126 |
| | Contrast | G4 vs G3 | -0.3059 | 0.3279 | 0.6559 | | -0.9594 | 0.3476 |
| | Mean_CI | G1 | | | | | | |
| | Mean_CI | G2 | | | | | | |
| | Mean_CI | G3 | | | | | | |
| | Mean_CI | G4 | | | | | | |
| **MEP** | **Source** | **Parameter** | **Estimate** | **Pvalue** | **fdr_p** | **OverallP** | **DIfM_LCL** | **DiffM_UCL** |
| | Intercept | Intercept | 2.8312 | 0.0000 | 0.0000 | | | |
| | Group | Group G2 | 0.1752 | 0.5892 | 0.6547 | 0.5105 | | |
| | Group | Group G3 | 0.4462 | 0.2153 | 0.4332 | | | |
| | Group | Group G4 | 0.3906 | 0.2166 | 0.4332 | | | |
| | Group | Group G1 | 0.0000 | | | | | |
| | Contrast | G2 vs G1 | 0.1752 | 0.5892 | 0.6547 | | -0.5127 | 0.8631 |
| | Contrast | G3 vs G1 | 0.4462 | 0.2153 | 0.4332 | | -0.2969 | 1.1892 |
| | Contrast | G4 vs G1 | 0.3906 | 0.2166 | 0.4332 | | -0.2620 | 1.0432 |
| | Contrast | G3 vs G2 | 0.2710 | 0.4422 | 0.6547 | | -0.4720 | 1.0140 |
| | Contrast | G4 vs G2 | 0.2155 | 0.4857 | 0.6547 | | -0.4371 | 0.8681 |
| | Contrast | G4 vs G3 | -0.0555 | 0.8676 | 0.8676 | | -0.7660 | 0.6549 |
| | Mean_CI | G1 | | | | | | |
| | Mean_CI | G2 | | | | | | |
| | Mean_CI | G3 | | | | | | |
| | Mean_CI | G4 | | | | | | |
| **CMP** | **Source** | **Parameter** | **Estimate** | **Pvalue** | **fdr_p** | **OverallP** | **DIfM_LCL** | **DiffM_UCL** |
| | Intercept | Intercept | 3.8325 | 0.0000 | 0.0000 | | | |
| | Group | Group G2 | -0.0821 | 0.8518 | 0.8518 | 0.1134 | | |
| | Group | Group G3 | -0.3550 | 0.4596 | 0.6566 | | | |
| | Group | Group G4 | -0.9790 | **0.0336** | 0.1120 | | | |
| | Group | Group G1 | 0.0000 | | | | | |
| | Contrast | G2 vs G1 | -0.0821 | 0.8518 | 0.8518 | | -1.0195 | 0.8553 |
| | Contrast | G3 vs G1 | -0.3550 | 0.4596 | 0.6566 | | -1.3676 | 0.6575 |
| | Contrast | G4 vs G1 | -0.9790 | **0.0336** | 0.1120 | | -1.8683 | -0.0897 |
| | Contrast | G3 vs G2 | -0.2729 | 0.5679 | 0.7099 | | -1.2854 | 0.7396 |
| | Contrast | G4 vs G2 | -0.8969 | **0.0484** | 0.1209 | | -1.7862 | -0.0075 |
| | Contrast | G4 vs G3 | -0.6240 | 0.1856 | 0.3712 | | -1.5921 | 0.3442 |

| | | |
|---|---|---|
| | Mean_CI | G1 |
| | Mean_CI | G2 |
| | Mean_CI | G3 |
| | Mean_CI | G4 |
| **Group Names** | | |
| MxCre | G1 | |
| NPM1 | G2 | |
| PTPN11 | G3 | |
| NPM1/PTPN11 | G4 | |

| ances | | | | | | |
|---|---|---|---|---|---|---|
| **GMean** | **GMean_LCL** | **GMean_UCL** | **Mean_O** | **Std_O** | **Min_O** | **Max_O** |
| 27.3970 | 17.5143 | 42.8560 | 27.6025 | 3.8616 | 22.86 | 32.24 |
| 25.3790 | 16.2243 | 39.6994 | 26.5350 | 9.2150 | 16.81 | 39.01 |
| 23.3861 | 13.9505 | 39.2037 | 23.5000 | 2.8736 | 21.14 | 26.7 |
| 17.2235 | 11.5432 | 25.6990 | 19.8560 | 11.0594 | 6.63 | 36.51 |
| **GMean** | **GMean_LCL** | **GMean_UCL** | **Mean_O** | **Std_O** | **Min_O** | **Max_O** |
| 16.9666 | 10.4315 | 27.5958 | 17.1775 | 3.1136 | 13.59 | 21.14 |
| 20.2147 | 12.4285 | 32.8788 | 22.3075 | 11.6722 | 11.8 | 38.35 |
| 26.5070 | 15.1158 | 46.4826 | 26.6833 | 3.7143 | 22.77 | 30.16 |
| 25.0753 | 16.2293 | 38.7430 | 29.0980 | 18.0524 | 12.99 | 58.03 |
| **GMean** | **GMean_LCL** | **GMean_UCL** | **Mean_O** | **Std_O** | **Min_O** | **Max_O** |

| | | | | | | |
|---|---|---|---|---|---|---|
| 46.1778 | 23.7991 | 89.5995 | 46.1875 | 1.0937 | 45.2 | 47.38 |
| 42.5366 | 21.9225 | 82.5345 | 42.9850 | 7.1334 | 34.43 | 51.76 |
| 32.3774 | 15.0603 | 69.6062 | 32.8400 | 6.5090 | 25.5 | 37.91 |
| 17.3485 | 9.5892 | 31.3866 | 23.2780 | 14.5544 | 3 | 40.94 |

| ANOVA Model for Log(Cell) for 7 Substance | | | | | | | | | |
|---|---|---|---|---|---|---|---|---|---|
| **Substance** | **Source** | **Parameter** | **Estimate** | **Pvalue** | **fdr_p** | **OverallP** | **DIfM_LCL** | **DiffM_UCL** | **GMean** |
| **HSC** | Intercept | Intercept | -1.3884 | 0.0116 | 0.0232 | | | | |
| | Group | Group G2 | -0.0621 | 0.9281 | 0.9281 | **0.0001** | | | |
| | Group | Group G3 | 0.8231 | 0.2593 | 0.3242 | | | | |
| | Group | Group G4 | 2.8991 | **0.0001** | **0.0005** | | | | |
| | Group | Group G1 | 0.0000 | | | | | | |
| | Contrast | G2 vs G1 | -0.0621 | 0.9281 | 0.9281 | | -1.4757 | 1.3516 | |
| | Contrast | G3 vs G1 | 0.8231 | 0.2593 | 0.3242 | | -0.6534 | 2.2996 | |
| | Contrast | G4 vs G1 | 2.8991 | **0.0001** | **0.0005** | | 1.5969 | 4.2012 | |
| | Contrast | G3 vs G2 | 0.8852 | 0.2070 | 0.3242 | | -0.5285 | 2.2988 | |
| | Contrast | G4 vs G2 | 2.9612 | **0.0001** | **0.0005** | | 1.7307 | 4.1916 | |
| | Contrast | G4 vs G3 | 2.0760 | **0.0033** | **0.0082** | | 0.7738 | 3.3781 | |
| | Mean_CI | G1 | | | | | | | 0.2495 |
| | Mean_CI | G2 | | | | | | | 0.2345 |
| | Mean_CI | G3 | | | | | | | 0.5682 |
| | Mean_CI | G4 | | | | | | | 4.5299 |
| **MPP1** | **Source** | **Parameter** | **Estimate** | **Pvalue** | **fdr_p** | **OverallP** | **DIfM_LCL** | **DiffM_UCL** | **GMean** |
| | Intercept | Intercept | 0.3414 | 0.6241 | 0.7801 | | | | |
| | Group | Group G2 | 0.2216 | 0.8214 | 0.8214 | **0.0009** | | | |
| | Group | Group G3 | -0.6481 | 0.5437 | 0.7767 | | | | |
| | Group | Group G4 | -4.2561 | **0.0005** | **0.0018** | | | | |
| | Group | Group G1 | 0.0000 | | | | | | |
| | Contrast | G2 vs G1 | 0.2216 | 0.8214 | 0.8214 | | -1.8701 | 2.3132 | |
| | Contrast | G3 vs G1 | -0.6481 | 0.5437 | 0.7767 | | -2.9073 | 1.6111 | |
| | Contrast | G4 vs G1 | -4.2561 | **0.0005** | **0.0018** | | -6.2404 | -2.2718 | |
| | Contrast | G3 vs G2 | -0.8696 | 0.4180 | 0.7767 | | -3.1288 | 1.3896 | |
| | Contrast | G4 vs G2 | -4.4777 | **0.0004** | **0.0018** | | -6.4619 | -2.4934 | |
| | Contrast | G4 vs G3 | -3.6080 | **0.0034** | **0.0085** | | -5.7682 | -1.4478 | |
| | Mean_CI | G1 | | | | | | | 1.4070 |
| | Mean_CI | G2 | | | | | | | 1.7559 |
| | Mean_CI | G3 | | | | | | | 0.7359 |
| | Mean_CI | G4 | | | | | | | 0.0199 |
| **MPP2** | **Source** | **Parameter** | **Estimate** | **Pvalue** | **fdr_p** | **OverallP** | **DIfM_LCL** | **DiffM_UCL** | **GMean** |
| | Intercept | Intercept | -1.1042 | 0.2413 | 0.3196 | | | | |
| | Group | Group G2 | 1.3153 | 0.3196 | 0.3196 | **0.0203** | | | |
| | Group | Group G3 | -1.5938 | 0.2667 | 0.3196 | | | | |
| | Group | Group G4 | -2.9905 | **0.0285** | 0.0950 | | | | |
| | Group | Group G1 | 0.0000 | | | | | | |
| | Contrast | G2 vs G1 | 1.3153 | 0.3196 | 0.3196 | | -1.4446 | 4.0752 | |
| | Contrast | G3 vs G1 | -1.5938 | 0.2667 | 0.3196 | | -4.5748 | 1.3873 | |
| | Contrast | G4 vs G1 | -2.9905 | **0.0285** | 0.0950 | | -5.6088 | -0.3722 | |
| | Contrast | G3 vs G2 | -2.9091 | 0.0549 | 0.1373 | | -5.8901 | 0.0720 | |
| | Contrast | G4 vs G2 | -4.3058 | **0.0038** | **0.0376** | | -6.9241 | -1.6875 | |
| | Contrast | G4 vs G3 | -1.3967 | 0.3067 | 0.3196 | | -4.2471 | 1.4537 | |

| | | | | | | | | | |
|---|---|---|---|---|---|---|---|---|---|
| | Mean_CI | G1 | | | | | | | 0.3315 |
| | Mean_CI | G2 | | | | | | | 1.2350 |
| | Mean_CI | G3 | | | | | | | 0.0673 |
| | Mean_CI | G4 | | | | | | | 0.0167 |
| **MPP3** | **Source** | **Parameter** | **Estimate** | **Pvalue** | **fdr_p** | **OverallP** | **DIfM_LCL** | **DiffM_UCL** | **GMean** |
| | Intercept | Intercept | 3.5849 | 0.0000 | 0.0000 | | | | |
| | Group | Group G2 | 0.3164 | 0.1153 | 0.2883 | 0.1946 | | | |
| | Group | Group G3 | 0.0559 | 0.7861 | 0.7861 | | | | |
| | Group | Group G4 | -0.0850 | 0.6396 | 0.7861 | | | | |
| | Group | Group G1 | 0.0000 | | | | | | |
| | Contrast | G2 vs G1 | 0.3164 | 0.1153 | 0.2883 | | -0.0897 | 0.7226 | |
| | Contrast | G3 vs G1 | 0.0559 | 0.7861 | 0.7861 | | -0.3828 | 0.4945 | |
| | Contrast | G4 vs G1 | -0.0850 | 0.6396 | 0.7861 | | -0.4702 | 0.3003 | |
| | Contrast | G3 vs G2 | -0.2606 | 0.2199 | 0.4399 | | -0.6992 | 0.1781 | |
| | Contrast | G4 vs G2 | -0.4014 | **0.0424** | 0.2122 | | -0.7867 | -0.0161 | |
| | Contrast | G4 vs G3 | -0.1408 | 0.4785 | 0.7861 | | -0.5603 | 0.2786 | |
| | Mean_CI | G1 | | | | | | | 36.0509 |
| | Mean_CI | G2 | | | | | | | 49.4704 |
| | Mean_CI | G3 | | | | | | | 38.1229 |
| | Mean_CI | G4 | | | | | | | 33.1147 |
| **MPP4** | **Source** | **Parameter** | **Estimate** | **Pvalue** | **fdr_p** | **OverallP** | **DIfM_LCL** | **DiffM_UCL** | **GMean** |
| | Intercept | Intercept | 3.1925 | 0.0001 | 0.0015 | | | | |
| | Group | Group G2 | -1.2601 | 0.1534 | 0.2224 | 0.1303 | | | |
| | Group | Group G3 | 0.0927 | 0.9191 | 0.9191 | | | | |
| | Group | Group G4 | -1.6485 | 0.0574 | 0.1602 | | | | |
| | Group | Group G1 | 0.0000 | | | | | | |
| | Contrast | G2 vs G1 | -1.2601 | **0.1534** | 0.2224 | | -3.0613 | 0.5412 | |
| | Contrast | G3 vs G1 | 0.0927 | 0.9191 | 0.9191 | | -1.8529 | 2.0382 | |
| | Contrast | G4 vs G1 | -1.6485 | 0.0574 | 0.1602 | | -3.3573 | 0.0603 | |
| | Contrast | G3 vs G2 | 1.3528 | 0.1557 | 0.2224 | | -0.5928 | 3.2983 | |
| | Contrast | G4 vs G2 | -0.3884 | 0.6294 | 0.7867 | | -2.0972 | 1.3204 | |
| | Contrast | G4 vs G3 | -1.7411 | 0.0641 | 0.1602 | | -3.6015 | 0.1192 | |
| | Mean_CI | G1 | | | | | | | 24.3481 |
| | Mean_CI | G2 | | | | | | | 6.9059 |
| | Mean_CI | G3 | | | | | | | 26.7125 |
| | Mean_CI | G4 | | | | | | | 4.6832 |
| **MPP5** | **Source** | **Parameter** | **Estimate** | **Pvalue** | **fdr_p** | **OverallP** | **DIfM_LCL** | **DiffM_UCL** | **GMean** |
| | Intercept | Intercept | 2.5689 | 0.0000 | 0.0000 | | | | |
| | Group | Group G2 | -0.2278 | 0.4797 | 0.5330 | **0.0025** | | | |
| | Group | Group G3 | -0.3421 | 0.3304 | 0.4720 | | | | |
| | Group | Group G4 | 1.0138 | **0.0051** | **0.0101** | | | | |
| | Group | Group G1 | 0.0000 | | | | | | |
| | Contrast | G2 vs G1 | -0.2278 | 0.4797 | 0.5330 | | -0.9080 | 0.4525 | |
| | Contrast | G3 vs G1 | -0.3421 | 0.3304 | 0.4720 | | -1.0768 | 0.3926 | |

| | | | | | | | | | |
|---|---|---|---|---|---|---|---|---|---|
| | Contrast | G4 vs G1 | 1.0138 | **0.0051** | **0.0101** | | 0.3685 | 1.6591 | |
| | Contrast | G3 vs G2 | -0.1143 | 0.7404 | 0.7404 | | -0.8491 | 0.6204 | |
| | Contrast | G4 vs G2 | 1.2416 | **0.0012** | **0.0042** | | 0.5962 | 1.8869 | |
| | Contrast | G4 vs G3 | 1.3559 | **0.0012** | **0.0042** | | 0.6533 | 2.0584 | |
| | Mean_CI | G1 | | | | | | | 13.0510 |
| | Mean_CI | G2 | | | | | | | 10.3927 |
| | Mean_CI | G3 | | | | | | | 9.2699 |
| | Mean_CI | G4 | | | | | | | 35.9691 |

| **MPP6** | **Source** | **Parameter** | **Estimate** | **Pvalue** | **fdr_p** | **OverallP** | **DIfM_LCL** | **DiffM_UCL** | **GMean** |
|---|---|---|---|---|---|---|---|---|---|
| | Intercept | Intercept | -0.3039 | 0.4451 | 0.5003 | | | | |
| | Group | Group G2 | -0.7258 | 0.2072 | 0.2960 | 0.0179 | | | |
| | Group | Group G3 | 0.4086 | 0.5003 | 0.5003 | | | | |
| | Group | Group G4 | 1.2243 | **0.0354** | 0.1178 | | | | |
| | Group | Group G1 | 0.0000 | | | | | | |
| | Contrast | G2 vs G1 | -0.7258 | 0.2072 | 0.2960 | | -1.9118 | 0.4602 | |
| | Contrast | G3 vs G1 | 0.4086 | 0.5003 | 0.5003 | | -0.8724 | 1.6897 | |
| | Contrast | G4 vs G1 | 1.2243 | **0.0354** | 0.1178 | | 0.0991 | 2.3495 | |
| | Contrast | G3 vs G2 | 1.1344 | 0.0777 | 0.1942 | | -0.1466 | 2.4155 | |
| | Contrast | G4 vs G2 | 1.9501 | **0.0026** | **0.0264** | | 0.8249 | 3.0753 | |
| | Contrast | G4 vs G3 | 0.8157 | 0.1724 | 0.2960 | | -0.4092 | 2.0406 | |
| | Mean_CI | G1 | | | | | | | 0.7379 |
| | Mean_CI | G2 | | | | | | | 0.3571 |
| | Mean_CI | G3 | | | | | | | 1.1104 |
| | Mean_CI | G4 | | | | | | | 2.5104 |

| s | | | | | | | | |
|---|---|---|---|---|---|---|---|---|
| **GMean_LCL** | **GMean_UCL** | **Mean_O** | **Std_O** | **Min_O** | **Max_O** | | **Group Names** | |
| | | | | | | | MxCre | G1 |
| | | | | | | | NPM1 | G2 |
| | | | | | | | PTPN11 | G3 |
| | | | | | | | NPM1/PTPN11 | G4 |
| 0.0878 | 0.7087 | 0.4920 | 0.5557 | 0.06 | 1.28 | | | |
| 0.0904 | 0.6081 | 0.4400 | 0.4760 | 0.03 | 1.28 | | | |
| 0.2000 | 1.6141 | 0.6560 | 0.4216 | 0.36 | 1.34 | | | |
| 2.0803 | 9.8639 | 6.8444 | 6.2190 | 0.92 | 19.24 | | | |
| **GMean_LCL** | **GMean_UCL** | **Mean_O** | **Std_O** | **Min_O** | **Max_O** | | | |
| 0.3206 | 6.1745 | 1.4850 | 0.6097 | 1.01 | 2.38 | | | |
| 0.4001 | 7.7059 | 1.8525 | 0.6817 | 1.15 | 2.55 | | | |
| 0.1334 | 4.0599 | 0.9867 | 0.7550 | 0.23 | 1.74 | | | |
| 0.0053 | 0.0749 | 0.1000 | 0.1904 | 0 | 0.44 | | | |
| **GMean_LCL** | **GMean_UCL** | **Mean_O** | **Std_O** | **Min_O** | **Max_O** | | | |

| | | | | | |
|---|---|---|---|---|---|
| 0.0471 | 2.3334 | 0.3500 | 0.1273 | 0.19 | 0.49 |
| 0.1754 | 8.6941 | 1.2750 | 0.3816 | 0.88 | 1.79 |
| 0.0071 | 0.6411 | 0.3900 | 0.3900 | 0 | 0.78 |
| 0.0029 | 0.0954 | 0.0300 | 0.0265 | 0 | 0.07 |
| **GMean_LCL** | **GMean_UCL** | **Mean_O** | **Std_O** | **Min_O** | **Max_O** |
| 27.0521 | 48.0431 | 36.3650 | 5.4126 | 29.39 | 41.9 |
| 37.1219 | 65.9266 | 52.1050 | 20.7314 | 36.24 | 82.3 |
| 27.3639 | 53.1123 | 38.3833 | 5.4831 | 33.01 | 43.97 |
| 25.6137 | 42.8124 | 34.1600 | 8.8950 | 20.9 | 43.72 |
| **GMean_LCL** | **GMean_UCL** | **Mean_O** | **Std_O** | **Min_O** | **Max_O** |
| 6.8127 | 87.0184 | 26.5250 | 12.3928 | 15.89 | 40.86 |
| 1.9323 | 24.6811 | 12.9275 | 11.6681 | 0.76 | 26.64 |
| 6.1376 | 116.2603 | 26.7500 | 1.7432 | 24.82 | 28.21 |
| 1.4990 | 14.6316 | 9.4740 | 11.8315 | 0.78 | 29.84 |
| **GMean_LCL** | **GMean_UCL** | **Mean_O** | **Std_O** | **Min_O** | **Max_O** |

| | | | | | |
|---|---|---|---|---|---|
| 8.0677 | 21.1127 | 13.5200 | 4.0845 | 9.17 | 18.21 |
| 6.4244 | 16.8122 | 11.1875 | 4.4539 | 5.4 | 15.85 |
| 5.3194 | 16.1543 | 9.3200 | 1.2020 | 8.2 | 10.59 |
| 23.3930 | 55.3062 | 40.6840 | 20.2715 | 17.55 | 63.56 |
| **GMean_LCL** | **GMean_UCL** | **Mean_O** | **Std_O** | **Min_O** | **Max_O** |
| 0.3190 | 1.7070 | 0.7775 | 0.2775 | 0.45 | 1.05 |
| 0.1544 | 0.8261 | 0.5850 | 0.5708 | 0.06 | 1.39 |
| 0.4216 | 2.9245 | 1.3200 | 0.9677 | 0.6 | 2.42 |
| 1.1857 | 5.3150 | 2.6580 | 1.1188 | 1.79 | 4.61 |
