## Supplemental Data Table 2 for "*PTPN11* Mutation Clonal Hierarchy in Acute Myeloid Leukemia"

| | p_val | avg_log2FC | pct.1 | pct.2 | p_val_adj |
|---|---|---|---|---|---|
| CD11b | 0.000167 | -23.0054 | 0.952 | 0.961 | 7.52E-03 |
| CD117 | 1.30E-10 | -8.09248 | 0.99 | 0.998 | 5.85E-09 |
| CD34 | 6.24E-16 | -7.16729 | 0.952 | 0.996 | 2.81E-14 |
| CD13 | 1.18E-05 | -4.73694 | 0.952 | 0.978 | 5.30E-04 |
| CD123 | 1.63E-05 | -4.28044 | 0.981 | 0.996 | 7.35E-04 |
| CD45RO | 3.64E-06 | 20.11137 | 0.924 | 0.832 | 1.64E-04 |
| CD2 | 2.48E-08 | 21.35332 | 0.895 | 0.856 | 1.12E-06 |
| CD3 | 2.89E-08 | 27.78945 | 0.914 | 0.909 | 1.30E-06 |
| CD5 | 3.83E-15 | 61.1806 | 0.943 | 0.866 | 1.72E-13 |

| | p_val | avg_log2FC | pct.1 | pct.2 | p_val_adj |
|---|---|---|---|---|---|
| FcÎµRI | 4.94E-07 | -19.0985 | 0.811 | 0.763 | 2.22E-05 |
| CD163 | 4.18E-09 | -8.80566 | 0.822 | 0.863 | 1.88E-07 |
| CD304 | 1.56E-06 | -6.44639 | 0.795 | 0.848 | 7.02E-05 |
| CD13 | 1.88E-53 | -5.50123 | 0.84 | 0.947 | 8.45E-52 |
| CD11b | 4.55E-41 | -5.03943 | 0.834 | 0.919 | 2.05E-39 |
| CD11c | 1.04E-46 | -4.24348 | 0.819 | 0.926 | 4.69E-45 |
| CD64 | 1.01E-62 | -3.90375 | 0.854 | 0.934 | 4.56E-61 |
| CD14 | 6.97E-57 | -3.82401 | 0.848 | 0.924 | 3.14E-55 |
| CD117 | 2.88E-31 | -3.69545 | 0.831 | 0.888 | 1.29E-29 |
| CD71 | 2.22E-27 | -3.01269 | 0.915 | 0.932 | 1.00E-25 |
| CD33 | 8.26E-94 | -2.45791 | 0.825 | 0.955 | 3.71E-92 |
| CD141 | 4.81E-55 | -1.91909 | 0.9 | 0.959 | 2.17E-53 |
| CD123 | 7.56E-19 | -0.93159 | 0.851 | 0.902 | 3.40E-17 |
| HLA.DR | 4.50E-24 | -0.33235 | 0.856 | 0.908 | 2.02E-22 |
| CD1c | 2.71E-09 | 0.307402 | 0.781 | 0.745 | 1.22E-07 |
| CD16 | 6.93E-05 | 0.344416 | 0.838 | 0.813 | 0.003117 |
| CD45RA | 9.94E-07 | 0.780933 | 0.94 | 0.958 | 4.47E-05 |
| CD30 | 6.94E-08 | 1.510063 | 0.832 | 0.805 | 3.12E-06 |
| CD303 | 1.28E-08 | 3.043252 | 0.805 | 0.848 | 5.76E-07 |
| CD38 | 3.22E-15 | 3.191929 | 0.923 | 0.963 | 1.45E-13 |
| CD45 | 6.03E-39 | 5.585566 | 0.981 | 0.95 | 2.71E-37 |
| CD4 | 1.40E-10 | 6.641649 | 0.926 | 0.948 | 6.29E-09 |
| CD69 | 0.000932 | 6.739503 | 0.86 | 0.874 | 0.041926 |
| CD7 | 1.07E-81 | 6.875381 | 0.926 | 0.83 | 4.82E-80 |
| CD8 | 5.98E-43 | 8.127403 | 0.843 | 0.76 | 2.69E-41 |
| CD22 | 4.20E-05 | 8.311878 | 0.787 | 0.772 | 0.00189 |
| CD2 | 3.69E-169 | 8.629881 | 0.927 | 0.751 | 1.66E-167 |
| CD45RO | 3.99E-29 | 8.681505 | 0.849 | 0.757 | 1.80E-27 |
| CD25 | 1.31E-17 | 9.860483 | 0.826 | 0.755 | 5.90E-16 |
| CD5 | 1.00E-152 | 11.74603 | 0.92 | 0.751 | 4.50E-151 |
| CD3 | 1.48E-151 | 12.16819 | 0.922 | 0.765 | 6.65E-150 |
| CD19 | 2.27E-17 | 14.16559 | 0.817 | 0.761 | 1.02E-15 |
| CD62L | 1.67E-63 | 16.6225 | 0.892 | 0.791 | 7.50E-62 |

| | p_val | avg_log2FC | pct.1 | pct.2 | p_val_adj |
|---|---|---|---|---|---|
| CD62P | 6.47E-10 | -74.034 | 0.837 | 0.896 | 2.91E-08 |
| CD10 | 7.92E-11 | -29.0203 | 0.881 | 0.93 | 3.56E-09 |
| CD33 | 5.79E-20 | -28.3327 | 0.856 | 0.933 | 2.61E-18 |
| CD123 | 1.50E-07 | -17.4379 | 0.886 | 0.911 | 6.76E-06 |
| CD25 | 0.000219 | -14.9389 | 0.846 | 0.808 | 0.00987 |
| CD13 | 0.000525 | -14.8578 | 0.851 | 0.88 | 0.023633 |
| CD45RO | 0.000331 | -12.0434 | 0.881 | 0.857 | 0.014908 |
| HLA.DR | 8.52E-09 | -11.312 | 0.863 | 0.893 | 3.83E-07 |
| CD38 | 1.24E-09 | -6.37813 | 0.942 | 0.96 | 5.60E-08 |
| CD34 | 1.49E-63 | -5.94478 | 0.944 | 0.982 | 6.69E-62 |
| CD117 | 1.62E-71 | -4.9546 | 0.984 | 0.987 | 7.28E-70 |
| CD56 | 6.26E-47 | -4.38174 | 0.961 | 0.977 | 2.82E-45 |
| CD49d | 1.89E-37 | -4.33614 | 0.967 | 0.991 | 8.52E-36 |
| CD69 | 2.88E-06 | -2.37784 | 0.954 | 0.968 | 0.00013 |
| CD71 | 6.14E-59 | -1.90988 | 0.965 | 0.993 | 2.76E-57 |
| CD45 | 2.06E-40 | -1.55982 | 0.954 | 0.911 | 9.26E-39 |
| CD141 | 2.27E-41 | -1.50958 | 0.991 | 0.995 | 1.02E-39 |
| CD2 | 5.69E-86 | -1.25106 | 0.968 | 0.895 | 2.56E-84 |
| CD303 | 7.54E-13 | 0.583664 | 0.937 | 0.95 | 3.39E-11 |
| CD64 | 2.95E-05 | 1.346678 | 0.921 | 0.947 | 0.001328 |
| CD44 | 1.70E-08 | 1.550645 | 0.954 | 0.955 | 7.64E-07 |
| CD4 | 4.72E-44 | 6.325615 | 0.94 | 0.914 | 2.13E-42 |
| CD7 | 4.74E-26 | 6.527313 | 0.977 | 0.983 | 2.13E-24 |
| CD62L | 9.99E-18 | 8.779748 | 0.907 | 0.862 | 4.49E-16 |
| CD3 | 4.05E-107 | 14.52209 | 0.956 | 0.851 | 1.82E-105 |
| CD5 | 9.35E-132 | 15.84183 | 0.954 | 0.834 | 4.21E-130 |
| CD8 | 1.05E-18 | 15.96206 | 0.897 | 0.875 | 4.73E-17 |

| | p_val | avg_log2FC | pct.1 | pct.2 | p_val_adj |
|---|---|---|---|---|---|
| CD33 | 0.000841 | 24.59283 | 0.989 | 1 | 0.037837 |
| HLA.DR | 0.000374 | 15.57278 | 0.999 | 1 | 0.01681 |
| CD11c | 0.001242 | 14.92564 | 0.997 | 1 | 0.055909 |
| CD11b | 0.000832 | 12.15693 | 0.993 | 1 | 0.037437 |
| CD64 | 0.000134 | 8.447968 | 0.999 | 1 | 0.006019 |
| CD4 | 0.000457 | 5.070246 | 1 | 1 | 0.020545 |
| CD25 | 0.000369 | 3.449765 | 0.975 | 1 | 0.016623 |

| | p_val | avg_log2FC | pct.1 | pct.2 | p_val_adj |
|---|---|---|---|---|---|
| FcÎµRI | 1.03E-05 | -27.2696 | 0.927 | 0.871 | 0.000466 |
| CD34 | 3.96E-06 | -23.6064 | 0.912 | 0.911 | 0.000178 |
| CD13 | 3.87E-06 | -18.3381 | 0.901 | 0.89 | 0.000174 |
| CD163 | 8.93E-13 | -17.395 | 0.914 | 0.925 | 4.02E-11 |
| CD123 | 2.61E-99 | -17.3069 | 0.947 | 0.972 | 1.17E-97 |
| CD1c | 5.34E-05 | -14.558 | 0.897 | 0.866 | 0.002404 |
| CD117 | 1.20E-19 | -13.7346 | 0.934 | 0.93 | 5.40E-18 |
| CD64 | 8.07E-125 | -10.9498 | 0.933 | 0.978 | 3.63E-123 |
| CD14 | 7.22E-26 | -10.2885 | 0.941 | 0.954 | 3.25E-24 |
| CD33 | 4.84E-199 | -9.50785 | 0.937 | 0.997 | 2.18E-197 |
| CD11b | 5.52E-65 | -7.93074 | 0.951 | 0.986 | 2.48E-63 |
| CD11c | 8.41E-139 | -7.11434 | 0.946 | 0.991 | 3.78E-137 |
| CD141 | 3.59E-53 | -3.04884 | 0.951 | 0.98 | 1.62E-51 |
| CD304 | 6.12E-39 | -2.00392 | 0.88 | 0.964 | 2.75E-37 |
| CD44 | 2.34E-20 | -1.66709 | 0.987 | 0.995 | 1.05E-18 |
| CD49d | 2.07E-19 | -1.04268 | 0.99 | 0.995 | 9.31E-18 |
| HLA.DR | 1.95E-76 | 0.376206 | 0.961 | 0.989 | 8.76E-75 |
| CD45 | 1.25E-09 | 0.906023 | 0.994 | 0.991 | 5.61E-08 |
| CD71 | 2.24E-12 | 1.037756 | 0.972 | 0.954 | 1.01E-10 |
| CD45RA | 9.01E-21 | 3.370141 | 0.983 | 0.951 | 4.05E-19 |
| CD62L | 2.05E-11 | 6.51996 | 0.936 | 0.894 | 9.22E-10 |
| CD83 | 0.000299 | 8.635459 | 0.948 | 0.897 | 0.013456 |
| CD90 | 4.28E-13 | 9.336799 | 0.932 | 0.845 | 1.93E-11 |
| CD16 | 5.01E-11 | 11.51974 | 0.971 | 0.93 | 2.26E-09 |
| CD4 | 3.73E-15 | 11.89679 | 0.934 | 0.986 | 1.68E-13 |
| CD30 | 1.04E-13 | 12.34151 | 0.939 | 0.87 | 4.68E-12 |
| CD69 | 2.81E-19 | 17.05909 | 0.953 | 0.957 | 1.26E-17 |
| CD38 | 1.52E-31 | 18.45349 | 0.985 | 0.995 | 6.83E-30 |
| CD7 | 5.74E-133 | 19.94096 | 0.983 | 0.875 | 2.58E-131 |
| CD8 | 6.75E-54 | 20.54982 | 0.95 | 0.874 | 3.04E-52 |
| CD22 | 1.98E-08 | 23.24748 | 0.907 | 0.867 | 8.91E-07 |
| CD2 | 1.05E-160 | 25.38616 | 0.982 | 0.826 | 4.72E-159 |
| CD5 | 6.00E-125 | 29.91522 | 0.965 | 0.821 | 2.70E-123 |
| CD56 | 2.04E-22 | 34.94687 | 0.936 | 0.89 | 9.20E-21 |
| CD3 | 4.51E-115 | 41.64502 | 0.962 | 0.852 | 2.03E-113 |
| CD25 | 1.28E-14 | 45.15368 | 0.891 | 0.817 | 5.76E-13 |
| CD19 | 1.71E-39 | 61.20265 | 0.932 | 0.812 | 7.69E-38 |

| | p_val | avg_log2FC | pct.1 | pct.2 | p_val_adj |
|---|---|---|---|---|---|
| CD62L | 4.08E-12 | 13.24725 | 0.78 | 0.938 | 1.84E-10 |
| CD25 | 9.82E-13 | 9.596002 | 0.786 | 0.943 | 4.42E-11 |
| CD45RO | 4.01E-23 | 7.075288 | 0.772 | 0.948 | 1.81E-21 |
| CD163 | 3.59E-07 | 5.793242 | 0.894 | 0.933 | 1.61E-05 |
| CD11b | 8.24E-24 | 5.442222 | 0.943 | 0.928 | 3.71E-22 |
| CD64 | 3.93E-42 | 5.079895 | 0.958 | 0.907 | 1.77E-40 |
| CD11c | 1.76E-37 | 4.918335 | 0.949 | 0.881 | 7.94E-36 |
| CD10 | 6.26E-05 | 4.305757 | 0.818 | 0.943 | 0.002818 |
| CD33 | 4.37E-49 | 3.63216 | 0.959 | 0.881 | 1.97E-47 |
| CD117 | 5.74E-13 | 2.874162 | 0.885 | 0.912 | 2.58E-11 |
| CD123 | 1.10E-32 | 2.860073 | 0.947 | 0.928 | 4.97E-31 |
| CD141 | 8.14E-24 | 2.681039 | 0.957 | 0.928 | 3.66E-22 |
| CD71 | 3.61E-15 | 2.599195 | 0.947 | 0.954 | 1.63E-13 |
| CD90 | 9.59E-05 | 2.509122 | 0.813 | 0.923 | 0.004316 |
| CD38 | 2.19E-23 | 0.961883 | 0.962 | 0.954 | 9.87E-22 |
| CD44 | 0.000537 | 0.92215 | 0.971 | 0.964 | 0.024175 |
| FcεRI | 1.13E-10 | 0.50214 | 0.792 | 0.938 | 5.09E-09 |

| | p_val | avg_log2FC | pct.1 | pct.2 | p_val_adj |
|---|---|---|---|---|---|
| CD14 | 3.42E-06 | -36.2868 | 0.845 | 0.764 | 0.000154 |
| CD11c | 5.33E-05 | -25.9681 | 0.845 | 0.881 | 0.002398 |
| CD62P | 0.000132 | -24.6421 | 0.866 | 0.794 | 0.00594 |
| CD16 | 2.85E-07 | -19.523 | 0.918 | 0.812 | 1.28E-05 |
| CD33 | 5.42E-14 | -19.4701 | 0.866 | 0.906 | 2.44E-12 |
| CD123 | 1.32E-29 | -18.2416 | 0.918 | 0.961 | 5.96E-28 |
| CD117 | 9.85E-25 | -9.84673 | 0.701 | 0.928 | 4.43E-23 |
| CD38 | 4.10E-15 | -9.15324 | 0.948 | 0.97 | 1.85E-13 |
| CD22 | 2.25E-05 | -8.6683 | 0.876 | 0.788 | 0.001013 |
| CD69 | 1.86E-12 | -7.14529 | 0.979 | 0.96 | 8.37E-11 |
| CD3 | 7.93E-37 | -6.51036 | 0.959 | 0.774 | 3.57E-35 |
| CD303 | 0.000139 | -3.63486 | 0.876 | 0.807 | 0.006265 |
| CD25 | 6.75E-06 | -0.40012 | 0.876 | 0.781 | 0.000304 |
| CD45 | 7.71E-13 | 1.919366 | 0.959 | 0.876 | 3.47E-11 |
| CD71 | 8.47E-10 | 1.986341 | 0.639 | 0.9 | 3.81E-08 |
| CD4 | 4.39E-21 | 2.90286 | 0.948 | 0.862 | 1.98E-19 |
| CD62L | 8.18E-08 | 3.594007 | 0.918 | 0.828 | 3.68E-06 |
| CD7 | 3.75E-40 | 7.675216 | 1 | 0.858 | 1.69E-38 |
| CD45RA | 2.54E-17 | 11.24771 | 0.938 | 0.8 | 1.14E-15 |
| CD2 | 5.81E-36 | 13.00682 | 0.948 | 0.791 | 2.61E-34 |
| CD5 | 2.73E-39 | 20.02831 | 0.969 | 0.774 | 1.23E-37 |
| CD19 | 6.25E-06 | 25.45253 | 0.866 | 0.799 | 0.000281 |
| CD1c | 4.47E-06 | 28.21082 | 0.835 | 0.76 | 0.000201 |
| CD8 | 2.09E-07 | 37.2605 | 0.876 | 0.809 | 9.39E-06 |

| | p_val | avg_log2FC | pct.1 | pct.2 | p_val_adj |
|---|---|---|---|---|---|
| CD62P | 6.12E-07 | -39.8394 | 0.785 | 0.876 | 2.75E-05 |
| CD11b | 4.07E-09 | -37.5724 | 0.817 | 0.864 | 1.83E-07 |
| CD64 | 6.58E-05 | -30.8926 | 0.817 | 0.89 | 0.002961 |
| CD123 | 4.30E-08 | -29.3085 | 0.839 | 0.966 | 1.94E-06 |
| CD25 | 1.47E-12 | -28.2486 | 0.806 | 0.864 | 6.61E-11 |
| CD45RO | 1.74E-10 | -26.2617 | 0.828 | 0.874 | 7.81E-09 |
| CD138 | 6.02E-09 | -22.4284 | 0.204 | 0.214 | 2.71E-07 |
| CD56 | 2.69E-13 | -20.9303 | 0.828 | 0.867 | 1.21E-11 |
| CD33 | 2.31E-18 | -20.7529 | 0.129 | 0.704 | 1.04E-16 |
| CD16 | 3.93E-08 | -20.4214 | 0.828 | 0.875 | 1.77E-06 |
| CD304 | 0.000226 | -19.4162 | 0.806 | 0.895 | 0.010188 |
| CD117 | 4.31E-16 | -17.7255 | 0.882 | 0.977 | 1.94E-14 |
| CD14 | 2.36E-12 | -17.6917 | 0.796 | 0.854 | 1.06E-10 |
| CD62L | 3.35E-11 | -15.5449 | 0.817 | 0.876 | 1.51E-09 |
| CD30 | 2.64E-13 | -13.6949 | 0.86 | 0.881 | 1.19E-11 |
| FcεRI | 8.63E-13 | -10.3142 | 0.828 | 0.854 | 3.88E-11 |
| CD303 | 1.58E-09 | -9.08395 | 0.796 | 0.864 | 7.11E-08 |
| CD163 | 4.04E-13 | -8.58966 | 0.828 | 0.866 | 1.82E-11 |
| CD10 | 1.10E-09 | -6.17826 | 0.817 | 0.874 | 4.93E-08 |
| CD34 | 2.23E-09 | -4.4356 | 0.796 | 0.861 | 1.00E-07 |
| CD71 | 1.71E-09 | 2.768412 | 0.441 | 0.772 | 7.71E-08 |
| CD90 | 6.17E-13 | 4.127591 | 0.828 | 0.859 | 2.78E-11 |
| CD45 | 1.79E-13 | 4.324924 | 0.935 | 0.944 | 8.07E-12 |
| CD141 | 0.000137 | 4.996263 | 0.892 | 0.978 | 0.006184 |
| HLA.DR | 0.000978 | 5.582545 | 0.892 | 0.924 | 0.043992 |
| CD8 | 6.16E-18 | 14.36452 | 0.849 | 0.857 | 2.77E-16 |
| CD19 | 2.01E-09 | 14.51262 | 0.871 | 0.922 | 9.05E-08 |
| CD44 | 3.27E-08 | 15.24334 | 0.925 | 0.94 | 1.47E-06 |
| CD38 | 2.35E-07 | 15.73419 | 0.892 | 0.975 | 1.06E-05 |
| CD4 | 1.97E-14 | 17.79284 | 0.892 | 0.914 | 8.86E-13 |
| CD22 | 1.02E-21 | 18.82814 | 0.871 | 0.862 | 4.58E-20 |
| CD7 | 1.05E-20 | 28.47069 | 0.925 | 0.904 | 4.71E-19 |
| CD2 | 1.12E-29 | 31.16601 | 0.914 | 0.848 | 5.04E-28 |
| CD1c | 1.09E-16 | 33.55945 | 0.839 | 0.856 | 4.92E-15 |
| CD45RA | 2.03E-35 | 40.19806 | 0.925 | 0.858 | 9.12E-34 |
| CD3 | 8.27E-22 | 48.21079 | 0.86 | 0.856 | 3.72E-20 |
| CD5 | 2.12E-26 | 102.3484 | 0.882 | 0.847 | 9.55E-25 |

| | p_val | avg_log2FC | pct.1 | pct.2 | p_val_adj |
|---|---|---|---|---|---|
| CD64 | 0.000205 | -10.4402 | 0.75 | 0.922 | 0.00923 |
| CD123 | 0.000793 | -9.48972 | 0.85 | 0.952 | 0.035693 |
| CD11c | 9.97E-05 | -9.44083 | 0.7 | 0.892 | 0.004485 |
| CD33 | 0.00014 | -9.06785 | 0.75 | 0.919 | 0.006311 |
| CD3 | 0.000229 | 18.81837 | 0.9 | 0.763 | 0.010327 |
| CD2 | 0.000576 | 38.79702 | 0.85 | 0.736 | 0.025924 |

| | p_val | avg_log2FC | pct.1 | pct.2 | p_val_adj |
|---|---|---|---|---|---|
| CD123 | 1.74E-11 | -10.0523 | 0.729 | 0.909 | 7.85E-10 |
| CD33 | 5.71E-12 | -6.88177 | 0.721 | 0.887 | 2.57E-10 |
| CD71 | 3.15E-13 | -5.67415 | 0.357 | 0.666 | 1.42E-11 |
| CD117 | 4.25E-08 | -5.64679 | 0.286 | 0.537 | 1.91E-06 |
| CD141 | 0.000396 | -4.91489 | 0.279 | 0.483 | 0.017803 |
| CD56 | 8.65E-07 | -4.42915 | 0.271 | 0.537 | 3.89E-05 |
| CD30 | 0.000342 | -3.65609 | 0.15 | 0.103 | 0.015412 |
| HLA.DR | 7.07E-06 | -3.14679 | 0.729 | 0.711 | 0.000318 |
| CD10 | 0.000107 | 1.029422 | 0.65 | 0.655 | 0.00482 |
| CD16 | 0.000129 | 1.498728 | 0.686 | 0.693 | 0.005805 |
| CD45RA | 2.06E-10 | 1.51833 | 0.393 | 0.203 | 9.25E-09 |
| CD45RO | 1.62E-09 | 3.503569 | 0.314 | 0.172 | 7.29E-08 |
| CD45 | 1.28E-11 | 6.349643 | 0.907 | 0.843 | 5.76E-10 |
| CD5 | 1.47E-39 | 6.527358 | 0.907 | 0.683 | 6.62E-38 |
| CD4 | 1.25E-13 | 6.809093 | 0.807 | 0.675 | 5.64E-12 |
| CD8 | 2.58E-18 | 11.46548 | 0.443 | 0.067 | 1.16E-16 |
| CD25 | 3.10E-06 | 11.68458 | 0.843 | 0.798 | 0.00014 |
| CD3 | 3.38E-38 | 12.46019 | 0.929 | 0.717 | 1.52E-36 |
| CD7 | 2.36E-26 | 13.99228 | 0.864 | 0.709 | 1.06E-24 |
| CD69 | 2.91E-07 | 16.21624 | 0.743 | 0.717 | 1.31E-05 |
| CD22 | 4.68E-05 | 19.69975 | 0.793 | 0.756 | 0.002105 |
| CD19 | 3.25E-07 | 20.73578 | 0.85 | 0.821 | 1.46E-05 |
| CD2 | 4.58E-38 | 21.0729 | 0.907 | 0.697 | 2.06E-36 |
| CD1c | 0.000285 | 24.99935 | 0.85 | 0.823 | 0.012845 |

| | p_val | avg_log2FC | pct.1 | pct.2 | p_val_adj |
|---|---|---|---|---|---|
| CD14 | 1.96E-12 | -45.3674 | 0.895 | 0.921 | 8.82E-11 |
| CD163 | 2.67E-28 | -23.8461 | 0.962 | 0.976 | 1.20E-26 |
| CD62L | 7.99E-41 | -23.4155 | 0.975 | 0.932 | 3.60E-39 |
| CD123 | 6.40E-84 | -17.3376 | 0.924 | 0.987 | 2.88E-82 |
| CD33 | 5.85E-102 | -16.77 | 0.92 | 0.987 | 2.63E-100 |
| CD13 | 3.50E-09 | -16.6622 | 0.849 | 0.928 | 1.58E-07 |
| CD304 | 2.29E-12 | -16.6145 | 0.824 | 0.903 | 1.03E-10 |
| CD64 | 1.24E-97 | -16.2246 | 0.895 | 0.987 | 5.57E-96 |
| CD117 | 5.71E-19 | -15.8411 | 0.975 | 0.985 | 2.57E-17 |
| CD49d | 8.04E-19 | -13.1952 | 0.983 | 0.992 | 3.62E-17 |
| CD11b | 5.09E-75 | -10.7258 | 0.92 | 0.984 | 2.29E-73 |
| CD138 | 6.94E-08 | -3.76903 | 0.933 | 0.965 | 3.12E-06 |
| CD34 | 1.94E-10 | -2.62811 | 0.912 | 0.947 | 8.75E-09 |
| CD38 | 9.32E-26 | -1.35313 | 0.979 | 0.99 | 4.20E-24 |
| CD11c | 1.75E-101 | -1.17719 | 0.937 | 0.988 | 7.87E-100 |
| CD45 | 5.21E-18 | 2.522263 | 0.983 | 0.99 | 2.34E-16 |
| CD25 | 0.000417 | 4.181922 | 0.882 | 0.838 | 0.018764 |
| HLA.DR | 1.79E-55 | 4.210674 | 0.958 | 0.988 | 8.06E-54 |
| CD141 | 5.35E-09 | 4.890293 | 0.996 | 0.993 | 2.41E-07 |
| CD45RA | 1.35E-06 | 6.097652 | 0.979 | 0.987 | 6.06E-05 |
| CD71 | 3.75E-10 | 7.995727 | 0.992 | 0.991 | 1.69E-08 |
| CD4 | 0.000106 | 8.789058 | 0.975 | 0.992 | 0.004781 |
| CD7 | 1.02E-69 | 14.70695 | 0.987 | 0.944 | 4.58E-68 |
| CD8 | 4.35E-12 | 18.87003 | 0.937 | 0.903 | 1.96E-10 |
| CD5 | 1.69E-65 | 25.05066 | 0.979 | 0.878 | 7.58E-64 |
| CD2 | 2.53E-70 | 27.12998 | 0.962 | 0.835 | 1.14E-68 |
| CD3 | 2.15E-57 | 27.5123 | 0.979 | 0.887 | 9.68E-56 |
| CD19 | 5.56E-12 | 45.73689 | 0.903 | 0.812 | 2.50E-10 |
| CD1c | 1.03E-05 | 70.487 | 0.84 | 0.797 | 0.000463 |

| | p_val | avg_log2FC | pct.1 | pct.2 | p_val_adj |
|---|---|---|---|---|---|
| CD16 | 0.000107 | -46.4608 | 0.939 | 0.895 | 0.004802 |
| CD64 | 7.16E-11 | -38.0037 | 0.843 | 0.922 | 3.22E-09 |
| CD13 | 3.70E-55 | -14.2467 | 0.881 | 0.975 | 1.67E-53 |
| CD90 | 7.53E-06 | -14.1632 | 0.897 | 0.861 | 0.000339 |
| CD304 | 5.41E-12 | -11.8388 | 0.847 | 0.947 | 2.43E-10 |
| CD62P | 3.60E-24 | -10.8429 | 0.874 | 0.943 | 1.62E-22 |
| CD33 | 1.81E-70 | -10.2449 | 0.912 | 0.983 | 8.14E-69 |
| CD71 | 9.13E-50 | -9.26533 | 0.992 | 0.989 | 4.11E-48 |
| CD123 | 1.79E-57 | -7.64235 | 0.943 | 0.979 | 8.06E-56 |
| CD11c | 2.82E-59 | -7.34228 | 0.87 | 0.975 | 1.27E-57 |
| CD141 | 2.63E-33 | -5.93312 | 0.992 | 0.987 | 1.18E-31 |
| CD117 | 1.59E-54 | -3.94732 | 0.969 | 0.977 | 7.14E-53 |
| CD69 | 8.14E-11 | -0.44488 | 1 | 0.985 | 3.66E-09 |
| CD38 | 5.02E-51 | 2.401166 | 0.985 | 0.992 | 2.26E-49 |
| CD44 | 0.000724 | 4.093502 | 1 | 0.981 | 0.032577 |
| CD45RA | 4.01E-35 | 4.459874 | 1 | 0.981 | 1.80E-33 |
| HLA.DR | 4.03E-17 | 5.11003 | 0.908 | 0.966 | 1.81E-15 |
| CD45 | 3.49E-31 | 6.617422 | 1 | 0.985 | 1.57E-29 |
| CD22 | 0.000172 | 7.293446 | 0.889 | 0.882 | 0.007744 |
| CD25 | 0.000462 | 9.504228 | 0.935 | 0.899 | 0.020811 |
| CD7 | 1.87E-34 | 10.44943 | 0.977 | 0.949 | 8.43E-33 |
| CD19 | 3.25E-09 | 10.75365 | 0.916 | 0.869 | 1.46E-07 |
| CD3 | 5.37E-52 | 12.51682 | 0.95 | 0.827 | 2.41E-50 |
| CD4 | 8.69E-20 | 14.58687 | 0.973 | 0.949 | 3.91E-18 |
| CD5 | 4.39E-58 | 21.5091 | 0.973 | 0.823 | 1.97E-56 |
| CD2 | 5.34E-52 | 24.9829 | 0.973 | 0.857 | 2.40E-50 |
| CD62L | 1.19E-15 | 33.33802 | 0.966 | 0.932 | 5.38E-14 |
| CD8 | 8.27E-08 | 47.88799 | 0.897 | 0.84 | 3.72E-06 |
| CD1c | 5.61E-06 | 70.14279 | 0.854 | 0.827 | 0.000252 |

| | p_val | avg_log2FC | pct.1 | pct.2 | p_val_adj |
|---|---|---|---|---|---|
| CD123 | 2.87E-09 | -27.0937 | 0.854 | 0.905 | 1.29E-07 |
| CD38 | 2.58E-12 | -17.3621 | 0.897 | 0.941 | 1.16E-10 |
| CD33 | 1.87E-22 | -16.5041 | 0.851 | 0.935 | 8.42E-21 |
| CD117 | 1.80E-39 | -6.70166 | 0.9 | 0.98 | 8.12E-38 |
| CD163 | 4.45E-05 | -6.69019 | 0.793 | 0.756 | 0.002003 |
| CD14 | 1.93E-05 | -4.45624 | 0.759 | 0.735 | 0.000868 |
| CD11b | 4.17E-05 | -2.33222 | 0.762 | 0.74 | 0.001874 |
| CD138 | 7.49E-05 | 1.252989 | 0.816 | 0.788 | 0.003369 |
| CD45RO | 1.44E-06 | 2.241111 | 0.739 | 0.709 | 6.50E-05 |
| CD45 | 8.75E-08 | 3.027076 | 0.87 | 0.83 | 3.94E-06 |
| CD1c | 4.85E-05 | 4.785255 | 0.766 | 0.739 | 0.002185 |
| CD62L | 1.43E-07 | 8.534984 | 0.789 | 0.745 | 6.43E-06 |
| CD7 | 6.92E-11 | 15.60117 | 0.881 | 0.829 | 3.12E-09 |
| CD8 | 7.06E-10 | 18.43127 | 0.797 | 0.745 | 3.18E-08 |
| CD4 | 8.08E-11 | 21.13213 | 0.847 | 0.801 | 3.64E-09 |
| CD19 | 3.86E-05 | 24.78492 | 0.766 | 0.745 | 0.001739 |
| CD2 | 7.26E-31 | 29.21209 | 0.87 | 0.742 | 3.27E-29 |
| CD90 | 1.04E-06 | 30.37105 | 0.789 | 0.758 | 4.69E-05 |
| CD16 | 3.39E-06 | 31.36489 | 0.774 | 0.741 | 0.000153 |
| CD3 | 1.70E-26 | 32.0263 | 0.87 | 0.743 | 7.66E-25 |
| CD5 | 8.86E-27 | 33.84855 | 0.858 | 0.732 | 3.99E-25 |
