## Extended Data for "*PTPN11* Mutation Clonal Hierarchy in Acute Myeloid Leukemia"

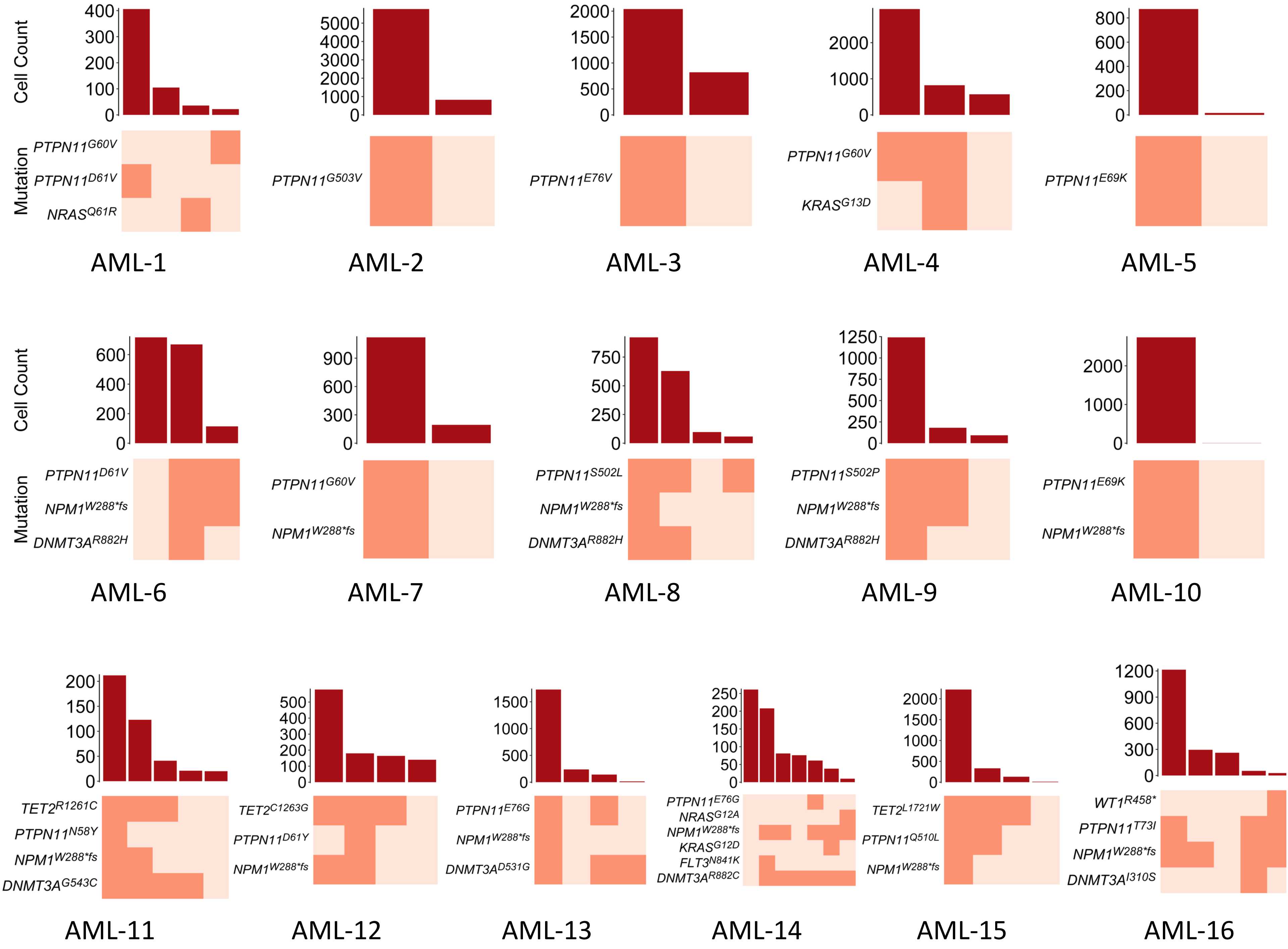

Cell Count
Mutation
AML-1
PTPN11G60V
PTPN11D61V
NRASQ61R
AML-2
PTPN11G503V
AML-3
PTPN11E76V
AML-4
PTPN11G60V
KRASG13D
AML-5
PTPN11E69K
AML-6
PTPN11D61V
NPM1W288*fs
DNMT3AR882H
AML-7
PTPN11G60V
NPM1W288*fs
AML-8
PTPN11S502L
NPM1W288*fs
DNMT3AR882H
AML-9
PTPN11S502P
NPM1W288*fs
DNMT3AR882H
AML-10
PTPN11E69K
NPM1W288*fs
AML-11
TET2R1261C
PTPN11N58Y
NPM1W288*fs
DNMT3AG543C
AML-12
TET2C1263G
PTPN11D61Y
NPM1W288*fs
AML-13
PTPN11E76G
NPM1W288*fs
DNMT3AD531G
AML-14
PTPN11E76G
NRASG12A
NPM1W288*fs
KRASG12D
FLT3N841K
DNMT3AR882C
AML-15
TET2L1721W
PTPN11Q510L
NPM1W288*fs
AML-16
WT1R458*
PTPN11T73I
NPM1W288*fs
DNMT3AI310S

#### Extended data figure 2a - CD11b$^{+}$/cKit$^{+}$ cells in the peripheral blood over time

| **Percent CD11b$^{+}$/cKit$^{+}$ cells** | | | | |
|---|---|---|---|---|
| **Week** | Mx1-Cre (n=7) | Npm1$^{cA}$ (n=6) | Ptpn11$^{E76K}$ (n=3) | Npm1$^{cA}$/Ptpn11$^{E76K}$ (n=12) |
| **4** | 0.186±0.176 | 0.095±0.048 | 0.187±0.161 | 2.586±4.060 |
| **8** | 0.091±0.053 | 0.113±0.135 | 0.443±0.190 | 3.436±1.910 |
| **12** | 0.404±0.441 | 0.164±0.124 | 0.395±0.460 | 6.270±3.758 |
| **16** | 0.170±0.189 | 0.158±0.102 | 1.450±0.000* | 12.427±5.892 |
| **CD11b$^{+}$/cKit$^{+}$ counts ($10^3$/µL)** | | | | |
| **Week** | Mx1-Cre (n=7) | Npm1$^{cA}$ (n=6) | Ptpn11$^{E76K}$ (n=3) | Npm1$^{cA}$/Ptpn11$^{E76K}$ (n=8) |
| **4** | 0.022±.023 | 0.010±.006 | 0.032±.026 | 1.397±2.618 |
| **8** | 0.008±.005 | 0.014±.020 | 0.077±.049 | 1.704±1.539 |
| **12** | 0.045±.062 | 0.013±.009 | 0.071±.076 | 4.735±4.030 |
| **16** | 0.017±.019 | 0.014±.008 | 0.183±0.000* | 33.207±28.283 |

* data reflects only 1 mouse

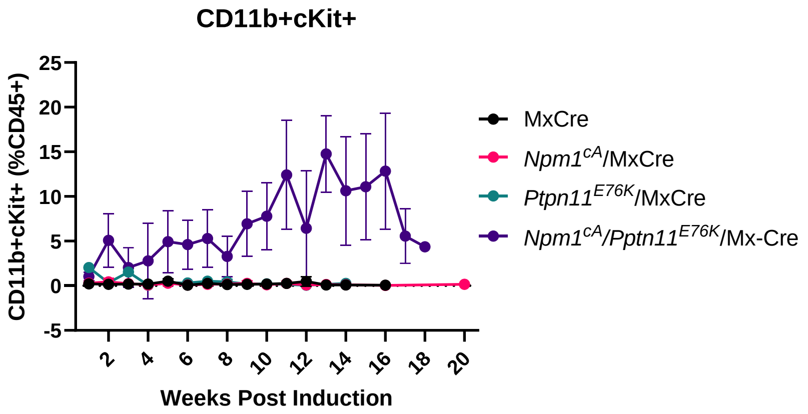


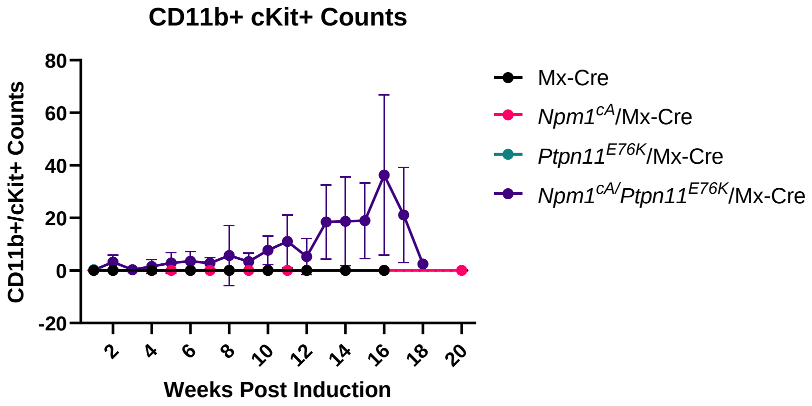

### Extended data figure 2b

S0140 (MxCre) 1 month post poly(I:C)

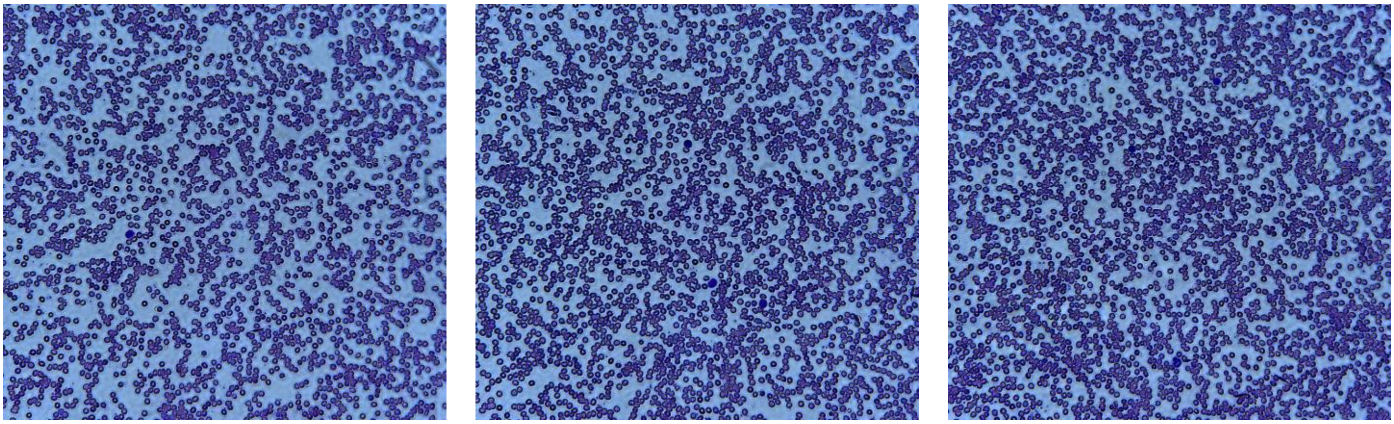

S0158 (*Ptpn11*$^{E57K}$/MxCre) 1 month post poly(I:C)

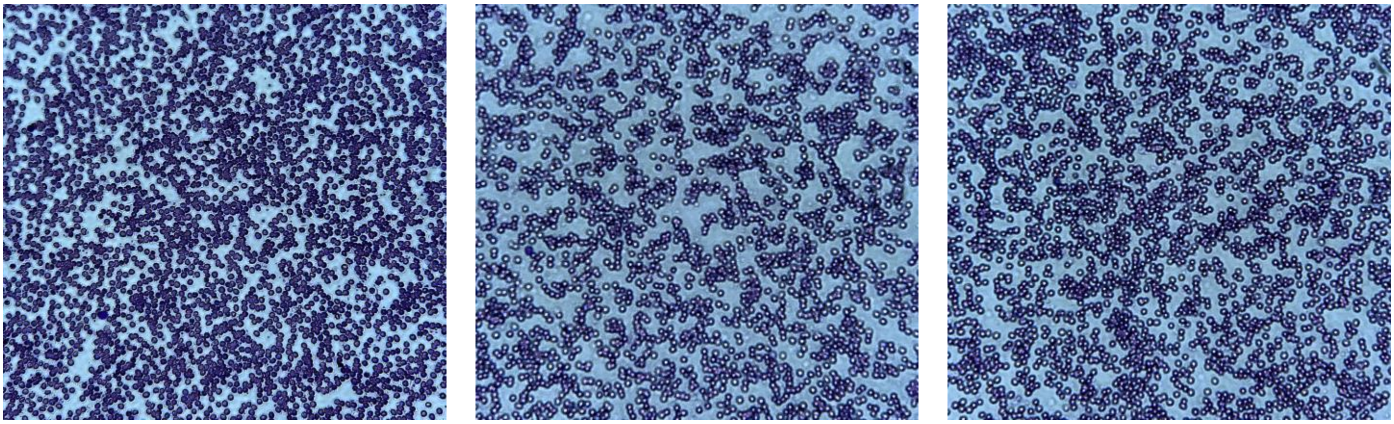

S0162 (*Ptpn11*$^{E57K}$/MxCre) 1 month post poly(I:C)

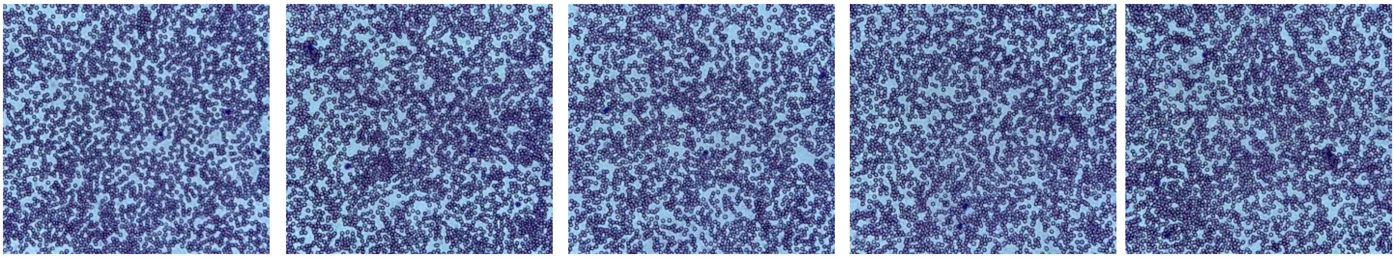

S0163 (*Ptpn11*$^{E57K}$/*Npm1*$^{cA}$/MxCre) 1 month post poly(I:C)

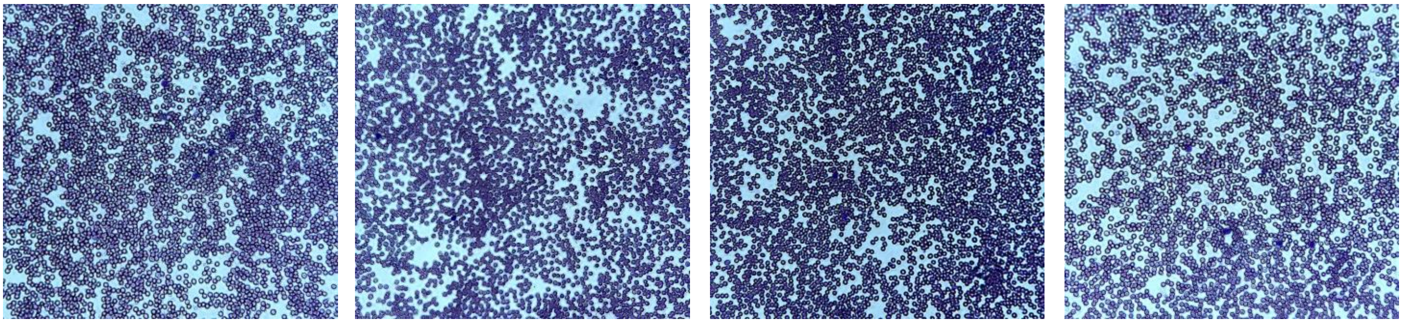

S0165 (*Ptpn11*$^{E57K}$/*Npm1*$^{cA}$/MxCre) 1 month post poly(I:C)

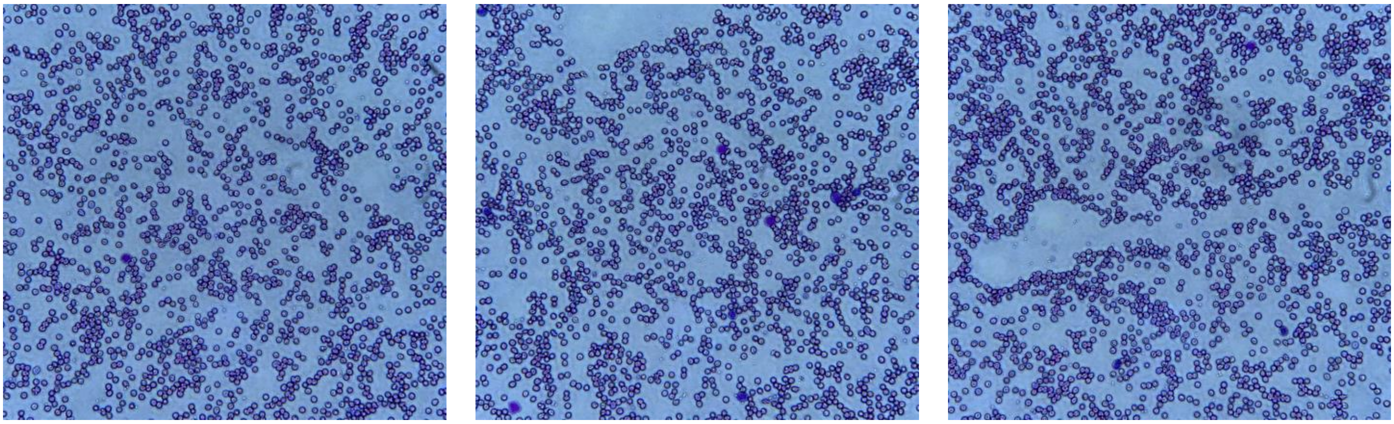

S0167 (*Ptpn11*$^{E57K}$/*Npm1*$^{cA}$/MxCre) 1 month post poly(I:C)

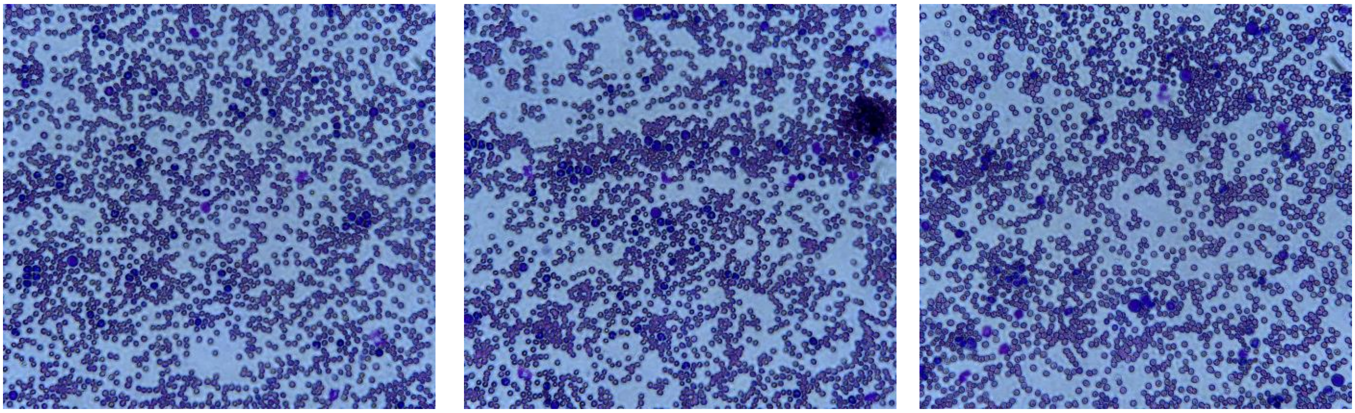

S0175 (*Ptpn11*$^{E57K}$/*Npm1*$^{cA}$/MxCre) 1 month post poly(I:C)

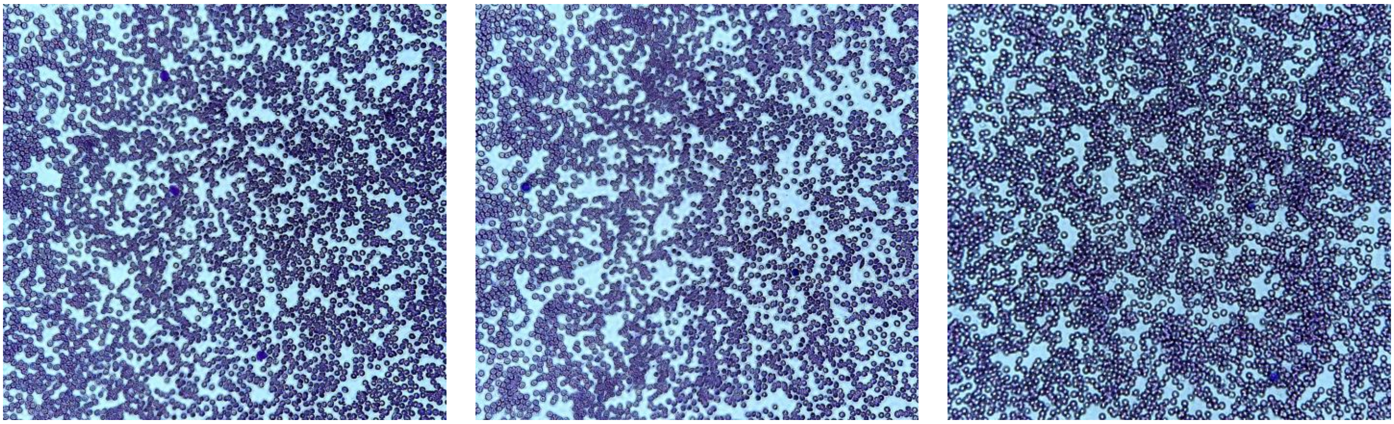

S0176 (*Ptpn11*$^{E57K}$/*Npm1*$^{cA}$/MxCre) 1 month post poly(I:C)

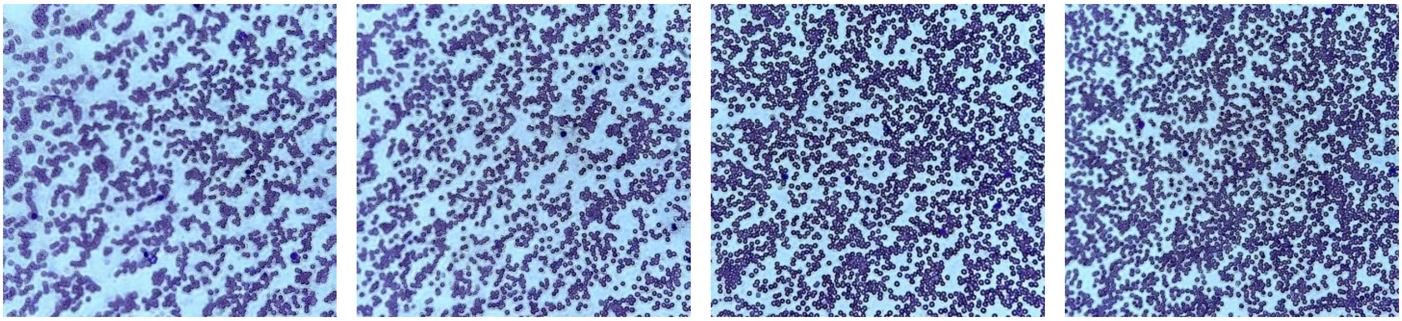

Extended data figure 2c

| Mouse ID | Sex | Genotype | age/cause of death | Neoplasia (Y or N) | Neoplasm Diagnosis | Spleen | Lymph Node | Liver | Lung | Kidney | GI | Bone marrow | Brain | Mesentery | Other changes |
|---|---|---|---|---|---|---|---|---|---|---|---|---|---|---|---|
| S0137/ S0167/ S0173 | F | *Ptpn11*$^{E76K}$/ *Npm1*$^{cA}$/ MxCre | 4 months, died | Y | myeloid malignancy | NA | Y | Y | N | Y, mild | N | Y | Y | N | markedly enlarged lymph nodes, some mild differentiation to bands, no obvious lymphoid cells, brain AML mostly within meninges, a bit autolyzed |
| S0168/ S0176 | F | *Ptpn11*$^{E76K}$/ *Npm1*$^{cA}$/ MxCre | 3 months, died | Y | myeloid malignancy | Y | Y | Y, subcapsular hemorrhage and necrosis | N | Y, mild | Y | Y | N | Y | Markedly enlarged lymph nodes, somewhat less differentiated than 173, small foci on endocardium |
| S0135/ S0165 | F | *Ptpn11*$^{E76K}$/ *Npm1*$^{cA}$/ MxCre | 5 months, died | Y | myeloid malignancy | Y | Y | Y | Y, numerous small foci in all sections | NA | N | Y | N | y | Disease also within the myocardium, within meninges |
| S0140 | F | MxCre | 5 months, control | N | N/A | N | N | N | N | N | N | N | N | N | Moderate, variably mature myeloid expansion within liver, lymph nodes, adrenal glands, spleen, bone marrow, and mesenteric fat. |
| S0037 | F | MxCre | 11.7 months, littermate control | N | N/A | N | N | N | N | N | N | N | N | N | Lymphoid and myeloid hyperplasia in lymph nodes and bone marrow. No evidence of neoplasm. extramedullary hematopoeisis in kindey and liver. BALT hyperplasia in lung. To confirm that this lymphoid hyperplasia is non-neoplastic, IHC would be recommended. May indicate underlying inflammatory process of uknown origin/etiology. One testis has a focus of fibrosis which does not appear to be neoplastic. |
| S0038 | F | *Ptpn11*$^{E76K}$/ MxCre | 11.7 months, died | Y | Myeloid malignancy | NA | large lymph node full of large, blast-like round cells, some of which are moderately differentiated with band morphology | numerous round, blast-like cells in sinusoids and surrounding blood vessels | same population surrounding vessels throughout the lung | very small foci of increased cellularity | N | yes, numerous large round blast cells with many mitotic figures which are often bizarre, lack of normal hematopoetic tissue | N | N | hemorrhage in spinal cord and neoplastic cells extend into spinal canal, consistent with history of hind limb paralysis. Myeloid malignancy, least differentiated of all mice. |

| Mouse ID | Sex | Genotype | age/cause of death | Neoplasia (Y or N) | Neoplasm Diagnosis | Spleen | Lymph Node | Liver | Lung | Kidney | GI | Bone marrow | Brain | Mesentery | Other changes |
|---|---|---|---|---|---|---|---|---|---|---|---|---|---|---|---|
| S0160/ S0175 | M | *Ptpn11*$^{E76K}$/ *Npm1*$^{cA}$/ MxCre | 6 months, died | Y | myeloid malignancy | very large, neoplastic myeloid cells | yes, myeloblastic cells enlarge lymph node, no apparent germinal centers | Y, very large, neoplastic myeloid cells within sinusoids and surrounding vessels | accumulations of neoplastic cells around vessels | Y, small foci of same neoplastic population throughout interstitium | lamina propria has increased cellularity | yes, large myeloblastic neoplastic cells, lack of normal hematopoietic tissue | Y, restricted to meninges | Y, scattered throughout mesentery, no obvious masses | Myeloid malignancy throughout most organs |
| S0189/ S0199 | M | MxCre | 5.4 months, littermate control | N | N/A | N | N | N | N | N | N | N | N | N | No evidence of neoplasia nor toxicity in any organs observed. Mild extramedullary hematopoesis in liver, normal M:E ratio in all bone marrow compartments with orderly maturation of lymphoid, myeloid, and erythroid lines. |
| S0194/ S0200 | F | *Ptpn11*$^{E76K}$/ *Npm1*$^{cA}$/ MxCre | 5 months, died | Y | myeloid malignancy | y, myeloblastic cells efface red and white pulp, no apparent germinal centers, lots of very large cells with bizarre mitotic figures, lots of MNGCs | y, myeloblastic cells enlarge lymph node, no apparent germinal centers, lots of very large cells with bizarre mitotic figures, lots of MNGCs | Y, very large, neoplastic myeloid cells within sinusoids and surrounding vessels | Y, neoplastic cells around vessles, in alveolar walls. one lobe has coagula of neuts, pyknotic debris, hemorrhage, fibrin in bronchus | Y, small foci of same neoplastic population throughout interstitium | N | Y, lack of normal hematopoeitic tissue | Y | Y | Neoplasm exends into meninges and also focally expands and disrupts the cornea |

### Extended figure 3A

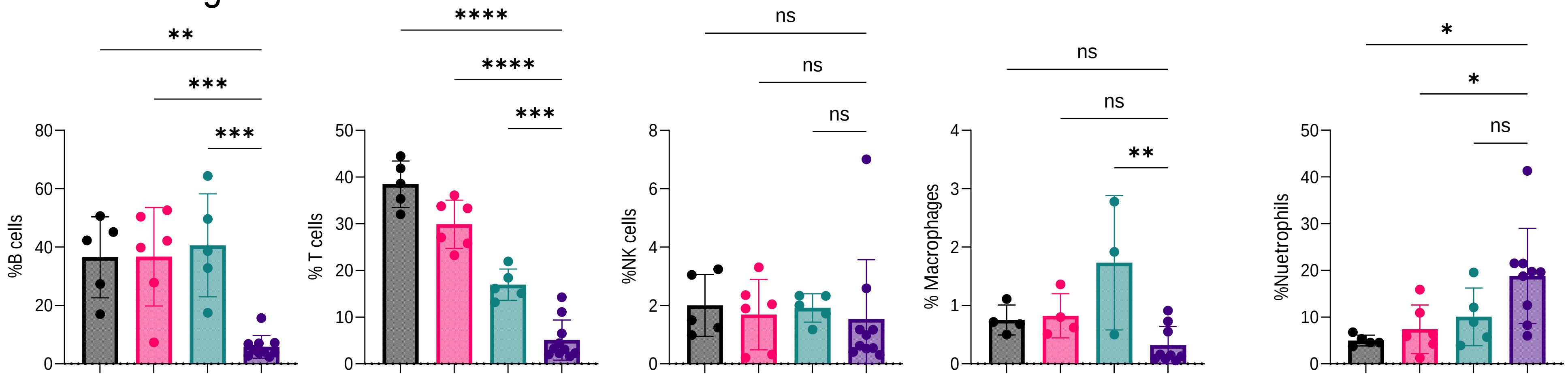


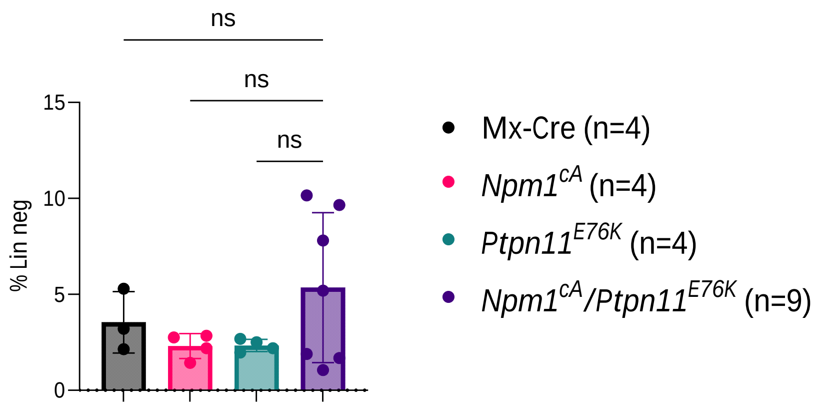


B

| | Percent pDCs | | | |
|---|---|---|---|---|
| **Week** | *Mx1-Cre* (n=7) | *$Npm1^{cA}$* (n=5) | *$Ptpn11^{E76K}$* (n=3) | *$Npm1^{cA}$/$Ptpn11^{E76K}$* (n=12) |
| **4** | .031±.027 | .020±.019 | .140±.154 | .155±.134 |
| **8** | .016±.021 | .010±.010 | .040±.028 | .460±.474 |
| **12** | .024±.017 | .032±.026 | .020±0 | 1.032±.706 |
| **16** | .004±.005 | 0±0 | -* | 3.236±1.831 |
| | **pDCs counts ($10^3$/µL)** | | | |
| **Week** | *Mx1-Cre* (n=7) | *$Npm1^{cA}$* (n=5) | *$Ptpn11^{E76K}$* (n=3) | *$Npm1^{cA}$/$Ptpn11^{E76K}$* (n=12) |
| **4** | .004±.004 | .002±.002 | .023±.026 | .052±.073 |
| **8** | .001±.002 | .001±.001 | .007±.007 | .190±.206 |
| **12** | .003±.002 | .002±.002 | .005±0 | .712±.556 |
| **16** | .000±.001 | .000±.000 | | 9.145±7.516 |

C

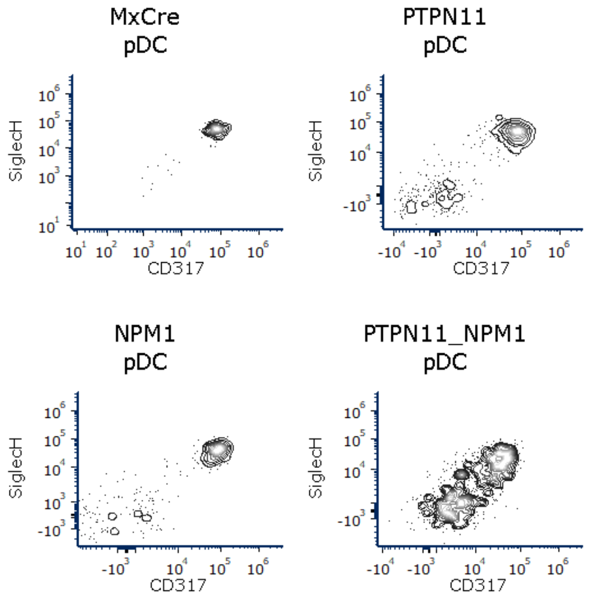


D

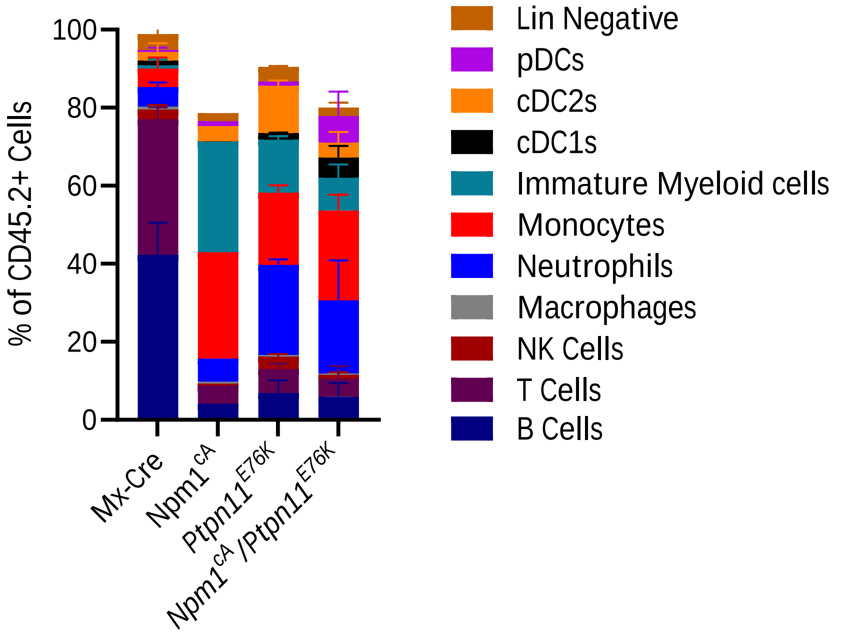

Extended data figure 4a

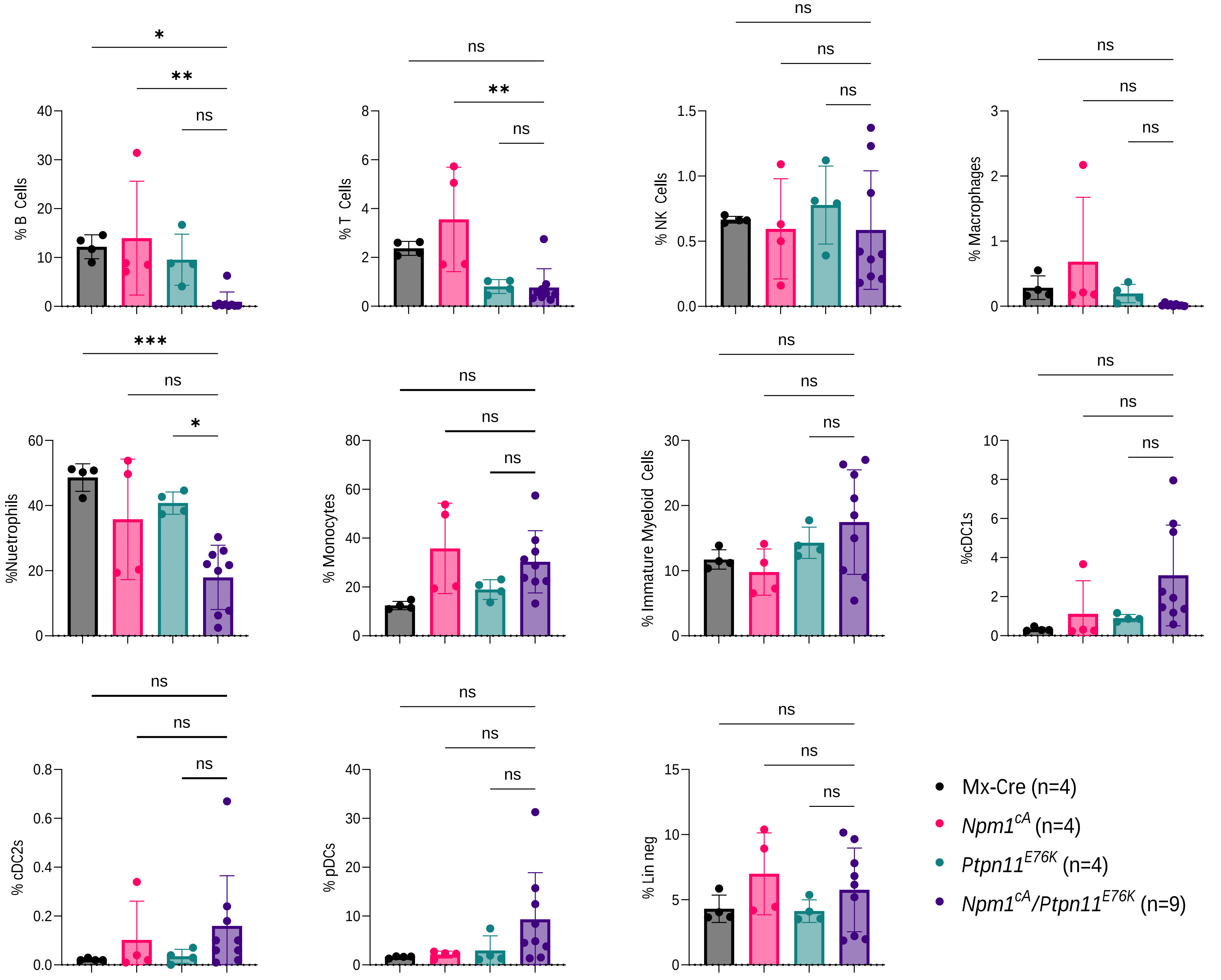

Extended data figure 4b

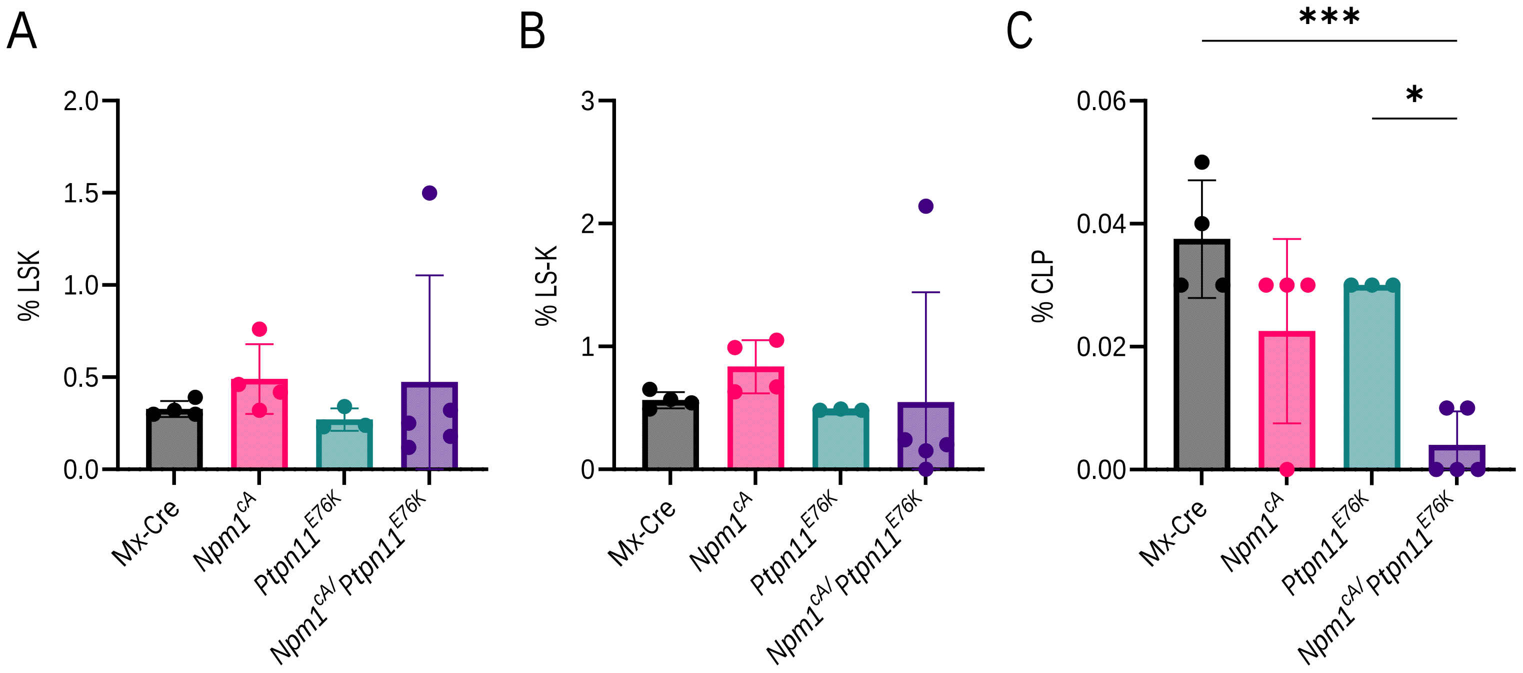

### Extended data Figure 6A

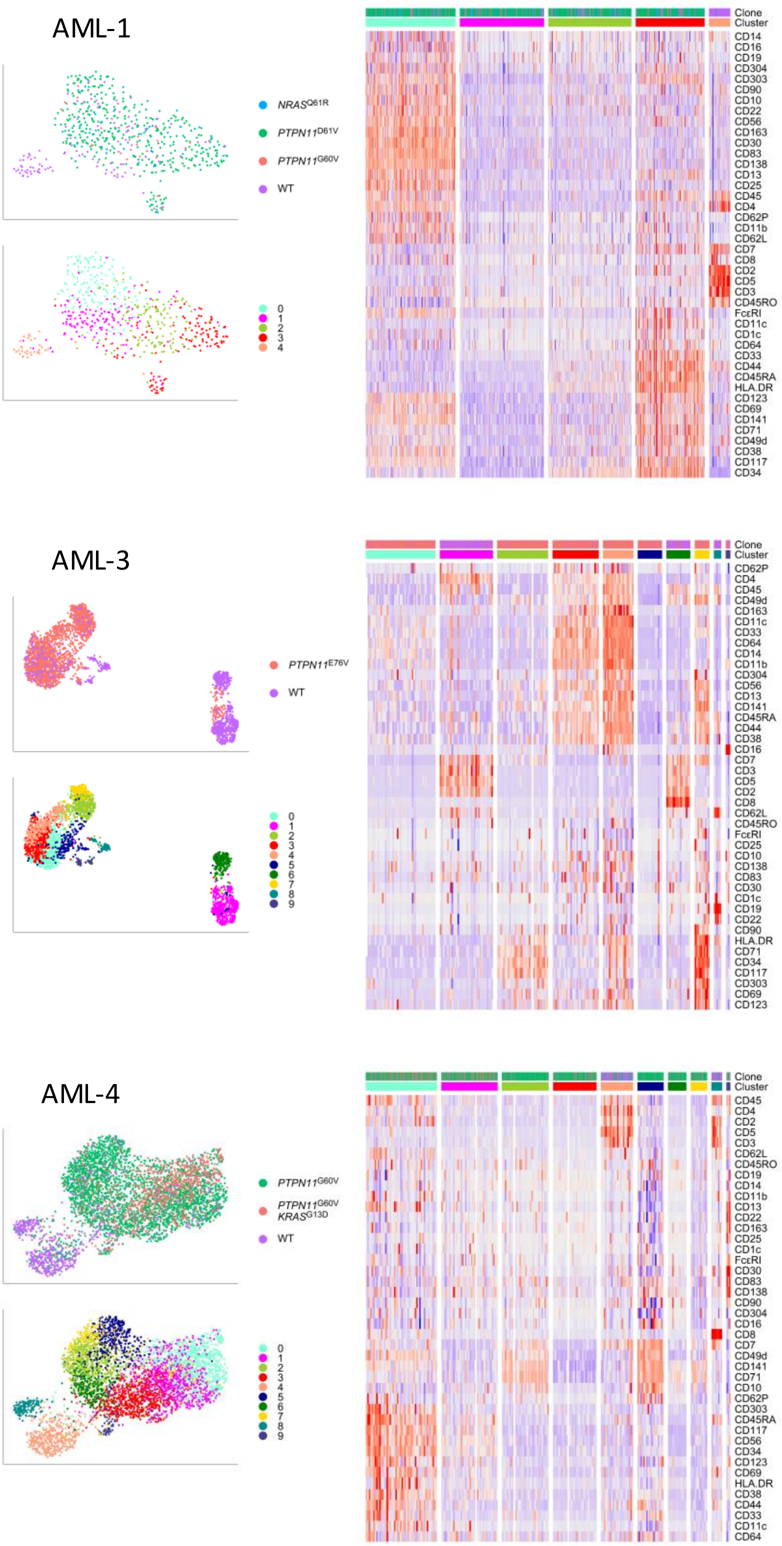

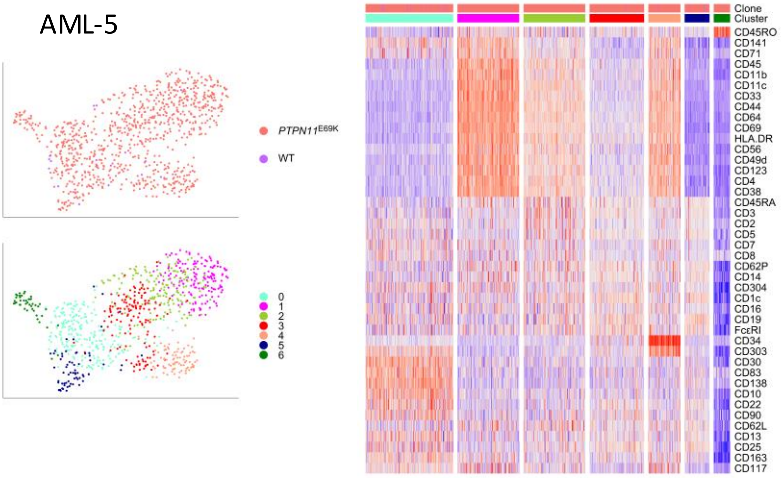


### Extended data Figure 6B

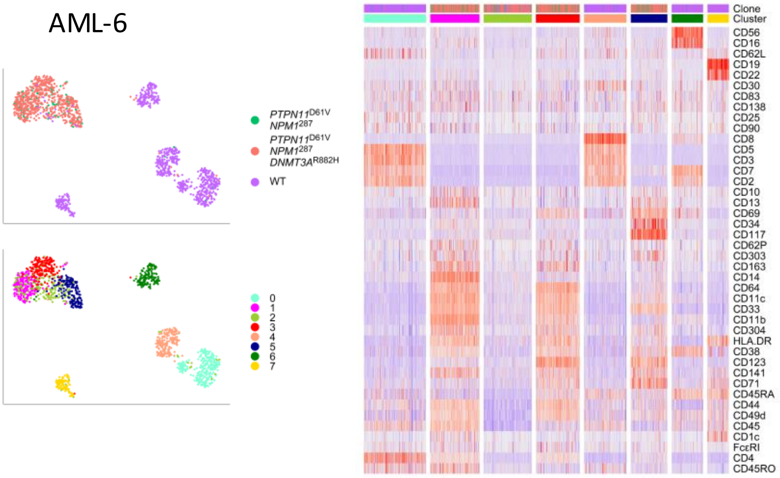


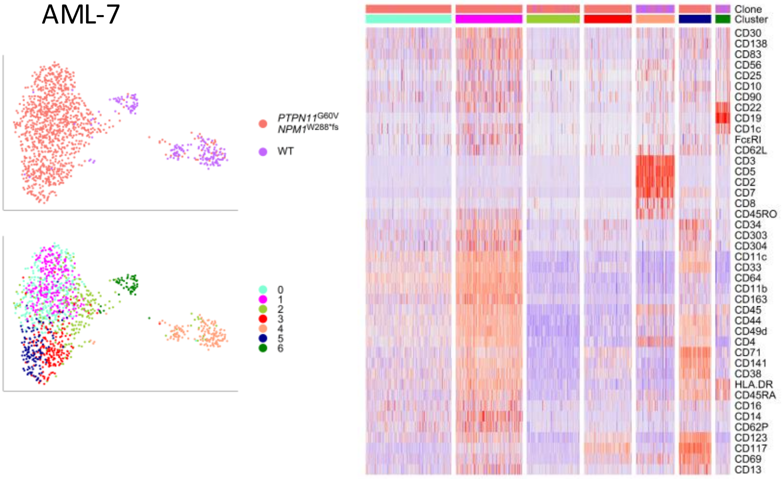

AML-8

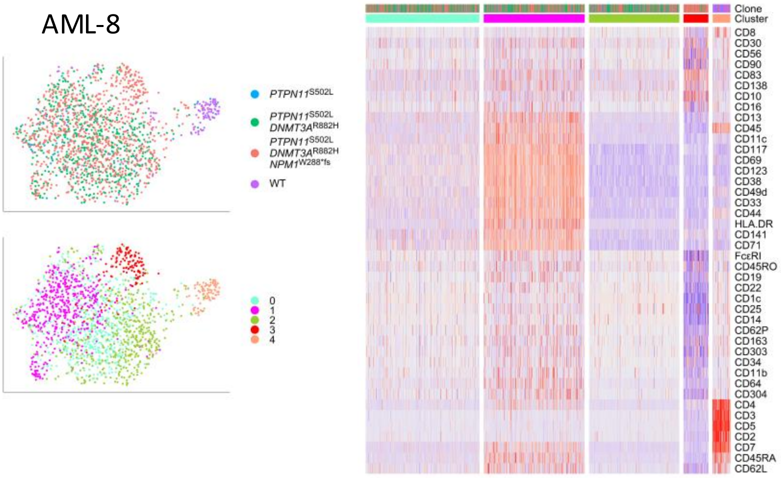


AML-9

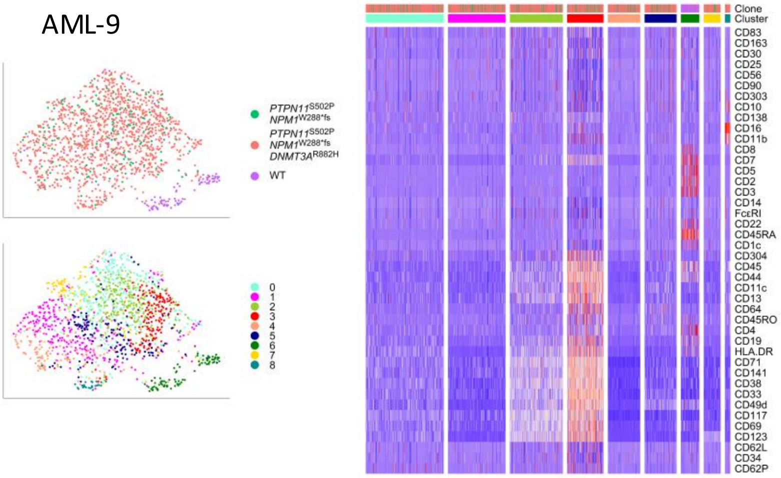


AML-10

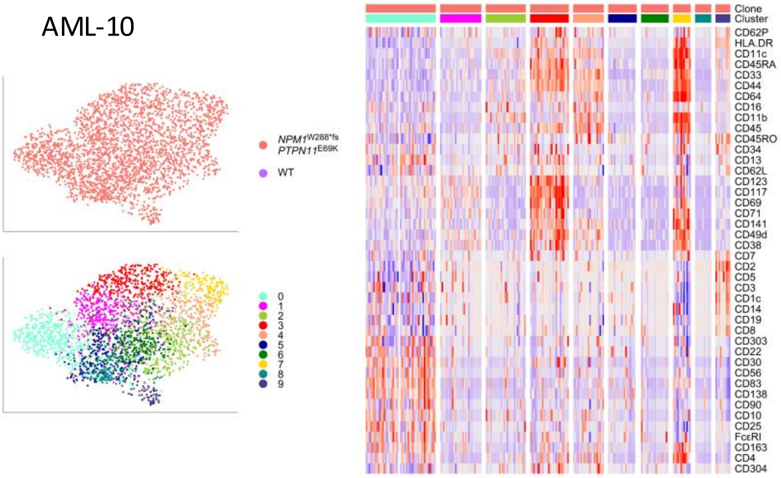

### Extended data Figure 6C

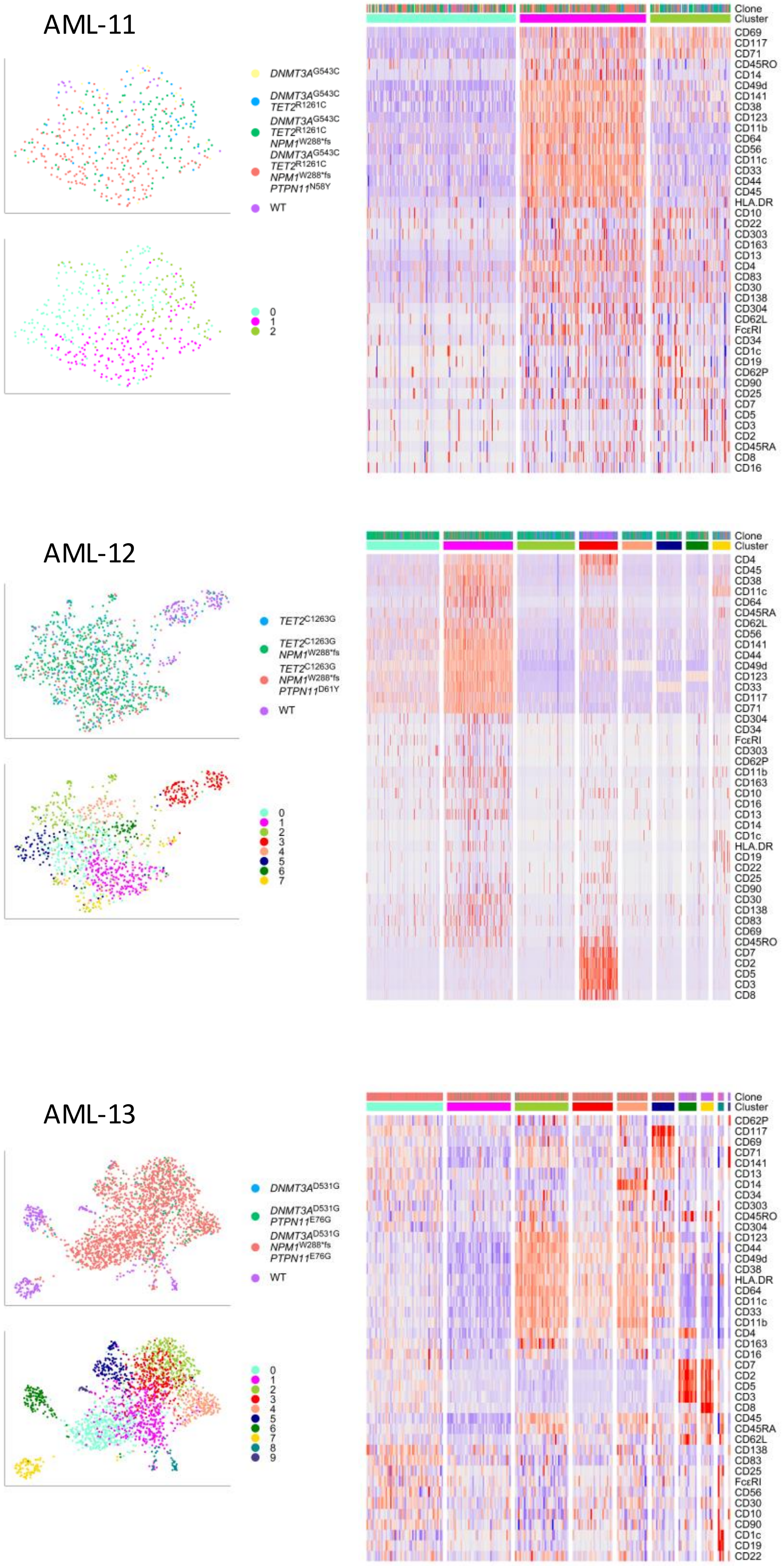

AML-14

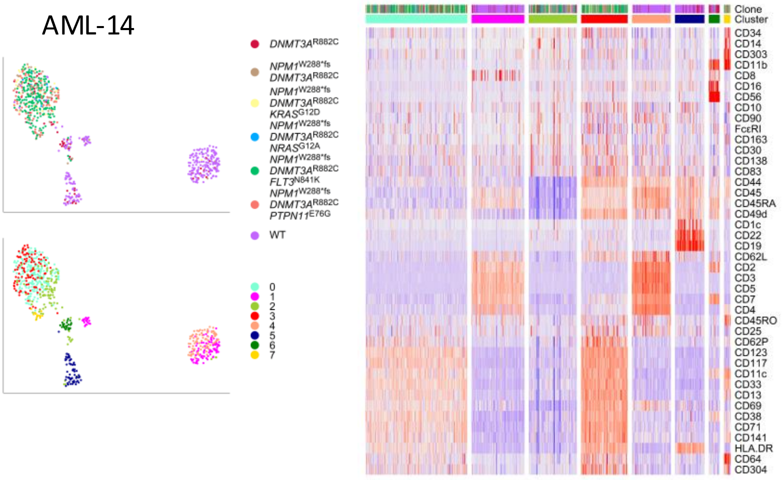


AML-15

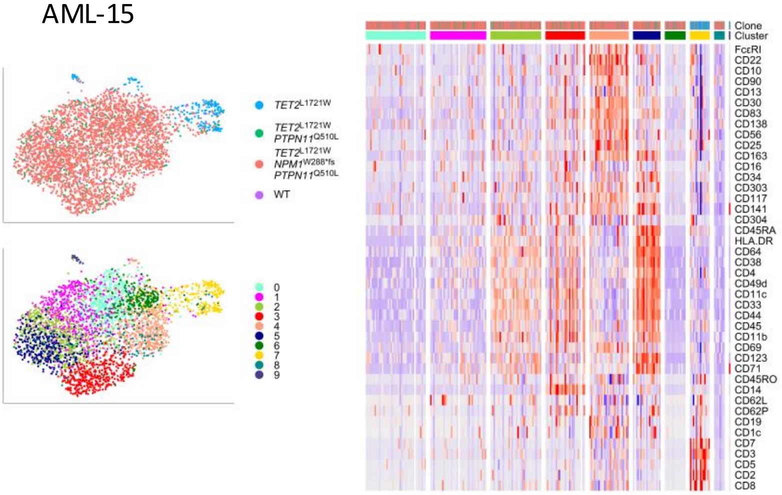


AML-16

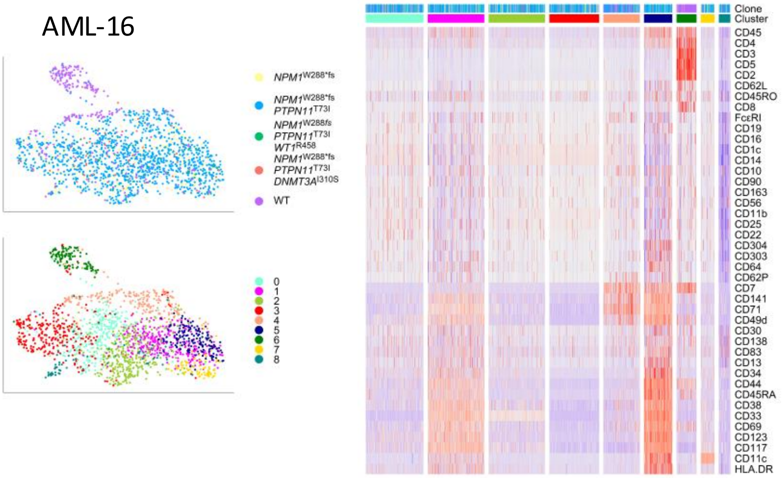
